# Nitrogen palaeo-isoscapes: Changing spatial gradients of faunal δ^15^N in late Pleistocene and early Holocene Europe

**DOI:** 10.1101/2022.05.05.490797

**Authors:** Hazel Reade, Jennifer A. Tripp, Delphine Frémondeau, Kerry L. Sayle, Thomas F.G. Higham, Martin Street, Rhiannon E. Stevens

**Author notes:** Corresponding author (HR). Department of Chemistry, University of San Francisco, San Francisco USA. Department of Evolutionary Anthropology, Faculty of Life Sciences, University of Vienna, Vienna, Austria (Thomas F.G. Higham). Human Evolution and Archaeological Sciences (HEAS), University of Vienna, A-1030, Vienna, Austria.

## Abstract

Nitrogen isotope (δ^15^N) analysis of animal tissue is widely used in archaeology and palaeoecology to investigate diet and ecological niche. Data interpretations require an understanding of nitrogen isotope compositions at the base of the food web (baseline δ^15^N). Significant variation in animal δ^15^N has been recognised at various spatiotemporal scales and linked to changes both in baseline δ^15^N and animal ecology. Isoscapes (models of isotope spatial variation) have proved a useful tool for investigating spatial variability in biogeochemical cycles in present-day marine and terrestrial ecosystems, but so far, their application to palaeo-data has been limited. Here, we present time-sliced nitrogen isoscapes for late Pleistocene and early Holocene Europe (c. 50,000 to 10,000 years BP) using herbivore collagen δ^15^N data. This period covers the Last Glacial-Interglacial Transition, during which significant variation in the terrestrial nitrogen cycle occurred. Our results show clear changes in spatial gradients of δ^15^N through time. Prediction of the lowest faunal δ^15^N values in northern latitudes after, rather than during, the Last Glacial Maximum is consistent with the Late Glacial Nitrogen Excursion (LGNE). We consider the potential of incorporating climatic covariate data into isoscape models but find their inclusion does not improve model performance. These findings have implications for investigating the drivers of the LGNE, which has been linked to increased landscape moisture and permafrost thaw, and for understanding changing isotopic baselines, which are fundamental for studies investigating diets, niche partitioning, and migration of higher trophic level animals.

## 1. Introduction

Nitrogen isotope ratio (^15^N/^14^N, expressed as δ^15^N) of biological tissue is frequently used in archaeology and palaeoecology to investigate dietary behaviours, ecological niche, and past food webs (1–3). Specifically, δ^15^N is used to infer information about trophic structures. Obtaining reliable estimations of faunal trophic position requires understanding of the isotope compositions at the base of the food web. In other words, knowledge of the plant and soil δ^15^N values upon which the fauna lived and fed (hereafter termed baseline δ^15^N). However, this information is not usually readily obtainable from archaeological or palaeontological contexts, where the preservation of plant and/or soil material suitable for analysis can be limited. Moreover, plant and soil δ^15^N is highly heterogeneous and is not static in space or time, complicating inferences of baseline δ^15^N available to fauna.

Many interconnected factors exert influence on plant and soil δ^15^N and nitrogen cycling in the terrestrial environment (4). These relate to climate, plant functional type, mycorrhizal associations, soil characteristics, and the availability of different forms of nitrogen (5–8). On global and continental scales strong, albeit indirect, relationships exist between plant δ^15^N and temperature and precipitation (6, 7). These relationships are also expressed over smaller spatial scales with strong altitudinal gradients (9, 10). Likewise, such spatial relationships are also represented in faunal δ^15^N values (9,11,12). However, differences in dietary and mobility behaviours between different species, populations, and individuals introduce additional variation into the faunal δ^15^N signal (1, 3). Indeed, while δ^15^N analysis of biological tissues is frequently used in archaeology and palaeoecology to investigate dietary behaviours and ecological niche, our ability to decipher environmental influence from feeding behaviour remains an ongoing challenge.

On long timescales (10^3^ to 10^5^ years) significant temporal variation has been identified in herbivore δ^15^N (13–25). This variation has been interpreted as representing changes to baseline δ^15^N in response to climatic and environmental drivers. Most notably, a large decrease and then rapid increase in herbivore δ^15^N occurred during the Late Glacial, between approximately 17,000- and 12,000-years before present (BP) (15–17,20–23,25,26). This trend occurs in multiple species, across a wide range of mid and high latitude environments and in recent years has been termed the Late Glacial Nitrogen Excursion (LGNE) (25). As the body of late Pleistocene herbivore δ^15^N data has grown, spatial and temporal asynchronicities in the LGNE are becoming increasingly apparent (3, 25). Similarly, significant differences in species-specific δ^15^N variation are also recognised (1, 3).

Through this increasing body of data, significant new opportunities to investigate spatiotemporal patterns in herbivore δ^15^N are emerging. Isoscape approaches (modelling of isotope spatial variation) have proved useful tools for investigating isotopic spatial variability in present-day marine and terrestrial ecosystems but are yet to be widely applied to palaeo-focused research. Here, we create time-sliced isoscape prediction maps of herbivore collagen δ^15^N through the late Pleistocene and early Holocene periods in Europe. Time-sliced spatial interpolation offers the potential to assess changing spatial gradients of δ^15^N through time. Combining this analysis with high resolution climate model data (27) opens up a significant new avenue of research through which the potential drivers of the LGNE can be investigated. Improved characterisation of spatiotemporal trends in herbivore δ^15^N may also ultimately contribute to more robust trophic structure analysis of archaeological and palaeontological materials. This is particularly important as many palaeo-focued studies use herbivore δ^15^N, in the absence of suitable plant samples, to infer baseline δ^15^N values for terrestrial food web analysis and in the interpretation of data from higher trophic level animals in relation to mobility, migration, and dietary research.

## 2 Materials and Methods

### 2.1 Data compilation

Newly generated and previously published herbivore collagen δ^15^N from late Pleistocene and early Holocene European contexts were compiled for latitudes between 35°N and 60°N and longitudes between 10°W and 30°E. Temporal scope was restricted to before the 8.2 ka BP climatic event (28), to avoid capturing human-influence on baseline δ^15^N that occurred through agricultural developments with the onset of the Neolithic (29, 30), and after 55 ka BP, which is the current approximate limit of radiocarbon dating and calibration (31). Data come from both archaeological and palaeontological assemblages. We believe the resultant compilation captures the majority of available δ^15^N data from the time period and geographical region, enabling major spatial and temporal trends in δ^15^N to be evaluated.

We did not include δ^15^N data which had a C/N atomic ratio <2.9 or >3.6, or where their publication indicated the result was unreliable. We also did not include mammoth, which have been shown to display anomalously high δ^15^N values when compared to other contemporaneous herbivore species (32–34), or smaller herbivores, such as those in the order *Rodentia*, which are underrepresented in the available data and are sensitive to capturing small-scale heterogeneities in baseline δ^15^N that act independently of large-scale climatic influences due to small home range size (35–37). Any published data where taxonomic identification was uncertain was not included. All antler and tooth samples from Cervid species (*Alces alces, Capreolus capreolus, Cervus elaphus, Dama dama, Megaloceros giganticus, Rangifer tarandus,* and *Rupicapra rupicapra*) were avoided to limit seasonal biasing in the data (38–41). Finally, any data which were identified as duplicate analyses on the same sample/individual animal were omitted.

Compiled data were divided into 7 temporal bins (Table 1); to minimise the potential of averaging data across different climatic states/environmental conditions, whilst not overly limiting the number of data included in each time bin, we base our time bins on known major climatic events (42, 43). We recognise that further climate events occurred within our selected bins, and that their expression is asynchronous across the region of study, but without a greater sample size and/or improvements in the accuracy to which sample age can be estimated, analysis at greater temporal resolution is not possible. In particular, the earliest of our two bins (early Oxygen Isotope Stage 3 (EOIS3) and late Oxygen Isotope Stage 3 (LOIS3)) capture data from multiple different climate states. For directly radiocarbon dated samples, calibration was performed using OxCal (version 4.4) (44) and the IntCal20 calibration curve (31). Dates were binned based on the median of the 95.4% probability calibrated age range. For samples where age is based on stratigraphic provenance (context dated samples), time bin assignment was based on the age of the assemblage given in the original publication of the data, or most recent age model for the site in cases where the chronological position of an assemblage had been subsequently revised. For data where a secure age assignment could not be made, or when an age assignment spanned the boundary between two temporal bins, the data were excluded. The resultant dataset contained 2,718 δ^15^N values, as reported in the Supporting Information (S1 Dataset). In total the data include 479 new δ^15^N values and 197 new radiocarbon dates. Methods of sample preparation and analysis for the newly generated isotope data and radiocarbon determinations are provided Supporting Information (Section 1 in S3 Supporting Information).

**Table 1.** Time bins and climate model time steps used in this study.

| Time bin (abbreviation) | Upper limit<br>(year cal BP) | Lower limit<br>(year cal BP) | Climate model time<br>step |
| --- | --- | --- | --- |
| Early Holocene (EH) | 11,650 | 8,190 | 11,000 |
| Younger Dryas (YD) | 12,850 | 11,650 | 12,000 |
| Late Glacial Interstadial (LGI) | 14,650 | 12,850 | 14,000 |
| Last Glacial Termination (LGT) | 19,500 | 14,650 | 15,000 |
| Last Glacial Maximum (LGM) | 27,500 | 19,500 | 24,000 |
| Late Oxygen Isotope Stage 3 (LOIS3) | 39,850 | 27,500 | 36,000 |
| Early Oxygen Isotope Stage 3 (EOIS3) | 55,000 | 39,850 | 42,000 |

Elevation and bioclimatic covariate data were assembled for each δ^15^N sample, based on its geographic origin and time bin assignment. Elevation data was extracted from the Global Multi-Resolution Terrain Elevation Data 2010 (GMTED2010) model (45). As this elevation data is relative to present day sea level, it does not account for temporal changes in sea level or isostatic changes related to the growth and melting of ice sheets. Bioclimatic data was extracted from 0.5° resolution, biased-corrected combined HadCM3 and HadAM3H time series climate simulations (27). Data for these variables is available at a temporal resolution of 1,000-years for 21,000 years BP to present and at 2,000-year intervals prior 21,000 years BP. The distribution of samples within each time bin was evaluated using 1000-year bins and the modelled time step most closely corresponding to the greatest prevalence of data within the upper and lower limit of the event boundary was selected (Table 1).

### 2.2 Data exploration and analysis

All statistical analyses were performed using the R programming language (version 4.0.4) (46); the R script is provided in the Supporting Information (S2 Script). The compiled data were first evaluated for potential species-based effects on δ^15^N related to diet, habitat preference, and ecology. While differences were identified between species, these were unsystematic, varying by location and time period (full analysis is reported in Section 2 in S3 Supporting Information). As such, no species-based data normalisation procedures were applied prior to geostatistical analysis, although we also consider scope for species-specific analysis where sample size permits in Discussion section 4.2.

Spatial structures in the data were evaluated by time bin using Anselin’s Local Moran’s I and Global Moran’s I tests. Coincident sample points were spatially jittered around 0.1° latitude and longitude. Spatial relationships were defined using inverse Euclidean distance. Row standardization was applied to the spatial weights to account for unequal sample distribution and corrections based on the False Discovery Rate were applied to cluster and outlier p-values to account for spatial dependency. Geostatistical analysis followed a linear mixed modelling approach, as described by (47, 48). Briefly, the method uses a linear mixed-effects model (GLMM) to describe, for each observation at a given sampling location, the value of the response variable (δ^15^N) to specified fixed effects, random effects and residual errors (47). Effects can include fixed spatial and bioclimatic covariates, environmental factors that vary spatially but are not considered fixed effect, and factors that differ between locations but that are not spatially correlated (47). The process involves fitting a residual dispersion model to the observed variances at each location, then fitting a mean model. The parameter estimates from these two models are used to compute a predicted value and residual variance for each location across the prediction area. Various models were tested, both with and without the inclusion of bioclimatic covariate data as fixed effects. Different combinations of fixed effects for testing were selected based on known empirical relationships between δ15N and environmental variables, and on interrogation of correlations between faunal δ15N and modelled climatic data, tested with Pearson’s correlation analysis (Section 3 in S3 Supporting Information). Model performance was evaluated using the conditional Akaike Information Criterion (cAIC). This analysis was conducted using the R package spaMM (49). For selected models, time binned δ^15^N isoscapes (interpolated prediction surfaces) were then drawn to visualise spatial variations.

## 3 Results

### 3.1 Data summary and spatial structure

Within the assembled data (n=2,718) mean δ^15^N is 4.1 ± 2.0‰, ranging from -0.9‰ to 11.9‰ (Table 2, Fig 1). Cluster and outlier analysis (Anselin’s Local Moran) demonstrates underlying spatial trends in the data, with higher δ^15^N values clustering at lower latitudes (e.g. northern Spain, Italy, and southwest France) and low δ^15^N values clustering at high latitudes and areas of significant elevation (e.g. Great Britain, Germany, and Alpine regions) (Fig 2). The pattern of spatial clustering varies between different time periods, as does the number of outliers (Fig 2, Table S3 in S3 Supporting Information). Outlying data in these instances are spatial outliers, rather than being outliers in the more standard sense of extreme observations. The spatial outliers could indicate incorrect time bin assignment based on uncertainties in age estimation or could represent true local variability in the faunal δ^15^N data produced by localised environmental variation and/or differences in animal ecology. The outliers may also be an artifact of our data aggregation procedure, particularly for the EOIS3 and LOIS3 time bins which span long periods; it is entirely possible that some samples in close geographical proximity to one another are temporally disparate, and thus combine data representing differing climatic/environmental states. As our interest is in investigating generalised continental-scale spatial patterns, the decision was taken to omit these outliers (n=186) from further analysis. All time bins displayed significant spatial autocorrelation (Global Moran I), indicating systematic spatial variation in the data (Table S3 in S3 Supporting Information). The strength of this spatial relationship varied in time, being strongest for the Last Glacial Maximum (LGM), Last Glacial Termination (LGT), Late Glacial Interstadial (LGI) and Younger Dryas (YD) time bins, and weaker for EOIS3, LOIS3, and early Holocene (EH) time bins.

**Figure 1.**
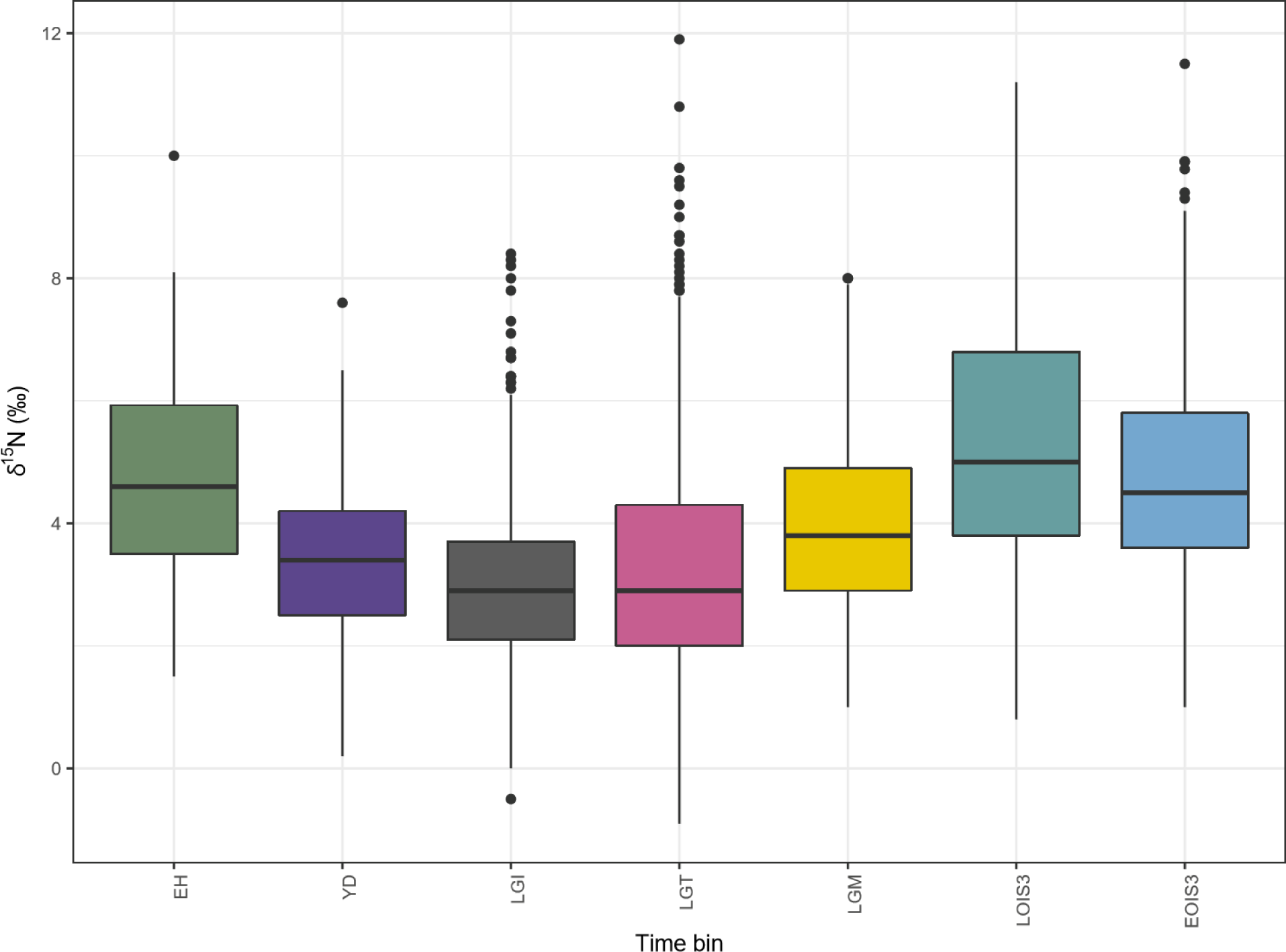
**Boxplot of faunal δ^15^N, plotted by time bin.**

**Figure 2.**
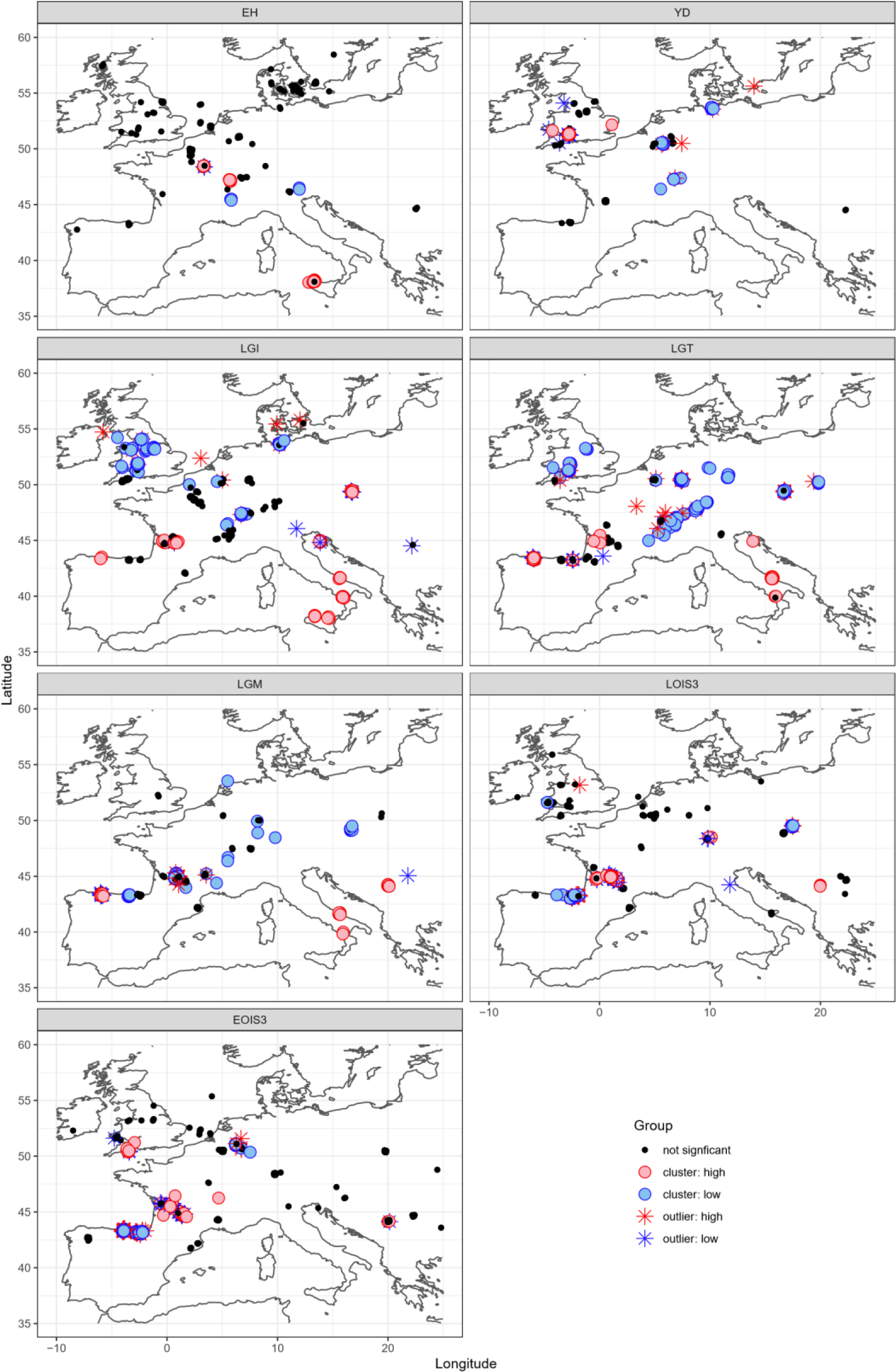
**Visualisation of cluster (circles) and outlier (stars) analysis. High δ^15^N values are indicated in red, low δ^15^N values are indicated in blue.**

**Table 2.** Summary of faunal δ^15^N by time bin

| Age Bin (abbreviation) | n | Mean | Standard deviation | Median | Minimum | Maximum |
| --- | --- | --- | --- | --- | --- | --- |
| Early Holocene (EH) | 176 | 4.7 | 1.6 | 4.6 | 1.5 | 10 |
| Younger Dryas (YD) | 133 | 3.4 | 1.3 | 3.4 | 0.2 | 7.6 |
| Late Glacial Interstadial (LGI) | 485 | 3.1 | 1.4 | 2.9 | -0.5 | 8.4 |
| Last Glacial Termination (LGT) | 602 | 3.3 | 1.9 | 2.9 | -0.9 | 11.9 |
| Last Glacial Maximum (LGM) | 339 | 4 | 1.5 | 3.8 | 1 | 8 |
| Late OIS 3 (LOIS3) | 466 | 5.3 | 2 | 5 | 0.8 | 11.2 |
| Early OIS 3 (EOIS3) | 517 | 4.7 | 1.6 | 4.5 | 1 | 11.5 |

### 3.2 Geostatistical analyses: Isoscapes

Spatial interpolation was first investigated without the inclusion of climate data as covariate fixed effects (Fig 3). Prediction surfaces show the development of a north-south gradient in δ^15^N during LOIS3, which becomes gradually more pronounced through the LGM, LGT and LGI. This contrasts to a lack of obvious spatial patterning of predicted δ^15^N in the earlier EOIS3 and later YD and EH time bins. Notably, the amplification of the latitudinal gradient, particularly during the LGT and LGI, appears to primarily be driven by a decrease in δ^15^N in northerly locations, rather than by an increase in δ^15^N in southerly locations. The lowest δ^15^N values, which are observed at northern latitudes only during the LGT and LGI, are absent in the LGM (the coldest part of the last glacial cycle), suggesting that the trend cannot be explained by temperature change alone.

**Figure 3.**
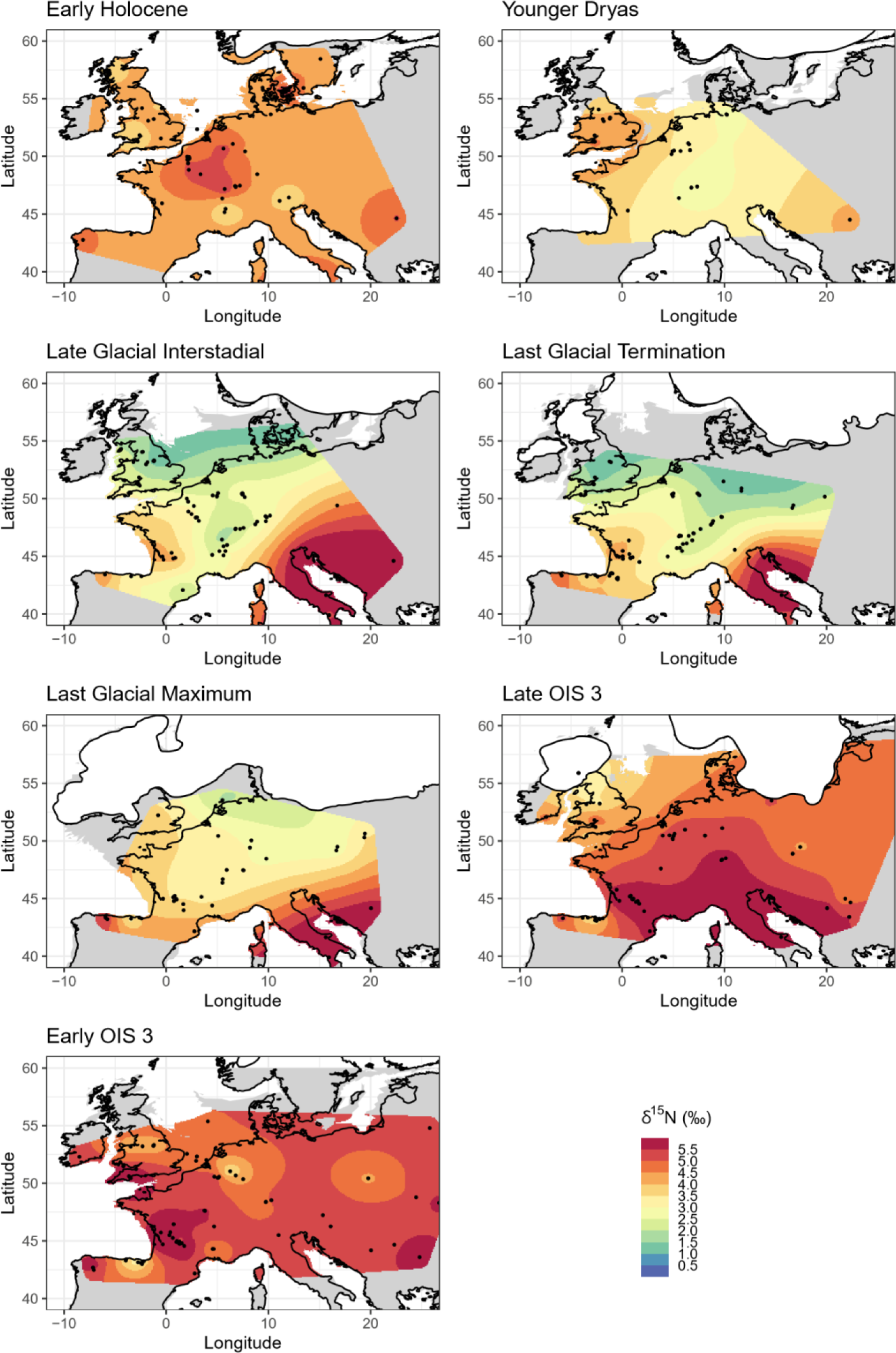
δ^15^N isoscape prediction surfaces, modelled using random effects only. Palaeocoastline data from (79), ice sheet extent from (80) and modern coastline from (81), as detailed in S4 Base maps for figures.

Model performance evaluated through prediction variance surfaces (Fig 4) show that variance ranges from >1.5‰ across much of the prediction area for LOIS3 to <1‰ for EOIS3, LGM, LGT, LGI. Variance is lowest closest to sample locations, but otherwise does not appear to be spatially structured. Comparing predicted δ^15^N values to observed δ^15^N values at sample locations shows that models for all time periods perform well, explaining between 47 and 91% of the variance, with Root Mean Squared Errors (RMSE) ranging from 0.40‰ to 1.11‰ (Fig 5).

**Figure 4.**
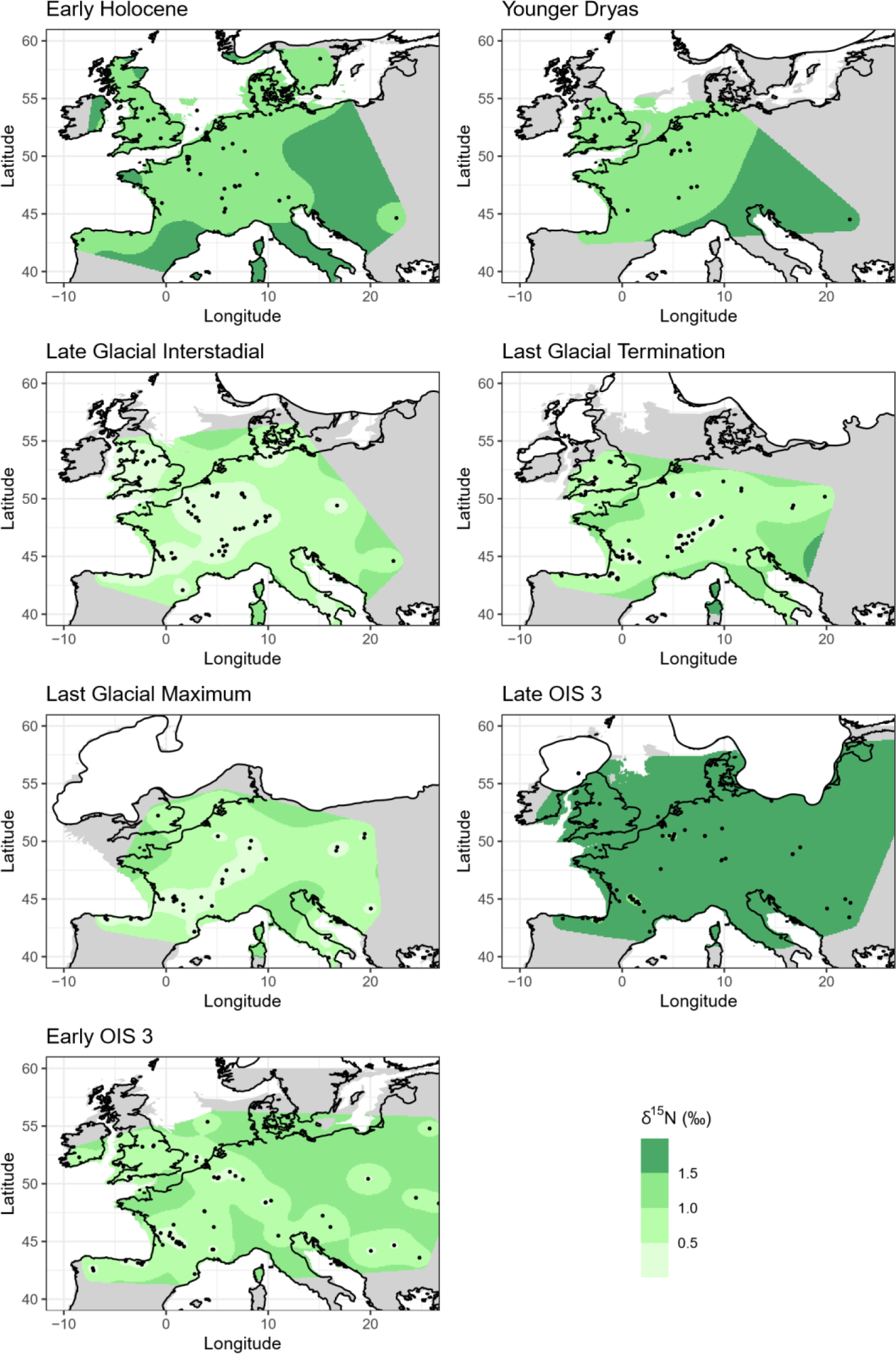
δ^15^N isoscape variance surfaces, modelled using random effects only. Base maps as described in. Fig 3 **caption.**

**Figure 5.**
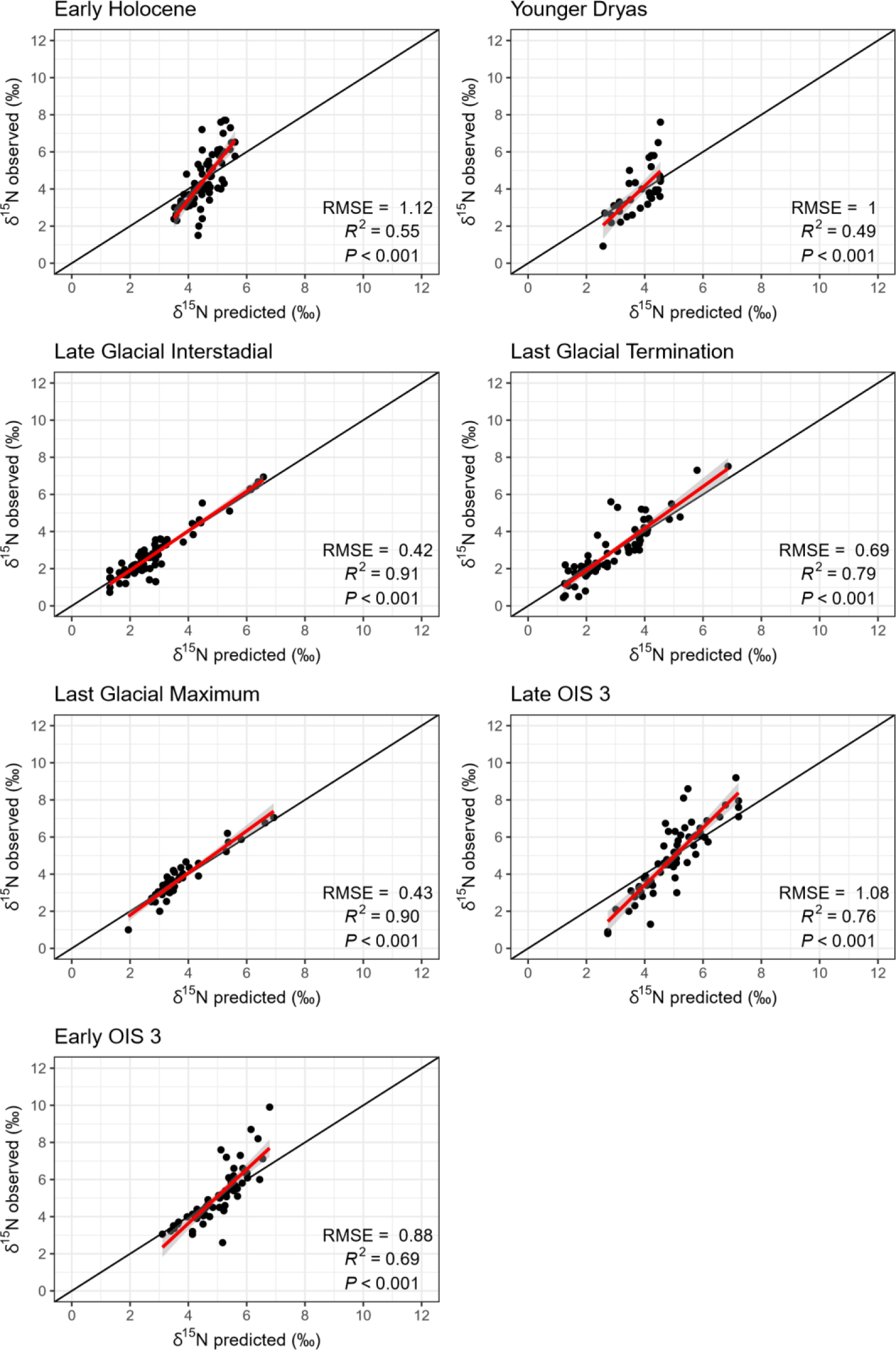
**Comparison of observed site mean δ^15^N versus model predicted δ^15^N, for the models using random effects only.**

To explore possible relationships between spatial variation in faunal δ^15^N and climate, correlations between site-mean δ^15^N and modelled terrestrial climatic variables from Beyer et al (2020), as well as elevation, were tested (Table S4.1 and S4.2 in S3 Supporting Information). Notable differences occur in correlations between δ^15^N and climatic variables across different time bins and no single variable showed significant correlation with δ^15^N in every time bin. Importantly, no significant correlations were identified for the early OIS 3. In selecting which variables and combinations of variables to test as fixed effects in the isoscape models, we considered known empirical relationships between δ^15^N and climate in the modern environment, issues of collinearity between climatic variables, and the strength of correlations between faunal δ^15^N and modelled climatic data (full analysis reported in Section 4 in S3 Supporting Information). From this analysis the variables selected as fixed effects for model testing were mean annual temperature (MAT), mean annual precipitation (MAP), temperature of the warmest quarter, precipitation of the warmest quarter and precipitation of the coldest quarter (Fig S4.2 – S4.6 in S3 Supporting Information). Different combinations of these variables were considered and model performance was evaluated using the cAIC (Table 3 and Section 5 in S3 Supporting Information). Given that the strength of correlation with covariate data differed by time bin, it was unsurprising to find that the best fit model also differed by time bin. For all time bins, models including bioclimatic fixed effects performed better than the model where no fixed effects were included, with the exception of the LGI. However, for all time bins the changes in the cAIC criterion between different models were for the most part extremely small (Table 3), suggesting most models performed similarly well, and the inclusion of bioclimatic variables did not significantly improve model performance for any time bin.

**Table 3.** Model fit results (cAIC) for faunal nitrogen isoscape prediction models. Fixed effects tested include mean annual temperature (MAT), mean annual precipitation (MAP), temperature of the direst quarter (temp.dry), temperature of the warmest quarter (temp.warm), and precipitation of the warmest quarter (precip.warm). A spatially-structured random effect following a Matérn correlation structure using latitude and longitude to compute distances between observations (spatial), and an uncorrelated random effect identical for all observations from the same location (site) were also included. Further fit results are given in Supplementary Information 4. The best performing model, based on cAIC, is highlighted in red.

| Model parameters | cAIC |  |  |  |  |  |  |
| --- | --- | --- | --- | --- | --- | --- | --- |
|  | EH | YD | LGI | LGT | LGM | LOIS3 | EOIS3 |
| No fixed effects + spatial + site | 213.3 | 77.9 | **110.5** | 132.6 | 54.1 | 170.2 | 181.2 |
| MAT + spatial + site | 212.7 | 78.0 | 110.8 | **131.8** | 54.2 | 170.8 | **181.0** |
| MAP + spatial + site | 210.9 | 77.9 | 110.9 | 132.9 | 54.3 | **169.3** | 181.9 |
| MAT + MAP + spatial + site | **210.3** | 78.1 | 111.2 | 132.4 | 54.3 | 169.3 | 181.1 |
| MAT + MAP + MAT:MAP + spatial + site | 210.9 | 78.3 | 111.6 | 132.5 | 54.8 | 169.7 | 182.0 |
| temp.warm + spatial + site | 211.9 | **77.7** | 110.7 | 131.9 | 56.6 | 170.4 | 181.8 |
| precip.warm + spatial + site | 211.4 | 77.9 | 110.8 | 133.1 | 54.0 | 169.4 | 181.6 |
| precip.cold + spatial + site | 211.9 | 78.0 | 111.1 | 132.9 | 53.7 | 169.9 | 182.2 |
| temp.warm + precip.warm + spatial + site | 210.8 | 77.8 | 111.1 | 132.2 | 56.4 | 169.8 | 182.2 |
| temp.warm + precip.cold + spatial + site | 211.2 | 78.0 | 111.2 | 132.1 | 55.6 | 170.0 | 182.7 |
| Precip.cold + precip.warm + spatial + site | 211.4 | 78.1 | 111.3 | 133.3 | **52.9** | 169.5 | 182.5 |
| temp.warm + precip.cold + precip.warm + spatial + site | 211.2 | 78.0 | 111.4 | 132.4 | 54.9 | 169.8 | 183.0 |

Figures 6, 7 and 8 present results from the best performing model incorporating climatic fixed effect(s) for each time bin. In very general terms, the best fit model incorporating climatic fixed effect(s) (Fig 6) predicted somewhat similar continental-scale spatial patterns of δ^15^N as the corresponding model without fixed effects (Fig 3), although with some notable differences. Importantly, the strength of expression of the north-south gradients in δ^15^N is more muted when climatic variables are incorporated, particularly for LOIS3, the LGT, and the LGI. Greater localised variation in δ^15^N is also apparent when climatic variables are incorporated, related to localised spatial climatic gradients, such as those that exist across areas of varying topography (e.g. the Alps mountain range). The prediction variance surfaces (Fig 7) show only minor differences when compared to those for models without fixed effect (Fig 4). When predicted and observed δ^15^N values are compared for models incorporating climatic fixed effects (Fig 8), all performed slightly worse than those without fixed effects (Fig 5), with r^2^ values ranging from 0.31 to 0.91 and RMSE ranging from 0.51‰ to 1.22‰.

**Figure 6.**
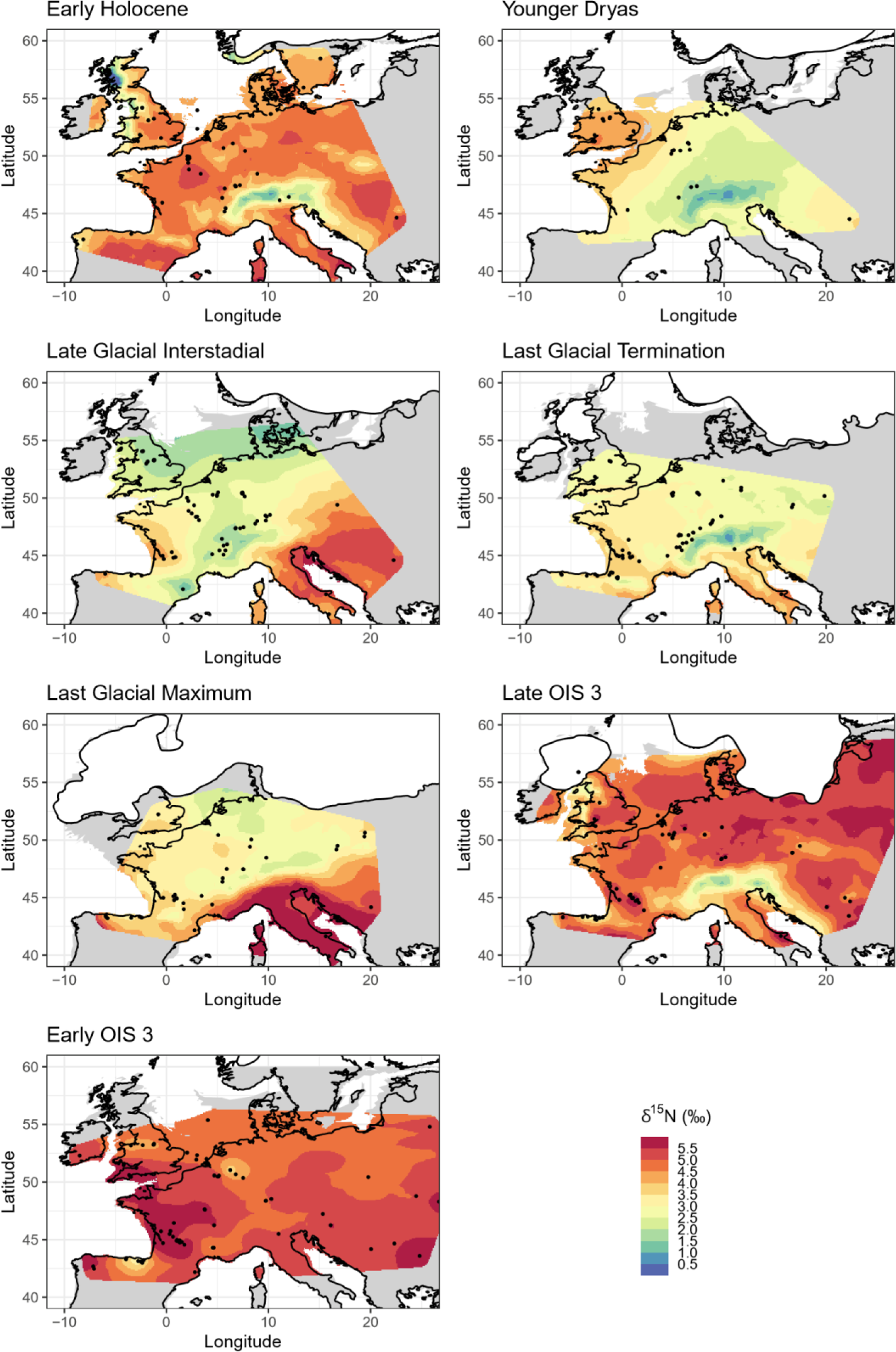
δ^15^N isoscape prediction surfaces, best performing model incorporating climatic fixed effect(s) for each time bin. Base maps as described in. Fig 3 **caption.**

**Figure 7.**
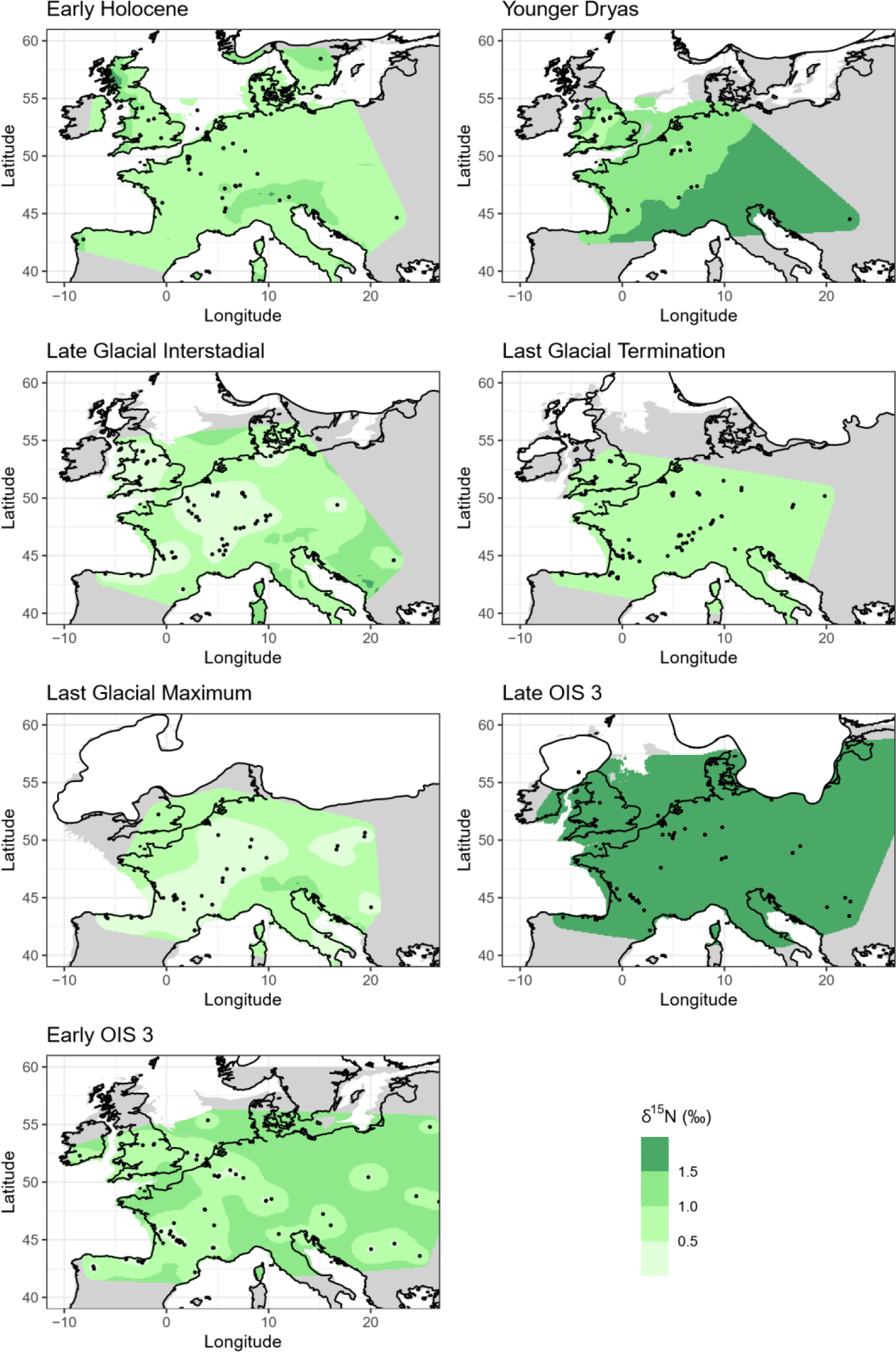
δ^15^N isoscape variance surfaces, best performing model incorporating climatic fixed effect(s) for each time bin. Base maps as described in. Fig 3 **caption.**

**Figure 8.**
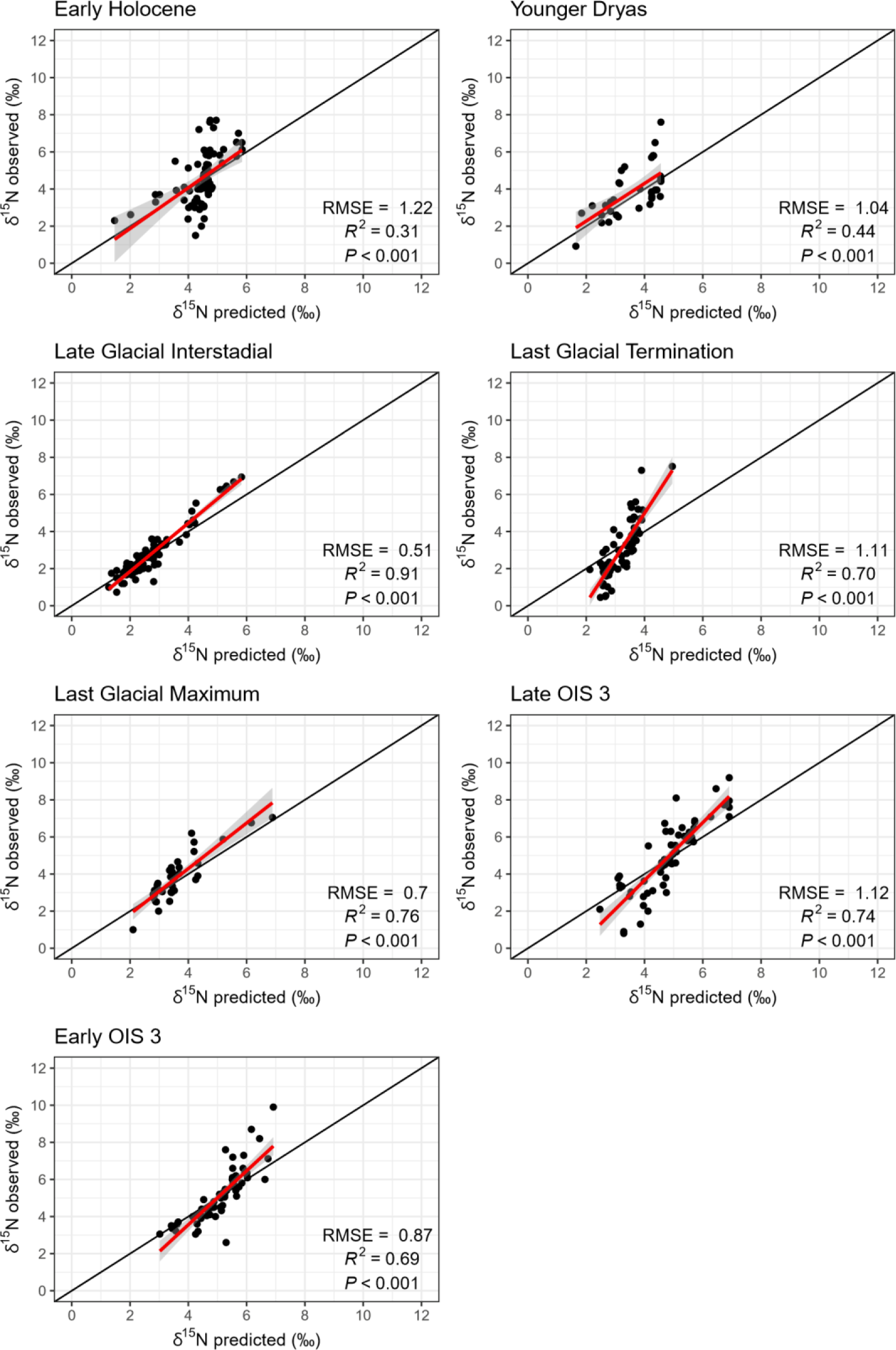
**Comparison of observed site mean δ^15^N versus model predicted δ^15^N, best performing model incorporating climatic fixed effect(s) for each time bin.**

## 4. Discussion

### 4.1 Isoscape mapping using faunal isotope data

This study represents the first (to our knowledge) study to create temporally-layered isoscapes using palaeo-data. While the application of isoscape approaches in modern terrestrial environmental and ecological research are now relatively widespread (50), the use of isoscape modelling in palaeo-focused research has so far been considerably more limited (51–55). In part, this can be attributed to the additional complexities that palaeo-isoscapes must contend with; while plant, soil or animal δ^15^N is a spatially and temporally continuous variable, our means of sampling such data is inescapably discretized (i.e. each sample represents a discrete temporal and spatial interval). The discretization is many orders of magnitude larger in fossil than in modern data ensembles, owing to the uncertainties in establishing calendar age estimates for fossil samples, the need to consider samples of different ages together as a single temporal unit, and the assumptions that must be made about the spatial resolution and provenance of the sample. As such, isoscapes constructed using palaeo-data will always have a certain level of unavoidable uncertainty inbuilt.

Likewise, probably owing to the complex nature of the terrestrial nitrogen cycle and relative data paucity compared to other environmental systems (e.g. oxygen and hydrogen in the hydrological cycle), the application of geostatistical approaches specifically toward modern terrestrial nitrogen isotope data have so far also been comparatively limited (5,56–58). Part of the difficulty in assessing regional/global scale gradients in δ^15^N is that the nitrogen isotope composition of soils and plants may be highly heterogeneous at very localised spatial and/or temporal scales (59). In this regard, relying on bone collagen data may actually be advantageous; herbivores act as natural integrators, providing a measure of ecosystem nitrogen that is spatially averaged over the extent of the animal’s home range, and temporally averaged over a number of years (temporal resolution depends on bone collagen turnover rate, but is typically in the order of several years). Therefore, while the use of faunal δ^15^N to trace changes in underlying environmental δ^15^N introduces noise from dietary and behavioural differences, it also offers a unique means to assess ecosystem-scale variation in δ^15^N, particularly in past environments, where other sampling opportunities are lacking or inadequate.

A recent study by Barrientos et al. (52) illustrated the potential of using archaeological bone collagen δ^15^N data in palaeo-isoscape mapping. The resultant Inverse Distance Weighted (IDW) isoscapes demonstrated how geostatistical approaches, rooted in community and trophic ecology, could be applicable to addressing archaeological questions (52). In this study we have progressed these ideas, demonstrating the possibility of applying more complex geostatistical methods, which, unlike a IDW approach, allow for errors and uncertainties to be quantified and covariate data to be incorporated. The approach followed here demonstrates a means to consider variations in spatial gradients in δ^15^N through time.

The prediction models presented provide a method to quantitively estimate site-averaged herbivore δ^15^N in the past, at locations where empirical data is absent. While such isoscape approaches should not replace efforts to establish local and time-specific baseline data through empirical sampling, they offer a complementary source of information through which past environments can be explored. The two approaches need to work hand-in-hand, as continued efforts to generate empirical data will ultimately lead to improvements in the predictive power of isoscape models. While the nature of fossil sample material dictates that there will always be limitations in the temporal and spatial accuracy of such an approach, the value of such predictive maps for investigating long term continental-scale changes in the terrestrial nitrogen cycle, and natural- and anthropogenically-driven impacts on said cycle, should not be understated.

### 4.2 Species-specific spatial gradients of δ^15^N variability

One of the primary challenges in understanding the variability present in fossil δ^15^N data is to distinguish between environmental effects and the effects of dietary and ecological differences between species, and the variability that can occur in both effects across space and time. Some previous studies of environmental change restricted analyses to single species to limit variability introduced by dietary ecology (e.g.(17, 23)). Others took data from more than one species with similar dietary characteristics and applied data normalisation/transformation procedures (e.g.(25)). The former severely limits the size of the data set available for analysis, while the latter, when used to infer environmental change, relies on the assumption that species’ dietary behaviour and isotope niche relative to one another have remained stable through time. Empirical evidence suggests this assumption is problematic (1,3,60,61). Indeed, species most capable of dietary flexibility are often most successful at adapting to changes in local environmental conditions (62, 63), and thus it is these species that are most abundant in the fossil record and remain present across major climate transitions. In this study we applied no correction or data transformation procedure to account for species-based differences, and instead considered only site mean δ^15^N values in the models thus far presented. Our primary reasoning for this is that while species-based differences are present in the data, neither the option of restricting analysis to a single species or applying species-based corrections to a multiple-species analysis were appropriate in this instance.

Nonetheless, it is important to consider the implications of our approach compared to model outputs when different species are considered independently. While the data is not of sufficient quantity to enable species-specific isoscapes to be constructed for all time bins and species, they can be considered in some contexts. Our data is dominated by 3 species: horse (25%), reindeer (24%) and red deer (30%), and while their geographical distribution varies considerably in time (Fig S6.1 in in S3 Supporting Information), data is of sufficient quantity and comparable geographic distribution to consider isoscape models for each of these species for the LGT and LGI time bins (Fig 9 and 10).

**Figure 9.**
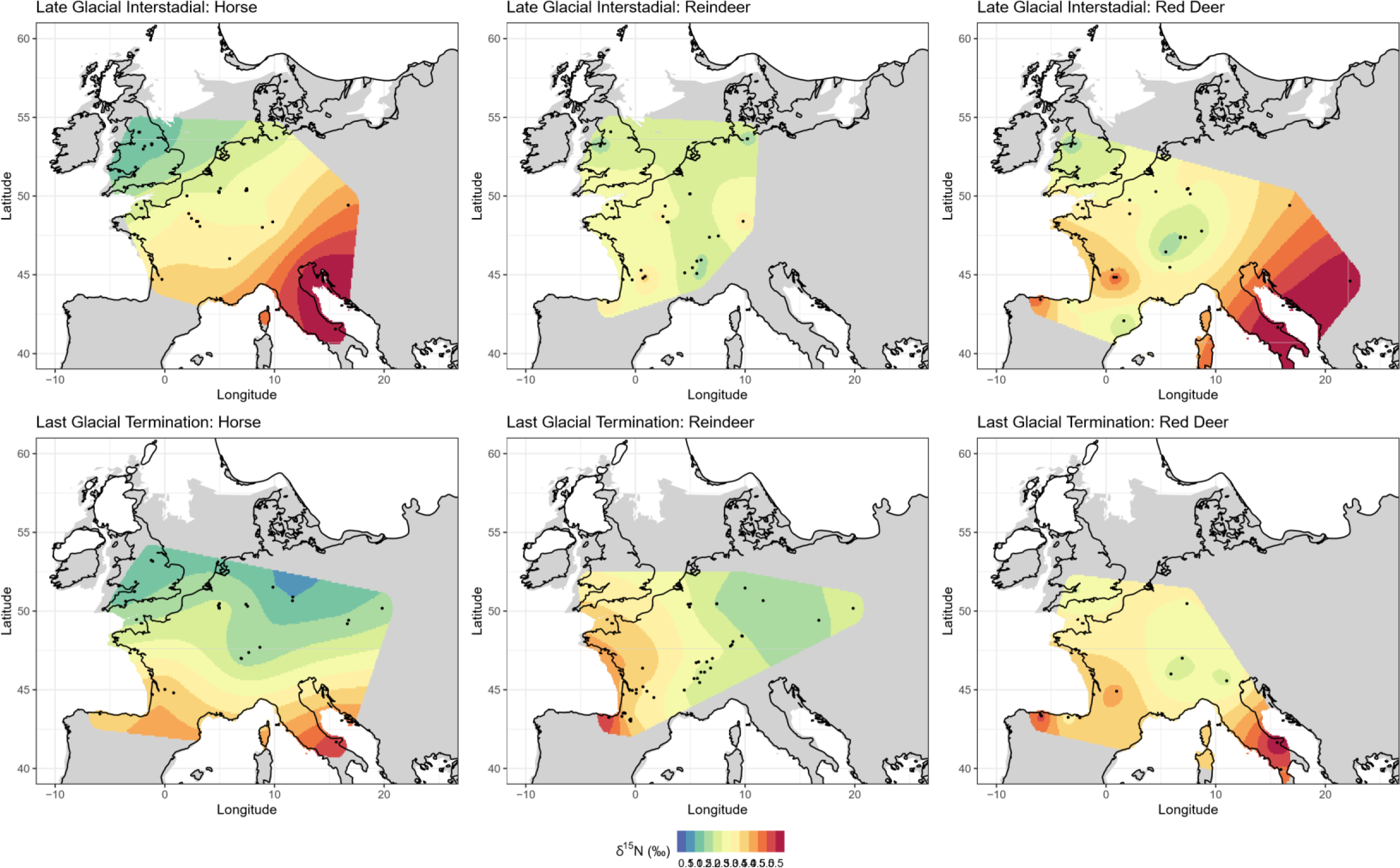
δ^15^N isoscape prediction surfaces for *Equus* sp., *Rangifer tarandus*, and *Cervus elaphus* for the Last Glacial Termination and Late Glacial Interstadial time bins. Linear mixed models were run without the addition of environmental covariate data. Base maps as described in Fig 3 caption.

**Figure 10.**
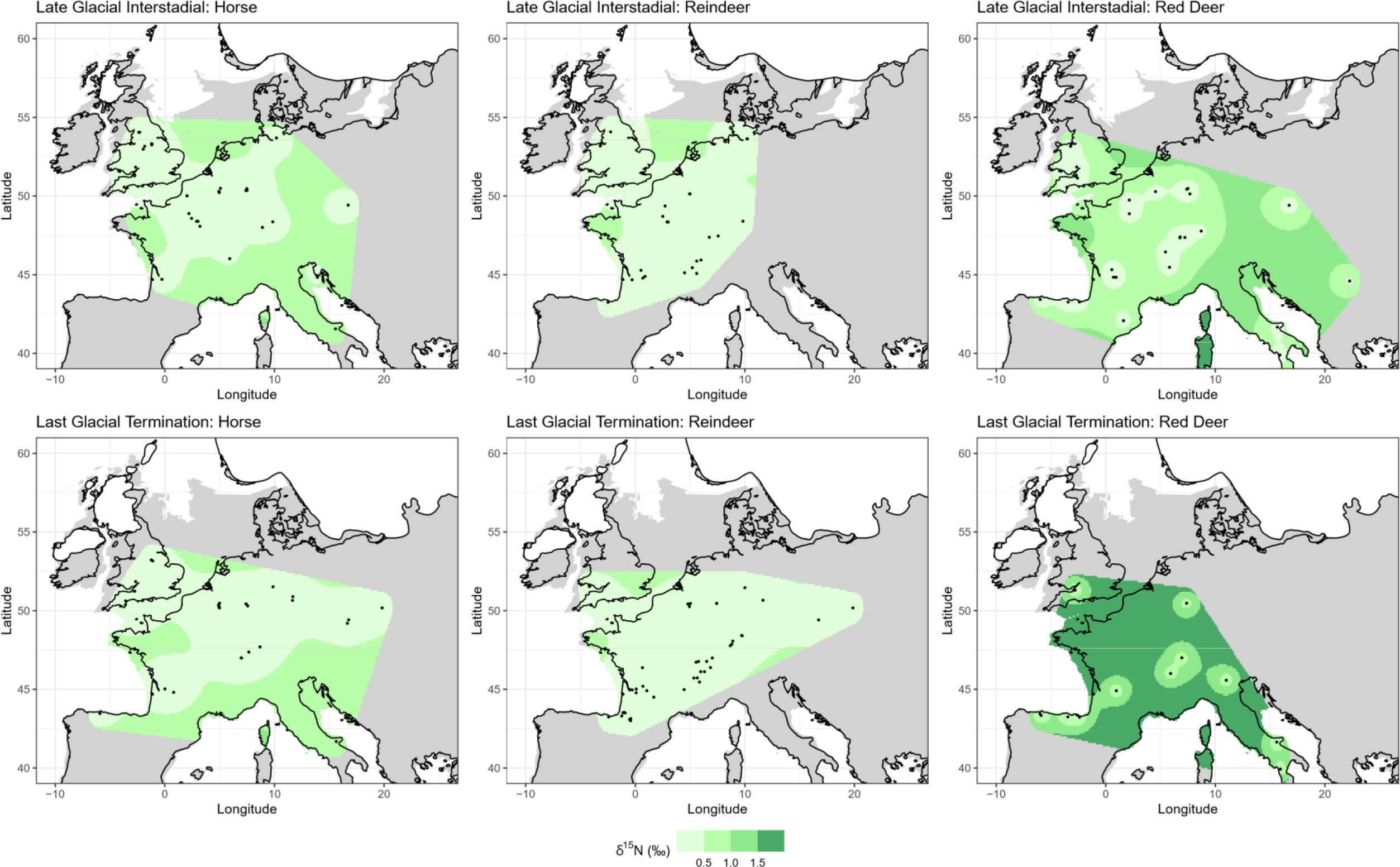
δ^15^N isoscape variance surfaces for *Equus* sp., *Rangifer tarandus*, and *Cervus elaphus* for the Last Glacial Termination and Late Glacial Interstadial time bins. Linear mixed models were run without the addition of environmental covariate data. Base maps as described in Fig 3 caption.

From this analysis horse can be seen to display the strongest spatial gradients in δ^15^N in both the LGT and LGI, with lowest values in the north and northwest of the interpolation area, and highest values in the south. North-south gradients in δ^15^N are also seen in the reindeer and red deer data but are more muted. While these differences likely stem from differences in dietary ecology and mobility (see Section 2 of S3 Supporting Information for more detailed discussion), the implications for this in our objective of understanding large-scale changing spatial gradients in δ^15^N requires consideration. Deciphering the relative contributions of dietary behaviour and environmental influence on the δ^15^N signal is extremely challenging, and it may not be possible to fully disentangle the two by measuring bulk collagen δ^15^N alone; as localised environmental conditions exert strong influence on feeding behaviours and diet, the two are inextricably linked. In the context of late Pleistocene northern Europe, tooth meso- and microwear analysis confirm horse most likely had a graze-dominated diet, red deer a browse-dominated diet, and reindeer a mixed diet (63). However, the extent to which these different plant types in the diet can be equated to isotopically distinct diets is debateable. As Schwartz-Narbonne et al. (3) discuss, while at the most generalised scale patterns of δ^15^N in tundra ecosystems can be summarised as shrub < lichen < herb < fungi, there are examples where this does not hold true in either space or time (64–67). If this generalised plant type δ^15^N pattern was taken at face value and applied to the late Pleistocene European herbivores, a pattern of red deer < reindeer < horse would be expected; in fact, the opposite pattern is identified.

Part of the difficulty in relating faunal δ^15^N to plant δ^15^N is the issue of scale. Plant δ^15^N is highly heterogenous and is related to N availability and a plant’s ability to utilise and acquire different forms of N, which are influenced by root depth and mycorrhizal association, as well as environmental factors which can vary on a sub-annual scale (4). In comparison, one collagen δ^15^N analysis represents a homogenised data point, averaging a multitude of plant δ^15^N values at a spatial scale equivalent to the animal’s home range (which can vary considerably between species) and a temporal scale of several years. A further consideration in interpreting the faunal signal, particularly that of reindeer, is the inclusion of lichen in the diet. Lichens fix nitrogen from the atmosphere, and therefore species consuming a significant proportion of lichen may display δ^15^N signatures decoupled from environmental-mediated changes in vegetation δ^15^N. However, the amount of lichen consumed, and its contribution to the reindeer bone collagen δ^15^N signal cannot be easily discerned. For example, significant differences in the amount of lichen incorporated into reindeer diets between the LGI and YD in northern Europe, based on tooth meso- and micro-wear analysis, did not translate to differences in bone collagen δ^15^N (68). As such, we decided that the exclusion of reindeer from the data, which would have significantly reduced the sample size, was not justified in this instance.

In the future, with ever increasing amounts of faunal isotope data and radiocarbon dates being published, it is hoped that species-specific geostatistical analyses can be further explored. Such investigations would undoubtedly be of great benefit to furthering our understanding the isotope ecology and niche overlap/partitioning of key herbivore species, and of the complex and competing influences of environment and ecology on faunal δ^15^N.

### 4.3 Evaluating spatial gradients and drivers of δ^15^N variability in the past

The results presented here provide the means to visualise the spatiotemporal character of changing faunal δ^15^N, and, when combined with the recent publication of high-resolution climate model data (27), interrogate potential links between faunal δ^15^N and climatic variables at a resolution not previously achievable.

Our results show spatial gradients in faunal δ^15^N appear in late OIS 3, strengthening during the Last Glacial Maximum, Last Glacial Termination and Late Glacial Interstadial. This compares to early OIS 3 and the Younger Dryas and Holocene where strong spatial gradients are absent. The amplification of the latitudinal gradient appears to reach its maximum during the Late Glacial Interstadial, when the lowest δ^15^N values (<2‰) are predicted. The fact that these lowest values occur during the Late Glacial Interstadial, a relatively warm climatic period, and not during the coldest part of the last glacial cycle, show that temperature is not the primary driver of variation. We draw attention to the location of the lowest predicted δ^15^N values, in regions that were either glaciated or were immediately proximal to the British, Scandinavian, and Alpine ice sheets during the LGM (Fig 3 and 6). It is noteworthy that δ^15^N values of <2‰ only occur within the zone of continuous permafrost that existed across Europe at the height of the last glacial (69), and their occurrence within this zone is only after the onset of deglaciation and thaw. The role of increased landscape moisture driven by increased precipitation and increased input of meltwater from icesheets and thawing permafrost has long been suggested as a driver of the LGNE (17,18,22,23,25), and the results presented here add further weight to this interpretation. This environmental change would have both altered the floral community in such landscapes and altered the form and source of nitrogen available to vegetation, with microbially-mediated changes in N cycling between pools of NO3^-^ (nitrate), NH4^+^ (ammonium), and N2 (elemental nitrogen) resulting in changing plant δ^15^N (70–72). Biogeochemical cycles and microbial activity in cold environments may be particularly sensitive to changes in soil moisture content, O2 status, and temperature (70,73,74). In this regard, the use of geostatistical interpolation to reconstruct changing δ^15^N spatiotemporal gradients may provide the means to further interrogate sub-continental scale processes of permafrost thaw and changing landscape moisture during the terminal Pleistocene in Europe. If increased data availability in coming years were to enable faunal isoscape mapping at an increased temporal resolution for the late glacial, it would certainly be of interest to compare these to contemporaneous maps of changes in the distribution of European permafrost.

Regarding the absence of spatial gradients in the early OIS 3, Younger Dryas and early Holocene time bins, while the previously discussed problems of data aggregation across multiple climatic events may explain the muted gradients in early OIS 3, this explanation is unsatisfactory for the early Holocene and Younger Dryas. A more plausible explanation, at least for the Early Holocene is that the reduced spatial gradient in δ^15^N is the result of less pronounced climatic gradients across Europe during this time period, as is evidenced in both proxy-based data and model simulations (75, 76). For the Younger Dryas, the expression of the rapid cooling event in European proxy archives is inconsistent (77, 78); this, coupled with the brevity of the event (c. 1200 years), potential lag in environmental response, and uncertainty in assigning faunal samples to such a narrow age bracket may go some way to explaining difficulties in understanding the resultant isoscape model.

Despite our finding that the inclusion of climatic covariate data did not improve isoscape model performance, our analysis nonetheless shows that palaeo-fauna δ^15^N, when averaged by location, is correlated with modelled temperature and precipitation, with relationships similar to those that are observed between modern plant and soil δ^15^N, and MAT and MAP (5–8). What is most striking about the investigated palaeo-fauna δ^15^N–climate correlations, is the strength of the relationship between site mean faunal δ^15^N and MAT (as well as temperature of the warmest quarter and precipitation of the warmest quarter) during the Last Glacial Maximum, Last Glacial Termination and Late Glacial Interstadial (fig. S4.2; S4.4 and S4.5). Such relationships are far stronger than for the other time periods considered and are also stronger than those observed for modern soil/plant δ^15^N – MAT relationships (6, 7). Interestingly, while the relationship between foliar and soil δ^15^N and MAT has been shown not to hold true in modern low temperature environments (<-0.5°C for foliar δ^15^N and <9.8°C for soil δ^15^N (6, 7), we do not observe such inflection points in our data, although it should be noted that few samples come from environments where MAT is predicted to be <0°C.

Also notable is the stronger correlation between site mean faunal δ^15^N and precipitation of the warmest month, compared to the relationship with MAP (Fig S4.3 and S4.5 in S3 Supporting Information). The relatively weak correlation between faunal δ^15^N and MAP for most time bins does not mirror those seen in the modern environment. In part, this may relate to the comparatively more complex nature of reconstructing palaeo-precipitation, and the poorer performance of precipitation models when compared to proxy data-based reconstructions, than for reconstructing temperature (27). However, this result may also represent the importance of the complex interplay of temperature and precipitation in determining δ^15^N. Further, the seasonal cycle of plant growth and N requirements/availability may be responsible for seasonally distinct relationships between δ^15^N and climate.

Keeping in mind the caveats of our analysis, that; 1) faunal δ^15^N is only indirectly related to climate, being mediated also by the interplay of species-specific characteristics and inter-species interactions; that 2) the climate data we are using is modelled output, not empirical measurements; and 3) the assembled data aggregates δ^15^N across multiple species and potentially disparate time periods, the presented results offer an intriguing insight into the spatial and temporal variability of δ^15^N in the past.

## 5. Conclusion

The isoscape models presented here represent the first (to our knowledge) attempt to create time-sliced maps of terrestrial δ^15^N gradients based on archaeological and palaeontological animal isotope data. In addition to compiling and critically evaluating previously published data, our analysis includes the publication of several hundred new faunal δ^15^N data and radiocarbon dates. The analysis presented here serves two main purposes; to investigate changes in spatial gradients of δ^15^N in late Pleistocene Europe, with a view to investigating the Late Glacial Nitrogen Excursion, and to demonstrate more broadly the application of isoscape approaches to palaeo-data with implications for how baseline data is understood and used in archaeological and palaeoecological research. Our results have shown clear changes in spatial gradients of δ^15^N through time, that are most likely related to changes in landscape moisture (particularly from increased input of meltwater from icesheets and thawing permafrost) that occurred after the Last Glacial Maximum. Our analysis found that the inclusion of climatic covariate data in the models did not significantly improve model performance, suggesting that the combination of the variables considered did not fully capture the drivers producing the observed spatial variation in the δ^15^N faunal data. Our results highlight the significant opportunities (and challenges) of applying isoscape approaches to faunal data. We demonstrate how data from multiple species of different ages can be combined to form data sets suitable for geostatistical interpolation. With the continued publication of faunal isotope data from archaeological and palaeontological assemblages, it is likely that in the coming years the accuracy and the temporal and spatial resolutions of such models can be much improved upon. Such models can make an important contribution to understanding baseline δ^15^N values for terrestrial food web analysis and in the interpretation of data from higher trophic level animals in relation to mobility, migration, and dietary research. Moreover, improved understanding of baseline δ^15^N in late Pleistocene and early Holocene contexts provides a background reference against which subsequent human impact on the nitrogen cycle and overall landscape health, such as through farming practices and deforestation, can be assessed.

## Supporting information

S4 Base maps for figures

S2 Script

## Acknowledgments

This work was made possible by the support of a great many colleagues and institutes, who facilitated access to, and permitted sampling of, numerous archaeological and palaeontological collections. We would like to thank Roger Jacobi, Chris Stringer and the Ancient Human Occupation of Britain project, Adrian Lister, Sonja Grimm, Sophy Charlton, Ian Barnes, Melissa Marr, Tom Lord, Terry O’Connor, Linda Wilson, Graham Mullan and the University of Bristol Spelaeological Society Museum, Barry Chandler and Torquay Museum, Jan Freedman and Plymouth City Museum & Art Gallery, Brian Lewarne and the Devon Kart Society, the Museum of Archaeology and Anthropology, Cambridge, the Potteries Museum & Art Gallery, Stoke-on-Trent, Buxton Museum & Art Gallery, Elizabeth Walker and the National Museum Cardiff, Rebecca Miller, the University of Liege, Mietje Germonpré, Annelise Folie and the Royal Belgian Institute of Natural Sciences, Petr Neruda, Zdenka Nerudova, Martina Robličková and the Moravian Museum, Alex Pryor, Jiri Svoboda, Nick Conard, Susan Münzel, Wells & Mendips Museum, Somerset, Spyridoula Pappa, Pip Brewer and the Natural History Museum London, Lucy Astill and Creswell Crags Museum and Prehistoric Gorge, Sheffield Museums, Thomas Terberger, Piotr Wotjal, Marta Poltowicz-Bobak, Dariusz Bobak, and the Institute of Systematic and Evolution of Animals, Polish Academy of Sciences.

## Supporting Information

S1 Dataset

S2 Script

S3 Supporting Information

S4 Base maps for figures

