## Supplementary material for "Nitrogen palaeo-isoscapes: Changing spatial gradients of faunal δ^15^N in late Pleistocene and early Holocene Europe": S3 Supporting Information

Nitrogen palaeo-isoscapes:

* Corresponding author

### 1 Sample preparation and isotope and radiocarbon analytical methods

Of the 479 newly generated $\delta^{15}$N data reported in this study, 68 came from pre-existing collagen samples and 411 came from newly collected material from which we extracted the collagen. The pre-existing collagen samples were originally produced as part of other research projects, for example those focusing on radiocarbon dating, and the leftover collagen was kindly donated to this study for the purpose of $\delta^{15}$N analysis.

For the newly collected material, a small (<1g) sample of bone or tooth dentine was taken from each specimen using a dental drill with either a small cutting wheel or tungsten burr attachment. This sample collection was conducted over a number of years and collagen extraction and isotope analysis was performed at different laboratories. The Supplementary Data File details which of the following collagen extraction methods were followed for each sample, and the analysing laboratory.

Collagen extracted at University College London (UCL) followed a modified version of the Oxford Radiocarbon Accelerator Unit (ORAU) collagen extraction procedures (AF and AG methods (1)), which is based on a modified version of the Longin method (2). For samples that had been, or were suspected to have been, conserved with PVA glue, a solvent extraction pre-treatment was used to remove the adhesive (denoted as AG* or AF* method). All samples were demineralised in 0.5 M hydrochloric acid (HCl) at 4°C and then thoroughly rinsed with ultrapure water. Unless otherwise indicated in the Supplementary Data File, all samples were then treated with 0.1 M sodium hydroxide (NaOH) for 30 minutes to remove humic acids, before being thoroughly rinsed. Samples were then gelatinised in pH 3 HCl solution at 75°C for 48h and filtered using a pre-cleaned Ezee-filter (AG method). For some samples, including all those radiocarbon dated, the filtrate was then passed through a pre-cleaned 15–30 kD ultrafilter, with the > 30 kD fraction collected and freeze-dried (AF method).

Collagen extracted at RLAHA, University of Oxford also followed a modified version of the Longin method (2), which is detailed in Stevens and Hedges (3). For samples that had been, or were suspected to have been, conserved with PVA glue, a solvent extraction pre-treatment was used to remove the adhesive (RLAHA Method 1 and 3 used this step). Solvent extraction involved heating the sample at 40°C for an hour in distilled water, then repeating the heating process using acetone, distilled water, methanol, and distilled water, respectively. Samples were then demineralised in 0.5 M aq. HCl at 4°C until the mineral fraction had dissolved and then rinsed three times with distilled water (RLAHA Methods 1-4 followed this step). Where sample amount was sufficient and where deemed necessary 0.1 M NaOH was added for 30 minutes to remove humic acids (RLAHA Method 1 and 2 used this step). Samples were then rinsed with distilled water, gelatinised in a pH 3 solution for 48 hours at 75 °C (RLAHA Methods 1-4 followed this step). The filtered supernate containing the soluble collagen was then collected, frozen, and lyophilized.

Collagen extracted at the University of Cambridge again followed a broadly similar method, which is detailed in Stevens et al (4). Samples were demineralised in 0.5 M aq. HCl at 4 °C until they had fully demineralised. Samples were then rinsed in distilled water and gelatinised by heating in pH 3.0 aqueous solution at 75 °C for 48 h. The liquid fraction containing the dissolved collagen was filtered off, frozen overnight at −20°C, then stored at −80°C for 4 h and finally lyophilised.

Isotopic analysis at the Scottish Universities Environment Research Centre (SUERC) was undertaken using a Delta V Advantage continuous-flow isotope ratio mass spectrometer coupled via a ConfloIV to an IsoLink Elemental Analyser (Thermo Scientific, Bremen). Between 1.2 and 1.5mg collagen was loaded into a tin capsule for continuous flow combustion and isotopic analysis. For every ten archaeological samples, three in-house standards that are calibrated to the International Atomic Energy Agency (IAEA) reference materials USGS40, USGS41, IAEA-N-1 were run (5). Results are reported as per mil (‰) relative to the internationally accepted standard AIR. Precision was determined to ± 0.2‰ for $\delta^{15}N$ based on repeated measurements of calibration standards.

Isotopic analysis at RLAHA, University of Oxford was undertaken using an automated Carlo Erba carbon and nitrogen elemental analyser coupled with a continuous flow isotope ratio-monitoring mass spectrometer (PDZ Europa Geo 20/20 mass spectrometer). Between 2.5 and 3.5 mg of collagen was loaded into a tin capsule for isotopic analysis. For every six samples, two in-house reference standards Nylon (Nylon 66, BDH, UK), and Alanine (L-Alanine, Fluka, UK) whose isotopic values were calibrated against IAEA standards IAEA-N-1 and IAEA-N-2 were run. Results are reported on the delta scale in per mil (‰) relative to the internationally accepted standard AIR. Where possible each sample was run in triplicate, and at least in duplicate with $\delta^{15}N$ analytical errors of ±0.2‰ based on repeated measurements of calibration standards.

Isotopic analysis at the Godwin Laboratory, University of Cambridge was undertaken using an automated elemental analyser (Costech Analytical, Valencia, CA, USA) coupled in continuous-flow mode to a Thermo Finnigan MAT253 isotope ratio mass spectrometer (Thermo Fisher Scientific, Bremen, Germany). Between 0.6 and 1 mg of collagen was loaded into a tin capsule for isotopic analysis. International (IAEA: caffeine and glutamic acid-USGS-40) standards with known isotopic values and in-house laboratory standards (nylon, alanine and bovine liver standards) calibrated to IAEA standards were interspersed throughout each analytical run. Results are reported using the delta scale in per mil (‰) relative to the internationally accepted standard AIR. Samples were analysed in duplicate with $\delta^{15}N$ analytical errors of ±0.2‰ based on repeated measurements of calibration standards.

In some instances, samples with previously published $\delta^{15}N$ values have subsequently been re-analysed, and the data from the latest analysis is reported here. For example, some samples with previously published $\delta^{15}N$ values from the RLAHA laboratory (3,6), have subsequently been reanalysed at the SUERC laboratory as part of ongoing research focusing on the simultaneous analyses of $\delta^{15}N$, $\delta^{13}C$, and $\delta^{34}S$. Duplicate data is not included in our data set and the Supplementary Data File clearly specifies which laboratory the reported data comes from. To ensure accurate data curation and archiving, we give previously published samples codes alongside our new analyses in the S1 Dataset.

Radiocarbon dating was performed at the Oxford Radiocarbon Accelerator Unit (ORAU) using their standard procedures (1). Approximately 5 mg of dry collagen per sample was weighed into pre-baked tin capsules and combusted using an elemental analyser coupled to an isotope ratio mass spectrometer, employing a splitter to allow for collection of the $CO_2$ (1,7). Samples were graphitised by reduction of collected $CO_2$ over an iron catalyst in an excess $H_2$ atmosphere at 560 °C (8,9). The $^{14}C$ dates were measured on the Oxford AMS system using a caesium ion source for ionisation of the solid graphite sample(10). To denote samples where collagen extraction took place at a laboratory other than ORAU, all measured dates were given “OxA-V-wwww-pp” numbers, where “wwww” indicates the wheel number, and “pp” is the position of the sample on the wheel (1). For samples where collagen was extracted at UCL, following the method outlined by Wood et al (11), background corrections were applied to our dates to account for inter-laboratory differences in background carbon. A full description of our correction methodology is detailed in Reade et al (12) . Corrected dates are denoted by the “C” at the end of the date code assigned by ORAU. All dates are given as uncalibrated radiocarbon dates ($^{14}C$ BP) and calibrated dates BP (cal. BP) in the Supplementary Data file. Date calibration was performed using OxCal 4.4 (13) and the IntCal20 dataset (14)

### 2 Inter-species comparisons

While herbivore $\delta^{15}N$ tracks that of the environmental baseline $\delta^{15}N$, differences in dietary behaviours between different species (and between different populations or individuals of the same species) introduce variation into the $\delta^{15}N$ signal. Herbivore feeding habits typically fall on a spectrum between graze-dominated and browse-dominated diets; the position of a species on this continuum is partly determined by dietary physiology and partly determined by environmental conditions and inter-species competition in a given location/time context. Broadly, higher $\delta^{15}N$ is associated with grazers and lower $\delta^{15}N$ with browsers within a given ecosystem, although this pattern is not consistent through space or time (15–17). Further variation may occur in herbivore $\delta^{15}N$ related to the consumption of different plant parts and different plant species within the graze (typically grasses, sedges, forbs) or browse (typically shrubs and tree foliage) groupings (18). Additionally, the proportion of different plant species consumed may not be represented in the bone collagen $\delta^{15}N$ in directly equivalent proportions. As bone collagen $\delta^{15}N$ predominately represents the $\delta^{15}N$ of dietary proteins, for a species consuming a mix of protein-rich and protein-poor plant types, it is the protein-rich species that will be greater represented in the bone collagen $\delta^{15}N$ signature (16).

The assembled data were evaluated for potential species-based effects related to diet, habitat preference, and ecology on $\delta^{15}N$. Each taxon was assigned to a dietary category (either grazer, browser, or mixed-feeder) based on prevailing understanding of dietary behaviour (see Schwartz-Narbonne et al (16) and references therein for detailed discussion). In summary, *Equus*, *Bos/Bison, Mammuthus* and *Coelodonta* were categorised as grazers, *Rangifer, Cervus elaphus*, *Megaloceros, Saiga, Ovibos* and *Rupicapra* were categorised as mixed feeders, and *Alces, Capra* and *Capreolus* were categorised as browsers. In making these categorisations the dietary behaviour that is considered most dominant, or most commonly evident in extant species, was selected. However, most species display considerable ecological flexibility, and dietary behaviours are strongly influenced by environmental conditions and inter-species competition (16,17).

Data was divided by time bin and $\delta^{15}N$ was compared between species and dietary category (Fig S2.1). Kruskal-Wallis tests indicate significant differences in $\delta^{15}N$ between species and in $\delta^{15}N$ between dietary categories (Table S2.1). This is not unexpected as the comparison makes no consideration of the spatial distribution of different species and thus data spanning a range of climatic and environmental zones are being compared.

**Table S2.1 Kruskall-Wallis test statistics comparing $\delta^{15}N$ between species, and $\delta^{15}N$ between dietary categories, for each time bin.**

| Time bin | Species Comparison | | Dietary Comparison | |
|---|---|---|---|---|
| | Test statistic | *p* | Test statistic | *p* |
| **EH** | 30.9 | <0.000 | 11.8 | 0.003 |
| **YD** | 11.3 | 0.003 | 12.5 | <0.000 |
| **LGI** | 88.0 | <0.000 | 22.2 | <0.000 |
| **LGT** | 274.0 | <0.000 | 77.3 | <0.000 |
| **LGM** | 57.3 | <0.000 | 5.91 | 0.052 |
| **LOIS3** | 25.6 | <0.000 | 17.0 | <0.000 |
| **EOIS3** | 49.9 | <0.000 | 20.5 | <0.000 |

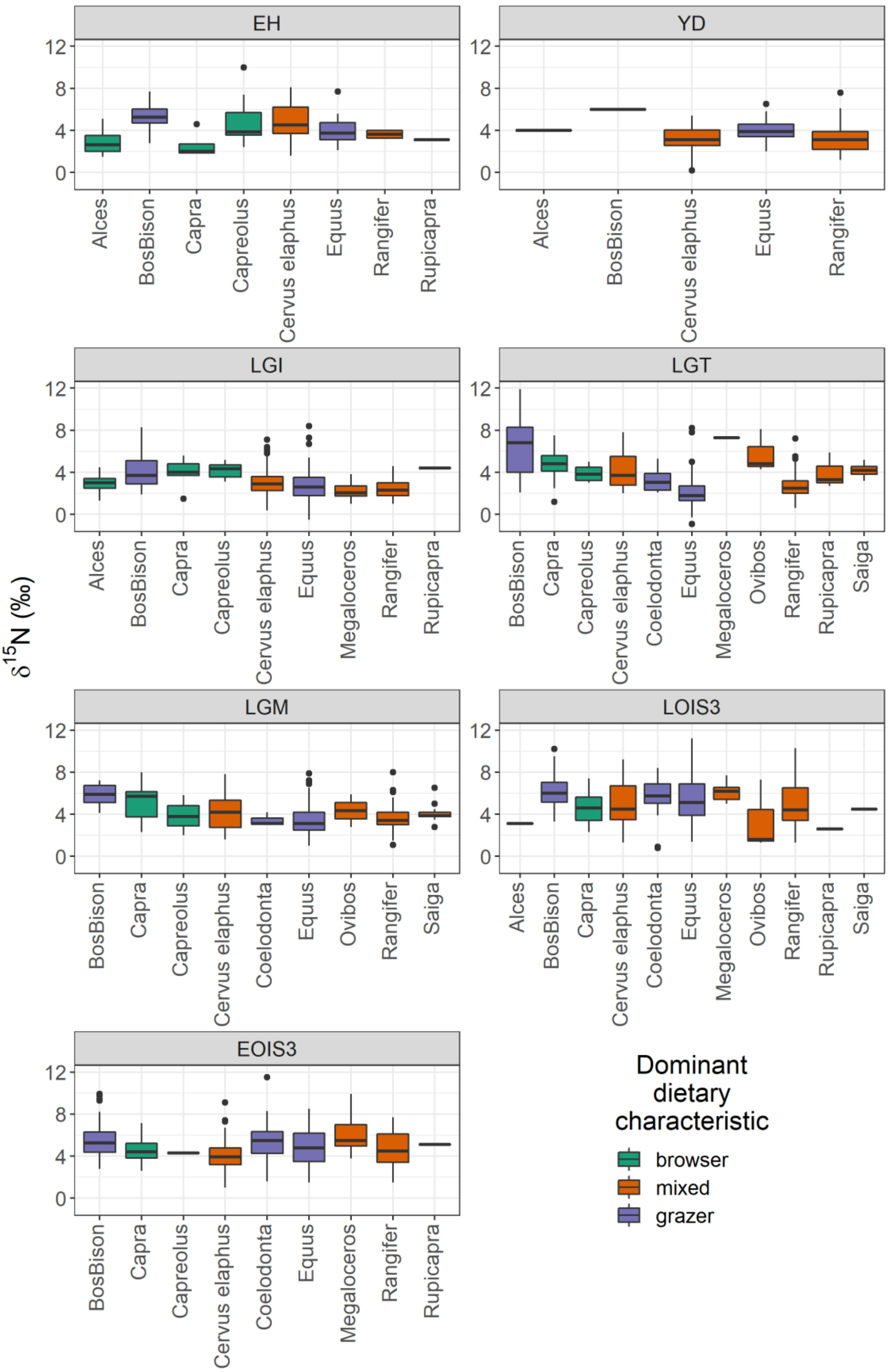


158

159 **Figure S2.1 Boxplots showing the range of $\delta^{15}N$ values for each species, divided by time**

160 **bin, and coloured by dominant dietary characteristic**

Differences between species can be more robustly examined by considering $\delta^{15}N$ variability within spatiotemporal clusters. Samples within the same time bin which originated from locations within 100km of one another were grouped together in clusters. Inter-species differences were then investigated for each spatiotemporal cluster where at least 2 different species were present, each with at least 3 data points. Within-cluster $\delta^{15}N$ was compared between species using Mann Whitney U tests where the number of species was 2 and using Kruskal-Wallis tests where the number of species were greater than 2. Of the 65 spatiotemporal clusters evaluated for species-based differences, 31 showed significant differences in $\delta^{15}N$ between species and 34 did not (Table S2.2), and no consistent pattern in space or time is apparent (Fig S2.2).

**Table S2.2 Summary and test statistics for comparisons of $\delta^{15}N$ between species by spatiotemporal clusters. Tests for difference were and Mann-Whitney (MW) where number of species = 2 and Kruskall-Wallis (KW) where n ≥ 3. Significance was taken to be *p* < 0.05 and significant differences are indicated with ***

| Early Holocene | | | | | | | | | | |
|---|---|---|---|---|---|---|---|---|---|---|
| Cluster | Faunal Category | n | $\delta^{15}N$ (‰) Mean | Median | s.d. | Min | Max | Test Statistic | P value | Test |
| 1 | BosBison | 3 | 6 | 6 | 0.1 | 6 | 6.1 | 12.0 | 0.050* | MW |
| | Capreolus | 4 | 3.4 | 3.5 | 0.4 | 2.9 | 3.8 | | | |
| 2 | BosBison | 9 | 5.2 | 5.2 | 0.6 | 4.7 | 6.6 | 18.0 | 0.456 | MW |
| | Equus | 3 | 4.2 | 2.7 | 3.1 | 2.1 | 7.7 | | | |
| 5 | Capra | 3 | 1.9 | 1.9 | 0.2 | 1.8 | 2.1 | 0.0 | 0.050* | MW |
| | Cervus elaphus | 4 | 3.8 | 4 | 0.4 | 3.2 | 4.1 | | | |
| 7 | BosBison | 5 | 5.4 | 5.2 | 0.5 | 4.9 | 6 | 2.5 | 0.29 | KW |

| | | | | | | | | | | |
|---|---|---|---|---|---|---|---|---|---|---|
| | Capreolus | 7 | 5.8 | 5.5 | 2.1 | 3.7 | 10 | | | |
| | Cervus elaphus | 5 | 6.3 | 6.8 | 1.5 | 3.7 | 7.4 | | | |
| 18 | BosBison | 3 | 5.5 | 5.3 | 0.4 | 5.2 | 5.9 | 24.0 | 0.012* | MW |
| | Cervus elaphus | 8 | 4.2 | 4.2 | 0.6 | 3.4 | 5 | | | |

Younger Dryas

| Cluster | Faunal Category | n | δ¹⁵N (‰) | | | | | Test Statistic | P value | Test |
|---|---|---|---|---|---|---|---|---|---|---|
| | | | **Mean** | **Median** | **s.d.** | **Min** | **Max** | | | |
| 1 | Equus | 12 | 4.4 | 4.4 | 0.7 | 3.6 | 5.8 | 128.0 | 0.583 | MW |
| | Rangifer | 19 | 4.3 | 4.1 | 1.3 | 1.8 | 7.6 | | | |
| 2 | Equus | 4 | 3.4 | 3.5 | 0.5 | 2.8 | 3.8 | 10.5 | 0.154 | MW |
| | Rangifer | 3 | 2.7 | 2.8 | 0.5 | 2.2 | 3.1 | | | |
| 3 | Equus | 3 | 3.7 | 4.3 | 1.1 | 2.5 | 4.4 | 38.0 | 0.073 | MW |
| | Rangifer | 15 | 2.5 | 2.4 | 0.9 | 1.3 | 5 | | | |
| 5 | Equus | 4 | 3.3 | 3.2 | 0.5 | 2.9 | 3.9 | 38.5 | 0.011* | MW |
| | Rangifer | 10 | 2 | 1.9 | 0.6 | 1.2 | 3 | | | |

Late Glacial Interstadial

| Cluster | Faunal Category | n | δ¹⁵N (‰) | | | | | Test Statistic | P value | Test |
|---|---|---|---|---|---|---|---|---|---|---|
| | | | **Mean** | **Median** | **s.d.** | **Min** | **Max** | | | |
| 1 | BosBison | 3 | 5.5 | 5.7 | 0.6 | 4.9 | 6 | 20.2 | <0.000* | KW |
| | Capreolus | 7 | 4.6 | 4.8 | 0.6 | 3.2 | 5.2 | | | |
| | Cervus elaphus | 11 | 4.5 | 4.5 | 0.9 | 2.9 | 5.5 | | | |
| | Equus | 7 | 3.8 | 3.6 | 0.5 | 3 | 4.4 | | | |
| | Rangifer | 19 | 3.5 | 3.4 | 0.7 | 2.3 | 4.6 | | | |
| 2 | Alces | 14 | 2.8 | 2.7 | 0.6 | 1.3 | 3.6 | 2.6 | 0.45 | KW |
| | BosBison | 13 | 3.1 | 2.9 | 0.6 | 2.1 | 4.2 | | | |
| | Cervus elaphus | 29 | 3.1 | 3.3 | 0.7 | 1.7 | 4.4 | | | |
| | Equus | 3 | 3.2 | 2.5 | 1.6 | 2.1 | 5.1 | | | |
| 4 | BosBison | 15 | 3.5 | 3.6 | 0.6 | 2.6 | 4.6 | 41.0 | 0.033* | MW |

| | | | | | | | | | | |
|---|---|---|---|---|---|---|---|---|---|---|
| | Equus | 3 | 1.8 | 1.8 | 1.4 | 0.4 | 3.1 | | | |
| 6 | Cervus elaphus | 6 | 3.4 | 3.5 | 0.4 | 2.8 | 4 | 0.9 | 0.623 | KW |
| | Equus | 11 | 3.3 | 3.2 | 1 | 2 | 5.4 | | | |
| | Rangifer | 8 | 3.1 | 3.1 | 0.5 | 2.3 | 3.9 | | | |
| 7 | Capra | 5 | 4.5 | 4.2 | 0.8 | 3.7 | 5.6 | 0.5 | 0.051 | MW |
| | Cervus elaphus | 3 | 5.8 | 5.8 | 0.2 | 5.6 | 6 | | | |
| 10 | Alces | 5 | 3.8 | 4 | 0.7 | 2.8 | 4.5 | 5.4 | 0.144 | KW |
| | BosBison | 11 | 4.7 | 4.9 | 1 | 2.4 | 5.8 | | | |
| | Cervus elaphus | 7 | 4 | 4.2 | 0.6 | 3.2 | 4.7 | | | |
| | Equus | 10 | 4.1 | 4.2 | 1.1 | 2.5 | 5.3 | | | |
| 12 | Equus | 4 | 2.2 | 2.4 | 0.6 | 1.5 | 2.7 | 53.5 | 0.221 | MW |
| | Rangifer | 19 | 1.8 | 1.8 | 0.5 | 1 | 3.2 | | | |
| 14 | Equus | 19 | 2.9 | 3 | 0.6 | 1.8 | 4 | 179.0 | 0.096 | MW |
| | Rangifer | 14 | 2.6 | 2.6 | 0.4 | 1.6 | 3.1 | | | |
| 15 | BosBison | 4 | 7.1 | 7.1 | 1 | 6 | 8 | 8.0 | 1 | MW |
| | Equus | 4 | 6.8 | 7 | 1.5 | 4.9 | 8.4 | | | |
| 19 | Cervus elaphus | 26 | 2.4 | 2.3 | 0.5 | 1.7 | 4.2 | 176.5 | <0.000* | MW |
| | Equus | 7 | 1.4 | 1.4 | 0.4 | 0.8 | 1.9 | | | |
| 20 | Capreolus | 8 | 4 | 4 | 0.5 | 3.2 | 4.6 | 36.5 | 0.019* | MW |
| | Cervus elaphus | 5 | 3 | 3.1 | 0.5 | 2.2 | 3.6 | | | |
| 22 | BosBison | 3 | 3 | 2.8 | 0.3 | 2.8 | 3.4 | 7.6 | 0.022* | KW |
| | Cervus elaphus | 3 | 1.8 | 1.8 | 0.5 | 1.3 | 2.2 | | | |
| | Rangifer | 13 | 1.6 | 1.5 | 0.4 | 1.3 | 2.5 | | | |
| 23 | Cervus elaphus | 17 | 2 | 2 | 0.7 | 0.4 | 3.1 | 50.0 | 0.567 | MW |
| | Rangifer | 7 | 2.2 | 2.3 | 0.5 | 1.3 | 2.8 | | | |
| 29 | Equus | 11 | 2.5 | 2.5 | 0.8 | 1 | 3.9 | 22.0 | 0.436 | MW |
| | Rangifer | 3 | 2.2 | 2.2 | 0.4 | 1.8 | 2.6 | | | |
| Last Glacial Termination | | | | | | | | | | |

| Cluster | Faunal Category | n | δ15N (‰) | | | | | Test Statistic | P value | Test |
|---|---|---|---|---|---|---|---|---|---|---|
| | | | Mean | Median | s.d. | Min | Max | | | |
| 1 | BosBison | 3 | 2.6 | 2.8 | 0.4 | 2.2 | 2.9 | 5.8 | 0.121 | KW |
| | Coelodonta | 3 | 2.7 | 2.9 | 0.6 | 2.1 | 3.2 | | | |
| | Equus | 75 | 2.1 | 1.9 | 0.9 | -0.9 | 3.7 | | | |
| | Rangifer | 32 | 1.9 | 1.9 | 0.6 | 0.6 | 3 | | | |
| 2 | BosBison | 4 | 3.4 | 2.8 | 1.7 | 2.1 | 5.7 | 7.1 | 0.029* | KW |
| | Equus | 13 | 1.6 | 1.6 | 0.6 | 0.8 | 2.4 | | | |
| | Rangifer | 14 | 2.2 | 2.1 | 1 | 1.1 | 4.3 | | | |
| 4 | Equus | 5 | 1.5 | 1.6 | 0.6 | 0.6 | 2.3 | 18.0 | 0.014* | MW |
| | Rangifer | 25 | 2.3 | 2.4 | 0.5 | 1.2 | 3.4 | | | |
| 6 | Capra | 28 | 5.3 | 5 | 1.1 | 3.9 | 7.5 | 13.6 | 0.001* | KW |
| | Cervus elaphus | 42 | 5.5 | 5.5 | 0.7 | 3.9 | 7.2 | | | |
| | Equus | 4 | 3.4 | 3.4 | 0.3 | 3.1 | 3.7 | | | |
| 7 | Equus | 4 | 3.7 | 3.7 | 0.9 | 2.7 | 4.6 | 5.8 | 0.056 | KW |
| | Rangifer | 9 | 3.4 | 3.3 | 0.6 | 2.5 | 4.2 | | | |
| | Saiga | 3 | 4.7 | 4.5 | 0.5 | 4.3 | 5.2 | | | |
| 8 | Equus | 8 | 1.7 | 1.8 | 0.3 | 1.2 | 2 | 0.0 | 0.008* | MW |
| | Rangifer | 4 | 2.6 | 2.5 | 0.4 | 2.2 | 3.1 | | | |
| 9 | Equus | 13 | 1.7 | 1.5 | 0.5 | 1.1 | 2.9 | 3.0 | 0.010* | MW |
| | Rangifer | 4 | 2.9 | 2.8 | 0.3 | 2.6 | 3.2 | | | |
| 10 | BosBison | 5 | 6 | 5.8 | 0.5 | 5.4 | 6.8 | 19.1 | 0.000* | KW |
| | Equus | 5 | 4.2 | 4.2 | 0.5 | 3.4 | 4.8 | | | |
| | Rangifer | 10 | 3.5 | 3.6 | 0.6 | 2.4 | 4.6 | | | |
| | Saiga | 19 | 4.2 | 4.1 | 0.5 | 3.2 | 5.1 | | | |
| 11 | BosBison | 24 | 8.2 | 8.1 | 1.4 | 6.1 | 11.9 | 311.0 | <0.000* | MW |
| | Cervus elaphus | 15 | 6.5 | 6.5 | 0.8 | 5 | 7.8 | | | |
| 13 | Cervus elaphus | 3 | 2.3 | 2.3 | 0.2 | 2.1 | 2.4 | 54.0 | 0.031* | MW |

| Cluster | Faunal Category | n | δ15N (‰) Mean | Median | s.d. | Min | Max | Test Statistic | P value | Test |
|---|---|---|---|---|---|---|---|---|---|---|
| | Equus | 20 | 1.2 | 1.1 | 0.8 | -0.3 | 3.2 | | | |
| 18 | Equus | 17 | 2.8 | 2.1 | 1.7 | 1.3 | 8.2 | 6.6 | 0.037* | KW |
| | Ovibos | 3 | 5.7 | 4.8 | 2.1 | 4.3 | 8.1 | | | |
| | Rangifer | 10 | 2.5 | 2.5 | 0.8 | 1.1 | 4.1 | | | |

Last Glacial Maximum

| Cluster | Faunal Category | n | $\delta^{15}N$ (‰) | | | | | Test Statistic | P value | Test |
|---|---|---|---|---|---|---|---|---|---|---|
| | | | Mean | Median | s.d. | Min | Max | | | |
| 2 | BosBison | 6 | 4.6 | 4.5 | 0.4 | 4.1 | 5.2 | 14.5 | 0.002* | MW |
| | Equus | 20 | 3.2 | 3.2 | 1.1 | 1 | 6 | | | |
| | Rangifer | 48 | 3.4 | 3.3 | 0.8 | 1.5 | 5.2 | | | |
| | Saiga | 10 | 3.9 | 3.9 | 0.6 | 2.8 | 5 | | | |
| 4 | Cervus elaphus | 46 | 3.2 | 2.8 | 1.3 | 1.6 | 7.8 | 15.5 | <0.000* | MW |
| | Rangifer | 6 | 5.7 | 5.4 | 1.2 | 4.7 | 8 | | | |
| 5 | Capra | 12 | 5.8 | 5.8 | 1.2 | 3.7 | 8 | 261.5 | 0.213 | MW |
| | Cervus elaphus | 35 | 5.3 | 5.3 | 1.1 | 3.1 | 7.2 | | | |
| 6 | Cervus elaphus | 7 | 4.8 | 4.6 | 0.5 | 4 | 5.7 | 35.0 | 0.005* | MW |
| | Equus | 5 | 3 | 3.2 | 0.3 | 2.6 | 3.3 | | | |
| 7 | BosBison | 3 | 6.9 | 6.8 | 0.2 | 6.8 | 7.1 | 0.3 | 0.843 | KW |
| | Cervus elaphus | 6 | 6.4 | 6.7 | 1.1 | 5 | 7.5 | | | |
| | Equus | 7 | 7 | 6.9 | 0.5 | 6.5 | 7.9 | | | |
| 8 | Equus | 4 | 2.7 | 2.7 | 0.9 | 1.8 | 3.7 | 3.5 | 0.14 | MW |
| | Rangifer | 5 | 3.7 | 3.7 | 0.5 | 2.9 | 4.3 | | | |
| 10 | BosBison | 8 | 6 | 6 | 0.6 | 5 | 6.9 | 17.5 | 0.305 | MW |
| | Equus | 3 | 5.5 | 5.2 | 0.5 | 5.2 | 6.1 | | | |
| 12 | Equus | 3 | 3.3 | 2.2 | 1.8 | 2.2 | 5.4 | 5.0 | 1 | MW |
| | Rangifer | 3 | 2.5 | 2.6 | 1.4 | 1.1 | 3.9 | | | |

Late OIS 3

| Cluster | Faunal Category | n | $\delta^{15}N$ (‰) | Test Statistic | P value | Test |
|---|---|---|---|---|---|---|

| | | | Mean | Median | s.d. | Min | Max | | | |
|---|---|---|---|---|---|---|---|---|---|---|
| 1 | BosBison | 4 | 5.4 | 5.0 | 1.5 | 4.3 | 7.5 | 6.4 | 0.094 | KW |
| | Cervus elaphus | 10 | 5.4 | 4.7 | 1.8 | 3.8 | 8.5 | | | |
| | Equus | 34 | 6.5 | 6.5 | 2.3 | 3.0 | 11.2 | | | |
| | Rangifer | 28 | 7.1 | 7.3 | 1.7 | 4.1 | 10.3 | | | |
| 2 | Cervus elaphus | 50 | 4.7 | 4.0 | 1.9 | 1.3 | 9.2 | 807.0 | 0.041* | MW |
| | Equus | 25 | 3.8 | 3.6 | 1.8 | 1.4 | 8.0 | | | |
| 3 | BosBison | 6 | 6.6 | 6.2 | 1.2 | 5.3 | 8.6 | 5.0 | 0.364 | MW |
| | Coelodonta | 3 | 7.5 | 7.1 | 0.8 | 7.1 | 8.4 | | | |
| 4 | BosBison | 5 | 4.6 | 4.2 | 1.5 | 3.3 | 6.8 | 10.7 | 0.005* | KW |
| | Equus | 23 | 5.1 | 5.0 | 1.5 | 2.5 | 9.8 | | | |
| | Rangifer | 24 | 4.0 | 4.0 | 1.1 | 2.6 | 8.0 | | | |
| 5 | Equus | 21 | 6.9 | 6.8 | 1.9 | 4.1 | 9.7 | 106.5 | 0.012* | MW |
| | Rangifer | 6 | 4.4 | 4.6 | 1.0 | 2.6 | 5.8 | | | |
| 6 | BosBison | 5 | 4.5 | 4.1 | 0.8 | 3.8 | 5.7 | 18.6 | <0.000* | KW |
| | Coelodonta | 7 | 5.9 | 5.9 | 1.2 | 4.3 | 7.3 | | | |
| | Equus | 15 | 5.8 | 5.8 | 1.0 | 3.8 | 7.5 | | | |
| | Rangifer | 19 | 3.9 | 3.4 | 1.5 | 2.1 | 8.1 | | | |
| 7 | BosBison | 8 | 5.5 | 5.6 | 0.6 | 4.6 | 6.1 | 19.0 | 0.67 | MW |
| | Rangifer | 4 | 5.3 | 4.9 | 1.8 | 3.7 | 7.6 | | | |
| 9 | Cervus elaphus | 5 | 4.8 | 4.6 | 1.4 | 3.2 | 7.1 | 4.1 | 0.126 | KW |
| | Equus | 7 | 4.8 | 3.9 | 2.1 | 2.1 | 7.9 | | | |
| | Rangifer | 23 | 6.1 | 6.1 | 1.5 | 3.4 | 8.6 | | | |
| 11 | BosBison | 7 | 6.8 | 6.1 | 1.7 | 5.0 | 9.3 | 0.6 | 0.759 | KW |
| | Capra | 3 | 5.9 | 6.0 | 1.5 | 4.4 | 7.3 | | | |
| | Cervus elaphus | 4 | 6.3 | 6.0 | 1.0 | 5.5 | 7.7 | | | |
| 12 | BosBison | 4 | 5.4 | 5.4 | 0.6 | 4.7 | 6.1 | 9.8 | 0.020* | KW |
| | Coelodonta | 3 | 5.3 | 5.1 | 0.9 | 4.6 | 6.3 | | | |

| Cluster | Faunal Category | n | δ15N (‰) Mean | Median | s.d. | Min | Max | Test Statistic | P value | Test |
|---|---|---|---|---|---|---|---|---|---|---|
| | Equus | 4 | 5.7 | 5.3 | 2.1 | 3.6 | 8.6 | | | |
| | Rangifer | 6 | 3.3 | 3.2 | 0.8 | 2.4 | 4.4 | | | |
| 16 | BosBison | 8 | 7.5 | 8.2 | 2.3 | 3.5 | 10.2 | 14.6 | 0.002* | KW |
| | Coelodonta | 3 | 5.4 | 5.4 | 0.1 | 5.3 | 5.5 | | | |
| | Equus | 3 | 4.3 | 3.8 | 1.9 | 2.6 | 6.4 | | | |
| | Rangifer | 8 | 2.6 | 2.8 | 0.8 | 1.3 | 3.6 | | | |

| Early OIS 3 | | | | | | | | | | |
|---|---|---|---|---|---|---|---|---|---|---|
| Cluster | Faunal Category | n | $\delta^{15}N$ (‰) | | | | | Test Statistic | P value | Test |
| | | | Mean | Median | s.d. | Min | Max | | | |
| 1 | Cervus elaphus | 3 | 3.6 | 3.8 | 0.4 | 3.1 | 3.9 | 2.8 | 0.252 | KW |
| | Equus | 3 | 4.4 | 4.8 | 2.0 | 2.2 | 6.2 | | | |
| | Rangifer | 3 | 5.1 | 5.2 | 0.3 | 4.7 | 5.3 | | | |
| 2 | Cervus elaphus | 45 | 3.5 | 3.2 | 1.2 | 1.9 | 9.1 | 348.0 | 0.245 | MW |
| | Equus | 19 | 4.1 | 3.6 | 1.7 | 1.5 | 7.6 | | | |
| 3 | BosBison | 4 | 5.3 | 5.5 | 0.8 | 4.1 | 6.0 | 10.2 | 0.017* | KW |
| | Cervus elaphus | 19 | 5.1 | 5.1 | 1.0 | 3.7 | 7.3 | | | |
| | Equus | 21 | 4.5 | 4.3 | 1.6 | 1.6 | 6.7 | | | |
| | Rangifer | 11 | 6.3 | 6.5 | 1.2 | 3.4 | 7.7 | | | |
| 4 | Cervus elaphus | 5 | 4.2 | 4.4 | 0.9 | 3.0 | 5.3 | 0.3 | 0.865 | KW |
| | Equus | 7 | 4.6 | 4.7 | 1.8 | 2.1 | 6.5 | | | |
| | Rangifer | 5 | 4.2 | 4.6 | 0.7 | 3.3 | 4.8 | | | |
| 5 | Coelodonta | 11 | 6.4 | 6.1 | 2.1 | 3.7 | 11.5 | 13.0 | 0.64 | MW |
| | Equus | 3 | 6.8 | 6.4 | 1.4 | 5.6 | 8.4 | | | |
| 6 | BosBison | 10 | 5.6 | 5.4 | 1.0 | 4.2 | 8.0 | 8.8 | 0.033* | KW |
| | Cervus elaphus | 5 | 4.3 | 4.2 | 1.0 | 3.0 | 5.4 | | | |
| | Equus | 6 | 5.2 | 5.6 | 1.3 | 2.9 | 6.4 | | | |
| | Rangifer | 3 | 6.4 | 6.3 | 0.2 | 6.3 | 6.6 | | | |
| 7 | BosBison | 5 | 7.6 | 8.1 | 1.3 | 6.3 | 9.3 | 8.6 | 0.013* | KW |

| | | | | | | | | | | |
|---|---|---|---|---|---|---|---|---|---|---|
| | Equus | 30 | 5.3 | 5.4 | 1.4 | 2.3 | 8.5 | | | |
| | Rangifer | 30 | 5.1 | 4.8 | 1.4 | 2.6 | 7.6 | | | |
| 8 | BosBison | 3 | 3.2 | 3.3 | 0.4 | 2.8 | 3.6 | 2.9 | 0.575 | KW |
| | Coelodonta | 5 | 3.6 | 4.2 | 1.1 | 2.2 | 4.8 | | | |
| | Equus | 18 | 4.0 | 3.7 | 1.8 | 1.7 | 7.3 | | | |
| | Megaloceros | 3 | 4.4 | 4.6 | 0.5 | 3.8 | 4.7 | | | |
| | Rangifer | 9 | 3.3 | 3.3 | 0.5 | 2.0 | 3.8 | | | |
| 11 | BosBison | 21 | 5.4 | 4.9 | 1.8 | 3.4 | 9.9 | 2.2 | 0.336 | KW |
| | Capra | 7 | 5.0 | 4.6 | 1.3 | 3.8 | 7.1 | | | |
| | Cervus elaphus | 6 | 4.3 | 4.1 | 1.5 | 2.7 | 6.6 | | | |
| 12 | BosBison | 9 | 5.0 | 4.8 | 0.9 | 4.3 | 7.0 | 10.3 | 0.016* | KW |
| | Coelodonta | 8 | 6.2 | 6.0 | 0.9 | 5.3 | 7.5 | | | |
| | Equus | 14 | 5.4 | 5.1 | 1.3 | 3.1 | 7.3 | | | |
| | Rangifer | 6 | 3.8 | 3.5 | 2.1 | 1.5 | 7.7 | | | |
| 13 | Capra | 5 | 5.0 | 4.4 | 1.2 | 3.6 | 6.6 | 4.5 | 0.451 | MW |
| | Cervus elaphus | 3 | 5.7 | 6.0 | 0.8 | 4.8 | 6.2 | | | |
| 17 | Equus | 3 | 4.2 | 4.8 | 1.6 | 2.3 | 5.4 | 7.0 | 1 | MW |
| | Rangifer | 5 | 4.8 | 4.4 | 1.3 | 3.2 | 6.8 | | | |

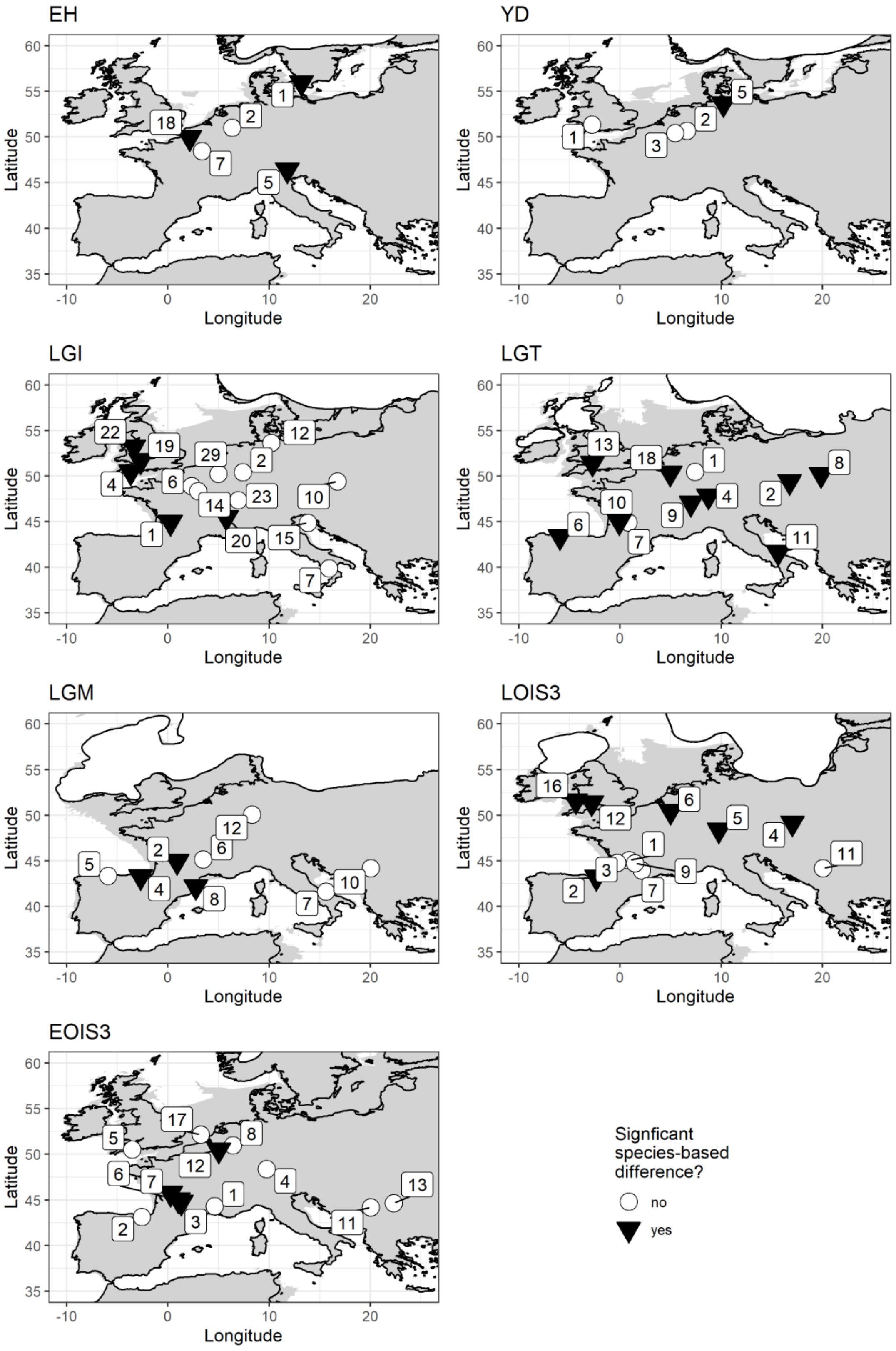


**Fig S2.2 Distribution of clustered samples. Each cluster contains samples from locations within a 100km search radii. Only clusters with at least two different species, with each**

**species containing a minimum of three $\delta^{15}N$ data points, are included. Symbols indicate clusters where $\delta^{15}N$ differs significantly ($p < 0.05$) between species (black triangles) and clusters where it does not (white circles). Cluster numbers correspond to those given in Table S2.2.**

The lack of consistent species-based differences can be most clearly demonstrated by considering horse and reindeer data; these two species comprise 49% of the total data set and both have wide geographical distributions across the study area (the exception being reindeer during the early Holocene). Horse and reindeer are known to have different dietary behaviours and thus would be expected to possess different $\delta^{15}N$ signatures when occupying the same environment; horse are typically considered to be predominantly grazers, while reindeer are considered to be mixed feeders consuming a range of both browse and graze, as well as lichen (19–22). These different plant types have different albeit overlapping and variable $\delta^{15}N$ compositions (16,23).

In our data, horse and reindeer occur together in 35 spatiotemporal clusters. The difference in mean $\delta^{15}N$ between these species within different spatiotemporal clusters ranges from -2.4‰ to +1.7‰ (mean = -0.3 ± 1.0 ‰), and a significant difference is identified in only 10 of the 35 clusters (Fig S4.3). The lack of systematic difference between these two species highlights the challenges faced when attempts are made to quantify and account for species-based differences. Indeed, while horse and reindeer and most commonly referred to as grazers and mixed-feeders respectively, a diversity of dietary behaviours are observed in extant populations of these species and evidenced in fossil assemblages (see discussion in Schwartz-Narbonne 2019). The lack of consistent differences between these two species most likely indicates dietary flexibility, with both species varying their diets relative to the availability of vegetation in their local environment.

Thus, in summary, while differences certainly occur in $\delta^{15}N$ between species and dietary behaviours, we judge that there is too much variation in inter-species differences to enable adequate data normalisation/correction. By avoiding the use of such a correction our data retains a certain degree of noise associated with species/dietary differences, which may increase uncertainties associated with the geostatistical interpolations presented in the main manuscript. However, we believe the approach provides a more faithful representation of average baseline $\delta^{15}N$ values and variability at the landscape scale. Species-specific geostatistical interpolations are considered in the discussion section of the main manuscript.

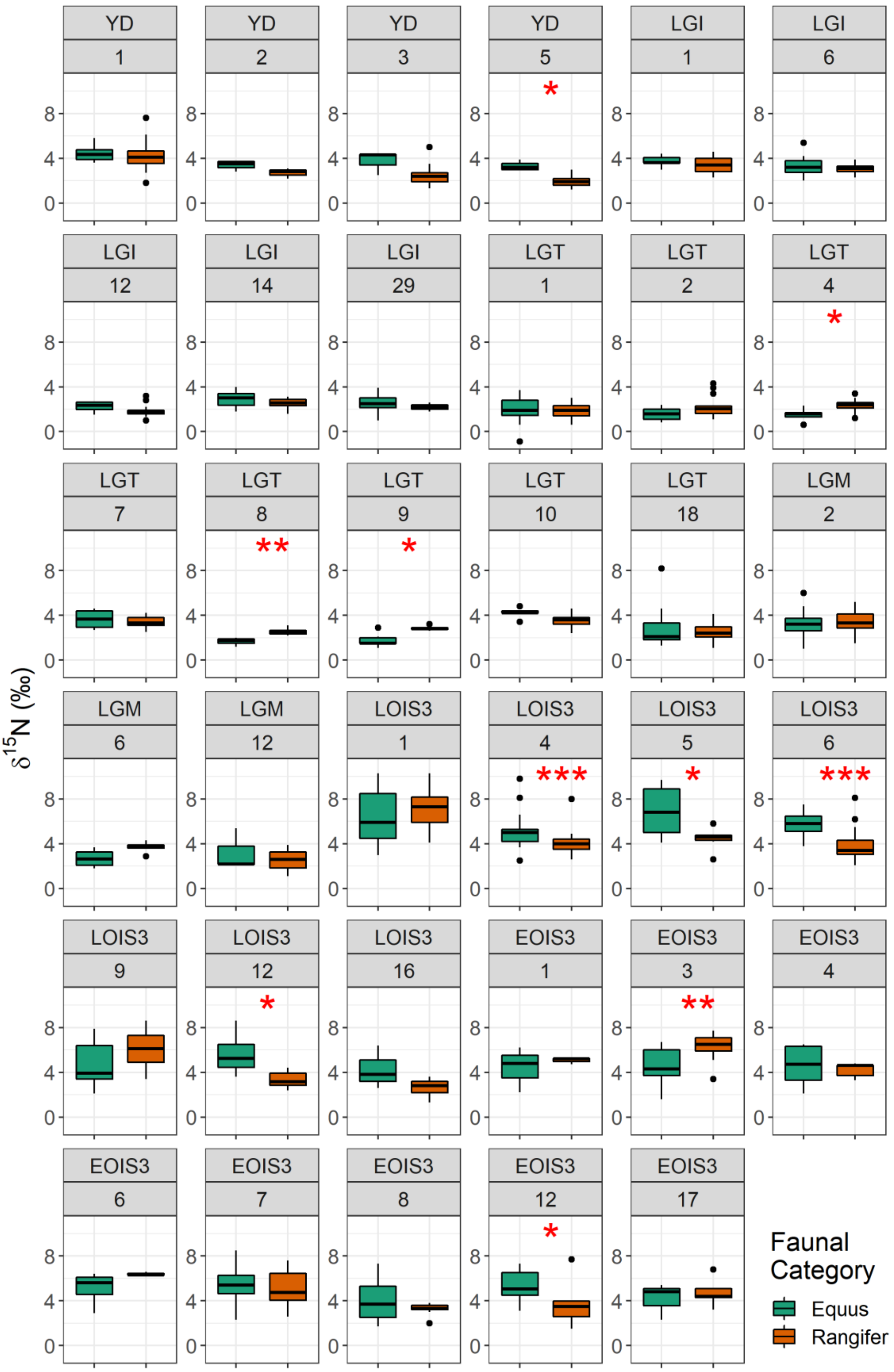


**Fig S2.3 Comparison of horse and reindeer $\delta^{15}$N values by spatiotemporal cluster.**

**Significance indicated by: ***$p$ < .001, ** $p$ < .01, *$p$ < .05.**

### 3 Spatial autocorrelation and outlier analysis

**Table S3.1 Global Moran’s I test statistics and number of spatial outliers identified for each time bin using Anselin’s Local Moran’s I.**

| Time bin | n | Number of spatial outliers | Global Moran’s I | | | |
|---|---|---|---|---|---|---|
| | | | Index | Expected Index | Z score | P value |
| EH | 176 | 2 | 0.308 | -0.006 | 9.5 | <0.000 |
| YD | 133 | 15 | 0.498 | -0.009 | 12.6 | <0.000 |
| LGI | 485 | 18 | 0.515 | -0.002 | 35.9 | <0.000 |
| LGT | 602 | 33 | 0.665 | -0.002 | 55.2 | <0.000 |
| LGM | 339 | 25 | 0.521 | -0.003 | 28.7 | <0.000 |
| LOIS3 | 466 | 33 | 0.305 | -0.002 | 20.2 | <0.000 |
| EOIS3 | 517 | 60 | 0.305 | -0.002 | 21.1 | <0.000 |

### 4 Correlations between faunal $\delta^{15}N$ and bioclimatic variables and covariate selection for geostatistical analyses

The bioclimatic variables from Beyer et al. (24) considered for inclusion as fixed effects in geostatistical analysis are given in Table S4.1. Many of these variables displayed significant correlation with faunal $\delta^{15}N$ (Table S4.2) and a high degree of collinearity with one another (Fig S4.1). In selecting which variables and combinations of variables to consider as fixed effects in the liner mixed-effects models, a number of aspects were considered. First, we considered variables which in the modern environment have demonstrated empirical relationships with $\delta^{15}N$; mean annual temperature (MAT), mean annual precipitation (MAP) and elevation (25–28). Based on investigated correlations (Table S4.2) we retained MAT and MAP and removed elevation, which showed significant correlation with $\delta^{15}N$ for only two time bins. While elevation is an important spatial variable across which $\delta^{15}N$ varies in the modern environment (25,28), there are challenges in estimating palaeo-elevations related to sea-level fluctuations and isostatic responses of the growth and melting of ice sheets. Moreover, our sample set is biased toward environments of <500m elevation, and uncertainties in species-based altitudinal mobility compound uncertainties. Second, we removed variables which provided largely redundant information and displayed high collinearity (Fig S4.1). This included removing precipitation amount of the wettest and driest months, which were highly correlated with precipitation amount of the wettest and driest quarters (correlation coefficients of 0.98 and 0.96, $p < 0.001$); removing minimum and maximum annual temperature which were highly correlated with temperature of the coldest and warmest quarters (correlation coefficients of 0.99 and 0.95, $p < 0.001$); and removing annual temperature range which was highly correlated with temperature seasonality (0.95, $p < 0.001$). Finally, of the remaining variables, those which displayed significant correlations with less than 50% of the time bins were removed. After this, the remaining variables, selected as fixed effects in model testing were: MAT, MAP, temperature of the warmest quarter, precipitation of the warmest quarter and precipitation of the coldest quarter (Fig S4.2 – S4.6).

**Table S4.1. Summary of covariate data considered in this study.**

| Variable | Unit |
|---|---|
| Annual mean temperature | °C |
| Temperature seasonality (standard deviation of monthly temperature) | °C |
| Minimum annual temperature | °C |
| Maximum annual temperature | °C |
| Temperature annual range (difference between minimum and maximum annual temperatures) | °C |
| Mean temperature of the wettest quarter | °C |
| Mean temperature of driest quarter | °C |
| Mean temperature of warmest quarter | °C |
| Mean temperature of coldest quarter | °C |
| Annual precipitation | mm $year^{-1}$ |
| Precipitation of wettest month | mm $month^{-1}$ |
| Precipitation of driest month | mm $month^{-1}$ |
| Precipitation seasonality (coefficient of variation of monthly precipitation) | - |
| Precipitation of wettest quarter | mm $quarter^{-1}$ |
| Precipitation of driest quarter | mm $quarter^{-1}$ |
| Precipitation of warmest quarter | mm $quarter^{-1}$ |
| Precipitation of coldest quarter | mm $quarter^{-1}$ |

**Table S4.2. Pearson's correlation coefficient indicating correlation between spatial and bioclimatic data and d15N values. Significance indicated by: *******p* < .0001, ****p* < .001, ** *p* < .01, **p* < .05.**

| | Test statistics | | | | | | | |
|---|---|---|---|---|---|---|---|---|
| | ALL | EH | YD | LGI | LGT | LGM | LOIS3 | EOIS3 |
| SPATIAL VARIABLES | | | | | | | | |
| Elevation | -0.20**** | -0.21 | -0.37* | -0.19 | -0.27* | -0.07 | 0.12 | -0.05 |
| BIOCLIMATIC VARIABLES | | | | | | | | |
| Annual mean temperature | 0.17*** | 0.18 | 0.12 | 0.71**** | 0.75**** | 0.69**** | 0.27* | 0.17 |
| Temperature seasonality (standard deviation of monthly temperature) | 0.18*** | 0.16 | -0.22 | 0.41*** | -0.16 | -0.27 | -0.1 | -0.07 |
| Minimum annual temperature | 0.09 | 0.09 | 0.34* | 0.47**** | 0.63**** | 0.63**** | 0.16 | 0.14 |
| Maximum annual temperature | 0.25**** | 0.32** | -0.21 | 0.69**** | 0.55**** | 0.60**** | 0.37** | 0.17 |
| Temperature annual range (difference between minimum and maximum annual temperatures) | 0.16*** | 0.16 | -0.28 | 0.37*** | -0.07 | -0.41* | 0 | -0.09 |
| Mean temperature of the wettest quarter | 0.02 | 0.22 | -0.28 | 0.18 | -0.37** | -0.22 | -0.14 | -0.01 |
| Mean temperature of driest quarter | 0.17*** | 0.04 | 0.32 | 0.47**** | 0.64**** | 0.61**** | 0.11 | 0.06 |
| Mean temperature of warmest quarter | 0.29**** | 0.33** | -0.12 | 0.75**** | 0.64**** | 0.69**** | 0.32* | 0.18 |
| Mean temperature of coldest quarter | 0.08 | 0.05 | 0.29 | 0.48**** | 0.65**** | 0.63**** | 0.2 | 0.12 |
| Mean annual precipitation | -0.24**** | -0.41*** | 0.17 | -0.31** | -0.08 | 0.12 | -0.46*** | -0.07 |

| | | | | | | | | |
|---|---|---|---|---|---|---|---|---|
| Precipitation of wettest month | -0.19**** | -0.36** | 0.26 | -0.13 | -0.03 | 0.15 | -0.49**** | -0.12 |
| Precipitation of driest month | -0.24**** | -0.50**** | 0.18 | -0.48**** | -0.15 | 0.04 | -0.36** | -0.01 |
| Precipitation seasonality (coefficient of variation of monthly precipitation) | 0.02 | 0.11 | 0.41* | 0.54**** | -0.24* | -0.11 | 0 | -0.12 |
| Precipitation of wettest quarter | -0.19**** | -0.36** | 0.26 | -0.22 | -0.05 | 0.14 | -0.47**** | -0.08 |
| Precipitation of driest quarter | -0.23**** | -0.47**** | 0.1 | -0.39*** | -0.1 | -0.01 | -0.32* | -0.01 |
| Precipitation of warmest quarter | -0.31**** | -0.36** | -0.05 | -0.44**** | - 0.48**** | -0.51** | -0.38** | -0.06 |
| Precipitation of coldest quarter | -0.13** | -0.38** | 0.3 | -0.24* | 0.19 | 0.42* | -0.36** | 0 |

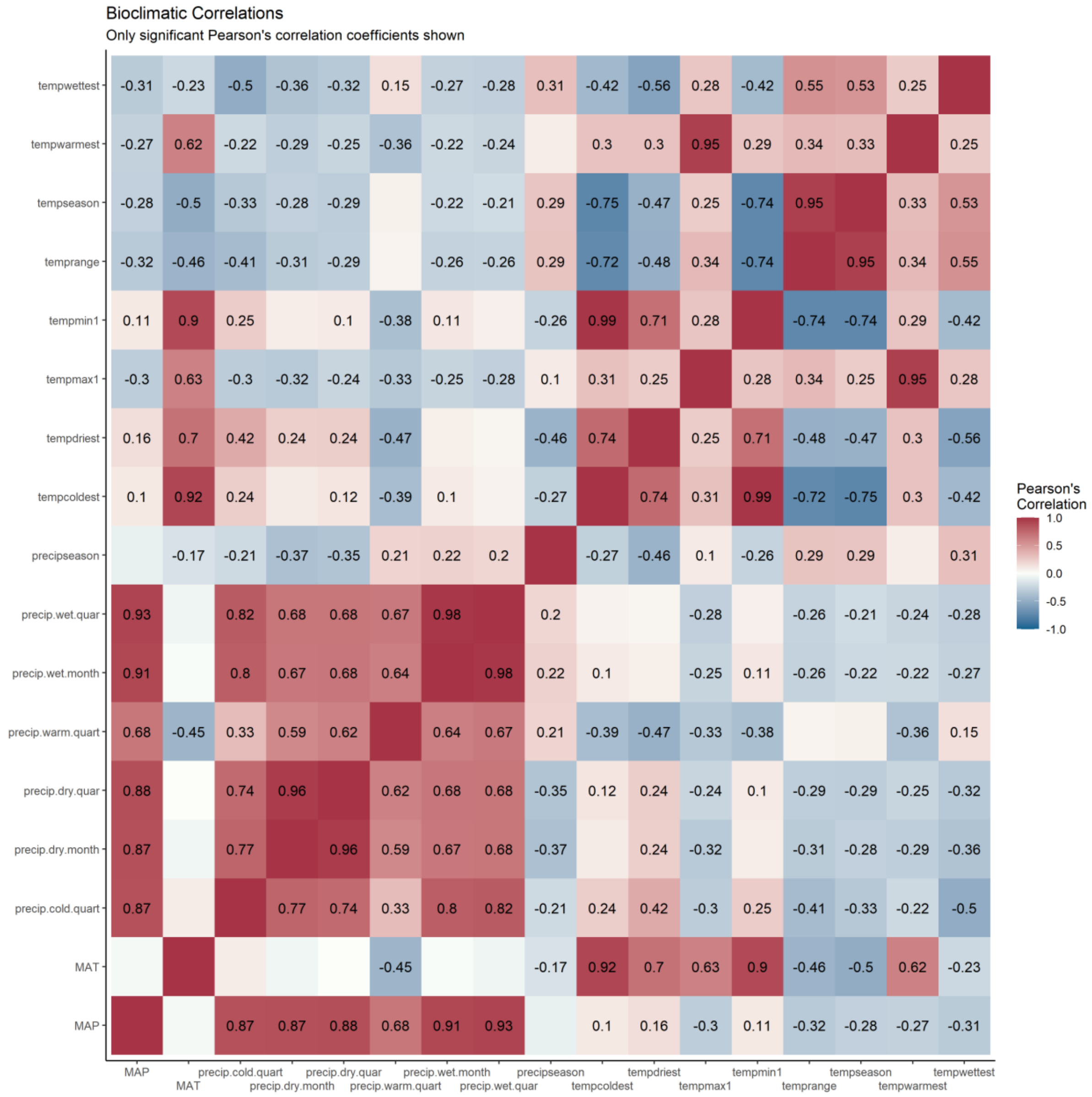

**Figure S4.1 Correlation matrix of bioclimatic variables from Beyer et al. (2020). Pearson's correlation test statistics is displayed only where correlation is significant at p > 0.05.**

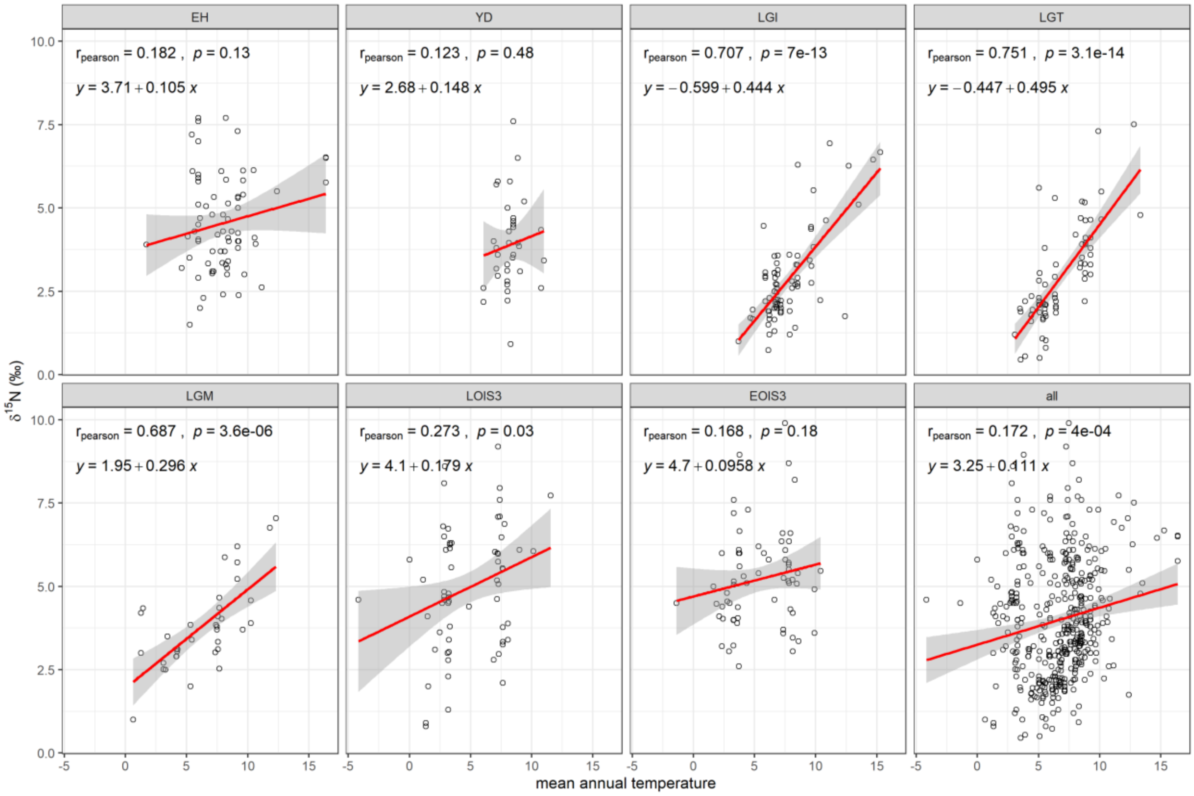


**Figure S4.2 Relationship between site mean faunal δ$^{15}$N and mean annual temperature as derived from the bioclimatic model outputs of Beyer et al. (2020).**

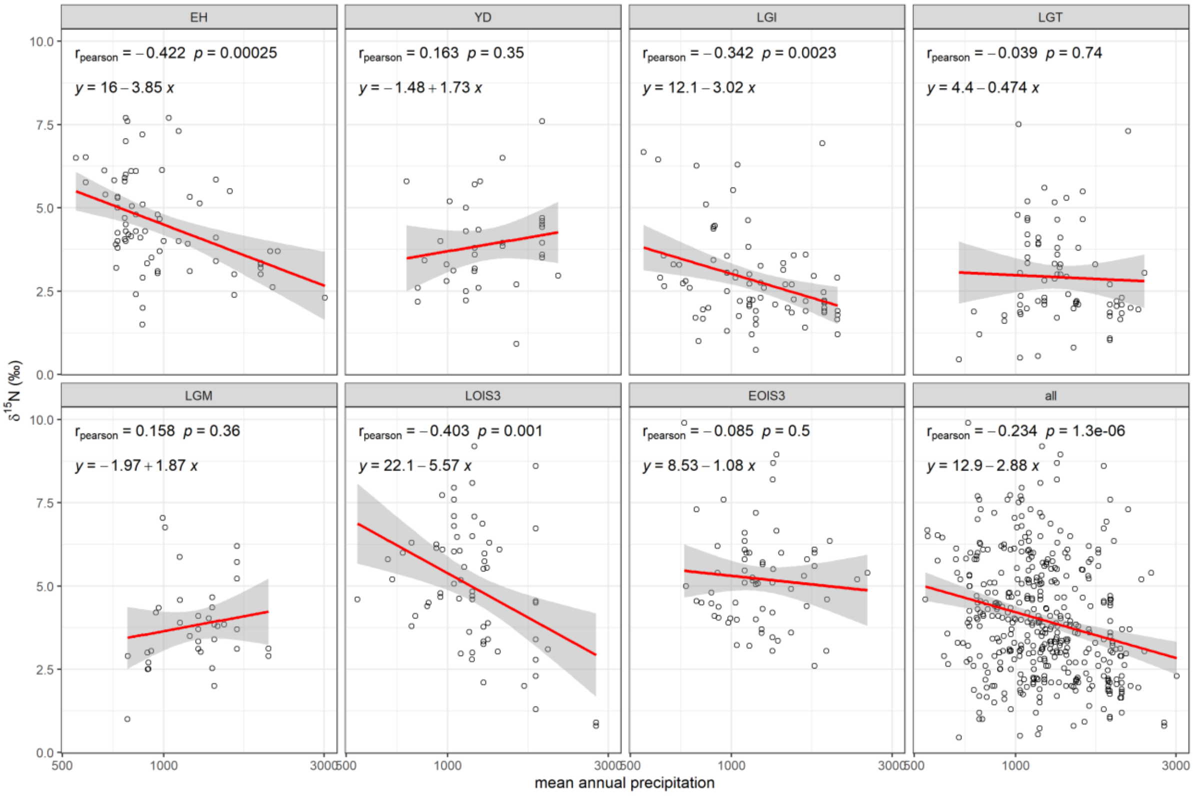


**Figure S4.3 Relationship between site mean faunal $\delta^{15}N$ and mean annual precipitation as derived from the bioclimatic model outputs of Beyer et al. (2020).**

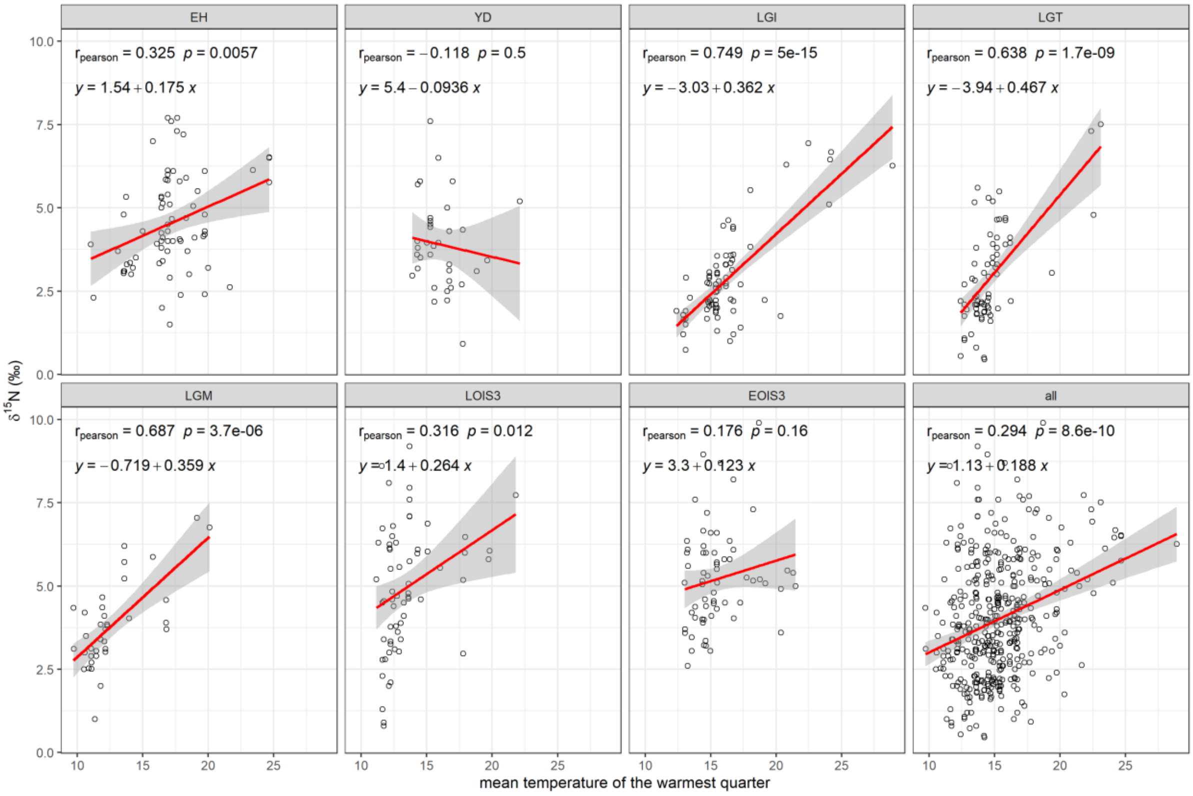


**Figure S4.4 Relationship between site mean faunal δ$^{15}$N and mean temperature of the warmest quarter as derived from the bioclimatic model outputs of Beyer et al. (2020).**

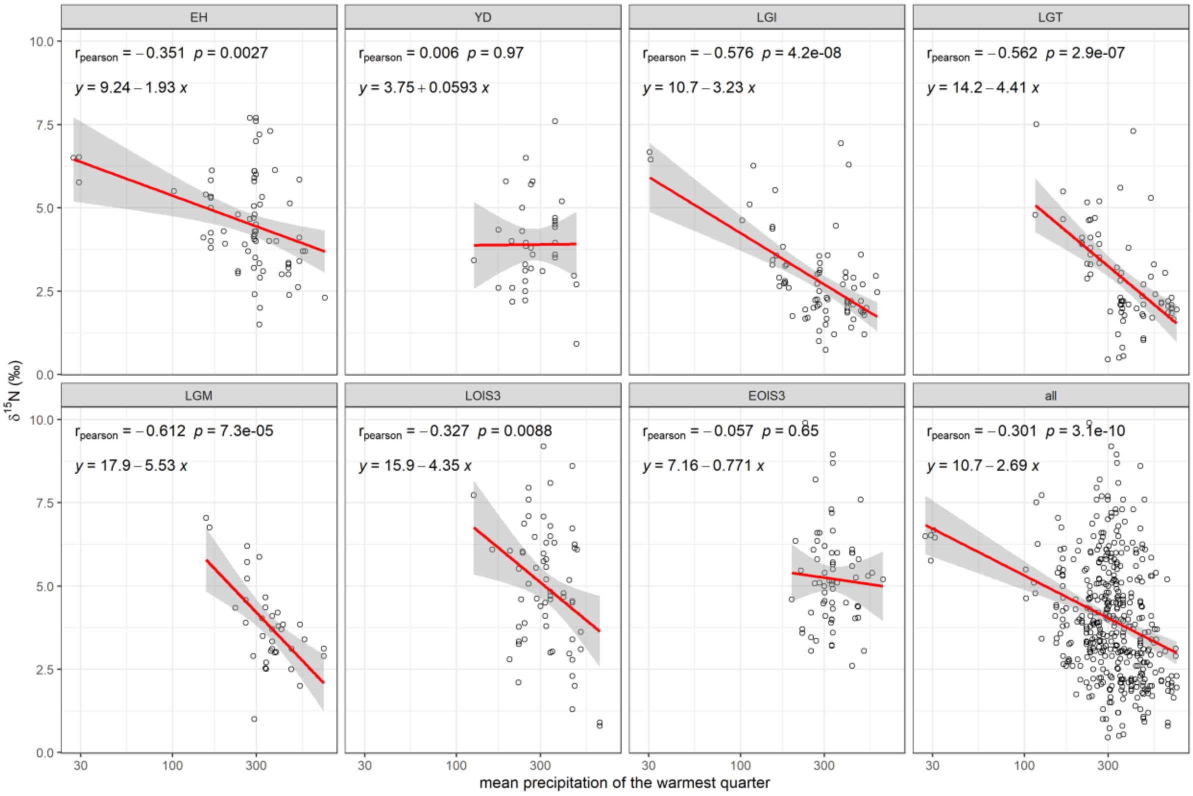


**Figure S4.5 Relationship between site mean faunal $\delta^{15}N$ and mean precipitation of the warmest quarter as derived from the bioclimatic model outputs of Beyer et al. (2020).**

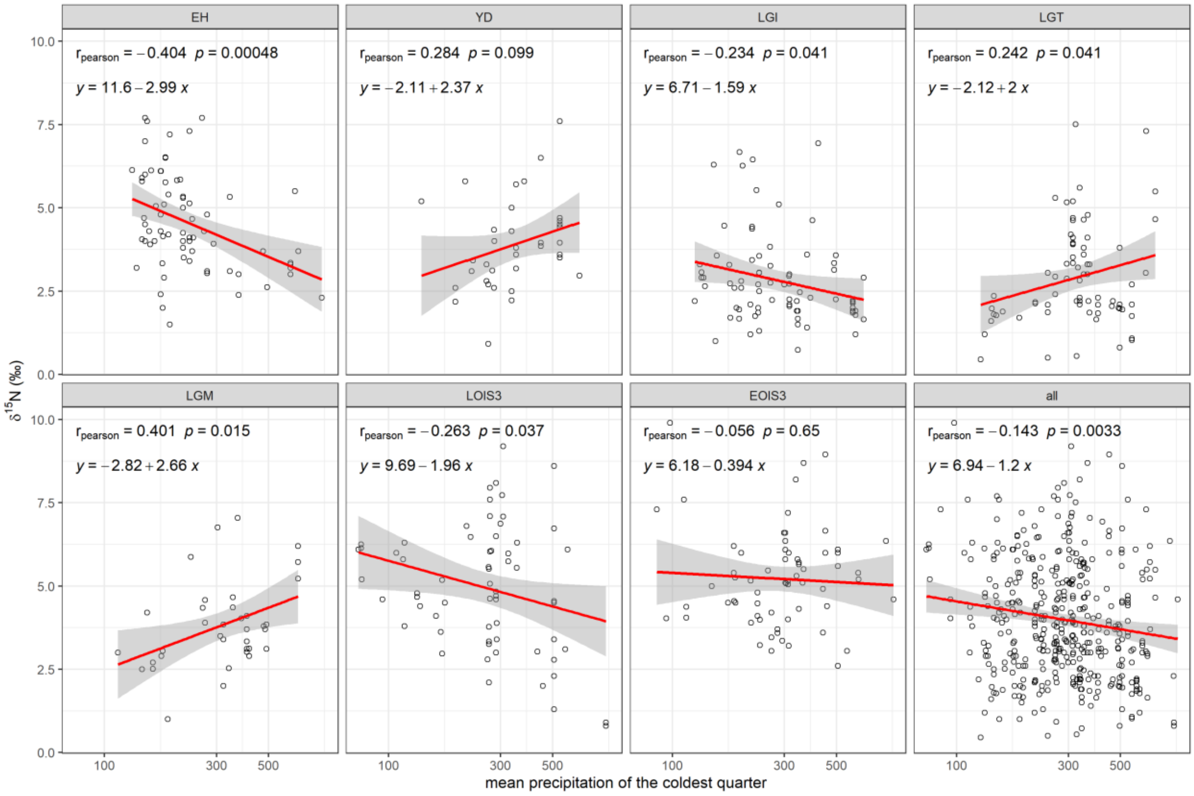


**Figure S4.6 Relationship between site mean faunal $\delta^{15}N$ and mean precipitation of the coldest quarter as derived from the bioclimatic model outputs of Beyer et al. (2020).**

### 5 Isoscape prediction model fitting

**Table S5.1 Models tested. Each model tested included/excluded a different combination of random effect, fixed effects, and order interaction terms.**

| Model number | Model |
|---|---|
| 1 | Intercept + spatial + uncorrelated |
| 2 | Intercept + MAT + spatial + uncorrelated |
| 3 | Intercept + MAP + spatial + uncorrelated |
| 4 | Intercept + MAT + MAP + spatial + uncorrelated |
| 5 | Intercept + MAT + MAP + MAT:MAP + spatial + uncorrelated |
| 6 | Intercept + temp.warmest.quart + spatial + uncorrelated |
| 7 | Intercept + precip.warmest.quart + spatial + uncorrelated |
| 8 | Intercept + precip.coldest.quart + spatial + uncorrelated |
| 9 | Intercept + temp.warmest.quart + precip.warmest.quart + spatial + uncorrelated |
| 10 | Intercept + temp.warmest.quart + precip.coldest.quart + spatial + uncorrelated |
| 11 | Intercept + precip.coldest.quart + precip.warmest.quart + spatial + uncorrelated |
| 12 | Intercept + temp.warmest.quart + precip.coldest.quart + precip.warmest.quart + spatial + uncorrelated |

**Table S5.2 Model fit results for each model tested (model number corresponds to those given in Table S5.1 and the effects specified therein). The top three performing models, based on cAIC criterion are indicated by ***(best), **(second best), *(third best), and the best performing model's cAIC, is highlighted in bold red italics.**

| | Model number | marginal AIC | conditional AIC | Effective df |
|---|---|---|---|---|
| **Early Holocene** | | | | |
| | 1 | 259.9 | 213.3 | 26.4 |
| | 2 | 258.0 | 212.7 | 27.0 |
| * | 3 | 253.1 | 210.9 | 28.3 |
| *** | 4 | 247.0 | ***210.3*** | 29.6 |
| | 5 | 248.4 | 210.9 | 29.3 |
| | 6 | 256.0 | 211.9 | 26.6 |
| | 7 | 251.8 | 211.4 | 28.1 |
| | 8 | 255.3 | 211.9 | 27.4 |
| ** | 9 | 251.3 | 210.8 | 28.0 |
| | 10 | 250.9 | 211.2 | 27.2 |
| | 11 | 250.4 | 211.4 | 28.0 |
| | 12 | 250.6 | 211.2 | 27.4 |

| | | | | |
|---|---|---|---|---|
| **Younger Dryas** | | | | |
| * | 1 | 125.2 | 77.9 | 7.3 |
| | 2 | 126.9 | 78.0 | 7.4 |
| | 3 | 126.8 | 77.9 | 7.3 |
| | 4 | 128.4 | 78.1 | 7.4 |
| | 5 | 130.5 | 78.3 | 7.1 |
| *** | 6 | 127.9 | ***77.7*** | 7.7 |
| | 7 | 126.3 | 77.9 | 7.4 |
| | 8 | 127.5 | 78.0 | 7.2 |
| ** | 9 | 128.3 | 77.8 | 7.6 |
| | 10 | 129.9 | 78.0 | 7.3 |
| | 11 | 128.7 | 78.1 | 7.1 |
| | 12 | 129.1 | 78.0 | 7.4 |
| **Late Glacial Interstadial** | | | | |
| *** | 1 | 198.6 | ***110.5*** | 29.1 |
| | 2 | 196.6 | 110.8 | 28.2 |
| | 3 | 200.3 | 110.9 | 28.7 |
| | 4 | 198.6 | 111.2 | 27.9 |
| | 5 | 200.7 | 111.6 | 27.7 |
| ** | 6 | 194.6 | 110.7 | 28.8 |
| * | 7 | 199.2 | 110.8 | 28.4 |
| | 8 | 201.5 | 111.1 | 28.8 |
| | 9 | 196.6 | 111.1 | 28.3 |
| | 10 | 196.2 | 111.2 | 28.7 |
| | 11 | 199.3 | 111.3 | 28.3 |
| | 12 | 197.9 | 111.4 | 28.2 |
| **Last Glacial Termination** | | | | |
| | 1 | 227.8 | 132.6 | 22.9 |
| *** | 2 | 201.2 | ***131.8*** | 22.7 |
| | 3 | 229.1 | 132.9 | 22.8 |
| | 4 | 201.1 | 132.4 | 22.8 |
| | 5 | 202.4 | 132.5 | 22.4 |
| ** | 6 | 211.6 | 131.9 | 24.0 |
| | 7 | 221.6 | 133.1 | 23.4 |
| | 8 | 227.7 | 132.9 | 22.7 |
| | 9 | 210.4 | 132.2 | 24.4 |
| * | 10 | 213.7 | 132.1 | 23.7 |
| | 11 | 218.3 | 133.3 | 23.7 |
| | 12 | 211.8 | 132.4 | 24.1 |
| **Last Glacial Maximum** | | | | |
| | 1 | 112.6 | 54.1 | 10.8 |
| | 2 | 101.5 | 54.2 | 11.8 |
| | 3 | 114.0 | 54.3 | 10.7 |
| | 4 | 104.0 | 54.3 | 11.7 |
| | 5 | 106.2 | 54.8 | 11.2 |
| | 6 | 97.2 | 56.6 | 12.7 |
| * | 7 | 111.7 | 54.0 | 11.0 |
| ** | 8 | 109.9 | 53.7 | 11.4 |
| | 9 | 98.4 | 56.4 | 12.3 |
| | 10 | 96.1 | 55.6 | 14.0 |

| | | | | |
|---|---|---|---|---|
| *** | 11 | 104.7 | ***52.9*** | 12.4 |
| | 12 | 98.3 | 54.9 | 13.9 |
| **Late OIS 3** | | | | |
| | 1 | 251.8 | 170.2 | 16.0 |
| | 2 | 246.5 | 170.8 | 16.5 |
| *** | 3 | 243.9 | ***169.3*** | 16.4 |
| ** | 4 | 241.5 | 169.3 | 17.0 |
| | 5 | 243.4 | 169.7 | 16.8 |
| | 6 | 244.7 | 170.4 | 17.0 |
| * | 7 | 243.7 | 169.4 | 16.8 |
| | 8 | 247.7 | 169.9 | 16.0 |
| | 9 | 243.5 | 169.8 | 17.0 |
| | 10 | 245.0 | 170.0 | 16.7 |
| | 11 | 244.6 | 169.5 | 16.6 |
| | 12 | 244.7 | 169.8 | 16.7 |
| **Early OIS 3** | | | | |
| * | 1 | 226.3 | 181.2 | 35.1 |
| *** | 2 | 227.2 | ***181.0*** | 34.4 |
| | 3 | 228.4 | 181.9 | 34.4 |
| ** | 4 | 229.2 | 181.1 | 33.5 |
| | 5 | 231.1 | 182.0 | 32.9 |
| | 6 | 228.1 | 181.8 | 34.2 |
| | 7 | 227.8 | 181.6 | 35.0 |
| | 8 | 228.3 | 182.2 | 34.8 |
| | 9 | 229.8 | 182.2 | 34.2 |
| | 10 | 230.1 | 182.7 | 34.0 |
| | 11 | 229.9 | 182.5 | 34.9 |
| | 12 | 232.1 | 183.0 | 34.0 |

289 # 6 Species-specific sample distribution

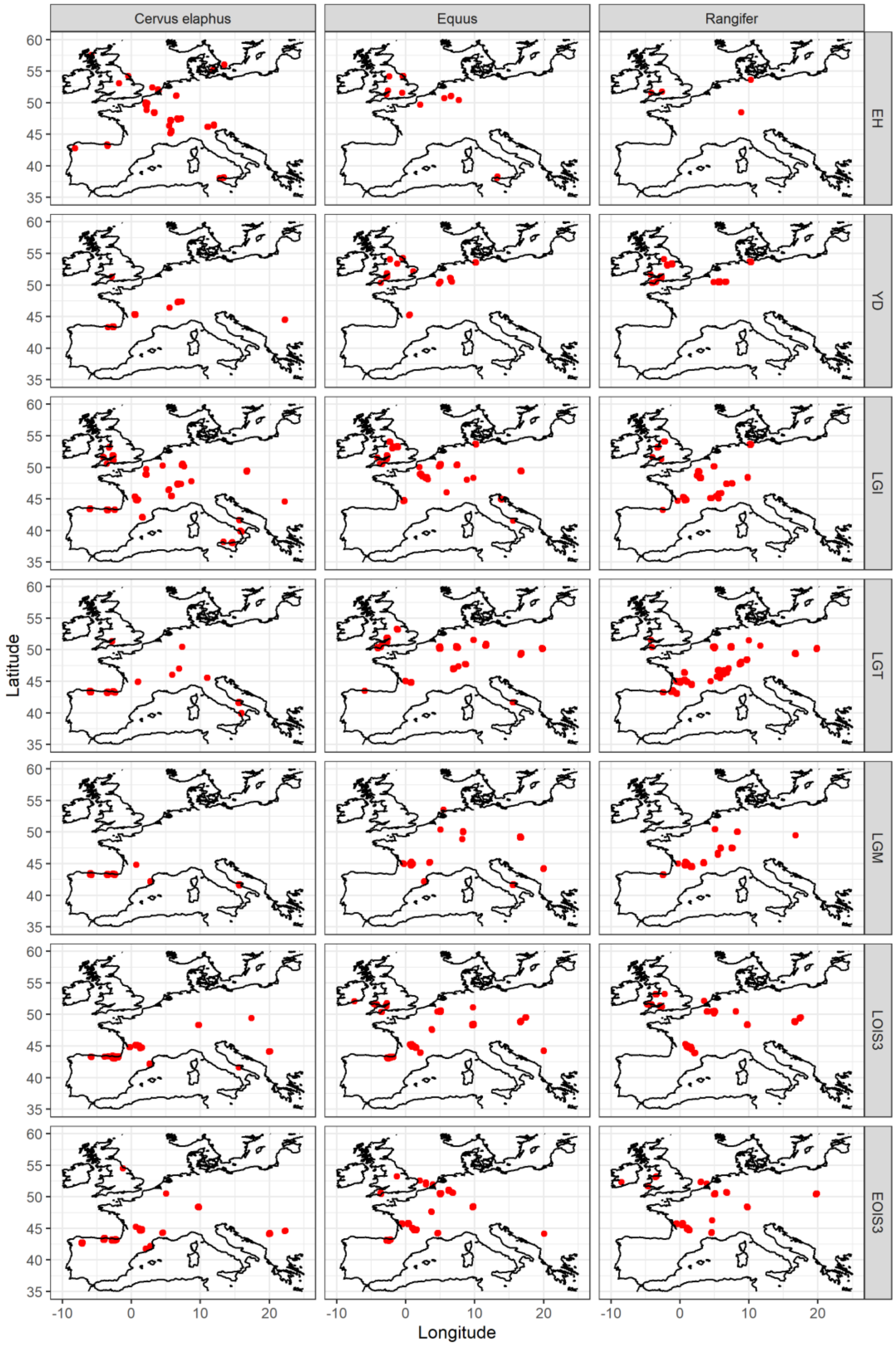


290

291 **Figure S6.1 Distribution of *Cervus elaphus, Equus* sp., and *Rangifer tarandus* samples in the**
292 **compiled data set for each time bin.**
