## Supplementary material for "Nitrogen palaeo-isoscapes: Changing spatial gradients of faunal δ^15^N in late Pleistocene and early Holocene Europe": S4 Base maps for figures: S4 Base maps README.txt

This folder contains processed base maps used in the figures contained within the manuscript.
 These are provided for convenience, so that the figures in this manuscript can be easily reproduced. For all other application users should download the original data.
Palaeocoastline data comes from:
Zickel, M., Becker, D., Verheul, J., Yener, Y., Willmes, C. (2016): Paleocoastlines GIS dataset. CRC806-Database, DOI: 10.5880/SFB806.20
Data were downloaded as a shp file from https://crc806db.uni-koeln.de/layer/show/327 (link last accessed 26th April 2022). Coastlines (palaeo sea-level, expressed in meters relative to modern sea level) used for each time bin are: EOIS3 -65, LOIS3 -80, LGM -120, LGT -80, LGI -65, YD -40, EH -25. Coastlines were processed and saved as individual shp files by decreasing resolution to 0.1 decimal degrees in order to reduce file size.
Ice sheet data comes from:
Hughes, Anna L C; Gyllencreutz, Richard; Lohne, Øystein S; Mangerud, Jan; Svendsen, John-Inge (2015): DATED-1: compilation of dates and time-slice reconstruction of the build-up and retreat of the last Eurasian (British-Irish, Scandinavian, Svalbard-Barents-Kara Seas) Ice Sheets 40-10 ka. Department of Earth Science, University of Bergen and Bjerknes Centre for Climate Research, PANGAEA
Data were downladed as shp files from https://doi.pangaea.de/10.1594/PANGAEA.848117?format=html#download (link last accessed 26th April 2022). Ice sheet shp timeslices used for each time bin are: EOIS3 TS_3835max_poly, LOIS3 TS_3230_poly, LGM TS21_mc, LGT TS16_mc, LGI TS14_mc, YD TS12_mc, and EH TS10_mc. Each shp was reprojected and saved under our time bin naming convention
Modern coastline data comes from:
Wessel, P., and W. H. F. Smith (1996), A global, self-consistent, hierarchical, high-resolution shoreline database, J. Geophys. Res., 101(B4), 8741-8743, doi:10.1029/96JB00104.
Data was downloaded from https://www.ngdc.noaa.gov/mgg/shorelines/ (link last accessed 26th April 2022). We use the GSHHS_l_L1.shp file, which provides a low resolution map of shorelines for continental land masses and ocean islands, except Antarctica the downloaded file was cropped and saved under the file name ‘eurmed.shp’.
Elevation data comes from:
Danielson, J. J., & Gesch, D. B. (2011). Global multi-resolution terrain elevation data 2010 (GMTED2010). https://doi.org/10.3133/ofr20111073
Data were downloaded as a tif file ("gmted2010_30mn.tif") from https://topotools.cr.usgs.gov/gmted_viewer/viewer.htm (link last accessed 26th April 2022). A cropped, lower resolution version is provided ("EURDEM01.tif", 310KB)
