## Supplementary material for "Nitrogen palaeo-isoscapes: Changing spatial gradients of faunal δ^15^N in late Pleistocene and early Holocene Europe": S2 Script

```
#### Nitrogen palaeo-isoscapes: Changing spatial gradients of faunal d15N in late Pleistocene and early Holocene Europe
#### Hazel Reade, Jennifer A. Tripp, Delphine Frémondeau, Kerry L. Sayle, Thomas F.G. Higham, Martin Street, Rhiannon E. Stevens

#### R script for running analysis contained within manuscript
#### Corresponding author: Hazel Reade

#### COMMENTS ON FILES REQUIRED ####

### this section details where original data used in the subsequent analysis and plots can be accessed

 ## data
  ## faunal isotope data is provided as Supporting Information (S1 Dataset) attached to this manuscript
  ## elevation data comes from the Danielson, J. J., & Gesch, D. B. (2011). Global multi-resolution terrain elevation data 2010
(GMTED2010). https://doi.org/10.3133/ofr20111073
    # data were downloaded as a tif file ("gmted2010_30mn.tif") from https://topotools.cr.usgs.gov/gmted_viewer/viewer.htm (link
last accessed 26th April 2022)
    # users should download this file directly (large file size of c. 700MB) for extracting elevation to point
    # a cropped, lower resolution version is also provide in the S4 Base maps folder ("EURDEM01.tif", 310KB)
  ## bioclimatic data comes from Beyer, R.M., Krapp, M. & Manica, A. Addendum: High-resolution terrestrial climate, bioclimate and
vegetation for the last 120,000 years. Sci Data 8, 262 (2021).
    # data was downloaded as a NetCDF file ("LateQuaternary_Environment.nc") from the Figshare data repository via
https://www.nature.com/articles/s41597-020-0552-1 (link last accessed 26th April 2022)
    # users should download this file directly (large file size of c. 1.3GB)

 ## base maps used in figures
  ## for convenience processed basemaps are provided in Supporting Information (S4 Base maps for figures)
  ## these are provided so that the figures in this manuscript can be easily reproduced.
  ## For all other application users should downlaod the original data.
  ## palaocoastlines use data from Zickel, M., Becker, D., Verheul, J., Yener, Y., Willmes, C. (2016): Paleocoastlines GIS dataset.
CRC806-Database, DOI: 10.5880/SFB806.20
    # data were downloaded as a shp file from https://crc806db.uni-koeln.de/layer/show/327 (link last accessed 26th April 2022)
    # coastline (palaeo sea-level, expressed in meters relative to modern sea level) used in plots are: EOIS3 -65, LOIS3 -80, LGM
-120, LGT -80, LGI -65, YD -40, EH -25
    # coastlines were processed and saved as individual shp files by decreasing resolution to 0.1 decimal degrees in order to reduce
file size, e.g.:
      # pcoast <- readOGR("paleocoastlines.shp") # load data
      # latlon <- CRS("+init=epsg:4326 +units=km") # define unprojected latlon coordinate system
      # proj4string(pcoast) <- latlon # add coordinate system info
      # pcoastLGM <- pcoast[pcoast$Sea_level==-120,] #select specific sea level for time period
      # pcoastLGM.simp <- gSimplify(pcoastLGM, tol=0.1) # simplfy coastline
      # pcoastLGM.simp.spdf <- as(pcoastLGM.simp, "SpatialPolygonsDataFrame") # convert back to spdf
      # writeOGR(pcoastLGM.simp.spdf, ".", "pcoastLGM.simp", driver = "ESRI Shapefile") # save as shp file
  ## ice sheet extent use data from Hughes, Anna L C; Gyllencreutz, Richard; Lohne, Øystein S; Mangerud, Jan; Svendsen, John-Inge
(2015): DATED-1: compilation of dates and time-slice reconstruction of the build-up and retreat of the last Eurasian (British-Irish,
Scandinavian, Svalbard-Barents-Kara Seas) Ice Sheets 40-10 ka. Department of Earth Science, University of Bergen and Bjerknes Centre
for Climate Research, PANGAEA
    # data were downladed as shp files from https://doi.pangaea.de/10.1594/PANGAEA.848117?format=html#download (link last accessed
26th April 2022)
    # ice sheet shp timeslices used for plotting are: EOIS3 TS_3835max_poly, LOIS3 TS_3230_poly, LGM TS21_mc, LGT TS16_mc, LGI
TS14_mc, YD TS12_mc, and EH TS10_mc
    # each shp file was reprojected and saved under our time bin naming convention, e.g.:
      # iceLGM <- readOGR("TS21_mc.shp") # load data
      # latlon <- CRS("+init=epsg:4326 +units=km") # define unprojected latlon coordinate system
      # iceLGM.trans <- spTransform(iceLGM,latlon) # transform projection
      # writeOGR(iceLGM.trans, ".", "iceLGM", driver = "ESRI Shapefile") # save as shp file
  ## plotted modern coastlines use data from Wessel, P., and W. H. F. Smith (1996), A global, self-consistent, hierarchical, high-
resolution shoreline database, J. Geophys. Res., 101(B4), 8741-8743, doi:10.1029/96JB00104.
    # data was downloaded from https://www.ngdc.noaa.gov/mgg/shorelines/ (link last accessed 26th April 2022)
    # the GSHHS_l_L1.shp file was used which provides a low resolution map of shorelines for continental land masses and ocean
islands, except Antarctica
    # the downloaded file was cropped and saved as follows:
      # eurmed <- readOGR("GSHHS_l_L1.shp")
      # eurmed<- crop(eurmed, extent(-10, 30, 35, 60))
      # writeOGR(eurmed, ".", "eurmed", driver = "ESRI Shapefile") # save as shp file

#### LOAD REQUIRED PACKAGES ####
library(ggplot2)
library(openxlsx)
library(dplyr)
library(raster)
library(rgdal)
library(ncdf4)
library(grid)
library(gtable)
library(cowplot)
library(gridExtra)
library(geosphere)
library(ggpubr)
library(rstatix)
library(devtools)
install_github("ahb108/sparch")
library(sparch)
library(spdep)
library(gstat)
library(automap)
library(RColorBrewer)
library(spaMM)
library(ggpmisc)
library(Hmisc)
library(knitr)
library(tidyverse, warn.conflict=F)
library(ggrepel)


#### LOAD REQUIRED FILES ####
```

```
# we recommend downloading and saving all data files and base map files to an appropriate folder

# setwd("folder name") # specify folder where files are saved

## load data files:
  # isotope data
    final_isoscape_data <- read.xlsx("S1 Dataset.xlsx", sheet=1, colNames =TRUE)
    # columns may need renaming to prevent errors:
      colnames(final_isoscape_data) # check column names and rename as necessary, e.g:
      final_isoscape_data <- final_isoscape_data %>% rename(MedianDate = `Median.Date.cal.BP.(2.sigma)`)
      final_isoscape_data <- final_isoscape_data %>% rename(d15Ncoll = "????15N.(???)")
      final_isoscape_data <- final_isoscape_data %>% rename(finalagebin = Assigned.Isoscape.Age.Bin)
      final_isoscape_data <- final_isoscape_data %>% rename(d15NRef = "????15N.reference")
  # elevation data
    elev <- raster("gmted2010_30mn.tif", verbose=FALSE)
  # climate data
    beyerdata <- "LateQuaternary_Environment.nc"

# load base maps:
eurmed <- readOGR("eurmed.shp") # current coastline
iceEOIS3 <- readOGR("iceEOIS3.shp") # EOIS3 ice
iceLOIS3 <- readOGR("iceLOIS3.shp") # LOIS3 ice
iceLGM <- readOGR("iceLGM.shp") # LGM ice
iceGS2 <- readOGR("iceGS2.shp") # LGT ice
iceGI1<- readOGR("iceGI1.shp") # LGI ice
iceGS1 <- readOGR("iceGS1.shp") # YD ice
iceEH <- readOGR("iceEH.shp") # EH ice
pcoastEOIS3.simp <- readOGR("pcoastEOIS3.simp.shp.shp") # EOIS3 coastline
pcoastLOIS3.simp <- readOGR("pcoastLOIS3.simp.shp.shp") # LOIS3 coastline
pcoastLGM.simp <- readOGR("pcoastLGM.simp.shp.shp") # LGM coastline
pcoastGS2.simp <- readOGR("pcoastGS2.simp.shp.shp") # LGT coastline
pcoastGI1.simp <- readOGR("pcoastGI1.simp.shp.shp") # LGI coastline
pcoastGS1.simp <- readOGR("pcoastGS1.simp.shp.shp") # YD coastline
pcoastEH.simp <- readOGR("pcoastEH.simp.shp.shp") # EH coastline


#### ASSEMBLE COVARIATE DATA ####
  ## extract elevation data (note this is based on modern DEM)
    names(elev) <- 'elevation'
    coordinates(final_isoscape_data) = ~ Longitude + Latitude
    elevation <- raster::extract(elev, final_isoscape_data) # extract values
    final_isoscape_data <- as.data.frame(final_isoscape_data)
    final_isoscape_data <- cbind(final_isoscape_data, 'elevation' = elevation)
    colnames(final_isoscape_data)
    rm(elev) # remove elevation file from work environment

  ## extract climate data from Beyer et al., 2020
    # first check distribution of samples:
      # by 1000-year bin widths
        ggplot() +
        geom_histogram(data= final_isoscape_data, aes(MedianDate), binwidth = 1000) +
        facet_wrap(~ finalagebin, scales = "free")
      # by 2000-year bin widths
        ggplot() +
        geom_histogram(data= final_isoscape_data, aes(MedianDate), binwidth = 2000) +
        facet_wrap(~ finalagebin, scales = "free")

  ## file climate data as raster bricks (one per variable)
    MAT <- brick(beyerdata, varname = "BIO1")  # mean annual temperature
    tempseason <- brick(beyerdata, varname = "BIO4")  # temperature seasonality
    tempmin <- brick(beyerdata, varname = "BIO5") # min annual temp
    tempmax <- brick(beyerdata, varname = "BIO6") # max annual temp
    temprange <- brick(beyerdata, varname = "BIO7")  # temp annual range
    tempwettest <- brick(beyerdata, varname = "BIO8")  # mean temp of wettest quarter
    tempdriest <- brick(beyerdata, varname = "BIO9") # mean temp of direst quarter
    tempwarmest <- brick(beyerdata, varname = "BIO10") # mean temp of warmest quarter
    tempcoldest <- brick(beyerdata, varname = "BIO11")  # mean temp of coldest quarter
    MAP <- brick(beyerdata, varname = "BIO12") # mean annual precipitation
    precip.wet.month <- brick(beyerdata, varname = "BIO13") # precipitation of wettest month
    precip.dry.month <- brick(beyerdata, varname = "BIO14") # precipitation of driest month
    precipseason <- brick(beyerdata, varname = "BIO15") # precipitation seasonality
    precip.wet.quar <- brick(beyerdata, varname = "BIO16") # precipitation of wettest quarter
    precip.dry.quar <- brick(beyerdata, varname = "BIO17") # precipitation of driest quarter
    precip.warm.quart <- brick(beyerdata, varname = "BIO18") # precipitation of warmest quarter
    precip.cold.quart <- brick(beyerdata, varname = "BIO19") # precipitation of coldest quarter

  # create a raster stack for each time period of interest (time slices selected based on distibution plots)
    stack.11k <- stack(MAP[[61]], MAT[[61]], precip.cold.quart[[61]], precip.dry.month[[61]],  precip.dry.quar[[61]],
precip.warm.quart[[61]], precip.wet.month[[61]],  precip.wet.quar[[61]], precipseason[[61]], tempcoldest[[61]], tempdriest[[61]],
tempmax[[61]], tempmin[[61]], temprange[[61]], tempseason[[61]], tempwarmest[[61]], tempwettest[[61]])
    stack.12k <- stack(MAP[[60]], MAT[[60]], precip.cold.quart[[60]], precip.dry.month[[60]],  precip.dry.quar[[60]],
precip.warm.quart[[60]], precip.wet.month[[60]],  precip.wet.quar[[60]], precipseason[[60]], tempcoldest[[60]], tempdriest[[60]],
tempmax[[60]], tempmin[[60]], temprange[[60]], tempseason[[60]], tempwarmest[[60]], tempwettest[[60]])
    stack.14k <- stack(MAP[[58]], MAT[[58]], precip.cold.quart[[58]], precip.dry.month[[58]],  precip.dry.quar[[58]],
precip.warm.quart[[58]], precip.wet.month[[58]],  precip.wet.quar[[58]], precipseason[[58]], tempcoldest[[58]], tempdriest[[58]],
tempmax[[58]], tempmin[[58]], temprange[[58]], tempseason[[58]], tempwarmest[[58]], tempwettest[[58]])
    stack.15k <- stack(MAP[[57]], MAT[[57]], precip.cold.quart[[57]], precip.dry.month[[57]],  precip.dry.quar[[57]],
precip.warm.quart[[57]], precip.wet.month[[57]],  precip.wet.quar[[57]], precipseason[[57]], tempcoldest[[57]], tempdriest[[57]],
tempmax[[57]], tempmin[[57]], temprange[[57]], tempseason[[57]], tempwarmest[[57]], tempwettest[[57]])
    stack.24k <- stack(MAP[[49]], MAT[[49]], precip.cold.quart[[49]], precip.dry.month[[49]],  precip.dry.quar[[49]],
precip.warm.quart[[49]], precip.wet.month[[49]],  precip.wet.quar[[49]], precipseason[[49]], tempcoldest[[49]], tempdriest[[49]],
tempmax[[49]], tempmin[[49]], temprange[[49]], tempseason[[49]], tempwarmest[[49]], tempwettest[[49]])
    stack.36k <- stack(MAP[[43]], MAT[[43]], precip.cold.quart[[43]], precip.dry.month[[43]],  precip.dry.quar[[43]],
precip.warm.quart[[43]], precip.wet.month[[43]],  precip.wet.quar[[43]], precipseason[[43]], tempcoldest[[43]], tempdriest[[43]],
```

```
tempmax[[43]], tempmin[[43]], temprange[[43]], tempseason[[43]], tempwarmest[[43]], tempwettest[[43]])
    stack.42k <- stack(MAP[[40]], MAT[[40]], precip.cold.quart[[40]], precip.dry.month[[40]],  precip.dry.quar[[40]],
precip.warm.quart[[40]], precip.wet.month[[40]],  precip.wet.quar[[40]], precipseason[[40]], tempcoldest[[40]], tempdriest[[40]],
tempmax[[40]], tempmin[[40]], temprange[[40]], tempseason[[40]], tempwarmest[[40]], tempwettest[[40]])

  # set names for the raster stacks
    names(stack.11k) <- c("MAP", "MAT", "precip.cold.quart", "precip.dry.month",
                      "precip.dry.quar", "precip.warm.quart", "precip.wet.month",  "precip.wet.quar",
                      "precipseason", "tempcoldest", "tempdriest", "tempmax", "tempmin", "temprange", "tempseason",
                      "tempwarmest", "tempwettest")
    names(stack.12k) <- c("MAP", "MAT", "precip.cold.quart", "precip.dry.month",
                      "precip.dry.quar", "precip.warm.quart", "precip.wet.month",  "precip.wet.quar",
                      "precipseason", "tempcoldest", "tempdriest", "tempmax", "tempmin", "temprange", "tempseason",
                      "tempwarmest", "tempwettest")
    names(stack.14k) <- c("MAP", "MAT", "precip.cold.quart", "precip.dry.month",
                      "precip.dry.quar", "precip.warm.quart", "precip.wet.month",  "precip.wet.quar",
                      "precipseason", "tempcoldest", "tempdriest", "tempmax", "tempmin", "temprange", "tempseason",
                      "tempwarmest", "tempwettest")
    names(stack.15k) <- c("MAP", "MAT", "precip.cold.quart", "precip.dry.month",
                      "precip.dry.quar", "precip.warm.quart", "precip.wet.month",  "precip.wet.quar",
                      "precipseason", "tempcoldest", "tempdriest", "tempmax", "tempmin", "temprange", "tempseason",
                      "tempwarmest", "tempwettest")
    names(stack.24k) <- c("MAP", "MAT", "precip.cold.quart", "precip.dry.month",
                      "precip.dry.quar", "precip.warm.quart", "precip.wet.month",  "precip.wet.quar",
                      "precipseason", "tempcoldest", "tempdriest", "tempmax", "tempmin", "temprange", "tempseason",
                      "tempwarmest", "tempwettest")
    names(stack.36k) <- c("MAP", "MAT", "precip.cold.quart", "precip.dry.month",
                      "precip.dry.quar", "precip.warm.quart", "precip.wet.month",  "precip.wet.quar",
                      "precipseason", "tempcoldest", "tempdriest", "tempmax", "tempmin", "temprange", "tempseason",
                      "tempwarmest", "tempwettest")
    names(stack.42k) <- c("MAP", "MAT", "precip.cold.quart", "precip.dry.month",
                      "precip.dry.quar", "precip.warm.quart", "precip.wet.month",  "precip.wet.quar",
                      "precipseason", "tempcoldest", "tempdriest", "tempmax", "tempmin", "temprange", "tempseason",
                      "tempwarmest", "tempwettest")
  # crop raster stacks
    stack.11k <- crop(stack.11k, extent(-10, 30, 35, 60))
    stack.12k <- crop(stack.12k, extent(-10, 30, 35, 60))
    stack.14k <- crop(stack.14k, extent(-10, 30, 35, 60))
    stack.15k <- crop(stack.15k, extent(-10, 30, 35, 60))
    stack.24k <- crop(stack.24k, extent(-10, 30, 35, 60))
    stack.36k <- crop(stack.36k, extent(-10, 30, 35, 60))
    stack.42k <- crop(stack.42k, extent(-10, 30, 35, 60))

  # divide isotope data by time bin
    EH <- final_isoscape_data %>%
          subset(finalagebin == "EH")
    GS1 <- final_isoscape_data %>%
            subset(finalagebin == "YD")
    GI1 <- final_isoscape_data %>%
            subset(finalagebin == "LGI")
    GS2 <- final_isoscape_data %>%
            subset(finalagebin == "LGT")
    LGM <-  final_isoscape_data %>%
            subset(finalagebin == "LGM")
    LOIS3 <-  final_isoscape_data %>%
              subset(finalagebin == "LOIS3")
    EOIS3 <-  final_isoscape_data %>%
              subset(finalagebin == "EOIS3")

  # convert data to spdf
    latlon <- CRS("+init=epsg:4326 +units=km") # unprojected latlon coordinate system
    coordinates(EH) <- ~Longitude+Latitude
    proj4string(EH) <- latlon
    coordinates(GS1) <- ~Longitude+Latitude
    proj4string(GS1) <- latlon
    coordinates(GI1) <- ~Longitude+Latitude
    proj4string(GI1) <- latlon
    coordinates(GS2) <- ~Longitude+Latitude
    proj4string(GS2) <- latlon
    coordinates(LGM) <- ~Longitude+Latitude
    proj4string(LGM) <- latlon
    coordinates(LOIS3) <- ~Longitude+Latitude
    proj4string(LOIS3) <- latlon
    coordinates(EOIS3) <- ~Longitude+Latitude
    proj4string(EOIS3) <- latlon

  # extract data for each stage and merge back to original data frame
    raster_data_11k <- raster::extract(stack.11k, EH, buffer = 100000, fun=max, na.rm=TRUE)
    EH <- cbind(EH, raster_data_11k)
    raster_data_12k <- raster::extract(stack.12k, GS1, buffer = 100000, fun=max, na.rm=TRUE)
    GS1 <- cbind(GS1, raster_data_12k )
    raster_data_14k <- raster::extract(stack.14k, GI1, buffer = 100000, fun=max, na.rm=TRUE)
    GI1 <- cbind(GI1, raster_data_14k )
    raster_data_15k <- raster::extract(stack.15k, GS2, buffer = 100000, fun=max, na.rm=TRUE)
    GS2 <- cbind(GS2, raster_data_15k )
    raster_data_24k <- raster::extract(stack.24k, LGM, buffer = 100000, fun=max, na.rm=TRUE)
    LGM <- cbind(LGM, raster_data_24k )
    raster_data_36k <- raster::extract(stack.36k, LOIS3, buffer = 100000, fun=max, na.rm=TRUE)
    LOIS3 <- cbind(LOIS3, raster_data_36k )
    raster_data_42k <- raster::extract(stack.42k, EOIS3, buffer = 100000, fun=max, na.rm=TRUE)
    EOIS3 <- cbind(EOIS3, raster_data_42k )

  # reassemble data frame:
    # convert spdfs to dfs
      EH <- as.data.frame(EH)
```

```
      GS1 <- as.data.frame(GS1)
      GI1 <- as.data.frame(GI1)
      GS2 <- as.data.frame(GS2)
      LGM <- as.data.frame(LGM)
      LOIS3 <- as.data.frame(LOIS3)
      EOIS3 <- as.data.frame(EOIS3)
    # combine time bins
      final_isoscape_data <- rbind(EH, GS1, GI1, GS2, LGM, LOIS3, EOIS3)

## at this point you may want to save a copy of the combined data set:
  # write.csv(final_isoscape_data, file="final_isoscape_data.csv")

#### FUNCTIONS FOR SUBESQUENT PLOTTING AND ANALYSIS ####
## function to shift legend into empty facet panels in plots (see https://stackoverflow.com/questions/54438495/shift-legend-into-
empty-facets-of-a-faceted-plot-in-ggplot2) ##
shift_legend <- function(p){

  # check if p is a valid object
  if(!"gtable" %in% class(p)){
    if("ggplot" %in% class(p)){
      gp <- ggplotGrob(p) # convert to grob
    } else {
      message("This is neither a ggplot object nor a grob generated from ggplotGrob. Returning original plot.")
      return(p)
    }
  } else {
    gp <- p
  }

  # check for unfilled facet panels
  facet.panels <- grep("^panel", gp[["layout"]][["name"]])
  empty.facet.panels <- sapply(facet.panels, function(i) "zeroGrob" %in% class(gp[["grobs"]][[i]]))
  empty.facet.panels <- facet.panels[empty.facet.panels]
  if(length(empty.facet.panels) == 0){
    message("There are no unfilled facet panels to shift legend into. Returning original plot.")
    return(p)
  }

  # establish extent of unfilled facet panels (including any axis cells in between)
  empty.facet.panels <- gp[["layout"]][empty.facet.panels, ]
  empty.facet.panels <- list(min(empty.facet.panels[["t"]]), min(empty.facet.panels[["l"]]),
                             max(empty.facet.panels[["b"]]), max(empty.facet.panels[["r"]]))
  names(empty.facet.panels) <- c("t", "l", "b", "r")

  # extract legend & copy over to location of unfilled facet panels
  guide.grob <- which(gp[["layout"]][["name"]] == "guide-box")
  if(length(guide.grob) == 0){
    message("There is no legend present. Returning original plot.")
    return(p)
  }
  gp <- gtable_add_grob(x = gp,
                        grobs = gp[["grobs"]][[guide.grob]],
                        t = empty.facet.panels[["t"]],
                        l = empty.facet.panels[["l"]],
                        b = empty.facet.panels[["b"]],
                        r = empty.facet.panels[["r"]],
                        name = "new-guide-box")

  # squash the original guide box's row / column (whichever applicable)
  # & empty its cell
  guide.grob <- gp[["layout"]][guide.grob, ]
  if(guide.grob[["l"]] == guide.grob[["r"]]){
    gp <- gtable_squash_cols(gp, cols = guide.grob[["l"]])
  }
  if(guide.grob[["t"]] == guide.grob[["b"]]){
    gp <- gtable_squash_rows(gp, rows = guide.grob[["t"]])
  }
  gp <- gtable_remove_grobs(gp, "guide-box")

  return(gp)
}
## kruskal-wallis function
      krus_test <- function(x, ...) {
        result <- kruskal.test(x, ...)

        tibble(
          p.value = result$p.value,
          statistic = result$statistic
        )
      }
## mann whitney function
      wilc_test <- function(x, ...) {
        result <- wilcox.test(x, ...)

        tibble(
          p.value = result$p.value,
          statistic = result$statistic
          # and any other values you want to capture from the return
        )
      }
## RMSE function (measure of the differences between observed and predicted, or an average deviation of the predicted vs observed
values)
      modelRMSE <- function(o, p) {
        sqrt(sum((o-p)^2) / (length(o)))
      }
```

```
#### INTER-SPECIES COMPARISONS (S3 SUPPORTING INFORMATION) ####

  ## reload data if necessary: isoscape_data <- read.csv("final_isoscape_data.csv", header = TRUE) ##

  ## Fig S2.1 ####

  # define order of plots and set variables as factors where needed
    isoscape_data$Faunal.Category <- factor(isoscape_data$Faunal.Category, levels = c("Alces", "BosBison", "Capra", "Capreolus",
"Cervus elaphus", "Coelodonta",
                                                                                  "Equus", "Megaloceros", "Ovibos", "Rangifer",
"Rupicapra", "Saiga"))
    isoscape_data$finalagebin  <- factor(isoscape_data$finalagebin, levels = c("EH", "YD", "LGI", "LGT", "LGM", "LOIS3", "EOIS3"))
    isoscape_data$Trophic.Position <- factor(isoscape_data$Trophic.Position, levels = c("browser", "mixed", "grazer"))

  # plot all data by species, dietary category and age category
  (species.tro.pos <- ggplot() +
    geom_boxplot(data=isoscape_data, aes(x=Faunal.Category, y=d15Ncoll, fill=Trophic.Position)) +
    scale_fill_brewer(palette = "Dark2", name = "Dominant\ndietary\ncharacteristic")+
    facet_wrap(~finalagebin, nrow=4, scales="free") +
    ylim(-1,12) +
   theme_bw()+
    ylab(expression(paste(delta^{15}, "N (\u2030)"))) +
    theme(text = element_text(size=16),
        legend.title.align=0.5,
        axis.text.x = element_text(angle=90, hjust=1, vjust = 0.25),
        axis.title.x = element_blank()))

  # save output
  pdf(file="Figure_SI2_1.pdf", width = 10, height = 15)
  grid.draw(shift_legend(species.tro.pos))
  dev.off()

  ## TEST FOR STATISTICAL DIFFERENCES BETWEEN ALL SPECIES/DIETARY CHARACTERISTICS, WHOLE GEOGRAPHIC AREA, BY TIME BIN (TABLE S2.1)
####

    # SPECIES

    # set up dataframes

    # select only species in each age bracket where there are at least 3 data points (for kruskal-wallis test):
      isoscape_data.3 <- isoscape_data %>%
      dplyr::group_by(finalagebin, Faunal.Category) %>%
      filter(n() >= 3)

    # run kruskal-wallis test
      kruskall.wallis.output.spec <- isoscape_data.3 %>%
      group_by(finalagebin) %>%
      group_modify(~ krus_test(.x$d15Ncoll ~ .x$Faunal.Category))

    ## DIET

    # set up dataframes

    # select only dietary groups in each age bracket where there are at least 3 data points:
      isoscape_data.3.tro <- isoscape_data %>%
      dplyr::group_by(finalagebin, Trophic.Position) %>%
      filter(n() >= 3)

    # run kruskal-wallis test
      kruskall.wallis.output.tro <- isoscape_data.3.tro %>%
      group_by(finalagebin) %>%
      group_modify(~ krus_test(.x$d15Ncoll ~ .x$Trophic.Position))

    # combine output (Table S2.1)
      TableS2.1 <- cbind (kruskall.wallis.output.spec, kruskall.wallis.output.tro)

  ## DIFFERENCES BETWEEN SPECIES BY GEOGRAPHICAL CLUSTER AND TIME BIN (TABLE S2.2 AND FIGURE S2.2) ####

  ## DIVIDE DATA INTO GEOGRAPHICAL CLUSTER##
    ## NOTE: EACH TIME CLUSTERS ARE DEFINED THEY ARE NUMBER DIFFERENTLY, BUT THE ACTUAL CLUSTERS REMAIN IDENTICAL
    ## THIS MEANS THAT THE NUMBERING AND ORDERING OF CLUSTERS IN TABLE S2.2 and FIG2.2 VARIES EACH TIME THE CODE IS RUN
    ## HOWEVER THIS DOES NOT CHANGE THE RESULTS

    # first divide data in time binned dataframes
      isoscape_data.df.EH <- isoscape_data[isoscape_data$finalagebin == "EH", ]
      isoscape_data.df.GS1 <- isoscape_data[isoscape_data$finalagebin == "YD", ]
      isoscape_data.df.GI1 <- isoscape_data[isoscape_data$finalagebin == "LGI", ]
      isoscape_data.df.GS2 <- isoscape_data[isoscape_data$finalagebin == "LGT", ]
      isoscape_data.df.LGM <- isoscape_data[isoscape_data$finalagebin == "LGM", ]
      isoscape_data.df.LOIS3 <- isoscape_data[isoscape_data$finalagebin == "LOIS3", ]
      isoscape_data.df.EOIS3 <- isoscape_data[isoscape_data$finalagebin == "EOIS3", ]

    # create geographical clusters for each time bin
      latlon <- CRS("+init=epsg:4326 +units=km") # unprojected latlon coordinate system
      # EH
        # create spdf from df
        isoscape_data.spdf.EH <- isoscape_data.df.EH
        coordinates(isoscape_data.spdf.EH) <- ~Longitude+Latitude
        # add coordinate system info to spdf
        proj4string(isoscape_data.spdf.EH) <- latlon # add this info to the datasets definition
        # use the distm function to generate a geodesic distance matrix in meters
        mdistEH <- distm(isoscape_data.spdf.EH)
        # cluster all points using a hierarchical clustering approach
        hcEH <- hclust(as.dist(mdistEH), method="complete")
        # define the distance threshold, in this case 100km
```

```
  d=100000
  # define clusters based on a tree "height" cutoff "d" and add them to the SpDataFrame
  isoscape_data.spdf.EH$clust <- cutree(hcEH, h=d)
  # convert back to df
  isoscape_data.EH.clust <- as.data.frame(isoscape_data.spdf.EH)

# YD
  # create spdf from df
  isoscape_data.GS1.spdf <- isoscape_data.df.GS1
  coordinates(isoscape_data.GS1.spdf) <- ~Longitude+Latitude
  # add coordinate system info to spdf
  proj4string(isoscape_data.GS1.spdf) <- latlon # add this info to the datasets definition
  # use the distm function to generate a geodesic distance matrix in meters
  mdistGS1 <- distm(isoscape_data.GS1.spdf)
  # cluster all points using a hierarchical clustering approach
  hcGS1 <- hclust(as.dist(mdistGS1), method="complete")
  # define the distance threshold, in this case 100km (100000m)
  d=100000
  # define clusters based on a tree "height" cutoff "d" and add them to the SpDataFrame
  isoscape_data.GS1.spdf$clust <- cutree(hcGS1, h=d)
  # convert back to df
  isoscape_data.GS1.clust <- as.data.frame(isoscape_data.GS1.spdf)

# LGI
  # create spdf from df
  isoscape_data.GI1.spdf <- isoscape_data.df.GI1
  coordinates(isoscape_data.GI1.spdf) <- ~Longitude+Latitude
  # add coordinate system info to spdf
  proj4string(isoscape_data.GI1.spdf) <- latlon # add this info to the datasets definition
  # use the distm function to generate a geodesic distance matrix in meters
  mdistGI1 <- distm(isoscape_data.GI1.spdf)
  # cluster all points using a hierarchical clustering approach
  hcGI1 <- hclust(as.dist(mdistGI1), method="complete")
  # define the distance threshold, in this case 100km (100000m)
  d=100000
  # define clusters based on a tree "height" cutoff "d" and add them to the SpDataFrame
  isoscape_data.GI1.spdf$clust <- cutree(hcGI1, h=d)
  # convert back to df
  isoscape_data.GI1.clust <- as.data.frame(isoscape_data.GI1.spdf)

# LGT
  # create spdf from df
  isoscape_data.GS2.spdf <- isoscape_data.df.GS2
  coordinates(isoscape_data.GS2.spdf) <- ~Longitude+Latitude
  # add coordinate system info to spdf
  proj4string(isoscape_data.GS2.spdf) <- latlon # add this info to the datasets definition
  # use the distm function to generate a geodesic distance matrix in meters
  mdistGS2 <- distm(isoscape_data.GS2.spdf)
  # cluster all points using a hierarchical clustering approach
  hcGS2 <- hclust(as.dist(mdistGS2), method="complete")
  # define the distance threshold, in this case 100km (100000m)
  d=100000
  # define clusters based on a tree "height" cutoff "d" and add them to the SpDataFrame
  isoscape_data.GS2.spdf$clust <- cutree(hcGS2, h=d)
  # convert back to df
  isoscape_data.GS2.clust <- as.data.frame(isoscape_data.GS2.spdf)

#LGM
  # create spdf from df
  isoscape_data.LGM.spdf <- isoscape_data.df.LGM
  coordinates(isoscape_data.LGM.spdf) <- ~Longitude+Latitude
  # add coordinate system info to spdf
  proj4string(isoscape_data.LGM.spdf) <- latlon # add this info to the datasets definition
  # use the distm function to generate a geodesic distance matrix in meters
  mdistLGM <- distm(isoscape_data.LGM.spdf)
  # cluster all points using a hierarchical clustering approach
  hcLGM <- hclust(as.dist(mdistLGM), method="complete")
  # define the distance threshold, in this case 100km (100000m)
  d=100000
  # define clusters based on a tree "height" cutoff "d" and add them to the SpDataFrame
  isoscape_data.LGM.spdf$clust <- cutree(hcLGM, h=d)
  # convert back to df
  isoscape_data.LGM.clust <- as.data.frame(isoscape_data.LGM.spdf)

#LOIS3
  # create spdf from df
  isoscape_data.LOIS3.spdf <- isoscape_data.df.LOIS3
  coordinates(isoscape_data.LOIS3.spdf) <- ~Longitude+Latitude
  # add coordinate system info to spdf
  proj4string(isoscape_data.LOIS3.spdf) <- latlon # add this info to the datasets definition
  # use the distm function to generate a geodesic distance matrix in meters
  mdistLOIS3 <- distm(isoscape_data.LOIS3.spdf)
  # cluster all points using a hierarchical clustering approach
  hcLOIS3 <- hclust(as.dist(mdistLOIS3), method="complete")
  # define the distance threshold, in this case 100km (100000m)
  d=100000
  # define clusters based on a tree "height" cutoff "d" and add them to the SpDataFrame
  isoscape_data.LOIS3.spdf$clust <- cutree(hcLOIS3, h=d)
  # convert back to df
  isoscape_data.LOIS3.clust <- as.data.frame(isoscape_data.LOIS3.spdf)

#EOIS3
  # create spdf from df
  isoscape_data.EOIS3.spdf <- isoscape_data.df.EOIS3
  coordinates(isoscape_data.EOIS3.spdf) <- ~Longitude+Latitude
```

```
        # add coordinate system info to spdf
        proj4string(isoscape_data.EOIS3.spdf) <- latlon # add this info to the datasets definition
        # use the distm function to generate a geodesic distance matrix in meters
        mdistEOIS3 <- distm(isoscape_data.EOIS3.spdf)
        # cluster all points using a hierarchical clustering approach
        hcEOIS3 <- hclust(as.dist(mdistEOIS3), method="complete")
        # define the distance threshold, in this case 100km (100000m)
        d=100000
        # define clusters based on a tree "height" cutoff "d" and add them to the SpDataFrame
        isoscape_data.EOIS3.spdf$clust <- cutree(hcEOIS3, h=d)
        # convert back to df
        isoscape_data.EOIS3.clust <- as.data.frame(isoscape_data.EOIS3.spdf)

      # rejoin data frames
        final_isoscape_data <- rbind(isoscape_data.EH.clust, isoscape_data.EOIS3.clust, isoscape_data.GI1.clust,
isoscape_data.GS1.clust, isoscape_data.GS2.clust, isoscape_data.LGM.clust, isoscape_data.LOIS3.clust)

    ## SET UP DATA TO TEST AND PLOT SPECIES DIFFERENCES WITHIN GEOGRAPHICAL CLUSTERS ##

        # divide data by species, cluster and time bin, and retain only the groups that have 3 or more data points:
          isoscape_data_3.min <- final_isoscape_data  %>%
          dplyr::group_by(finalagebin, clust, Faunal.Category) %>%
          filter(n() >= 3)

        # then select clusters that have at 3 or more different species (for kruskall wallis tests):
          isoscape_data_3.spec <- isoscape_data_3.min %>%
          dplyr::group_by(finalagebin, clust) %>%
          filter(n_distinct(Faunal.Category) >= 3)

        # and then select clusters where there are two species (for Mann Whitney U tests)
          isoscape_data_2.spec <- isoscape_data_3.min %>%
          dplyr::group_by(finalagebin, clust) %>%
          filter(n_distinct(Faunal.Category) == 2)

        # select cases where within a cluster/age bracket there are AT LEAST two species
          isoscape_data_clust.min.2.spec <- isoscape_data_3.min%>%
          dplyr::group_by(finalagebin, clust) %>%
          filter(n_distinct(Faunal.Category) >= 2)

    ## SPECIES COMPARISONS BY CLUSTER (Table S2.2) ##

      # table of summary stats for all clusters
        cluster.summary.stats <- isoscape_data_clust.min.2.spec %>%
        dplyr::group_by(finalagebin, clust, Faunal.Category) %>%
        dplyr::summarize(Latitude = mean(Latitude),
                   Longitude = mean(Longitude),
                    mean = mean(d15Ncoll),
                   median = median(d15Ncoll),
                   stdev = sd(d15Ncoll),
                   min= min(d15Ncoll),
                   max= max(d15Ncoll),
                   n=n()) %>%
        ungroup()
      cluster.summary.stats

      # test for differences

        mann.whitney.output.spec <- isoscape_data_2.spec %>%
        group_by(finalagebin, clust) %>%
        group_modify(~ wilc_test(.x$d15Ncoll ~ .x$Faunal.Category))

        mann.whitney.output.spec['Test'] = 'Mann Whitney'

        kruskall.wallis.output.spec <- isoscape_data_3.spec %>%
        group_by(finalagebin, clust) %>%
        group_modify(~ krus_test(.x$d15Ncoll ~ .x$Faunal.Category))

        kruskall.wallis.output.spec['Test'] = 'Kruskall Wallis'

        species.comparison.table <- rbind(mann.whitney.output.spec, kruskall.wallis.output.spec)

        species.comparison.table <- species.comparison.table %>%
        mutate(sign = case_when(
        p.value < 0.05 ~ "*",
        p.value >= 0.05 ~ ""))

        species.comparison.table$finalagebin  <- factor(species.comparison.table$finalagebin,  levels = c("EH", "YD", "LGI", "LGT",
"LGM", "LOIS3", "EOIS3"))

        knitr::kable(species.comparison.table, digits = c(0, 0, 3, 1), col.names = gsub("[.]", " ",
names(species.comparison.table)))

        species.comparison.table <- as.data.frame(species.comparison.table)

      # combine two outputs and save file (creates Table S2.2)
        cluster.summary.stats.signif <- merge(cluster.summary.stats, species.comparison.table, by=c("finalagebin","clust"))
        write.csv(file="Table_S2_2.csv", cluster.summary.stats.signif)

  ##PLOT CLUSTERS (Figure S2.2) ##

  # create simple data frame showing significant and non-significant clusters to be used in plotting
  (clusters.for.plots <- cluster.summary.stats.signif %>%
  dplyr::group_by(finalagebin, clust, sign) %>%
  dplyr::summarize(Latitude = mean(Latitude),
                   Longitude = mean(Longitude)) %>%
```

```
  ungroup())

  ## EOIS3 plot ##
  (EOIS3clustmap <-
  ggplot()+
  geom_polygon(data=pcoastEOIS3.simp, aes(long, lat, group=group), fill="lightgrey") +
  geom_path(data=eurmed, aes(long, lat, group=group), colour="black") +
  geom_polygon(data=iceEOIS3, aes(long, lat, group=group), colour="black", fill="white") +
  geom_point(data = clusters.for.plots[clusters.for.plots$finalagebin == "EOIS3", ], aes(Longitude, Latitude, shape=factor(sign),
colour=factor(sign), fill=factor(sign), size=factor(sign))) +
  geom_label_repel(data = clusters.for.plots[clusters.for.plots$finalagebin == "EOIS3", ], aes(Longitude, Latitude, label=clust))+
  scale_shape_manual(values = c(21, 25), labels=c("no","yes"))+
  scale_colour_manual(values = c("black", "black"), labels=c("no","yes")) +
  scale_fill_manual(values = c("white", "black"), labels=c("no","yes")) +
  scale_size_manual(values= c(5,5), labels=c("no","yes")) +
  coord_cartesian(xlim = c(-10,25), ylim = c(35, 60)) +
  xlab("Longitude") +
  ylab("Latitude") +
  ggtitle("EOIS3")+
  theme_bw() +
  theme(legend.position="right") +
  labs(shape = "Signficant\nspecies-based\ndifference?") +
  labs(colour = "Signficant\nspecies-based\ndifference?") +
  labs(fill = "Signficant\nspecies-based\ndifference?") +
  labs(size = "Signficant\nspecies-based\ndifference?"))

  ## LOIS3 plot ##
  (LOIS3clustmap <- ggplot()+
  geom_polygon(data=pcoastLOIS3.simp, aes(long, lat, group=group), fill="lightgrey") +
  geom_path(data=eurmed, aes(long, lat, group=group), colour="black") +
  geom_polygon(data=iceLOIS3, aes(long, lat, group=group), colour="black", fill="white") +
  geom_point(data = clusters.for.plots[clusters.for.plots$finalagebin == "LOIS3", ], aes(Longitude, Latitude, shape=factor(sign),
colour=factor(sign), fill=factor(sign), size=factor(sign))) +
  geom_label_repel(data = clusters.for.plots[clusters.for.plots$finalagebin == "LOIS3", ], aes(Longitude, Latitude, label=clust))+
  scale_shape_manual(values = c(21, 25))+
  scale_colour_manual(values = c("black", "black")) +
  scale_fill_manual(values = c("white", "black")) +
  scale_size_manual(values= c(5,5)) +
  coord_cartesian(xlim = c(-10,25), ylim = c(35, 60)) +
  xlab("Longitude") +
  ylab("Latitude") +
  ggtitle("LOIS3")+
  theme_bw() +
  theme(legend.position="none"))

  ## LGM plot ##
  (LGMclustmap <- ggplot()+
  geom_polygon(data=pcoastLGM.simp, aes(long, lat, group=group), fill="lightgrey") +
  geom_path(data=eurmed, aes(long, lat, group=group), colour="black") +
  geom_polygon(data=iceLGM, aes(long, lat, group=group), colour="black", fill="white") +
  geom_point(data = clusters.for.plots[clusters.for.plots$finalagebin == "LGM", ], aes(Longitude, Latitude, shape=factor(sign),
colour=factor(sign), fill=factor(sign), size=factor(sign))) +
  geom_label_repel(data = clusters.for.plots[clusters.for.plots$finalagebin == "LGM", ], aes(Longitude, Latitude, label=clust))+
  scale_shape_manual(values = c(21, 25))+
  scale_colour_manual(values = c("black", "black")) +
  scale_fill_manual(values = c("white", "black")) +
  scale_size_manual(values= c(5,5)) +
  coord_cartesian(xlim = c(-10,25), ylim = c(35, 60)) +
  xlab("Longitude") +
  ylab("Latitude") +
  ggtitle("LGM")+
  theme_bw() +
  theme(legend.position="none"))

  ## LGT plot ##
  (GS2clustmap <- ggplot()+
  geom_polygon(data=pcoastGS2.simp, aes(long, lat, group=group), fill="lightgrey") +
  geom_path(data=eurmed, aes(long, lat, group=group), colour="black") +
  geom_polygon(data=iceGS2, aes(long, lat, group=group), colour="black", fill="white") +
  geom_point(data = clusters.for.plots[clusters.for.plots$finalagebin == "LGT", ], aes(Longitude, Latitude, shape=factor(sign),
colour=factor(sign), fill=factor(sign), size=factor(sign))) +
  geom_label_repel(data = clusters.for.plots[clusters.for.plots$finalagebin == "LGT", ], aes(Longitude, Latitude, label=clust))+
  scale_shape_manual(values = c(21, 25))+
  scale_colour_manual(values = c("black", "black")) +
  scale_fill_manual(values = c("white", "black")) +
  scale_size_manual(values= c(5,5)) +
  coord_cartesian(xlim = c(-10,25), ylim = c(35, 60)) +
  xlab("Longitude") +
  ylab("Latitude") +
  ggtitle("LGT")+
  theme_bw() +
  theme(legend.position="none"))


  ## LGI plot ##
  (GI1clustmap <- ggplot()+
  geom_polygon(data=pcoastGI1.simp, aes(long, lat, group=group), fill="lightgrey") +
  geom_path(data=eurmed, aes(long, lat, group=group), colour="black") +
  geom_polygon(data=iceGI1, aes(long, lat, group=group), colour="black", fill="white") +
  geom_point(data = clusters.for.plots[clusters.for.plots$finalagebin == "LGI", ], aes(Longitude, Latitude, shape=factor(sign),
colour=factor(sign), fill=factor(sign), size=factor(sign))) +
  geom_label_repel(data = clusters.for.plots[clusters.for.plots$finalagebin == "LGI", ], aes(Longitude, Latitude, label=clust))+
  scale_shape_manual(values = c(21, 25))+
  scale_colour_manual(values = c("black", "black")) +
  scale_fill_manual(values = c("white", "black")) +
```

```
  scale_size_manual(values= c(5,5)) +
  coord_cartesian(xlim = c(-10,25), ylim = c(35, 60)) +
  xlab("Longitude") +
  ylab("Latitude") +
  ggtitle("LGI")+
  theme_bw() +
  theme(legend.position="none"))

  ## YD plot ##
  (GS1clustmap <- ggplot()+
  geom_polygon(data=pcoastGS1.simp, aes(long, lat, group=group), fill="lightgrey") +
  geom_path(data=eurmed, aes(long, lat, group=group), colour="black") +
  geom_polygon(data=iceGS1, aes(long, lat, group=group), colour="black", fill="white") +
  geom_point(data = clusters.for.plots[clusters.for.plots$finalagebin == "YD", ], aes(Longitude, Latitude, shape=factor(sign),
colour=factor(sign), fill=factor(sign), size=factor(sign))) +
  geom_label_repel(data = clusters.for.plots[clusters.for.plots$finalagebin == "YD", ], aes(Longitude, Latitude, label=clust))+
  scale_shape_manual(values = c(21, 25))+
  scale_colour_manual(values = c("black", "black")) +
  scale_fill_manual(values = c("white", "black")) +
  scale_size_manual(values= c(5,5)) +
  coord_cartesian(xlim = c(-10,25), ylim = c(35, 60)) +
  xlab("Longitude") +
  ylab("Latitude") +
  ggtitle("YD")+
  theme_bw() +
  theme(legend.position="none"))

  ## EH plot ##
  (EHclustmap <- ggplot()+
  geom_polygon(data=pcoastEH.simp, aes(long, lat, group=group), fill="lightgrey") +
  geom_path(data=eurmed, aes(long, lat, group=group), colour="black") +
  geom_polygon(data=iceEH, aes(long, lat, group=group), colour="black", fill="white") +
  geom_point(data = clusters.for.plots[clusters.for.plots$finalagebin == "EH", ], aes(Longitude, Latitude, shape=factor(sign),
colour=factor(sign), fill=factor(sign), size=factor(sign))) +
  geom_label_repel(data = clusters.for.plots[clusters.for.plots$finalagebin == "EH", ], aes(Longitude, Latitude, label=clust))+
  scale_shape_manual(values = c(21, 25))+
  scale_colour_manual(values = c("black", "black")) +
  scale_fill_manual(values = c("white", "black")) +
  scale_size_manual(values= c(5,5)) +
  coord_cartesian(xlim = c(-10,25), ylim = c(35, 60)) +
  xlab("Longitude") +
  ylab("Latitude") +
  ggtitle("EH")+
  theme_bw() +
  theme(legend.position="none"))

# get legends
legend <- get_legend(EOIS3clustmap)
# remove legend from EOIS3clustmap
EOIS3clustmap <- EOIS3clustmap + theme(legend.position="none")
# combine and save plots

# combine plots ##
pdf(file="Figure_SI2_2.pdf", width = 10, height = 15)
grid.arrange(EHclustmap, GS1clustmap, GI1clustmap,
             GS2clustmap, LGMclustmap, LOIS3clustmap,
             EOIS3clustmap, legend, ncol=2, nrow=4)
dev.off()


  ## HORSE AND REINDEER COMPARISON (FIGURE S2.3)####

  # assemble clusters containing horse and reindeer:
    isoscape_data_3.spec.horren <- isoscape_data_3.spec %>%
      filter(Faunal.Category == 'Equus' | Faunal.Category == 'Rangifer')
    isoscape_data_2.spec.horren <- isoscape_data_2.spec %>%
      filter(Faunal.Category == 'Equus' | Faunal.Category == 'Rangifer')
    horse_reindeer <- rbind(isoscape_data_3.spec.horren,isoscape_data_2.spec.horren)
    horse_reindeer <- horse_reindeer %>%
      dplyr::group_by(finalagebin, clust) %>%
      filter(n_distinct(Faunal.Category) == 2)

  # define order of plots
    horse_reindeer$finalagebin  <- factor(horse_reindeer$finalagebin, levels = c("EH", "YD", "LGI", "LGT", "LGM", "LOIS3", "EOIS3"))

  # plot horse v reindeer
    col <- brewer.pal(n = 3, name = 'Dark2')
    (horse.reindeer.boxplot <- ggboxplot(horse_reindeer, x="Faunal.Category", y="d15Ncoll", fill="Faunal.Category") +
    facet_wrap(~finalagebin + clust, nrow=6, scales = "free") +
    scale_fill_manual(values=col, name = "Faunal\nCategory")+
    ylim(-1,11)+
    theme_bw()+
    ylab(expression(paste(delta^{15}, "N (\u2030)"))) +
    theme(text = element_text(size=16),
        axis.text.x = element_blank(),
        axis.title.x = element_blank()))

  # add labels to plots to show significant differences
    horse.reindeer.boxplot <- horse.reindeer.boxplot + stat_compare_means(aes(label = ..p.signif..), , size = 10, label.x = 1.5,
label.y = 8, colour = "red", hide.ns = TRUE)
    horse.reindeer.boxplot
pdf(file="Figure_SI2_3.pdf", width = 10, height = 12.5)
grid.draw(shift_legend(horse.reindeer.boxplot))
dev.off()
```

```
#### BASIC SUMMARY STATS FOR ISOTOPE DATA (Table 2 and Figure 1 ) ####

  ## Table 2
    data.summary.stats <- final_isoscape_data %>%
      dplyr::group_by(finalagebin) %>%
      dplyr::summarize(n=n(),
                    mean = round(mean(d15Ncoll), digits = 1),
                    stdev = round(sd(d15Ncoll), digits = 1),
                    median = median(d15Ncoll),
                    min= min(d15Ncoll),
                    max= max(d15Ncoll)) %>%
        ungroup()
    write.csv(data.summary.stats, file="summary.stats.csv")

  ## FIGURE 1
    # set order of plots
      final_isoscape_data$finalagebin  <- factor(final_isoscape_data$finalagebin, levels = c("EH", "YD", "LGI", "LGT", "LGM",
"LOIS3", "EOIS3"))
    # Set up a colour palette:
      cbbPalette <- c("#698B69", "#5D478B", "#5C5C5C", "#CD6090", "#EEC900", "#5F9EA0", "#6CA6CD")
    #assign colour to each timebin
      names(cbbPalette) <- levels(isoscape_data$finalagebin)
    # create plot
      (summary_boxplot <- ggplot() +
      geom_boxplot(data=isoscape_data, aes(x=finalagebin, y=d15Ncoll, fill=finalagebin)) +
      scale_fill_manual(values = cbbPalette) +
      theme_bw()+
      theme(text = element_text(size=10),
        axis.text.x = element_text(angle=90, hjust=1)) +
      theme(legend.position="none") +
      xlab("Time bin")+
      ylab(expression(paste(delta^{15}, "N (\u2030)"))))
    #save output as pdf
      ggsave(file="Figure_1.pdf", summary_boxplot, width = 20, height = 15, units = c("cm"), dpi = 300)



#### LOCAL AND GLOBAL MORAN I STATS BY TIME BIN (Table S3) ####

  ## set up dataframe
  data.spdf <- final_isoscape_data
  coordinates(data.spdf) <- ~Longitude+Latitude
  proj4string(data.spdf) <- latlon # add coordinate system info to isotope data

  ## EARLY HOLOCENE INTERSTADIAL LOCAL AND GLOBAL MORAN I ####
  EH <- data.spdf[data.spdf@data$finalagebin=="LGI",] # select data
  EH.jit <- spJitter(EH, 0.1) # spatially jitter data to avoid coincident points
  # create an inverse distance weights matrix:
  coords <- sp::coordinates(EH.jit) # get coordinates from spdf
  dist.mat <- as.matrix(dist(coords, method = "euclidean")) #create distance matrix
  # calcualte inverse:
  dist.mat.inve <- 1 / dist.mat # 1 / d_{ij}
  diag(dist.mat.inve) <- 0 # 0 in the diagonal
  dist.mat.inve[!is.finite(dist.mat.inve)] <- 1 # just in case coincident points still occur, give a weight of 1 to avoid zero
division
  # standardized inverse weigth matrix:
  dist.mat.inve.EH <- mat2listw(dist.mat.inve, style = "W")
  # calculate local moran I (clusters and outliers)
  resI <- localmoran(EH.jit$d15Ncoll, dist.mat.inve.EH, alternative = "two.sided")
  resIpadjust <- (p.adjust(resI[,5], method="fdr")) #adjut p value for false discovery rate
  # reassemble data frame:
  EH.localMI.jit <- cbind(EH.jit, resI)
  EH.localMI.jit.df <- as.data.frame(EH.localMI.jit)
  EH.localMI.jit.df <- cbind(EH.localMI.jit.df, "PValue" = resIpadjust)
  # divide data into quadrants for plotting clusters and outliers
  # center d15N around its mean
  cDV <- EH.jit@data$d15Ncoll - mean(EH.jit@data$d15Ncoll)
  # A positive value for Ii indicates that the unit is surrounded by units with similar values.
  quadrant <- vector(mode="numeric",length=nrow(EH.localMI.jit.df))
  quadrant[cDV>0 & EH.localMI.jit.df$Ii>0] <- 1
  quadrant[cDV<0 & EH.localMI.jit.df$Ii>0] <- 2
  quadrant[cDV>0 & EH.localMI.jit.df$Ii<0] <- 3
  quadrant[cDV<0 & EH.localMI.jit.df$Ii<0] <- 4
  # set a statistical significance level for the local Moran's
  signif <- 0.05
  # place data with non-significant Moran's in the category "5"
  quadrant[EH.localMI.jit.df$PValue > signif] <- 5
  EH.df.jit.moran <- cbind(EH.localMI.jit.df, cDV, quadrant)
  # remove outliers and create new IDW matrix for Global Moran
  EH.jit.data_for_interpolation <- EH.df.jit.moran %>%
    filter(quadrant != 3 & quadrant != 4)
  EH.jit.data_for_interpolation.spdf <- EH.jit.data_for_interpolation
  coordinates(EH.jit.data_for_interpolation.spdf) <- ~Longitude+Latitude
  proj4string(EH.jit.data_for_interpolation.spdf) <- latlon # add coordinate system info to isotope data
  coords <- sp::coordinates(EH.jit.data_for_interpolation.spdf) # get coordinates from spdf
  dist.mat <- as.matrix(dist(coords, method = "euclidean")) #create distance matrix
  # calcualte inverse:
  dist.mat.inve <- 1 / dist.mat # 1 / d_{ij}
  diag(dist.mat.inve) <- 0 # 0 in the diagonal
  dist.mat.inve[!is.finite(dist.mat.inve)] <- 1
  # standardized inverse weigth matrix:
  dist.mat.inve.EH <- mat2listw(dist.mat.inve, style = "W")
  # calculate global moran I
  EH.gm.jit<- moran.test(EH.jit.data_for_interpolation$d15Ncoll, listw = dist.mat.inve.EH,
```

```
                          alternative = 'two.sided')
  EH.gm.jit.df <- tidy(EH.gm.jit)
  ## YOUNGER DRAYS INTERSTADIAL LOCAL AND GLOBAL MORAN I ####
  GS1 <- data.spdf[data.spdf@data$finalagebin=="LGI",] # select data
  GS1.jit <- spJitter(GS1, 0.1) # spatially jitter data to avoid coincident points
  # create an inverse distance weights matrix:
  coords <- sp::coordinates(GS1.jit) # get coordinates from spdf
  dist.mat <- as.matrix(dist(coords, method = "euclidean")) #create distance matrix
  # calcualte inverse:
  dist.mat.inve <- 1 / dist.mat # 1 / d_{ij}
  diag(dist.mat.inve) <- 0 # 0 in the diagonal
  dist.mat.inve[!is.finite(dist.mat.inve)] <- 1 # just in case coincident points still occur, give a weight of 1 to avoid zero
division
  # standardized inverse weigth matrix:
  dist.mat.inve.GS1 <- mat2listw(dist.mat.inve, style = "W")
  # calculate local moran I (clusters and outliers)
  resI <- localmoran(GS1.jit$d15Ncoll, dist.mat.inve.GS1, alternative = "two.sided")
  resIpadjust <- (p.adjust(resI[,5], method="fdr")) #adjut p value for false discovery rate
  # reassemble data frame:
  GS1.localMI.jit <- cbind(GS1.jit, resI)
  GS1.localMI.jit.df <- as.data.frame(GS1.localMI.jit)
  GS1.localMI.jit.df <- cbind(GS1.localMI.jit.df, "PValue" = resIpadjust)
  # divide data into quadrants for plotting clusters and outliers
  # center d15N around its mean
  cDV <- GS1.jit@data$d15Ncoll - mean(GS1.jit@data$d15Ncoll)
  # A positive value for Ii indicates that the unit is surrounded by units with similar values.
  quadrant <- vector(mode="numeric",length=nrow(GS1.localMI.jit.df))
  quadrant[cDV>0 & GS1.localMI.jit.df$Ii>0] <- 1
  quadrant[cDV<0 & GS1.localMI.jit.df$Ii>0] <- 2
  quadrant[cDV>0 & GS1.localMI.jit.df$Ii<0] <- 3
  quadrant[cDV<0 & GS1.localMI.jit.df$Ii<0] <- 4
  # set a statistical significance level for the local Moran's
  signif <- 0.05
  # place data with non-significant Moran's in the category "5"
  quadrant[GS1.localMI.jit.df$PValue > signif] <- 5
  GS1.df.jit.moran <- cbind(GS1.localMI.jit.df, cDV, quadrant)
  # remove outliers and create new IDW matrix for Global Moran
  GS1.jit.data_for_interpolation <- GS1.df.jit.moran %>%
    filter(quadrant != 3 & quadrant != 4)
  GS1.jit.data_for_interpolation.spdf <- GS1.jit.data_for_interpolation
  coordinates(GS1.jit.data_for_interpolation.spdf) <- ~Longitude+Latitude
  proj4string(GS1.jit.data_for_interpolation.spdf) <- latlon # add coordinate system info to isotope data
  coords <- sp::coordinates(GS1.jit.data_for_interpolation.spdf) # get coordinates from spdf
  dist.mat <- as.matrix(dist(coords, method = "euclidean")) #create distance matrix
  # calcualte inverse:
  dist.mat.inve <- 1 / dist.mat # 1 / d_{ij}
  diag(dist.mat.inve) <- 0 # 0 in the diagonal
  dist.mat.inve[!is.finite(dist.mat.inve)] <- 1
  # standardized inverse weigth matrix:
  dist.mat.inve.GS1 <- mat2listw(dist.mat.inve, style = "W")
  # calculate global moran I
  GS1.gm.jit<- moran.test(GS1.jit.data_for_interpolation$d15Ncoll, listw = dist.mat.inve.GS1,
                           alternative = 'two.sided')
  GS1.gm.jit.df <- tidy(GS1.gm.jit)
  ## LATE GLACIAL INTERSTADIAL LOCAL AND GLOBAL MORAN I ####
  GI1 <- data.spdf[data.spdf@data$finalagebin=="LGI",] # select data
  GI1.jit <- spJitter(GI1, 0.1) # spatially jitter data to avoid coincident points
  # create an inverse distance weights matrix:
  coords <- sp::coordinates(GI1.jit) # get coordinates from spdf
  dist.mat <- as.matrix(dist(coords, method = "euclidean")) #create distance matrix
  # calcualte inverse:
  dist.mat.inve <- 1 / dist.mat # 1 / d_{ij}
  diag(dist.mat.inve) <- 0 # 0 in the diagonal
  dist.mat.inve[!is.finite(dist.mat.inve)] <- 1 # just in case coincident points still occur, give a weight of 1 to avoid zero
division
  # standardized inverse weigth matrix:
  dist.mat.inve.GI1 <- mat2listw(dist.mat.inve, style = "W")
  # calculate local moran I (clusters and outliers)
  resI <- localmoran(GI1.jit$d15Ncoll, dist.mat.inve.GI1, alternative = "two.sided")
  resIpadjust <- (p.adjust(resI[,5], method="fdr")) #adjut p value for false discovery rate
  # reassemble data frame:
  GI1.localMI.jit <- cbind(GI1.jit, resI)
  GI1.localMI.jit.df <- as.data.frame(GI1.localMI.jit)
  GI1.localMI.jit.df <- cbind(GI1.localMI.jit.df, "PValue" = resIpadjust)
  # divide data into quadrants for plotting clusters and outliers
  # center d15N around its mean
  cDV <- GI1.jit@data$d15Ncoll - mean(GI1.jit@data$d15Ncoll)
  # A positive value for Ii indicates that the unit is surrounded by units with similar values.
  quadrant <- vector(mode="numeric",length=nrow(GI1.localMI.jit.df))
  quadrant[cDV>0 & GI1.localMI.jit.df$Ii>0] <- 1
  quadrant[cDV<0 & GI1.localMI.jit.df$Ii>0] <- 2
  quadrant[cDV>0 & GI1.localMI.jit.df$Ii<0] <- 3
  quadrant[cDV<0 & GI1.localMI.jit.df$Ii<0] <- 4
  # set a statistical significance level for the local Moran's
  signif <- 0.05
  # place data with non-significant Moran's in the category "5"
  quadrant[GI1.localMI.jit.df$PValue > signif] <- 5
  GI1.df.jit.moran <- cbind(GI1.localMI.jit.df, cDV, quadrant)
  # remove outliers and create new IDW matrix for Global Moran
  GI1.jit.data_for_interpolation <- GI1.df.jit.moran %>%
    filter(quadrant != 3 & quadrant != 4)
  GI1.jit.data_for_interpolation.spdf <- GI1.jit.data_for_interpolation
  coordinates(GI1.jit.data_for_interpolation.spdf) <- ~Longitude+Latitude
  proj4string(GI1.jit.data_for_interpolation.spdf) <- latlon # add coordinate system info to isotope data
  coords <- sp::coordinates(GI1.jit.data_for_interpolation.spdf) # get coordinates from spdf
```

```
  dist.mat <- as.matrix(dist(coords, method = "euclidean")) #create distance matrix
  # calcualte inverse:
  dist.mat.inve <- 1 / dist.mat # 1 / d_{ij}
  diag(dist.mat.inve) <- 0 # 0 in the diagonal
  dist.mat.inve[!is.finite(dist.mat.inve)] <- 1
  # standardized inverse weigth matrix:
  dist.mat.inve.GI1 <- mat2listw(dist.mat.inve, style = "W")
  # calculate global moran I
  GI1.gm.jit<- moran.test(GI1.jit.data_for_interpolation$d15Ncoll, listw = dist.mat.inve.GI1,
                          alternative = 'two.sided')
  GI1.gm.jit.df <- tidy(GI1.gm.jit)
  ## LAST GLACIAL TERMINATION LOCAL AND GLOBAL MORAN I ####
  GS2 <- data.spdf[data.spdf@data$finalagebin=="LGI",] # select data
  GS2.jit <- spJitter(GS2, 0.1) # spatially jitter data to avoid coincident points
  # create an inverse distance weights matrix:
  coords <- sp::coordinates(GS2.jit) # get coordinates from spdf
  dist.mat <- as.matrix(dist(coords, method = "euclidean")) #create distance matrix
  # calcualte inverse:
  dist.mat.inve <- 1 / dist.mat # 1 / d_{ij}
  diag(dist.mat.inve) <- 0 # 0 in the diagonal
  dist.mat.inve[!is.finite(dist.mat.inve)] <- 1 # just in case coincident points still occur, give a weight of 1 to avoid zero
division
  # standardized inverse weigth matrix:
  dist.mat.inve.GS2 <- mat2listw(dist.mat.inve, style = "W")
  # calculate local moran I (clusters and outliers)
  resI <- localmoran(GS2.jit$d15Ncoll, dist.mat.inve.GS2, alternative = "two.sided")
  resIpadjust <- (p.adjust(resI[,5], method="fdr")) #adjut p value for false discovery rate
  # reassemble data frame:
  GS2.localMI.jit <- cbind(GS2.jit, resI)
  GS2.localMI.jit.df <- as.data.frame(GS2.localMI.jit)
  GS2.localMI.jit.df <- cbind(GS2.localMI.jit.df, "PValue" = resIpadjust)
  # divide data into quadrants for plotting clusters and outliers
  # center d15N around its mean
  cDV <- GS2.jit@data$d15Ncoll - mean(GS2.jit@data$d15Ncoll)
  # A positive value for Ii indicates that the unit is surrounded by units with similar values.
  quadrant <- vector(mode="numeric",length=nrow(GS2.localMI.jit.df))
  quadrant[cDV>0 & GS2.localMI.jit.df$Ii>0] <- 1
  quadrant[cDV<0 & GS2.localMI.jit.df$Ii>0] <- 2
  quadrant[cDV>0 & GS2.localMI.jit.df$Ii<0] <- 3
  quadrant[cDV<0 & GS2.localMI.jit.df$Ii<0] <- 4
  # set a statistical significance level for the local Moran's
  signif <- 0.05
  # place data with non-significant Moran's in the category "5"
  quadrant[GS2.localMI.jit.df$PValue > signif] <- 5
  GS2.df.jit.moran <- cbind(GS2.localMI.jit.df, cDV, quadrant)
  # remove outliers and create new IDW matrix for Global Moran
  GS2.jit.data_for_interpolation <- GS2.df.jit.moran %>%
    filter(quadrant != 3 & quadrant != 4)
  GS2.jit.data_for_interpolation.spdf <- GS2.jit.data_for_interpolation
  coordinates(GS2.jit.data_for_interpolation.spdf) <- ~Longitude+Latitude
  proj4string(GS2.jit.data_for_interpolation.spdf) <- latlon # add coordinate system info to isotope data
  coords <- sp::coordinates(GS2.jit.data_for_interpolation.spdf) # get coordinates from spdf
  dist.mat <- as.matrix(dist(coords, method = "euclidean")) #create distance matrix
  # calcualte inverse:
  dist.mat.inve <- 1 / dist.mat # 1 / d_{ij}
  diag(dist.mat.inve) <- 0 # 0 in the diagonal
  dist.mat.inve[!is.finite(dist.mat.inve)] <- 1
  # standardized inverse weigth matrix:
  dist.mat.inve.GS2 <- mat2listw(dist.mat.inve, style = "W")
  # calculate global moran I
  GS2.gm.jit<- moran.test(GS2.jit.data_for_interpolation$d15Ncoll, listw = dist.mat.inve.GS2,
                          alternative = 'two.sided')
  GS2.gm.jit.df <- tidy(GS2.gm.jit)
  ## LAST GLACIAL MAXIMUM LOCAL AND GLOBAL MORAN I ####
  LGM <- data.spdf[data.spdf@data$finalagebin=="LGI",] # select data
  LGM.jit <- spJitter(LGM, 0.1) # spatially jitter data to avoid coincident points
  # create an inverse distance weights matrix:
  coords <- sp::coordinates(LGM.jit) # get coordinates from spdf
  dist.mat <- as.matrix(dist(coords, method = "euclidean")) #create distance matrix
  # calcualte inverse:
  dist.mat.inve <- 1 / dist.mat # 1 / d_{ij}
  diag(dist.mat.inve) <- 0 # 0 in the diagonal
  dist.mat.inve[!is.finite(dist.mat.inve)] <- 1 # just in case coincident points still occur, give a weight of 1 to avoid zero
division
  # standardized inverse weigth matrix:
  dist.mat.inve.LGM <- mat2listw(dist.mat.inve, style = "W")
  # calculate local moran I (clusters and outliers)
  resI <- localmoran(LGM.jit$d15Ncoll, dist.mat.inve.LGM, alternative = "two.sided")
  resIpadjust <- (p.adjust(resI[,5], method="fdr")) #adjut p value for false discovery rate
  # reassemble data frame:
  LGM.localMI.jit <- cbind(LGM.jit, resI)
  LGM.localMI.jit.df <- as.data.frame(LGM.localMI.jit)
  LGM.localMI.jit.df <- cbind(LGM.localMI.jit.df, "PValue" = resIpadjust)
  # divide data into quadrants for plotting clusters and outliers
  # center d15N around its mean
  cDV <- LGM.jit@data$d15Ncoll - mean(LGM.jit@data$d15Ncoll)
  # A positive value for Ii indicates that the unit is surrounded by units with similar values.
  quadrant <- vector(mode="numeric",length=nrow(LGM.localMI.jit.df))
  quadrant[cDV>0 & LGM.localMI.jit.df$Ii>0] <- 1
  quadrant[cDV<0 & LGM.localMI.jit.df$Ii>0] <- 2
  quadrant[cDV>0 & LGM.localMI.jit.df$Ii<0] <- 3
  quadrant[cDV<0 & LGM.localMI.jit.df$Ii<0] <- 4
  # set a statistical significance level for the local Moran's
  signif <- 0.05
  # place data with non-significant Moran's in the category "5"
```

```
  quadrant[LGM.localMI.jit.df$PValue > signif] <- 5
  LGM.df.jit.moran <- cbind(LGM.localMI.jit.df, cDV, quadrant)
  # remove outliers and create new IDW matrix for Global Moran
  LGM.jit.data_for_interpolation <- LGM.df.jit.moran %>%
    filter(quadrant != 3 & quadrant != 4)
  LGM.jit.data_for_interpolation.spdf <- LGM.jit.data_for_interpolation
  coordinates(LGM.jit.data_for_interpolation.spdf) <- ~Longitude+Latitude
  proj4string(LGM.jit.data_for_interpolation.spdf) <- latlon # add coordinate system info to isotope data
  coords <- sp::coordinates(LGM.jit.data_for_interpolation.spdf) # get coordinates from spdf
  dist.mat <- as.matrix(dist(coords, method = "euclidean")) #create distance matrix
  # calcualte inverse:
  dist.mat.inve <- 1 / dist.mat # 1 / d_{ij}
  diag(dist.mat.inve) <- 0 # 0 in the diagonal
  dist.mat.inve[!is.finite(dist.mat.inve)] <- 1
  # standardized inverse weigth matrix:
  dist.mat.inve.LGM <- mat2listw(dist.mat.inve, style = "W")
  # calculate global moran I
  LGM.gm.jit<- moran.test(LGM.jit.data_for_interpolation$d15Ncoll, listw = dist.mat.inve.LGM,
                          alternative = 'two.sided')
  LGM.gm.jit.df <- tidy(LGM.gm.jit)
  ## LATE OIS 3 LOCAL AND GLOBAL MORAN I ####
  LOIS3 <- data.spdf[data.spdf@data$finalagebin=="LGI",] # select data
  LOIS3.jit <- spJitter(LOIS3, 0.1) # spatially jitter data to avoid coincident points
  # create an inverse distance weights matrix:
  coords <- sp::coordinates(LOIS3.jit) # get coordinates from spdf
  dist.mat <- as.matrix(dist(coords, method = "euclidean")) #create distance matrix
  # calcualte inverse:
  dist.mat.inve <- 1 / dist.mat # 1 / d_{ij}
  diag(dist.mat.inve) <- 0 # 0 in the diagonal
  dist.mat.inve[!is.finite(dist.mat.inve)] <- 1 # just in case coincident points still occur, give a weight of 1 to avoid zero
division
  # standardized inverse weigth matrix:
  dist.mat.inve.LOIS3 <- mat2listw(dist.mat.inve, style = "W")
  # calculate local moran I (clusters and outliers)
  resI <- localmoran(LOIS3.jit$d15Ncoll, dist.mat.inve.LOIS3, alternative = "two.sided")
  resIpadjust <- (p.adjust(resI[,5], method="fdr")) #adjut p value for false discovery rate
  # reassemble data frame:
  LOIS3.localMI.jit <- cbind(LOIS3.jit, resI)
  LOIS3.localMI.jit.df <- as.data.frame(LOIS3.localMI.jit)
  LOIS3.localMI.jit.df <- cbind(LOIS3.localMI.jit.df, "PValue" = resIpadjust)
  # divide data into quadrants for plotting clusters and outliers
  # center d15N around its mean
  cDV <- LOIS3.jit@data$d15Ncoll - mean(LOIS3.jit@data$d15Ncoll)
  # A positive value for Ii indicates that the unit is surrounded by units with similar values.
  quadrant <- vector(mode="numeric",length=nrow(LOIS3.localMI.jit.df))
  quadrant[cDV>0 & LOIS3.localMI.jit.df$Ii>0] <- 1
  quadrant[cDV<0 & LOIS3.localMI.jit.df$Ii>0] <- 2
  quadrant[cDV>0 & LOIS3.localMI.jit.df$Ii<0] <- 3
  quadrant[cDV<0 & LOIS3.localMI.jit.df$Ii<0] <- 4
  # set a statistical significance level for the local Moran's
  signif <- 0.05
  # place data with non-significant Moran's in the category "5"
  quadrant[LOIS3.localMI.jit.df$PValue > signif] <- 5
  LOIS3.df.jit.moran <- cbind(LOIS3.localMI.jit.df, cDV, quadrant)
  # remove outliers and create new IDW matrix for Global Moran
  LOIS3.jit.data_for_interpolation <- LOIS3.df.jit.moran %>%
    filter(quadrant != 3 & quadrant != 4)
  LOIS3.jit.data_for_interpolation.spdf <- LOIS3.jit.data_for_interpolation
  coordinates(LOIS3.jit.data_for_interpolation.spdf) <- ~Longitude+Latitude
  proj4string(LOIS3.jit.data_for_interpolation.spdf) <- latlon # add coordinate system info to isotope data
  coords <- sp::coordinates(LOIS3.jit.data_for_interpolation.spdf) # get coordinates from spdf
  dist.mat <- as.matrix(dist(coords, method = "euclidean")) #create distance matrix
  # calcualte inverse:
  dist.mat.inve <- 1 / dist.mat # 1 / d_{ij}
  diag(dist.mat.inve) <- 0 # 0 in the diagonal
  dist.mat.inve[!is.finite(dist.mat.inve)] <- 1
  # standardized inverse weigth matrix:
  dist.mat.inve.LOIS3 <- mat2listw(dist.mat.inve, style = "W")
  # calculate global moran I
  LOIS3.gm.jit<- moran.test(LOIS3.jit.data_for_interpolation$d15Ncoll, listw = dist.mat.inve.LOIS3,
                            alternative = 'two.sided')
  LOIS3.gm.jit.df <- tidy(LOIS3.gm.jit)

  ## EARLY OIS 3 LOCAL AND GLOBAL MORAN I ####
  EOIS3 <- data.spdf[data.spdf@data$finalagebin=="LGI",] # select data
  EOIS3.jit <- spJitter(EOIS3, 0.1) # spatially jitter data to avoid coincident points
  # create an inverse distance weights matrix:
  coords <- sp::coordinates(EOIS3.jit) # get coordinates from spdf
  dist.mat <- as.matrix(dist(coords, method = "euclidean")) #create distance matrix
  # calcualte inverse:
  dist.mat.inve <- 1 / dist.mat # 1 / d_{ij}
  diag(dist.mat.inve) <- 0 # 0 in the diagonal
  dist.mat.inve[!is.finite(dist.mat.inve)] <- 1 # just in case coincident points still occur, give a weight of 1 to avoid zero
division
  # standardized inverse weigth matrix:
  dist.mat.inve.EOIS3 <- mat2listw(dist.mat.inve, style = "W")
  # calculate local moran I (clusters and outliers)
  resI <- localmoran(EOIS3.jit$d15Ncoll, dist.mat.inve.EOIS3, alternative = "two.sided")
  resIpadjust <- (p.adjust(resI[,5], method="fdr")) #adjut p value for false discovery rate
  # reassemble data frame:
  EOIS3.localMI.jit <- cbind(EOIS3.jit, resI)
  EOIS3.localMI.jit.df <- as.data.frame(EOIS3.localMI.jit)
  EOIS3.localMI.jit.df <- cbind(EOIS3.localMI.jit.df, "PValue" = resIpadjust)
  # divide data into quadrants for plotting clusters and outliers
  # center d15N around its mean
```

```
  cDV <- EOIS3.jit@data$d15Ncoll - mean(EOIS3.jit@data$d15Ncoll)
  # A positive value for Ii indicates that the unit is surrounded by units with similar values.
  quadrant <- vector(mode="numeric",length=nrow(EOIS3.localMI.jit.df))
  quadrant[cDV>0 & EOIS3.localMI.jit.df$Ii>0] <- 1
  quadrant[cDV<0 & EOIS3.localMI.jit.df$Ii>0] <- 2
  quadrant[cDV>0 & EOIS3.localMI.jit.df$Ii<0] <- 3
  quadrant[cDV<0 & EOIS3.localMI.jit.df$Ii<0] <- 4
  # set a statistical significance level for the local Moran's
  signif <- 0.05
  # place data with non-significant Moran's in the category "5"
  quadrant[EOIS3.localMI.jit.df$PValue > signif] <- 5
  EOIS3.df.jit.moran <- cbind(EOIS3.localMI.jit.df, cDV, quadrant)
  # remove outliers and create new IDW matrix for Global Moran
  EOIS3.jit.data_for_interpolation <- EOIS3.df.jit.moran %>%
    filter(quadrant != 3 & quadrant != 4)
  EOIS3.jit.data_for_interpolation.spdf <- EOIS3.jit.data_for_interpolation
  coordinates(EOIS3.jit.data_for_interpolation.spdf) <- ~Longitude+Latitude
  proj4string(EOIS3.jit.data_for_interpolation.spdf) <- latlon # add coordinate system info to isotope data
  coords <- sp::coordinates(EOIS3.jit.data_for_interpolation.spdf) # get coordinates from spdf
  dist.mat <- as.matrix(dist(coords, method = "euclidean")) #create distance matrix
  # calcualte inverse:
  dist.mat.inve <- 1 / dist.mat # 1 / d_{ij}
  diag(dist.mat.inve) <- 0 # 0 in the diagonal
  dist.mat.inve[!is.finite(dist.mat.inve)] <- 1
  # standardized inverse weigth matrix:
  dist.mat.inve.EOIS3 <- mat2listw(dist.mat.inve, style = "W")
  # calculate global moran I
  EOIS3.gm.jit<- moran.test(EOIS3.jit.data_for_interpolation$d15Ncoll, listw = dist.mat.inve.EOIS3,
                            alternative = 'two.sided')
  EOIS3.gm.jit.df <- tidy(EOIS3.gm.jit)




  ## COMBINE LOCAL AND GLOABL MORAN OUTPUTS FOR ALL TIME BINS (DATA FOR TABLE S3.1) ####
  # NOTE:
    # BECAUSE THE SPATIAL JITTERING UNDERTAKEN TO AVOID COINCIDENT SAMPLE POINTS IS RANDOMISED, THE JITTERING IS DIFFERENT EACH TIME
THE CODE IS RUN
    # THIS MEANS THAT THE NUMBER OF OUTLIERS IDENTIFIED AND REMOVED MAY VARY SLIGHTLY, WITH KNOCK ON EFFECT FOR CALCULATION OF
GLOBAL MORAN I AND FOR HOW CLUSTERS AND OUTLIERS APPEAR IN FIGURE 2
    # THE OUTLIERS IDENTIFED IN OUR ANALYSIS ARE INDICATED IN S1 DATASET IN COLUMN 'OUTLIER'
    # DIFFERENCES ARE MINOR AND DO NOT SIGNIFANTLY ALTER SUBSEQUENT INTERPOLATION
  # LOCAL MORAN I RESULTS FOR ALL TIME BINS #
  local.moran.results.jit <- rbind(EH.df.jit.moran, GS1.df.jit.moran, GI1.df.jit.moran, GS2.df.jit.moran, LGM.df.jit.moran,
LOIS3.df.jit.moran, EOIS3.df.jit.moran)
  # GLOBAL MORAN I RESULTS FOR ALL TIME BINS #
  global.moran.results.jit <- rbind(EH.gm.jit.df, GS1.gm.jit.df, GI1.gm.jit.df, GS2.gm.jit.df, LGM.gm.jit.df, LOIS3.gm.jit.df,
EOIS3.gm.jit.df)

  ## PLOT CLUSTERS AND OUTLIERS (FIGURE 2) ####
  # NOTE:
    # BECAUSE THE SPATIAL JITTERING UNDERTAKEN TO AVOID COINCIDENT SAMPLE POINTS IS RANDOMISED, THE JITTERING IS DIFFERENT EACH TIME
THE CODE IS RUN
    # THIS MEANS THAT THE NUMBER OF OUTLIERS IDENTIFIED AND REMOVED MAY VARY SLIGHTLY, WITH KNOCK ON EFFECT FOR CALCULATION OF
GLOBAL MORAN I AND FOR HOW CLUSTERS AND OUTLIERS APPEAR IN FIGURE 2
    # THE OUTLIERS IDENTIFED IN OUR ANALYSIS ARE INDICATED IN S1 DATASET IN COLUMN 'OUTLIER'
    # DIFFERENCES ARE MINOR AND DO NOT SIGNIFANTLY ALTER SUBSEQUENT INTERPOLATION

  # define order of plots
  local.moran.results.jit$finalagebin  <- factor(local.moran.results.jit$finalagebin, levels = c("EH", "YD", "LGI", "LGT", "LGM",
"LOIS3", "EOIS3"))

  #set qudrant as factor
  local.moran.results.jit <- local.moran.results.jit %>%
    filter(quadrant != 0)
  local.moran.results.jit$quadrant <- factor(local.moran.results.jit$quadrant, levels = c(5, 1, 2, 3, 4))

  (cluster.plot <- ggplot() +
      geom_path(data=eurmed, aes(long, lat, group=group), colour="grey34") +
      geom_point(data=local.moran.results.jit, aes(x=Longitude, y=Latitude, color=quadrant, size=quadrant, shape=quadrant,
fill=quadrant)) +
      scale_color_manual(name = "Group",
                         labels = c("not signficant", "cluster: high", "cluster: low", "outlier: high", "outlier: low"),
                         values = c("black", "red", "blue", "red", "blue")) +
      scale_fill_manual(name = "Group",
                        labels = c("not signficant", "cluster: high", "cluster: low", "outlier: high", "outlier: low"),
                        values = c("black", "lightpink", "skyblue2", "red", "blue")) +
      scale_size_manual(name = "Group",
                        labels = c("not signficant", "cluster: high", "cluster: low", "outlier: high", "outlier: low"),
                        values = c(2, 4, 4, 4, 4)) +
      scale_shape_manual(name = "Group",
                         labels = c("not signficant", "cluster: high", "cluster: low", "outlier: high", "outlier: low"),
                         values = c(16, 21, 21, 8, 8)) +
      facet_wrap(~ finalagebin, ncol=2) +
      xlim(-10,25) + ylim(35,60) +
      xlab("Longitude") + ylab("Latitude") +
      theme_bw())


  pdf(file="Figure_2.pdf", width = 10, height = 15)
  grid.draw(shift_legend(cluster.plot))
  dev.off()
```

```
  png(file="Figure_2.png", width = 20, height = 30, units="cm", res=300)
  grid.draw(shift_legend(cluster.plot))
  dev.off()

#### INTERPOLATION ####
  # FOR INTREPOLATION HERE WE USE THE s1 DATASET FILTERED BY 'OUTLIER'
    isoscape_data <- final_isoscape_data %>%
      filter(Outlier. == "n")
  # keep only the relevant columns
    isoscape_data <- final_isoscape_data %>%
    subset(select = c("Site", "Latitude", "Longitude", "Faunal.Category", "finalagebin", "d15Ncoll", "elevation","MAP", "MAT",
                    "precip.cold.quart", "precip.dry.month", "precip.dry.quar", "precip.warm.quart",
                    "precip.wet.month", "precip.wet.quar", "precipseason", "tempcoldest",
                    "tempdriest", "tempmax", "tempmin", "temprange",
                    "tempseason", "tempwarmest","tempwettest"))

  # remove data with NAs (there shouldn't be any, but just in case)
    isoscape_data <- na.omit(isoscape_data)

  # check structure of df and ammend as necessary
    str(isoscape_data)
    isoscape_data <- isoscape_data %>% rename(SiteName = Site)
    isoscape_data$SiteName <- as.factor(isoscape_data$SiteName) #change columns as needed
    isoscape_data$FaunalCatCombi <- as.factor(isoscape_data$FaunalCatCombi) #change columns as needed
    isoscape_data$finalagebin <- as.factor(isoscape_data$finalagebin) #change columns as needed
    isoscape_data$elevation <- as.numeric(isoscape_data$elevation)

  # create dataframe for site-averaged data (calculate average of environmental covariates at each location):
    d15N.data.aggre <- isoscape_data %>%
    dplyr::group_by(SiteName, finalagebin, add=TRUE) %>%
    dplyr::summarise(n_source_value = n(),
                   mean_source_value = mean(d15Ncoll),
                   var_source_value = var(d15Ncoll),
                   elevation = mean(elevation),
                   lat = mean(Latitude),
                   long = mean(Longitude),
                   MAT = mean(MAT),
                   MAP = mean(MAP),
                   precip.wet.quar = mean(precip.wet.quar),
                   tempdriest = mean(tempdriest),
                   precip.cold.quart = mean(precip.cold.quart),
                   precip.dry.month = mean(precip.dry.month),
                   precip.dry.quar = mean(precip.dry.quar),
                   precip.warm.quart = mean(precip.warm.quart),
                   precip.wet.month = mean(precip.wet.month),
                   precipseason = mean(precipseason),
                   tempcoldest = mean(tempcoldest),
                   tempmax = mean(tempmax),
                   tempmin = mean(tempmin),
                   temprange = mean(temprange),
                   tempseason = mean(tempseason),
                   tempwarmest = mean(tempwarmest),
                   tempwettest = mean(tempwettest)) %>%
  ungroup()

  # set var_source_value to NA where it equals zero so code works
    d15N.data.aggre$var_source_value[d15N.data.aggre$var_source_value == 0] <- NA

  # split df by age bin
    EH <- d15N.data.aggre %>% filter(finalagebin == "EH")
    GS1 <- d15N.data.aggre %>% filter(finalagebin == "YD")
    GI1 <- d15N.data.aggre %>% filter(finalagebin == "LGI")
    GS2 <- d15N.data.aggre %>% filter(finalagebin == "LGT")
    LGM <- d15N.data.aggre %>% filter(finalagebin == "LGM")
    LOIS3 <- d15N.data.aggre %>% filter(finalagebin == "LOIS3")
    EOIS3 <- d15N.data.aggre %>% filter(finalagebin == "EOIS3")

  ## MIXED MODEL INTERPOLATION - NO FIXED EFFECTS ####

  # Fit the residual dispersion model (following Courtiol and Rousset 2017) #
    EH.dispfit <- fitme(formula = var_source_value ~ 1 + Matern(1|long + lat) + (1|SiteName),
                    family = Gamma(link = log), data = EH, fixed = list(phi = 2),
                    prior.weights = n_source_value - 1, control.dist = list(dist.method = "Earth"), method = "REML")
    GS1.dispfit <- fitme(formula = var_source_value ~ 1 + Matern(1|long + lat) + (1|SiteName),
                     family = Gamma(link = log), data = GS1, fixed = list(phi = 2),
                     prior.weights = n_source_value - 1, control.dist = list(dist.method = "Earth"), method = "REML")
    GI1.dispfit <- fitme(formula = var_source_value ~ 1 + Matern(1|long + lat) + (1|SiteName),
                     family = Gamma(link = log), data = GI1, fixed = list(phi = 2),
                     prior.weights = n_source_value - 1, control.dist = list(dist.method = "Earth"), method = "REML")
    GS2.dispfit <- fitme(formula = var_source_value ~ 1 + Matern(1|long + lat) + (1|SiteName),
                     family = Gamma(link = log), data = GS2, fixed = list(phi = 2),
                     prior.weights = n_source_value - 1, control.dist = list(dist.method = "Earth"), method = "REML")
    LGM.dispfit <- fitme(formula = var_source_value ~ 1 + Matern(1|long + lat) + (1|SiteName),
                     family = Gamma(link = log), data = LGM, fixed = list(phi = 2),
                     prior.weights = n_source_value - 1, control.dist = list(dist.method = "Earth"), method = "REML")
    LOIS3.dispfit <- fitme(formula = var_source_value ~ 1 + Matern(1|long + lat) + (1|SiteName),
                       family = Gamma(link = log), data = LOIS3, fixed = list(phi = 2),
                       prior.weights = n_source_value - 1, control.dist = list(dist.method = "Earth"), method = "REML")
    EOIS3.dispfit <- fitme(formula = var_source_value ~ 1 + Matern(1|long + lat) + (1|SiteName),
                       family = Gamma(link = log), data = EOIS3, fixed = list(phi = 2),
                       prior.weights = n_source_value - 1, control.dist = list(dist.method = "Earth"), method = "REML")

    # predict Ïg of the expected square of the residual error in each location using the fit of the residual dispersion model:
    EH$disp <- predict(EH.dispfit, newdata = EH)[,1]
    GS1$disp <- predict(GS1.dispfit, newdata = GS1)[,1]
```

```
GI1$disp <- predict(GI1.dispfit, newdata = GI1)[,1]
GS2$disp <- predict(GS2.dispfit, newdata = GS2)[,1]
LGM$disp <- predict(LGM.dispfit, newdata = LGM)[,1]
LOIS3$disp <- predict(LOIS3.dispfit, newdata = LOIS3)[,1]
EOIS3$disp <- predict(EOIS3.dispfit, newdata = EOIS3)[,1]

# Fit mean models (no fixed effects, intercept only model with spatial and uncorrelated random effect)
EHmeanfit1 <- fitme(formula = mean_source_value ~ 1 + Matern(1|long + lat) + (1|SiteName),
                family = gaussian(link = identity), data = EH,
                resid.model = list(formula= ~ 0 + offset(disp), family = Gamma(link = identity)),
                prior.weights = n_source_value, control.dist = list(dist.method = "Earth"), method = "REML")
GS1meanfit1 <- fitme(formula = mean_source_value ~ 1 + Matern(1|long + lat) + (1|SiteName),
                 family = gaussian(link = identity), data = GS1,
                 resid.model = list(formula= ~ 0 + offset(disp), family = Gamma(link = identity)),
                 prior.weights = n_source_value, control.dist = list(dist.method = "Earth"), method = "REML")
GI1meanfit1 <- fitme(formula = mean_source_value ~ 1 + Matern(1|long + lat) + (1|SiteName),
                 family = gaussian(link = identity), data = GI1,
                 resid.model = list(formula= ~ 0 + offset(disp), family = Gamma(link = identity)),
                 prior.weights = n_source_value, control.dist = list(dist.method = "Earth"), method = "REML")
GS2meanfit1 <- fitme(formula = mean_source_value ~ 1 + Matern(1|long + lat) + (1|SiteName),
                 family = gaussian(link = identity), data = GS2,
                 resid.model = list(formula= ~ 0 + offset(disp), family = Gamma(link = identity)),
                 prior.weights = n_source_value, control.dist = list(dist.method = "Earth"), method = "REML")
LGMmeanfit1 <- fitme(formula = mean_source_value ~ 1 + Matern(1|long + lat) + (1|SiteName),
                 family = gaussian(link = identity), data = LGM,
                 resid.model = list(formula= ~ 0 + offset(disp), family = Gamma(link = identity)),
                 prior.weights = n_source_value, control.dist = list(dist.method = "Earth"), method = "REML")
LOIS3meanfit1 <- fitme(formula = mean_source_value ~ 1 + Matern(1|long + lat) + (1|SiteName),
                   family = gaussian(link = identity), data = LOIS3,
                   resid.model = list(formula= ~ 0 + offset(disp), family = Gamma(link = identity)),
                   prior.weights = n_source_value, control.dist = list(dist.method = "Earth"), method = "REML")
EOIS3meanfit1 <- fitme(formula = mean_source_value ~ 1 + Matern(1|long + lat) + (1|SiteName),
                   family = gaussian(link = identity), data = EOIS3,
                   resid.model = list(formula= ~ 0 + offset(disp), family = Gamma(link = identity)),
                   prior.weights = n_source_value, control.dist = list(dist.method = "Earth"), method = "REML")


# SET UP EMPTY RASTERS TO PREDICT INTO ####

  EH.raster <- raster(ncol=400, nrow=250, xmn=-10, xmx=30, ymn=35, ymx=60)
  projection(EH.raster) <- "+proj=longlat +datum=WGS84"
  EH_data_pred <- as.data.frame(EH.raster , xy = TRUE)
  EH.pred_locs <- cbind(EH_data_pred$x, EH_data_pred$y)
  EH.pred_pts <- SpatialPoints(EH.pred_locs)
  EH_data_pred$lat <- EH_data_pred$y
  EH_data_pred$long <- EH_data_pred$x
  EH_data_pred$SiteName <- EH_data_pred %>% group_indices(long, lat)

  GS1.raster <- raster(ncol=400, nrow=250, xmn=-10, xmx=30, ymn=35, ymx=60)
  projection(GS1.raster) <- "+proj=longlat +datum=WGS84"
  GS1_data_pred <- as.data.frame(GS1.raster , xy = TRUE)
  GS1.pred_locs <- cbind(GS1_data_pred$x, GS1_data_pred$y)
  GS1.pred_pts <- SpatialPoints(GS1.pred_locs)
  GS1_data_pred$lat <- GS1_data_pred$y
  GS1_data_pred$long <- GS1_data_pred$x
  GS1_data_pred$SiteName <- GS1_data_pred %>% group_indices(long, lat)

  GI1.raster <- raster(ncol=400, nrow=250, xmn=-10, xmx=30, ymn=35, ymx=60)
  projection(GI1.raster) <- "+proj=longlat +datum=WGS84"
  GI1_data_pred <- as.data.frame(GI1.raster , xy = TRUE)
  GI1.pred_locs <- cbind(GI1_data_pred$x, GI1_data_pred$y)
  GI1.pred_pts <- SpatialPoints(GI1.pred_locs)
  GI1_data_pred$lat <- GI1_data_pred$y
  GI1_data_pred$long <- GI1_data_pred$x
  GI1_data_pred$SiteName <- GI1_data_pred %>% group_indices(long, lat)

  GS2.raster <- raster(ncol=400, nrow=250, xmn=-10, xmx=30, ymn=35, ymx=60)
  projection(GS2.raster) <- "+proj=longlat +datum=WGS84"
  GS2_data_pred <- as.data.frame(GS2.raster , xy = TRUE)
  GS2.pred_locs <- cbind(GS2_data_pred$x, GS2_data_pred$y)
  GS2.pred_pts <- SpatialPoints(GS2.pred_locs)
  GS2_data_pred$lat <- GS2_data_pred$y
  GS2_data_pred$long <- GS2_data_pred$x
  GS2_data_pred$SiteName <- GS2_data_pred %>% group_indices(long, lat)

  LGM.raster <- raster(ncol=400, nrow=250, xmn=-10, xmx=30, ymn=35, ymx=60)
  projection(LGM.raster) <- "+proj=longlat +datum=WGS84"
  LGM_data_pred <- as.data.frame(LGM.raster , xy = TRUE)
  LGM.pred_locs <- cbind(LGM_data_pred$x, LGM_data_pred$y)
  LGM.pred_pts <- SpatialPoints(LGM.pred_locs)
  LGM_data_pred$lat <- LGM_data_pred$y
  LGM_data_pred$long <- LGM_data_pred$x
  LGM_data_pred$SiteName <- LGM_data_pred %>% group_indices(long, lat)

  LOIS3.raster <- raster(ncol=400, nrow=250, xmn=-10, xmx=30, ymn=35, ymx=60)
  projection(LOIS3.raster) <- "+proj=longlat +datum=WGS84"
  LOIS3_data_pred <- as.data.frame(LOIS3.raster , xy = TRUE)
  LOIS3.pred_locs <- cbind(LOIS3_data_pred$x, LOIS3_data_pred$y)
  LOIS3.pred_pts <- SpatialPoints(LOIS3.pred_locs)
  LOIS3_data_pred$lat <- LOIS3_data_pred$y
  LOIS3_data_pred$long <- LOIS3_data_pred$x
  LOIS3_data_pred$SiteName <- LOIS3_data_pred %>% group_indices(long, lat)

  EOIS3.raster <- raster(ncol=400, nrow=250, xmn=-10, xmx=30, ymn=35, ymx=60)
  projection(EOIS3.raster) <- "+proj=longlat +datum=WGS84"
```

```
    EOIS3_data_pred <- as.data.frame(EOIS3.raster , xy = TRUE)
    EOIS3.pred_locs <- cbind(EOIS3_data_pred$x, EOIS3_data_pred$y)
    EOIS3.pred_pts <- SpatialPoints(EOIS3.pred_locs)
    EOIS3_data_pred$lat <- EOIS3_data_pred$y
    EOIS3_data_pred$long <- EOIS3_data_pred$x
    EOIS3_data_pred$SiteName <- EOIS3_data_pred %>% group_indices(long, lat)

  # PREDICT ISOSCAPES ####

# EH
# Using env_data_pred and dispfit, we predict the residual dispersion variance for each prediction location:
EH_data_pred$disp <- predict(EH.dispfit, newdata = EH_data_pred)[, 1]
# We can then use the env_data_pred and meanfit to compute the predicted isotope value over all possible observations:
EH.predict <- predict(EHmeanfit1, newdata = EH_data_pred,
                      variances = list(predVar = TRUE, residVar = TRUE))
# Then extract the point predictions, variance and residual variance:
EH.pred <- EH_data_pred[, c("lat", "long")]
EH.pred$pred <- EH.predict[, 1]
EH.pred$predVar <- attr(EH.predict, "predVar")
EH.pred$residVar <- attr(EH.predict, "residVar")

# GS1
# Using env_data_pred and dispfit, we predict the residual dispersion variance for each prediction location:
GS1_data_pred$disp <- predict(GS1.dispfit, newdata = GS1_data_pred)[, 1]
# We can then use the env_data_pred and meanfit to compute the predicted isotope value over all possible observations:
GS1.predict <- predict(GS1meanfit1, newdata = GS1_data_pred,
                       variances = list(predVar = TRUE, residVar = TRUE))
# Then extract the point predictions, variance and residual variance:
GS1.pred <- GS1_data_pred[, c("lat", "long")]
GS1.pred$pred <- GS1.predict[, 1]
GS1.pred$predVar <- attr(GS1.predict, "predVar")
GS1.pred$residVar <- attr(GS1.predict, "residVar")

# GI1
# Using env_data_pred and dispfit, we predict the residual dispersion variance for each prediction location:
GI1_data_pred$disp <- predict(GI1.dispfit, newdata = GI1_data_pred)[, 1]
# We can then use the env_data_pred and meanfit to compute the predicted isotope value over all possible observations:
GI1.predict <- predict(GI1meanfit1, newdata = GI1_data_pred,
                       variances = list(predVar = TRUE, residVar = TRUE))
# Then extract the point predictions, variance and residual variance:
GI1.pred <- GI1_data_pred[, c("lat", "long")]
GI1.pred$pred <- GI1.predict[, 1]
GI1.pred$predVar <- attr(GI1.predict, "predVar")
GI1.pred$residVar <- attr(GI1.predict, "residVar")

# GS2
# Using env_data_pred and dispfit, we predict the residual dispersion variance for each prediction location:
GS2_data_pred$disp <- predict(GS2.dispfit, newdata = GS2_data_pred)[, 1]
# We can then use the env_data_pred and meanfit to compute the predicted isotope value over all possible observations:
GS2.predict <- predict(GS2meanfit1, newdata = GS2_data_pred,
                       variances = list(predVar = TRUE, residVar = TRUE))
# Then extract the point predictions, variance and residual variance:
GS2.pred <- GS2_data_pred[, c("lat", "long")]
GS2.pred$pred <- GS2.predict[, 1]
GS2.pred$predVar <- attr(GS2.predict, "predVar")
GS2.pred$residVar <- attr(GS2.predict, "residVar")

# LGM
# Using env_data_pred and dispfit, we predict the residual dispersion variance for each prediction location:
LGM_data_pred$disp <- predict(LGM.dispfit, newdata = LGM_data_pred)[, 1]
# We can then use the env_data_pred and meanfit to compute the predicted isotope value over all possible observations:
LGM.predict <- predict(LGMmeanfit1, newdata = LGM_data_pred,
                       variances = list(predVar = TRUE, residVar = TRUE))
# Then extract the point predictions, variance and residual variance:
LGM.pred <- LGM_data_pred[, c("lat", "long")]
LGM.pred$pred <- LGM.predict[, 1]
LGM.pred$predVar <- attr(LGM.predict, "predVar")
LGM.pred$residVar <- attr(LGM.predict, "residVar")

# LOIS3
# Using env_data_pred and dispfit, we predict the residual dispersion variance for each prediction location:
LOIS3_data_pred$disp <- predict(LOIS3.dispfit, newdata = LOIS3_data_pred)[, 1]
# We can then use the env_data_pred and meanfit to compute the predicted isotope value over all possible observations:
LOIS3.predict <- predict(LOIS3meanfit1, newdata = LOIS3_data_pred,
                         variances = list(predVar = TRUE, residVar = TRUE))
# Then extract the point predictions, variance and residual variance:
LOIS3.pred <- LOIS3_data_pred[, c("lat", "long")]
LOIS3.pred$pred <- LOIS3.predict[, 1]
LOIS3.pred$predVar <- attr(LOIS3.predict, "predVar")
LOIS3.pred$residVar <- attr(LOIS3.predict, "residVar")

# EOIS3
# Using env_data_pred and dispfit, we predict the residual dispersion variance for each prediction location:
EOIS3_data_pred$disp <- predict(EOIS3.dispfit, newdata = EOIS3_data_pred)[, 1]
# We can then use the env_data_pred and meanfit to compute the predicted isotope value over all possible observations:
EOIS3.predict <- predict(EOIS3meanfit1, newdata = EOIS3_data_pred,
                         variances = list(predVar = TRUE, residVar = TRUE))
# Then extract the point predictions, variance and residual variance:
EOIS3.pred <- EOIS3_data_pred[, c("lat", "long")]
EOIS3.pred$pred <- EOIS3.predict[, 1]
EOIS3.pred$predVar <- attr(EOIS3.predict, "predVar")
EOIS3.pred$residVar <- attr(EOIS3.predict, "residVar")

  # SET UP INTERPOLATION AREA FOR PLOTTING ####
```

```
# crop palaeocoastlines to plot area
pcoastEH.simp <- crop(pcoastEH.simp, extent(-10, 30, 35, 60))
pcoastGS1.simp <- crop(pcoastGS1.simp, extent(-10, 30, 35, 60))
pcoastGI1.simp <- crop(pcoastGI1.simp, extent(-10, 30, 35, 60))
pcoastGS2.simp <- crop(pcoastGS2.simp, extent(-10, 30, 35, 60))
pcoastLGM.simp <- crop(pcoastLGM.simp, extent(-10, 30, 35, 60))
pcoastLOIS3.simp <- crop(pcoastLOIS3.simp, extent(-10, 30, 35, 60))
pcoastEOIS3.simp <- crop(pcoastEOIS3.simp, extent(-10, 30, 35, 60))

# create interpolation area

# EH
# first create a buffered convex hull around points
EH.x <- EH$long
EH.y <- EH$lat
EH.d15N.sample_xy <- cbind(EH.x,EH.y)
EH.d15N.sample_pts <- SpatialPoints(EH.d15N.sample_xy)
EH.ch <- chull(EH.d15N.sample_xy)
EH.d15N.sample_bound <- EH.d15N.sample_xy[c(EH.ch, EH.ch[1]), ]  # closed polygon
EH.outer <- SpatialPolygons(list(Polygons(list(Polygon(EH.d15N.sample_bound)), ID=1)))
EH.outer <- gBuffer(EH.outer, 100000, byid=TRUE)
# then define interpolation area
EH.plotarea <- gDifference(EH.outer, pcoastEH.simp)
EH.interpol.area <- gDifference(EH.outer, EH.plotarea)

# GS1
# first create a buffered convex hull around points
GS1.x <- GS1$long
GS1.y <- GS1$lat
GS1.d15N.sample_xy <- cbind(GS1.x,GS1.y)
GS1.d15N.sample_pts <- SpatialPoints(GS1.d15N.sample_xy)
GS1.ch <- chull(GS1.d15N.sample_xy)
GS1.d15N.sample_bound <- GS1.d15N.sample_xy[c(GS1.ch, GS1.ch[1]), ]  # closed polygon
GS1.outer <- SpatialPolygons(list(Polygons(list(Polygon(GS1.d15N.sample_bound)), ID=1)))
GS1.outer <- gBuffer(GS1.outer, 100000, byid=TRUE)
# then define interpolation area
GS1.plotarea <- gDifference(GS1.outer, pcoastGS1.simp)
GS1.interpol.area <- gDifference(GS1.outer, GS1.plotarea)

# GI1
# first create a buffered convex hull around points
GI1.x <- GI1$long
GI1.y <- GI1$lat
GI1.d15N.sample_xy <- cbind(GI1.x,GI1.y)
GI1.d15N.sample_pts <- SpatialPoints(GI1.d15N.sample_xy)
GI1.ch <- chull(GI1.d15N.sample_xy)
GI1.d15N.sample_bound <- GI1.d15N.sample_xy[c(GI1.ch, GI1.ch[1]), ]  # closed polygon
GI1.outer <- SpatialPolygons(list(Polygons(list(Polygon(GI1.d15N.sample_bound)), ID=1)))
GI1.outer <- gBuffer(GI1.outer, 100000, byid=TRUE)
# then define interpolation area
GI1.plotarea <- gDifference(GI1.outer, pcoastGI1.simp)
GI1.interpol.area <- gDifference(GI1.outer, GI1.plotarea)

# GS2
# first create a buffered convex hull around points
GS2.x <- GS2$long
GS2.y <- GS2$lat
GS2.d15N.sample_xy <- cbind(GS2.x,GS2.y)
GS2.d15N.sample_pts <- SpatialPoints(GS2.d15N.sample_xy)
GS2.ch <- chull(GS2.d15N.sample_xy)
GS2.d15N.sample_bound <- GS2.d15N.sample_xy[c(GS2.ch, GS2.ch[1]), ]  # closed polygon
GS2.outer <- SpatialPolygons(list(Polygons(list(Polygon(GS2.d15N.sample_bound)), ID=1)))
GS2.outer <- gBuffer(GS2.outer, 100000, byid=TRUE)
# then define interpolation area
GS2.plotarea <- gDifference(GS2.outer, pcoastGS2.simp)
GS2.interpol.area <- gDifference(GS2.outer, GS2.plotarea)

# LGM
# first create a buffered convex hull around points
LGM.x <- LGM$long
LGM.y <- LGM$lat
LGM.d15N.sample_xy <- cbind(LGM.x,LGM.y)
LGM.d15N.sample_pts <- SpatialPoints(LGM.d15N.sample_xy)
LGM.ch <- chull(LGM.d15N.sample_xy)
LGM.d15N.sample_bound <- LGM.d15N.sample_xy[c(LGM.ch, LGM.ch[1]), ]  # closed polygon
LGM.outer <- SpatialPolygons(list(Polygons(list(Polygon(LGM.d15N.sample_bound)), ID=1)))
LGM.outer <- gBuffer(LGM.outer, 100000, byid=TRUE)
# then define interpolation area
LGM.plotarea <- gDifference(LGM.outer, pcoastLGM.simp)
LGM.interpol.area <- gDifference(LGM.outer, LGM.plotarea)

# LOIS3
# first create a buffered convex hull around points
LOIS3.x <- LOIS3$long
LOIS3.y <- LOIS3$lat
LOIS3.d15N.sample_xy <- cbind(LOIS3.x,LOIS3.y)
LOIS3.d15N.sample_pts <- SpatialPoints(LOIS3.d15N.sample_xy)
LOIS3.ch <- chull(LOIS3.d15N.sample_xy)
LOIS3.d15N.sample_bound <- LOIS3.d15N.sample_xy[c(LOIS3.ch, LOIS3.ch[1]), ]  # closed polygon
LOIS3.outer <- SpatialPolygons(list(Polygons(list(Polygon(LOIS3.d15N.sample_bound)), ID=1)))
LOIS3.outer <- gBuffer(LOIS3.outer, 100000, byid=TRUE)
# then define interpolation area
LOIS3.plotarea <- gDifference(LOIS3.outer, pcoastLOIS3.simp)
LOIS3.interpol.area <- gDifference(LOIS3.outer, LOIS3.plotarea)
```

```
# EOIS3
# first create a buffered convex hull around points
EOIS3.x <- EOIS3$long
EOIS3.y <- EOIS3$lat
EOIS3.d15N.sample_xy <- cbind(EOIS3.x,EOIS3.y)
EOIS3.d15N.sample_pts <- SpatialPoints(EOIS3.d15N.sample_xy)
EOIS3.ch <- chull(EOIS3.d15N.sample_xy)
EOIS3.d15N.sample_bound <- EOIS3.d15N.sample_xy[c(EOIS3.ch, EOIS3.ch[1]), ]  # closed polygon
EOIS3.outer <- SpatialPolygons(list(Polygons(list(Polygon(EOIS3.d15N.sample_bound)), ID=1)))
EOIS3.outer <- gBuffer(EOIS3.outer, 100000, byid=TRUE)
# then define interpolation area
EOIS3.plotarea <- gDifference(EOIS3.outer, pcoastEOIS3.simp)
EOIS3.interpol.area <- gDifference(EOIS3.outer, EOIS3.plotarea)

    # CREATE PREDICTION SURFACES ####

# EH
# tranform prediction surfaces into raster
EH.pred <- EH.pred[,c('long', 'lat', 'pred', 'predVar', 'residVar')]
EH.pred.rast <- rasterFromXYZ(EH.pred)  #Convert first two columns as lon-lat and third as value
# crop prediction values to interpolation area
EH.cr <- crop(EH.pred.rast, extent(EH.interpol.area), snap="out")
EH.fr <- rasterize(EH.interpol.area, EH.cr)
EH.lr <- raster::mask(x=EH.cr, mask=EH.fr)
# then convert raster to dataframe:
EH.pred.rast.df  <- as.data.frame(EH.lr, xy=TRUE)
# remove data with NAs
EH.pred.rast.df <- na.omit(EH.pred.rast.df)

# GS1
# tranform prediction surfaces into raster
GS1.pred <- GS1.pred[,c('long', 'lat', 'pred', 'predVar', 'residVar')]
GS1.pred.rast <- rasterFromXYZ(GS1.pred)  #Convert first two columns as lon-lat and third as value
# crop prediction values to interpolation area
GS1.cr <- crop(GS1.pred.rast, extent(GS1.interpol.area), snap="out")
GS1.fr <- rasterize(GS1.interpol.area, GS1.cr)
GS1.lr <- raster::mask(x=GS1.cr, mask=GS1.fr)
# then convert raster to dataframe:
GS1.pred.rast.df  <- as.data.frame(GS1.lr, xy=TRUE)
# remove data with NAs
GS1.pred.rast.df <- na.omit(GS1.pred.rast.df)

# GI1
# tranform prediction surfaces into raster
GI1.pred <- GI1.pred[,c('long', 'lat', 'pred', 'predVar', 'residVar')]
GI1.pred.rast <- rasterFromXYZ(GI1.pred)  #Convert first two columns as lon-lat and third as value
# crop prediction values to interpolation area
GI1.cr <- crop(GI1.pred.rast, extent(GI1.interpol.area), snap="out")
GI1.fr <- rasterize(GI1.interpol.area, GI1.cr)
GI1.lr <- raster::mask(x=GI1.cr, mask=GI1.fr)
# then convert raster to dataframe:
GI1.pred.rast.df  <- as.data.frame(GI1.lr, xy=TRUE)
# remove data with NAs
GI1.pred.rast.df <- na.omit(GI1.pred.rast.df)

# GS2
# tranform prediction surfaces into raster
GS2.pred <- GS2.pred[,c('long', 'lat', 'pred', 'predVar', 'residVar')]
GS2.pred.rast <- rasterFromXYZ(GS2.pred)  #Convert first two columns as lon-lat and third as value
# crop prediction values to interpolation area
GS2.cr <- crop(GS2.pred.rast, extent(GS2.interpol.area), snap="out")
GS2.fr <- rasterize(GS2.interpol.area, GS2.cr)
GS2.lr <- raster::mask(x=GS2.cr, mask=GS2.fr)
# then convert raster to dataframe:
GS2.pred.rast.df  <- as.data.frame(GS2.lr, xy=TRUE)
# remove data with NAs
GS2.pred.rast.df <- na.omit(GS2.pred.rast.df)

# LGM
# tranform prediction surfaces into raster
LGM.pred <- LGM.pred[,c('long', 'lat', 'pred', 'predVar', 'residVar')]
LGM.pred.rast <- rasterFromXYZ(LGM.pred)  #Convert first two columns as lon-lat and third as value
# crop prediction values to interpolation area
LGM.cr <- crop(LGM.pred.rast, extent(LGM.interpol.area), snap="out")
LGM.fr <- rasterize(LGM.interpol.area, LGM.cr)
LGM.lr <- raster::mask(x=LGM.cr, mask=LGM.fr)
# then convert raster to dataframe:
LGM.pred.rast.df  <- as.data.frame(LGM.lr, xy=TRUE)
# remove data with NAs
LGM.pred.rast.df <- na.omit(LGM.pred.rast.df)

# LOIS3
# tranform prediction surfaces into raster
LOIS3.pred <- LOIS3.pred[,c('long', 'lat', 'pred', 'predVar', 'residVar')]
LOIS3.pred.rast <- rasterFromXYZ(LOIS3.pred)  #Convert first two columns as lon-lat and third as value
# crop prediction values to interpolation area
LOIS3.cr <- crop(LOIS3.pred.rast, extent(LOIS3.interpol.area), snap="out")
LOIS3.fr <- rasterize(LOIS3.interpol.area, LOIS3.cr)
LOIS3.lr <- raster::mask(x=LOIS3.cr, mask=LOIS3.fr)
# then convert raster to dataframe:
LOIS3.pred.rast.df  <- as.data.frame(LOIS3.lr, xy=TRUE)
# remove data with NAs
LOIS3.pred.rast.df <- na.omit(LOIS3.pred.rast.df)

# EOIS3
```

```
# tranform prediction surfaces into raster
EOIS3.pred <- EOIS3.pred[,c('long', 'lat', 'pred', 'predVar', 'residVar')]
EOIS3.pred.rast <- rasterFromXYZ(EOIS3.pred)  #Convert first two columns as lon-lat and third as value
# crop prediction values to interpolation area
EOIS3.cr <- crop(EOIS3.pred.rast, extent(EOIS3.interpol.area), snap="out")
EOIS3.fr <- rasterize(EOIS3.interpol.area, EOIS3.cr)
EOIS3.lr <- raster::mask(x=EOIS3.cr, mask=EOIS3.fr)
# then convert raster to dataframe:
EOIS3.pred.rast.df  <- as.data.frame(EOIS3.lr, xy=TRUE)
# remove data with NAs
EOIS3.pred.rast.df <- na.omit(EOIS3.pred.rast.df)

    # PLOT ISOSCAPES (FIGURE 3 and 4) ####
# set up colour schemes
col <- rev(brewer.pal(11,"Spectral"))
col.var <- c("honeydew1", "palegreen", "mediumseagreen", "seagreen4")

# EH prediciton surface
(EHmappred <- ggplot()+
  geom_polygon(data=pcoastEH.simp, aes(long, lat, group=group), fill="lightgrey") +
  geom_tile(data = EH.pred.rast.df, mapping=aes(x=x, y=y, fill=pred)) +
  scale_fill_stepsn(colours = col, limits = c(0,6), breaks=c(0.5,1,1.5,2,2.5,3,3.5,4,4.5,5,5.5)) +
  geom_path(data=eurmed, aes(long, lat, group=group), colour="black") +
  geom_polygon(data=iceEH, aes(long, lat, group=group), colour="black", fill="white") +
  geom_point(data = EH, aes(long, lat), colour="black", size=0.5) +
  coord_cartesian(xlim = c(-10,25), ylim = c(40, 60)) +
  xlab("Longitude") +
  ylab("Latitude") +
  labs(title="Early Holocene",
       fill=expression(paste(delta^{15}, "N (\u2030)"))) +
  theme_bw() +
  theme(legend.position="right"))

# EH prediciton variance
(EHmapvar <- ggplot()+
  geom_polygon(data=pcoastEH.simp, aes(long, lat, group=group), fill="lightgrey") +
  geom_tile(data = EH.pred.rast.df, mapping=aes(x=x, y=y, fill=predVar)) +
  scale_fill_stepsn(colours = col.var, breaks=c(0.5,1,1.5), limits = c(0, 3))+
  geom_path(data=eurmed, aes(long, lat, group=group), colour="black") +
  geom_polygon(data=iceEH, aes(long, lat, group=group), colour="black", fill="white") +
  geom_point(data = EH, aes(long, lat), colour="black", size=0.5) +
  coord_cartesian(xlim = c(-10,25), ylim = c(40, 60)) +
  xlab("Longitude") +
  ylab("Latitude") +
  labs(title="Early Holocene",
       fill=expression(paste(delta^{15}, "N (\u2030)"))) +
  theme(legend.position="right") +
  theme_bw())

# GS1 prediciton surface
(GS1mappred <- ggplot()+
  geom_polygon(data=pcoastGS1.simp, aes(long, lat, group=group), fill="lightgrey") +
  geom_tile(data = GS1.pred.rast.df, mapping=aes(x=x, y=y, fill=pred)) +
  scale_fill_stepsn(colours = col, limits = c(0,6), breaks=c(0.5,1,1.5,2,2.5,3,3.5,4,4.5,5,5.5)) +
  geom_path(data=eurmed, aes(long, lat, group=group), colour="black") +
  geom_polygon(data=iceGS1, aes(long, lat, group=group), colour="black", fill="white") +
  geom_point(data = GS1, aes(long, lat), colour="black", size=0.5) +
  coord_cartesian(xlim = c(-10,25), ylim = c(40, 60)) +
  xlab("Longitude") +
  ylab("Latitude") +
  labs(title="Younger Dryas",
       fill=expression(paste(delta^{15}, "N (\u2030)"))) +
  theme_bw() +
  theme(legend.position="none"))


# GS1 prediciton variance
(GS1mapvar <- ggplot()+
  geom_polygon(data=pcoastGS1.simp, aes(long, lat, group=group), fill="lightgrey") +
  geom_tile(data = GS1.pred.rast.df, mapping=aes(x=x, y=y, fill=predVar)) +
  scale_fill_stepsn(colours = col.var, breaks=c(0.5,1,1.5), limits = c(0, 3))+
  geom_path(data=eurmed, aes(long, lat, group=group), colour="black") +
  geom_polygon(data=iceGS1, aes(long, lat, group=group), colour="black", fill="white") +
  geom_point(data = GS1, aes(long, lat), colour="black", size=0.5) +
  coord_cartesian(xlim = c(-10,25), ylim = c(40, 60)) +
  xlab("Longitude") +
  ylab("Latitude") +
  labs(title="Younger Dryas",
       fill=expression(paste(delta^{15}, "N (\u2030)"))) +
  theme_bw() +
  theme(legend.position="none"))


# GI1 prediciton surface
(GI1mappred <- ggplot()+
  geom_polygon(data=pcoastGI1.simp, aes(long, lat, group=group), fill="lightgrey") +
  geom_tile(data = GI1.pred.rast.df, mapping=aes(x=x, y=y, fill=pred)) +
  scale_fill_stepsn(colours = col, limits = c(0,6), breaks=c(0.5,1,1.5,2,2.5,3,3.5,4,4.5,5,5.5)) +
  geom_path(data=eurmed, aes(long, lat, group=group), colour="black") +
  geom_polygon(data=iceGI1, aes(long, lat, group=group), colour="black", fill="white") +
  geom_point(data = GI1, aes(long, lat), colour="black", size=0.5) +
  coord_cartesian(xlim = c(-10,25), ylim = c(40, 60)) +
  xlab("Longitude") +
  ylab("Latitude") +
  labs(title="Late Glacial Interstadial",
```

```
      fill=expression(paste(delta^{15}, "N (\u2030)"))) +
  theme_bw() +
  theme(legend.position="none"))

# GI1 prediciton variance
(GI1mapvar <- ggplot()+
  geom_polygon(data=pcoastGI1.simp, aes(long, lat, group=group), fill="lightgrey") +
  geom_tile(data = GI1.pred.rast.df, mapping=aes(x=x, y=y, fill=predVar)) +
  scale_fill_stepsn(colours = col.var, breaks=c(0.5,1,1.5), limits = c(0, 3))+
  geom_path(data=eurmed, aes(long, lat, group=group), colour="black") +
  geom_polygon(data=iceGI1, aes(long, lat, group=group), colour="black", fill="white") +
  geom_point(data = GI1, aes(long, lat), colour="black", size=0.5) +
  coord_cartesian(xlim = c(-10,25), ylim = c(40, 60)) +
  xlab("Longitude") +
  ylab("Latitude") +
  labs(title="Late Glacial Interstadial",
      fill=expression(paste(delta^{15}, "N (\u2030)"))) +
  theme_bw() +
  theme(legend.position="none"))

# GS2 prediciton surface
(GS2mappred <- ggplot()+
  geom_polygon(data=pcoastGS2.simp, aes(long, lat, group=group), fill="lightgrey") +
  geom_tile(data = GS2.pred.rast.df, mapping=aes(x=x, y=y, fill=pred)) +
  scale_fill_stepsn(colours = col, limits = c(0,6), breaks=c(0.5,1,1.5,2,2.5,3,3.5,4,4.5,5,5.5)) +
  geom_path(data=eurmed, aes(long, lat, group=group), colour="black") +
  geom_polygon(data=iceGS2, aes(long, lat, group=group), colour="black", fill="white") +
  geom_point(data = GS2, aes(long, lat), colour="black", size=0.5) +
  coord_cartesian(xlim = c(-10,25), ylim = c(40, 60)) +
  xlab("Longitude") +
  ylab("Latitude") +
  labs(title="Last Glacial Termination",
      fill=expression(paste(delta^{15}, "N (\u2030)"))) +
  theme_bw() +
  theme(legend.position="none"))

# GS2 prediciton variance
(GS2mapvar <- ggplot()+
  geom_polygon(data=pcoastGS2.simp, aes(long, lat, group=group), fill="lightgrey") +
  geom_tile(data = GS2.pred.rast.df, mapping=aes(x=x, y=y, fill=predVar)) +
  scale_fill_stepsn(colours = col.var, breaks=c(0.5,1,1.5), limits = c(0, 3))+
  geom_path(data=eurmed, aes(long, lat, group=group), colour="black") +
  geom_polygon(data=iceGS2, aes(long, lat, group=group), colour="black", fill="white") +
  geom_point(data = GS2, aes(long, lat), colour="black", size=0.5) +
  coord_cartesian(xlim = c(-10,25), ylim = c(40, 60)) +
  xlab("Longitude") +
  ylab("Latitude") +
  labs(title="Last Glacial Termination",
      fill=expression(paste(delta^{15}, "N (\u2030)"))) +
  theme_bw() +
  theme(legend.position="none"))

# LGM prediciton surface
(LGMmappred <- ggplot()+
  geom_polygon(data=pcoastLGM.simp, aes(long, lat, group=group), fill="lightgrey") +
  geom_tile(data = LGM.pred.rast.df, mapping=aes(x=x, y=y, fill=pred)) +
  scale_fill_stepsn(colours = col, limits = c(0,6), breaks=c(0.5,1,1.5,2,2.5,3,3.5,4,4.5,5,5.5)) +
  geom_path(data=eurmed, aes(long, lat, group=group), colour="black") +
  geom_polygon(data=iceLGM, aes(long, lat, group=group), colour="black", fill="white") +
  geom_point(data = LGM, aes(long, lat), colour="black", size=0.5) +
  coord_cartesian(xlim = c(-10,25), ylim = c(40, 60)) +
  xlab("Longitude") +
  ylab("Latitude") +
  labs(title="Last Glacial Maximum",
      fill=expression(paste(delta^{15}, "N (\u2030)"))) +
  theme_bw() +
  theme(legend.position="none"))

# LGM prediciton variance
(LGMmapvar <- ggplot()+
  geom_polygon(data=pcoastLGM.simp, aes(long, lat, group=group), fill="lightgrey") +
  geom_tile(data = LGM.pred.rast.df, mapping=aes(x=x, y=y, fill=predVar)) +
  scale_fill_stepsn(colours = col.var, breaks=c(0.5,1,1.5), limits = c(0, 3))+
  geom_path(data=eurmed, aes(long, lat, group=group), colour="black") +
  geom_polygon(data=iceLGM, aes(long, lat, group=group), colour="black", fill="white") +
  geom_point(data = LGM, aes(long, lat), colour="black", size=0.5) +
  coord_cartesian(xlim = c(-10,25), ylim = c(40, 60)) +
  xlab("Longitude") +
  ylab("Latitude") +
  labs(title="Last Glacial Maximum",
      fill=expression(paste(delta^{15}, "N (\u2030)"))) +
  theme_bw() +
  theme(legend.position="none"))

# LOIS3 prediciton surface
(LOIS3mappred <- ggplot()+
  geom_polygon(data=pcoastLOIS3.simp, aes(long, lat, group=group), fill="lightgrey") +
  geom_tile(data = LOIS3.pred.rast.df, mapping=aes(x=x, y=y, fill=pred)) +
  scale_fill_stepsn(colours = col, limits = c(0,6), breaks=c(0.5,1,1.5,2,2.5,3,3.5,4,4.5,5,5.5)) +
  geom_path(data=eurmed, aes(long, lat, group=group), colour="black") +
  geom_polygon(data=iceLOIS3, aes(long, lat, group=group), colour="black", fill="white") +
  geom_point(data = LOIS3, aes(long, lat), colour="black", size=0.5) +
  coord_cartesian(xlim = c(-10,25), ylim = c(40, 60)) +
  xlab("Longitude") +
  ylab("Latitude") +
```

```
  labs(title="Late OIS 3",
       fill=expression(paste(delta^{15}, "N (\u2030)"))) +
  theme_bw() +
  theme(legend.position="none"))

# LOIS3 prediciton variance
(LOIS3mapvar <- ggplot()+
  geom_polygon(data=pcoastLOIS3.simp, aes(long, lat, group=group), fill="lightgrey") +
  geom_tile(data = LOIS3.pred.rast.df, mapping=aes(x=x, y=y, fill=predVar)) +
  scale_fill_stepsn(colours = col.var, breaks=c(0.5,1,1.5), limits = c(0, 3))+
  geom_path(data=eurmed, aes(long, lat, group=group), colour="black") +
  geom_polygon(data=iceLOIS3, aes(long, lat, group=group), colour="black", fill="white") +
  geom_point(data = LOIS3, aes(long, lat), colour="black", size=0.5) +
  coord_cartesian(xlim = c(-10,25), ylim = c(40, 60)) +
  xlab("Longitude") +
  ylab("Latitude") +
  labs(title="Late OIS 3",
       fill=expression(paste(delta^{15}, "N (\u2030)"))) +
  theme_bw() +
  theme(legend.position="none"))

# EOIS3 prediciton surface
(EOIS3mappred <- ggplot()+
  geom_polygon(data=pcoastEOIS3.simp, aes(long, lat, group=group), fill="lightgrey") +
  geom_tile(data = EOIS3.pred.rast.df, mapping=aes(x=x, y=y, fill=pred)) +
  scale_fill_stepsn(colours = col, limits = c(0,6), breaks=c(0.5,1,1.5,2,2.5,3,3.5,4,4.5,5,5.5)) +
  geom_path(data=eurmed, aes(long, lat, group=group), colour="black") +
  geom_polygon(data=iceEOIS3, aes(long, lat, group=group), colour="black", fill="white") +
  geom_point(data = EOIS3, aes(long, lat), colour="black", size=0.5) +
  coord_cartesian(xlim = c(-10,25), ylim = c(40, 60)) +
  xlab("Longitude") +
  ylab("Latitude") +
  labs(title="Early OIS 3",
       fill=expression(paste(delta^{15}, "N (\u2030)"))) +
  theme_bw() +
  theme(legend.position="none"))

# EOIS3 prediciton variance
(EOIS3mapvar <- ggplot()+
  geom_polygon(data=pcoastEOIS3.simp, aes(long, lat, group=group), fill="lightgrey") +
  geom_tile(data = EOIS3.pred.rast.df, mapping=aes(x=x, y=y, fill=predVar)) +
  scale_fill_stepsn(colours = col.var, breaks=c(0.5,1,1.5), limits = c(0, 3))+
  geom_path(data=eurmed, aes(long, lat, group=group), colour="black") +
  geom_polygon(data=iceEOIS3, aes(long, lat, group=group), colour="black", fill="white") +
  geom_point(data = EOIS3, aes(long, lat), colour="black", size=0.5) +
  coord_cartesian(xlim = c(-10,25), ylim = c(40, 60)) +
  xlab("Longitude") +
  ylab("Latitude") +
  labs(title="Early OIS 3",
       fill=expression(paste(delta^{15}, "N (\u2030)"))) +
  theme_bw() +
  theme(legend.position="none"))

# first get legends
legendPred <- get_legend(EHmappred)
# remove legend from EH ploy
EHmappred <- EHmappred + theme(legend.position="none")
# combine plots
predplots <- ggpubr::ggarrange(EHmappred,
                               GS1mappred,
                               GI1mappred,
                               GS2mappred,
                               LGMmappred,
                               LOIS3mappred,
                               EOIS3mappred,
                               legendPred, ncol=2, nrow=4)

ggsave(file="Figure_3.pdf", predplots, width = 20, height = 30, units = c("cm"), dpi = 300)
ggsave(file="Figure_3.png", predplots, width = 20, height = 30, units = c("cm"), dpi = 300)


# variance surfaces #

# first get legends
legendvar <- get_legend(EHmapvar)

EHmapvar <- EHmapvar + theme(legend.position="none")

varplots <- ggpubr::ggarrange(EHmapvar,
                              GS1mapvar,
                              GI1mapvar,
                              GS2mapvar,
                              LGMmapvar,
                              LOIS3mapvar,
                              EOIS3mapvar,
                              legendvar, ncol=2, nrow=4)

ggsave(file="Figure_4.pdf", varplots, width = 20, height = 30, units = c("cm"), dpi = 300)
ggsave(file="Figure_4.png", varplots, width = 20, height = 30, units = c("cm"), dpi = 300)


    # COMPARE PREDICTED TO MODELLED VALUES  (FIGURE 5)####

# turn each dataframe into an spdf
d15N.EH.aggre.spdf <- EH
```

```
coordinates(d15N.EH.aggre.spdf) <- ~long+lat
proj4string(d15N.EH.aggre.spdf) <- latlon
d15N.GS1.aggre.spdf <- GS1
coordinates(d15N.GS1.aggre.spdf) <- ~long+lat
proj4string(d15N.GS1.aggre.spdf) <- latlon
d15N.GI1.aggre.spdf <- GI1
coordinates(d15N.GI1.aggre.spdf) <- ~long+lat
proj4string(d15N.GI1.aggre.spdf) <- latlon
d15N.GS2.aggre.spdf <- GS2
coordinates(d15N.GS2.aggre.spdf) <- ~long+lat
proj4string(d15N.GS2.aggre.spdf) <- latlon
d15N.LGM.aggre.spdf <- LGM
coordinates(d15N.LGM.aggre.spdf) <- ~long+lat
proj4string(d15N.LGM.aggre.spdf) <- latlon
d15N.LOIS3.aggre.spdf <- LOIS3
coordinates(d15N.LOIS3.aggre.spdf) <- ~long+lat
proj4string(d15N.LOIS3.aggre.spdf) <- latlon
d15N.EOIS3.aggre.spdf <- EOIS3
coordinates(d15N.EOIS3.aggre.spdf) <- ~long+lat
proj4string(d15N.EOIS3.aggre.spdf) <- latlon

#overlay rasters and extract predicted values
EH_predicted_data <- raster::extract(EH.lr, d15N.EH.aggre.spdf) # extract data
d15N.EH.aggre.spdf <- cbind(d15N.EH.aggre.spdf, EH_predicted_data) # merge to the main data
colnames(d15N.EH.aggre.spdf@data)
# convert back to df and remove NAs
d15N.EH.aggre.df <- as.data.frame(d15N.EH.aggre.spdf)
d15N.EH.aggre.df <- d15N.EH.aggre.df %>% drop_na(pred)

# Calculate RMSE
rmse.EH <- round(modelRMSE(o = d15N.EH.aggre.df$mean_source_value, p = d15N.EH.aggre.df$pred), 2)
rmse.EH <- as.character(rmse.EH)
rmse.EH

#overlay rasters and extract predicted values
GS1_predicted_data <- raster::extract(GS1.lr, d15N.GS1.aggre.spdf) # extract data
d15N.GS1.aggre.spdf <- cbind(d15N.GS1.aggre.spdf, GS1_predicted_data) # merge to the main data
colnames(d15N.GS1.aggre.spdf@data)
# convert back to df and remove NAs
d15N.GS1.aggre.df <- as.data.frame(d15N.GS1.aggre.spdf)
d15N.GS1.aggre.df <- d15N.GS1.aggre.df %>% drop_na(pred)

# Calculate RMSE
rmse.GS1 <- round(modelRMSE(o = d15N.GS1.aggre.df$mean_source_value, p = d15N.GS1.aggre.df$pred), 2)
rmse.GS1 <- as.character(rmse.GS1)
rmse.GS1

#overlay rasters and extract predicted values
GI1_predicted_data <- raster::extract(GI1.lr, d15N.GI1.aggre.spdf) # extract data
d15N.GI1.aggre.spdf <- cbind(d15N.GI1.aggre.spdf, GI1_predicted_data) # merge to the main data
colnames(d15N.GI1.aggre.spdf@data)
# convert back to df and remove NAs
d15N.GI1.aggre.df <- as.data.frame(d15N.GI1.aggre.spdf)
d15N.GI1.aggre.df <- d15N.GI1.aggre.df %>% drop_na(pred)

# Calculate RMSE
rmse.GI1 <- round(modelRMSE(o = d15N.GI1.aggre.df$mean_source_value, p = d15N.GI1.aggre.df$pred), 2)
rmse.GI1 <- as.character(rmse.GI1)
rmse.GI1

#overlay rasters and extract predicted values
GS2_predicted_data <- raster::extract(GS2.lr, d15N.GS2.aggre.spdf) # extract data
d15N.GS2.aggre.spdf <- cbind(d15N.GS2.aggre.spdf, GS2_predicted_data) # merge to the main data
colnames(d15N.GS2.aggre.spdf@data)
# convert back to df and remove NAs
d15N.GS2.aggre.df <- as.data.frame(d15N.GS2.aggre.spdf)
d15N.GS2.aggre.df <- d15N.GS2.aggre.df %>% drop_na(pred)

# Calculate RMSE
rmse.GS2 <- round(modelRMSE(o = d15N.GS2.aggre.df$mean_source_value, p = d15N.GS2.aggre.df$pred), 2)
rmse.GS2 <- as.character(rmse.GS2)
rmse.GS2

#overlay rasters and extract predicted values
LGM_predicted_data <- raster::extract(LGM.lr, d15N.LGM.aggre.spdf) # extract data
d15N.LGM.aggre.spdf <- cbind(d15N.LGM.aggre.spdf, LGM_predicted_data) # merge to the main data
colnames(d15N.LGM.aggre.spdf@data)
# convert back to df and remove NAs
d15N.LGM.aggre.df <- as.data.frame(d15N.LGM.aggre.spdf)
d15N.LGM.aggre.df <- d15N.LGM.aggre.df %>% drop_na(pred)

# Calculate RMSE
rmse.LGM <- round(modelRMSE(o = d15N.LGM.aggre.df$mean_source_value, p = d15N.LGM.aggre.df$pred), 2)
rmse.LGM <- as.character(rmse.LGM)
rmse.LGM

#overlay rasters and extract predicted values
LOIS3_predicted_data <- raster::extract(LOIS3.lr, d15N.LOIS3.aggre.spdf) # extract data
d15N.LOIS3.aggre.spdf <- cbind(d15N.LOIS3.aggre.spdf, LOIS3_predicted_data) # merge to the main data
colnames(d15N.LOIS3.aggre.spdf@data)
# convert back to df and remove NAs
d15N.LOIS3.aggre.df <- as.data.frame(d15N.LOIS3.aggre.spdf)
d15N.LOIS3.aggre.df <- d15N.LOIS3.aggre.df %>% drop_na(pred)

# Calculate RMSE
```

```
rmse.LOIS3 <- round(modelRMSE(o = d15N.LOIS3.aggre.df$mean_source_value, p = d15N.LOIS3.aggre.df$pred), 2)
rmse.LOIS3 <- as.character(rmse.LOIS3)
rmse.LOIS3


#overlay rasters and extract predicted values
EOIS3_predicted_data <- raster::extract(EOIS3.lr, d15N.EOIS3.aggre.spdf) # extract data
d15N.EOIS3.aggre.spdf <- cbind(d15N.EOIS3.aggre.spdf, EOIS3_predicted_data) # merge to the main data
colnames(d15N.EOIS3.aggre.spdf@data)
# convert back to df and remove NAs
d15N.EOIS3.aggre.df <- as.data.frame(d15N.EOIS3.aggre.spdf)
d15N.EOIS3.aggre.df <- d15N.EOIS3.aggre.df %>% drop_na(pred)

# Calculate RMSE
rmse.EOIS3 <- round(modelRMSE(o = d15N.EOIS3.aggre.df$mean_source_value, p = d15N.EOIS3.aggre.df$pred), 2)
rmse.EOIS3 <- as.character(rmse.EOIS3)
rmse.EOIS3




(EH.pred.ob <- ggplot(data = d15N.EH.aggre.df, mapping=aes(x=pred, y=mean_source_value))+
  geom_point() +
  geom_abline(slope=1, intercept=0)+
  geom_smooth(method = "lm", se=TRUE, color="red", formula = y ~ x) +
  stat_poly_eq(aes(label = paste(..rr.label.., sep = "~~~")),
               label.x.npc = "right", label.y.npc = 0.15,
               formula = y ~ x, parse = TRUE, size = 4) +
  stat_poly_eq(aes(label = paste(..p.value.label.., sep = "~~~")),
               label.x.npc = "right", label.y.npc = "bottom",
               formula = y ~ x, parse = TRUE, size = 4) +
  annotate("text", x=10.5, y=3, label = paste("RMSE = ", rmse.EH))+
  ggtitle("Early Holocene")+
  labs(x = expression(paste(delta^{15}, "N predicted (\u2030)")),
       y = expression(paste(delta^{15}, "N observed (\u2030)"))) +
   scale_y_continuous(limits = c(0,12), breaks = seq(0, 12, by = 2)) +
   scale_x_continuous(limits = c(0,12),breaks = seq(0, 12, by = 2)) +
  theme_bw())

(GS1.pred.ob <- ggplot(data = d15N.GS1.aggre.df, mapping=aes(x=pred, y=mean_source_value))+
  geom_point() +
  geom_abline(slope=1, intercept=0)+
  geom_smooth(method = "lm", se=TRUE, color="red", formula = y ~ x) +
  stat_poly_eq(aes(label = paste(..rr.label.., sep = "~~~")),
               label.x.npc = "right", label.y.npc = 0.15,
               formula = y ~ x, parse = TRUE, size = 4) +
  stat_poly_eq(aes(label = paste(..p.value.label.., sep = "~~~")),
               label.x.npc = "right", label.y.npc = "bottom",
               formula = y ~ x, parse = TRUE, size = 4) +
  annotate("text", x=10.5, y=3, label = paste("RMSE = ", rmse.GS1))+
  ggtitle("Younger Dryas")+
  labs(x = expression(paste(delta^{15}, "N predicted (\u2030)")),
       y = expression(paste(delta^{15}, "N observed (\u2030)"))) +
    scale_y_continuous(limits = c(0,12), breaks = seq(0, 12, by = 2)) +
    scale_x_continuous(limits = c(0,12),breaks = seq(0, 12, by = 2)) +
  theme_bw())

(GI1.pred.ob <- ggplot(data = d15N.GI1.aggre.df, mapping=aes(x=pred, y=mean_source_value))+
  geom_point() +
  geom_abline(slope=1, intercept=0)+
  geom_smooth(method = "lm", se=TRUE, color="red", formula = y ~ x) +
  stat_poly_eq(aes(label = paste(..rr.label.., sep = "~~~")),
               label.x.npc = "right", label.y.npc = 0.15,
               formula = y ~ x, parse = TRUE, size = 4) +
  stat_poly_eq(aes(label = paste(..p.value.label.., sep = "~~~")),
               label.x.npc = "right", label.y.npc = "bottom",
               formula = y ~ x, parse = TRUE, size = 4) +
  annotate("text", x=10.5, y=3, label = paste("RMSE = ", rmse.GI1))+
  ggtitle("Late Glacial Interstadial")+
  labs(x = expression(paste(delta^{15}, "N predicted (\u2030)")),
       y = expression(paste(delta^{15}, "N observed (\u2030)"))) +
    scale_y_continuous(limits = c(0,12), breaks = seq(0, 12, by = 2)) +
    scale_x_continuous(limits = c(0,12),breaks = seq(0, 12, by = 2)) +
  theme_bw())

(GS2.pred.ob <- ggplot(data = d15N.GS2.aggre.df, mapping=aes(x=pred, y=mean_source_value))+
  geom_point() +
  geom_abline(slope=1, intercept=0)+
  geom_smooth(method = "lm", se=TRUE, color="red", formula = y ~ x) +
  stat_poly_eq(aes(label = paste(..rr.label.., sep = "~~~")),
               label.x.npc = "right", label.y.npc = 0.15,
               formula = y ~ x, parse = TRUE, size = 4) +
  stat_poly_eq(aes(label = paste(..p.value.label.., sep = "~~~")),
               label.x.npc = "right", label.y.npc = "bottom",
               formula = y ~ x, parse = TRUE, size = 4) +
  annotate("text",x=10.5, y=3, label = paste("RMSE = ", rmse.GS2))+
  ggtitle("Last Glacial Termination")+
  labs(x = expression(paste(delta^{15}, "N predicted (\u2030)")),
       y = expression(paste(delta^{15}, "N observed (\u2030)"))) +
    scale_y_continuous(limits = c(0,12), breaks = seq(0, 12, by = 2)) +
    scale_x_continuous(limits = c(0,12),breaks = seq(0, 12, by = 2)) +
  theme_bw())

(LGM.pred.ob <- ggplot(data = d15N.LGM.aggre.df, mapping=aes(x=pred, y=mean_source_value))+
  geom_point() +
```

```
  geom_abline(slope=1, intercept=0)+
  geom_smooth(method = "lm", se=TRUE, color="red", formula = y ~ x) +
  stat_poly_eq(aes(label = paste(..rr.label.., sep = "~~~")),
               label.x.npc = "right", label.y.npc = 0.15,
               formula = y ~ x, parse = TRUE, size = 4) +
  stat_poly_eq(aes(label = paste(..p.value.label.., sep = "~~~")),
               label.x.npc = "right", label.y.npc = "bottom",
               formula = y ~ x, parse = TRUE, size = 4) +
  annotate("text", x=10.5, y=3, label = paste("RMSE = ", rmse.LGM))+
  ggtitle("Last Glacial Maximum")+
  labs(x = expression(paste(delta^{15}, "N predicted (\u2030)")),
       y = expression(paste(delta^{15}, "N observed (\u2030)"))) +
    scale_y_continuous(limits = c(0,12), breaks = seq(0, 12, by = 2)) +
    scale_x_continuous(limits = c(0,12),breaks = seq(0, 12, by = 2)) +
  theme_bw())

(LOIS3.pred.ob <- ggplot(data = d15N.LOIS3.aggre.df, mapping=aes(x=pred, y=mean_source_value))+
  geom_point() +
  geom_abline(slope=1, intercept=0)+
  geom_smooth(method = "lm", se=TRUE, color="red", formula = y ~ x) +
  stat_poly_eq(aes(label = paste(..rr.label.., sep = "~~~")),
               label.x.npc = "right", label.y.npc = 0.15,
               formula = y ~ x, parse = TRUE, size = 4) +
  stat_poly_eq(aes(label = paste(..p.value.label.., sep = "~~~")),
               label.x.npc = "right", label.y.npc = "bottom",
               formula = y ~ x, parse = TRUE, size = 4) +
  annotate("text", x=10.5, y=3, label = paste("RMSE = ", rmse.LOIS3))+
  ggtitle("Late OIS 3")+
  labs(x = expression(paste(delta^{15}, "N predicted (\u2030)")),
       y = expression(paste(delta^{15}, "N observed (\u2030)"))) +
    scale_y_continuous(limits = c(0,12), breaks = seq(0, 12, by = 2)) +
    scale_x_continuous(limits = c(0,12),breaks = seq(0, 12, by = 2)) +
  theme_bw())


# plot predicted versus observed values
(EOIS3.pred.ob <- ggplot(data = d15N.EOIS3.aggre.df, mapping=aes(x=pred, y=mean_source_value))+
  geom_point() +
  geom_abline(slope=1, intercept=0)+
  geom_smooth(method = "lm", se=TRUE, color="red", formula = y ~ x) +
  stat_poly_eq(aes(label = paste(..rr.label.., sep = "~~~")),
               label.x.npc = "right", label.y.npc = 0.15,
               formula = y ~ x, parse = TRUE, size = 4) +
  stat_poly_eq(aes(label = paste(..p.value.label.., sep = "~~~")),
               label.x.npc = "right", label.y.npc = "bottom",
               formula = y ~ x, parse = TRUE, size = 4) +
  annotate("text", x=10.5, y=3, label = paste("RMSE = ", rmse.EOIS3))+
  ggtitle("Early OIS 3")+
  labs(x = expression(paste(delta^{15}, "N predicted (\u2030)")),
       y = expression(paste(delta^{15}, "N observed (\u2030)"))) +
  scale_y_continuous(limits = c(0,12), breaks = seq(0, 12, by = 2)) +
  scale_x_continuous(limits = c(0,12),breaks = seq(0, 12, by = 2)) +
  theme_bw())


pred.ob.comparison <- ggpubr::ggarrange(EH.pred.ob,
                                        GS1.pred.ob,
                                        GI1.pred.ob,
                                        GS2.pred.ob,
                                        LGM.pred.ob,
                                        LOIS3.pred.ob,
                                        EOIS3.pred.ob,
                                        ncol=2, nrow=4)

ggsave(file="Figure_5.pdf", pred.ob.comparison, width = 20, height = 30, units = c("cm"), dpi = 300)
ggsave(file="Figure_5.png", pred.ob.comparison, width = 20, height = 30, units = c("cm"), dpi = 300)


  ## CONSIDER CLIMATE-d15N RELATIONSHIPS FOR INCLUSION AS FIXED EFFECTS ####

## note tempmin and tempmax have names wrong way round in original data -> switch these
d15N.data.aggre <- rename(d15N.data.aggre, tempmin1 = tempmax)
d15N.data.aggre <- rename(d15N.data.aggre, tempmax1 = tempmin)

# create a correlation matrix table #
# x is a matrix containing the data
# method : correlation method. "pearson"" or "spearman"" is supported
# removeTriangle : remove upper or lower triangle
corstars <-function(x, method=c("pearson", "spearman"), removeTriangle=c("upper", "lower"),
                    result=c("none", "html", "latex")){
  #Compute correlation matrix
  require(Hmisc)
  x <- as.matrix(x)
  correlation_matrix<-rcorr(x, type=method[1])
  R <- correlation_matrix$r # Matrix of correlation coeficients
  p <- correlation_matrix$P # Matrix of p-value

  ## Define notions for significance levels; spacing is important.
  mystars <- ifelse(p < .0001, "****", ifelse(p < .001, "*** ", ifelse(p < .01, "**  ", ifelse(p < .05, "*   ", "    "))))

  ## trunctuate the correlation matrix to two decimal
  R <- format(round(cbind(rep(-1.11, ncol(x)), R), 2))[,-1]

  ## build a new matrix that includes the correlations with their apropriate stars
  Rnew <- matrix(paste(R, mystars, sep=""), ncol=ncol(x))
```

```
  diag(Rnew) <- paste(diag(R), " ", sep="")
  rownames(Rnew) <- colnames(x)
  colnames(Rnew) <- paste(colnames(x), "", sep="")

  ## remove upper triangle of correlation matrix
  if(removeTriangle[1]=="upper"){
    Rnew <- as.matrix(Rnew)
    Rnew[upper.tri(Rnew, diag = TRUE)] <- ""
    Rnew <- as.data.frame(Rnew)
  }

  ## remove lower triangle of correlation matrix
  else if(removeTriangle[1]=="lower"){
    Rnew <- as.matrix(Rnew)
    Rnew[lower.tri(Rnew, diag = TRUE)] <- ""
    Rnew <- as.data.frame(Rnew)
  }

  ## remove last column and return the correlation matrix
  Rnew <- cbind(Rnew[1:length(Rnew)-1])
  if (result[1]=="none") return(Rnew)
  else{
    if(result[1]=="html") print(xtable(Rnew), type="html")
    else print(xtable(Rnew), type="latex")
  }
}


correlation.matrix <- corstars(d15N.data.aggre[,c(4,6,9:25)], method = "pearson")
correlation.matrix

write.csv(correlation.matrix, file = "correlation_matrix_all_data.csv")


# correlation matrix by time period
EH.data <-  d15N.data.aggre %>%
  filter(finalagebin == 'EH')
correlation.matrix.EH <- corstars(EH.data[,c(4,6,9:25)], method = "pearson")
write.csv(correlation.matrix.EH, file = "correlation_matrix_EH.csv")

GS1.data <-  d15N.data.aggre %>%
  filter(finalagebin == 'YD')
correlation.matrix.GS1 <- corstars(GS1.data[,c(4,6,9:25)], method = "pearson")
write.csv(correlation.matrix.GS1, file = "correlation_matrix_GS1.csv")

GI1.data <-  d15N.data.aggre %>%
  filter(finalagebin == 'LGI')
correlation.matrix.GI1 <- corstars(GI1.data[,c(4,6,9:25)], method = "pearson")
write.csv(correlation.matrix.GI1, file = "correlation_matrix_GI1.csv")

GS2.data <-  d15N.data.aggre %>%
  filter(finalagebin == 'LGT')
correlation.matrix.GS2 <- corstars(GS2.data[,c(4,6,9:25)], method = "pearson")
write.csv(correlation.matrix.GS2, file = "correlation_matrix_GS2.csv")

LGM.data <-  d15N.data.aggre %>%
  filter(finalagebin == 'LGM')
correlation.matrix.LGM <- corstars(LGM.data[,c(4,6,9:25)], method = "pearson")
write.csv(correlation.matrix.LGM, file = "correlation_matrix_LGM.csv")

LOIS3.data <-  d15N.data.aggre %>%
  filter(finalagebin == 'LOIS3')
correlation.matrix.LOIS3 <- corstars(LOIS3.data[,c(4,6,9:25)], method = "pearson")
write.csv(correlation.matrix.LOIS3, file = "correlation_matrix_LOIS3.csv")

EOIS3.data <-  d15N.data.aggre %>%
  filter(finalagebin == 'EOIS3')
correlation.matrix.EOIS3 <- corstars(EOIS3.data[,c(4,6,9:25)], method = "pearson")
write.csv(correlation.matrix.EOIS3, file = "correlation_matrix_EOIS3.csv")

  ## EXAMINE COLLINEARITY ###
    # CORRELATION MATRIX PLOT ####
# from https://towardsdatascience.com/customizable-correlation-plots-in-r-b1d2856a4b05
correlation.data <- d15N.data.aggre[,c(9:25)]
cors <- function(df) {
  # turn all three matrices (r, n, and P into a data frame)
  M <- Hmisc::rcorr(as.matrix(df))
  # return the three data frames in a list return(Mdf)
  Mdf <- map(M, ~data.frame(.x))
}

formatted_cors <- function(df){
  cors(df) %>%
    map(~rownames_to_column(.x, var="measure1")) %>%
    map(~pivot_longer(.x, -measure1, "measure2")) %>%
    bind_rows(.id = "id") %>%
    pivot_wider(names_from = id, values_from = value) %>%
    mutate(sig_p = ifelse(P < .05, T, F), p_if_sig = ifelse(P <.05, P, NA), r_if_sig = ifelse(P <.05, r, NA))
}
formatted_cors(correlation.data) %>% head() %>% kable()

(correlationplot <- formatted_cors(correlation.data) %>%
  ggplot(aes(measure1, measure2, fill=r, label=round(r_if_sig,2))) +
  geom_tile() +
  labs(x = NULL, y = NULL, fill = "Pearson's\nCorrelation", title="Bioclimatic Correlations", subtitle="Only significant Pearson's
```

```
correlation coefficients shown") + scale_fill_gradient2(mid="#FBFEF9",low="#0C6291",high="#A63446", limits=c(-1,1)) +
  geom_text() +
  theme_classic() +
  scale_x_discrete(guide = guide_axis(n.dodge=2)))

ggsave(file="Figure_S3_1.pdf", correlationplot, width = 30, height = 30, units = c("cm"), dpi = 300)


    # RETAINED VARIABLES PLOTS WITH d15N####

# MAT by time period plot
(MAT.d15N.plot <-
   ggplot(d15N.data.aggre %>% bind_rows(d15N.data.aggre %>% mutate(finalagebin = "all")) ,
          aes(MAT, mean_source_value)) +
   geom_point(shape = 1) +
   ylab(expression(paste(delta^{15}, "N (\u2030)"))) +
   xlab("mean annual temperature") +
   facet_wrap(~fct_relevel(finalagebin, "EH", "YD", "LGI", "LGT", "LGM", "LOIS3", "EOIS3", "all", after = Inf), nrow=2) +
   stat_fit_glance(method = "cor.test",
                   label.y = "top",
                   method.args = list(formula = ~ x + y),
                   mapping = aes(label = sprintf('r[pearson]~"="~%.3f~", "~italic(p)~"="~%.2g',
                                                 after_stat(estimate), after_stat(p.value))),
                   parse = TRUE) +
   stat_poly_eq(aes(label = paste(..eq.label.., sep = "~~~")),
                label.x.npc = "left", label.y.npc = 0.85,
                formula = y ~ x, parse = TRUE) +
   geom_smooth(method = "lm", se=TRUE, color="red", formula = y ~ x) +
   theme_bw())


ggsave(file="Figure_S3_2.pdf", MAT.d15N.plot, width = 30, height = 20, units = c("cm"), dpi = 300)


# MAP by time period plot

(MAP.d15N.plot <-
    ggplot(d15N.data.aggre %>% bind_rows(d15N.data.aggre %>% mutate(finalagebin = "all")),
           aes(MAP, mean_source_value)) +
    geom_point(shape = 1) +
    ylab(expression(paste(delta^{15}, "N (\u2030)"))) +
    xlab("mean annual precipitation") +
    scale_x_continuous(trans='log10') +
    facet_wrap(~fct_relevel(finalagebin, "EH", "YD", "LGI", "LGT", "LGM", "LOIS3", "EOIS3", "all", after = Inf), nrow=2) +
    stat_fit_glance(method = "cor.test",
                    label.y = "top",
                    method.args = list(formula = ~ x + y),
                    mapping = aes(label = sprintf('r[pearson]~"="~%.3f~~italic(p)~"="~%.2g',
                                                  after_stat(estimate), after_stat(p.value))),
                    parse = TRUE) +
    stat_poly_eq(aes(label = paste(..eq.label.., sep = "~~~")),
                 label.x.npc = "left", label.y.npc = 0.85,
                 formula = y ~ x, parse = TRUE) +
    geom_smooth(method = "lm", se=TRUE, color="red", formula = y ~ x) +
    theme_bw())

ggsave(file="Figure_S3_3.pdf", MAP.d15N.plot, width = 30, height = 20, units = c("cm"), dpi = 300)




# temp.warmest by time period plot

(temp.warm.d15N.plot <-
    ggplot(d15N.data.aggre %>% bind_rows(d15N.data.aggre %>% mutate(finalagebin = "all")),
           aes(tempwarmest, mean_source_value)) +
    geom_point(shape = 1) +
    ylab(expression(paste(delta^{15}, "N (\u2030)"))) +
    xlab("mean temperature of the warmest quarter") +
    facet_wrap(~fct_relevel(finalagebin, "EH", "YD", "LGI", "LGT", "LGM", "LOIS3", "EOIS3", "all", after = Inf), nrow=2) +
    stat_fit_glance(method = "cor.test",
                    label.y = "top",
                    method.args = list(formula = ~ x + y),
                    mapping = aes(label = sprintf('r[pearson]~"="~%.3f~~italic(p)~"="~%.2g',
                                                  after_stat(estimate), after_stat(p.value))),
                    parse = TRUE) +
    stat_poly_eq(aes(label = paste(..eq.label.., sep = "~~~")),
                 label.x.npc = "left", label.y.npc = 0.85,
                 formula = y ~ x, parse = TRUE) +
    geom_smooth(method = "lm", se=TRUE, color="red", formula = y ~ x) +
    theme_bw())



ggsave(file="Figure_S3_4.pdf", temp.warm.d15N.plot, width = 30, height = 20, units = c("cm"), dpi = 300)


# precip.warm by time period plot

(precip.warm.d15N.plot <-
    ggplot(d15N.data.aggre %>% bind_rows(d15N.data.aggre %>% mutate(finalagebin = "all")),
           aes(precip.warm.quart, mean_source_value)) +
    geom_point(shape = 1) +
    scale_x_continuous(trans='log10') +
    ylab(expression(paste(delta^{15}, "N (\u2030)"))) +
```

```
    xlab("mean precipitation of the warmest quarter") +
    facet_wrap(~fct_relevel(finalagebin, "EH", "YD", "LGI", "LGT", "LGM", "LOIS3", "EOIS3", "all", after = Inf), nrow=2) +
    stat_fit_glance(method = "cor.test",
                    label.y = "top",
                    method.args = list(formula = ~ x + y),
                    mapping = aes(label = sprintf('r[pearson]~"="~%.3f~~italic(p)~"="~%.2g',
                                                  after_stat(estimate), after_stat(p.value))),
                    parse = TRUE) +
    stat_poly_eq(aes(label = paste(..eq.label.., sep = "~~~")),
                 label.x.npc = "left", label.y.npc = 0.85,
                 formula = y ~ x, parse = TRUE) +
    geom_smooth(method = "lm", se=TRUE, color="red", formula = y ~ x) +
    theme_bw())

ggsave(file="Figure_S3_5.pdf", precip.warm.d15N.plot, width = 30, height = 20, units = c("cm"), dpi = 300)



# precip.cold by time period plot

(precip.cold.d15N.plot <-
    ggplot(d15N.data.aggre %>% bind_rows(d15N.data.aggre %>% mutate(finalagebin = "all")),
           aes(precip.cold.quart, mean_source_value)) +
    geom_point(shape = 1) +
    scale_x_continuous(trans='log10') +
    ylab(expression(paste(delta^{15}, "N (\u2030)"))) +
    xlab("mean precipitation of the coldest quarter") +
    facet_wrap(~fct_relevel(finalagebin, "EH", "YD", "LGI", "LGT", "LGM", "LOIS3", "EOIS3", "all", after = Inf), nrow=2) +
    stat_fit_glance(method = "cor.test",
                    label.y = "top",
                    method.args = list(formula = ~ x + y),
                    mapping = aes(label = sprintf('r[pearson]~"="~%.3f~~italic(p)~"="~%.2g',
                                                  after_stat(estimate), after_stat(p.value))),
                    parse = TRUE) +
    stat_poly_eq(aes(label = paste(..eq.label.., sep = "~~~")),
                 label.x.npc = "left", label.y.npc = 0.85,
                 formula = y ~ x, parse = TRUE) +
    geom_smooth(method = "lm", se=TRUE, color="red", formula = y ~ x) +
    theme_bw())

ggsave(file="Figure_S3_6.pdf", precip.cold.d15N.plot, width = 30, height = 20, units = c("cm"), dpi = 300)



  ## MIXED MODEL INTERPOLATION WITH FIXED EFFECTS ####
    ## set up dataframe
      # based on the correlation testing we are only keeping the following environmental variables:
      # MAT, MAP, tempdriest, precip.warm.quart and precip.cold.quart
      # keep only the relevant columns
        d15N.data.aggre <- d15N.data.aggre %>%
          subset(select = c("SiteName", "lat", "long", "n_source_value", "mean_source_value",
                    "var_source_value","finalagebin",  "MAP", "MAT",
                    "tempwarmest", "precip.warm.quart", "precip.cold.quart"))


# then split df by age bin
EH <- d15N.data.aggre %>% filter(finalagebin == "EH")
GS1 <- d15N.data.aggre %>% filter(finalagebin == "YD")
GI1 <- d15N.data.aggre %>% filter(finalagebin == "LGI")
GS2 <- d15N.data.aggre %>% filter(finalagebin == "LGT")
LGM <- d15N.data.aggre %>% filter(finalagebin == "LGM")
LOIS3 <- d15N.data.aggre %>% filter(finalagebin == "LOIS3")
EOIS3 <- d15N.data.aggre %>% filter(finalagebin == "EOIS3")

#scale environmental variables for each dataframe
EH <- EH %>%
  mutate_at(c("MAP", "MAT",
              "tempwarmest", "precip.warm.quart", "precip.cold.quart"), ~(scale(.) %>% as.vector))
GS1 <- GS1 %>%
  mutate_at(c("MAP", "MAT",
              "tempwarmest", "precip.warm.quart", "precip.cold.quart"), ~(scale(.) %>% as.vector))
GI1 <- GI1 %>%
  mutate_at(c("MAP", "MAT",
              "tempwarmest", "precip.warm.quart", "precip.cold.quart"), ~(scale(.) %>% as.vector))
GS2 <- GS2 %>%
  mutate_at(c("MAP", "MAT",
              "tempwarmest", "precip.warm.quart", "precip.cold.quart"), ~(scale(.) %>% as.vector))
LGM <- LGM %>%
  mutate_at(c("MAP", "MAT",
              "tempwarmest", "precip.warm.quart", "precip.cold.quart"), ~(scale(.) %>% as.vector))
LOIS3 <- LOIS3  %>%
  mutate_at(c("MAP", "MAT",
              "tempwarmest", "precip.warm.quart", "precip.cold.quart"), ~(scale(.) %>% as.vector))
EOIS3 <- EOIS3 %>%
  mutate_at(c("MAP", "MAT",
              "tempwarmest", "precip.warm.quart", "precip.cold.quart"), ~(scale(.) %>% as.vector))


# Fit the residual dispersion model (following Courtiol and Rousset 2017) #
EH.dispfit <- fitme(formula = var_source_value ~ 1 + Matern(1|long + lat) + (1|SiteName),
                    family = Gamma(link = log), data = EH, fixed = list(phi = 2),
                    prior.weights = n_source_value - 1, control.dist = list(dist.method = "Earth"), method = "REML")
GS1.dispfit <- fitme(formula = var_source_value ~ 1 + Matern(1|long + lat) + (1|SiteName),
                     family = Gamma(link = log), data = GS1, fixed = list(phi = 2),
                     prior.weights = n_source_value - 1, control.dist = list(dist.method = "Earth"), method = "REML")
```

```
GI1.dispfit <- fitme(formula = var_source_value ~ 1 + Matern(1|long + lat) + (1|SiteName),
                     family = Gamma(link = log), data = GI1, fixed = list(phi = 2),
                     prior.weights = n_source_value - 1, control.dist = list(dist.method = "Earth"), method = "REML")
GS2.dispfit <- fitme(formula = var_source_value ~ 1 + Matern(1|long + lat) + (1|SiteName),
                     family = Gamma(link = log), data = GS2, fixed = list(phi = 2),
                     prior.weights = n_source_value - 1, control.dist = list(dist.method = "Earth"), method = "REML")
LGM.dispfit <- fitme(formula = var_source_value ~ 1 + Matern(1|long + lat) + (1|SiteName),
                     family = Gamma(link = log), data = LGM, fixed = list(phi = 2),
                     prior.weights = n_source_value - 1, control.dist = list(dist.method = "Earth"), method = "REML")
LOIS3.dispfit <- fitme(formula = var_source_value ~ 1 + Matern(1|long + lat) + (1|SiteName),
                       family = Gamma(link = log), data = LOIS3, fixed = list(phi = 2),
                       prior.weights = n_source_value - 1, control.dist = list(dist.method = "Earth"), method = "REML")
EOIS3.dispfit <- fitme(formula = var_source_value ~ 1 + Matern(1|long + lat) + (1|SiteName),
                       family = Gamma(link = log), data = EOIS3, fixed = list(phi = 2),
                       prior.weights = n_source_value - 1, control.dist = list(dist.method = "Earth"), method = "REML")

# predict Ïg of the expected square of the residual error in each location using the fit of the residual dispersion model:
EH$disp <- predict(EH.dispfit, newdata = EH)[,1]
GS1$disp <- predict(GS1.dispfit, newdata = GS1)[,1]
GI1$disp <- predict(GI1.dispfit, newdata = GI1)[,1]
GS2$disp <- predict(GS2.dispfit, newdata = GS2)[,1]
LGM$disp <- predict(LGM.dispfit, newdata = LGM)[,1]
LOIS3$disp <- predict(LOIS3.dispfit, newdata = LOIS3)[,1]
EOIS3$disp <- predict(EOIS3.dispfit, newdata = EOIS3)[,1]


    # EARLY HOLOCENE FIXED EFFECTS MIXED MODELS ####

# MAT
EHmeanfit2 <- fitme(formula = mean_source_value ~ 1 + MAT + Matern(1|long + lat) + (1|SiteName),
                    family = gaussian(link = identity), data = EH,
                    resid.model = list(formula= ~ 0 + offset(disp), family = Gamma(link = identity)),
                    prior.weights = n_source_value, control.dist = list(dist.method = "Earth"), method = "REML")
# MAP
EHmeanfit3 <- fitme(formula = mean_source_value ~ 1 + MAP + Matern(1|long + lat) + (1|SiteName),
                    family = gaussian(link = identity), data = EH,
                    resid.model = list(formula= ~ 0 + offset(disp), family = Gamma(link = identity)),
                    prior.weights = n_source_value, control.dist = list(dist.method = "Earth"), method = "REML")

# MAT + MAP
EHmeanfit4 <- fitme(formula = mean_source_value ~ 1 + MAT + MAP + Matern(1|long + lat) + (1|SiteName),
                    family = gaussian(link = identity), data = EH,
                    resid.model = list(formula= ~ 0 + offset(disp), family = Gamma(link = identity)),
                    prior.weights = n_source_value, control.dist = list(dist.method = "Earth"), method = "REML")

# MAT + MAP + MAT:MAP
EHmeanfit5 <- fitme(formula = mean_source_value ~ 1 + MAT + MAP + MAT:MAP+ Matern(1|long + lat) + (1|SiteName),
                    family = gaussian(link = identity), data = EH,
                    resid.model = list(formula= ~ 0 + offset(disp), family = Gamma(link = identity)),
                    prior.weights = n_source_value, control.dist = list(dist.method = "Earth"), method = "REML")

# tempwarmest
EHmeanfit6 <- fitme(formula = mean_source_value ~ 1 + tempwarmest + Matern(1|long + lat) + (1|SiteName),
                    family = gaussian(link = identity), data = EH,
                    resid.model = list(formula= ~ 0 + offset(disp), family = Gamma(link = identity)),
                    prior.weights = n_source_value, control.dist = list(dist.method = "Earth"), method = "REML")

# precipwarmest
EHmeanfit7 <- fitme(formula = mean_source_value ~ 1 + precip.warm.quart + Matern(1|long + lat) + (1|SiteName),
                    family = gaussian(link = identity), data = EH,
                    resid.model = list(formula= ~ 0 + offset(disp), family = Gamma(link = identity)),
                    prior.weights = n_source_value, control.dist = list(dist.method = "Earth"), method = "REML")


# precipcoldest
EHmeanfit8 <- fitme(formula = mean_source_value ~ 1 + precip.cold.quart + Matern(1|long + lat) + (1|SiteName),
                    family = gaussian(link = identity), data = EH,
                    resid.model = list(formula= ~ 0 + offset(disp), family = Gamma(link = identity)),
                    prior.weights = n_source_value, control.dist = list(dist.method = "Earth"), method = "REML")

# tempwarmest + precip.warm.quart
EHmeanfit9 <- fitme(formula = mean_source_value ~ 1 + tempwarmest + precip.warm.quart + Matern(1|long + lat) + (1|SiteName),
                    family = gaussian(link = identity), data = EH,
                    resid.model = list(formula= ~ 0 + offset(disp), family = Gamma(link = identity)),
                    prior.weights = n_source_value, control.dist = list(dist.method = "Earth"), method = "REML")

# tempwarmest + precip.cold.quart
EHmeanfit10 <- fitme(formula = mean_source_value ~ 1 + tempwarmest + precip.cold.quart + Matern(1|long + lat) + (1|SiteName),
                     family = gaussian(link = identity), data = EH,
                     resid.model = list(formula= ~ 0 + offset(disp), family = Gamma(link = identity)),
                     prior.weights = n_source_value, control.dist = list(dist.method = "Earth"), method = "REML")

# precip.cold.quart + precip.warm.quart
EHmeanfit11 <- fitme(formula = mean_source_value ~ 1 + precip.cold.quart + precip.warm.quart + Matern(1|long + lat) + (1|SiteName),
                     family = gaussian(link = identity), data = EH,
                     resid.model = list(formula= ~ 0 + offset(disp), family = Gamma(link = identity)),
                     prior.weights = n_source_value, control.dist = list(dist.method = "Earth"), method = "REML")


# tempwarmest + precip.cold.quart +precip.warm.quart
EHmeanfit12 <- fitme(formula = mean_source_value ~ 1 + tempwarmest  + precip.cold.quart + precip.warm.quart + Matern(1|long + lat) +
(1|SiteName),
                     family = gaussian(link = identity), data = EH,
                     resid.model = list(formula= ~ 0 + offset(disp), family = Gamma(link = identity)),
                     prior.weights = n_source_value, control.dist = list(dist.method = "Earth"), method = "REML")
```

```
AIC(EHmeanfit1)
AIC(EHmeanfit2)
AIC(EHmeanfit3)
AIC(EHmeanfit4)
AIC(EHmeanfit5)
AIC(EHmeanfit6)
AIC(EHmeanfit7)
AIC(EHmeanfit8)
AIC(EHmeanfit9)
AIC(EHmeanfit10)
AIC(EHmeanfit11)
AIC(EHmeanfit12)


    # YOUNGER DRYAS FIXED EFFECTS MIXED MODELS ####

# MAT
GS1meanfit2 <- fitme(formula = mean_source_value ~ 1 + MAT + Matern(1|long + lat) + (1|SiteName),
                     family = gaussian(link = identity), data = GS1,
                     resid.model = list(formula= ~ 0 + offset(disp), family = Gamma(link = identity)),
                     prior.weights = n_source_value, control.dist = list(dist.method = "Earth"), method = "REML")
# MAP
GS1meanfit3 <- fitme(formula = mean_source_value ~ 1 + MAP + Matern(1|long + lat) + (1|SiteName),
                     family = gaussian(link = identity), data = GS1,
                     resid.model = list(formula= ~ 0 + offset(disp), family = Gamma(link = identity)),
                     prior.weights = n_source_value, control.dist = list(dist.method = "Earth"), method = "REML")

# MAT + MAP
GS1meanfit4 <- fitme(formula = mean_source_value ~ 1 + MAT + MAP + Matern(1|long + lat) + (1|SiteName),
                     family = gaussian(link = identity), data = GS1,
                     resid.model = list(formula= ~ 0 + offset(disp), family = Gamma(link = identity)),
                     prior.weights = n_source_value, control.dist = list(dist.method = "Earth"), method = "REML")

# MAT + MAP + MAT:MAP
GS1meanfit5 <- fitme(formula = mean_source_value ~ 1 + MAT + MAP + MAT:MAP+ Matern(1|long + lat) + (1|SiteName),
                     family = gaussian(link = identity), data = GS1,
                     resid.model = list(formula= ~ 0 + offset(disp), family = Gamma(link = identity)),
                     prior.weights = n_source_value, control.dist = list(dist.method = "Earth"), method = "REML")

# tempwarmest
GS1meanfit6 <- fitme(formula = mean_source_value ~ 1 + tempwarmest + Matern(1|long + lat) + (1|SiteName),
                     family = gaussian(link = identity), data = GS1,
                     resid.model = list(formula= ~ 0 + offset(disp), family = Gamma(link = identity)),
                     prior.weights = n_source_value, control.dist = list(dist.method = "Earth"), method = "REML")

# precipwarmest
GS1meanfit7 <- fitme(formula = mean_source_value ~ 1 + precip.warm.quart + Matern(1|long + lat) + (1|SiteName),
                     family = gaussian(link = identity), data = GS1,
                     resid.model = list(formula= ~ 0 + offset(disp), family = Gamma(link = identity)),
                     prior.weights = n_source_value, control.dist = list(dist.method = "Earth"), method = "REML")


# precipcoldest
GS1meanfit8 <- fitme(formula = mean_source_value ~ 1 + precip.cold.quart + Matern(1|long + lat) + (1|SiteName),
                     family = gaussian(link = identity), data = GS1,
                     resid.model = list(formula= ~ 0 + offset(disp), family = Gamma(link = identity)),
                     prior.weights = n_source_value, control.dist = list(dist.method = "Earth"), method = "REML")

# tempwarmest + precip.warm.quart
GS1meanfit9 <- fitme(formula = mean_source_value ~ 1 + tempwarmest + precip.warm.quart + Matern(1|long + lat) + (1|SiteName),
                     family = gaussian(link = identity), data = GS1,
                     resid.model = list(formula= ~ 0 + offset(disp), family = Gamma(link = identity)),
                     prior.weights = n_source_value, control.dist = list(dist.method = "Earth"), method = "REML")

# tempwarmest + precip.cold.quart
GS1meanfit10 <- fitme(formula = mean_source_value ~ 1 + tempwarmest + precip.cold.quart + Matern(1|long + lat) + (1|SiteName),
                      family = gaussian(link = identity), data = GS1,
                      resid.model = list(formula= ~ 0 + offset(disp), family = Gamma(link = identity)),
                      prior.weights = n_source_value, control.dist = list(dist.method = "Earth"), method = "REML")

# precip.cold.quart + precip.warm.quart
GS1meanfit11 <- fitme(formula = mean_source_value ~ 1 + precip.cold.quart + precip.warm.quart + Matern(1|long + lat) + (1|SiteName),
                      family = gaussian(link = identity), data = GS1,
                      resid.model = list(formula= ~ 0 + offset(disp), family = Gamma(link = identity)),
                      prior.weights = n_source_value, control.dist = list(dist.method = "Earth"), method = "REML")


# tempwarmest + precip.cold.quart +precip.warm.quart
GS1meanfit12 <- fitme(formula = mean_source_value ~ 1 + tempwarmest  + precip.cold.quart + precip.warm.quart + Matern(1|long + lat)
+ (1|SiteName),
                      family = gaussian(link = identity), data = GS1,
                      resid.model = list(formula= ~ 0 + offset(disp), family = Gamma(link = identity)),
                      prior.weights = n_source_value, control.dist = list(dist.method = "Earth"), method = "REML")


AIC(GS1meanfit1)
AIC(GS1meanfit2)
AIC(GS1meanfit3)
AIC(GS1meanfit4)
AIC(GS1meanfit5)
AIC(GS1meanfit6)
AIC(GS1meanfit7)
AIC(GS1meanfit8)
```

```
AIC(GS1meanfit9)
AIC(GS1meanfit10)
AIC(GS1meanfit11)
AIC(GS1meanfit12)




    # LATE GLACIAL INTERSTADIAL FIXED EFFECTS MIXED MODELS ####

# MAT
GI1meanfit2 <- fitme(formula = mean_source_value ~ 1 + MAT + Matern(1|long + lat) + (1|SiteName),
                     family = gaussian(link = identity), data = GI1,
                     resid.model = list(formula= ~ 0 + offset(disp), family = Gamma(link = identity)),
                     prior.weights = n_source_value, control.dist = list(dist.method = "Earth"), method = "REML")
# MAP
GI1meanfit3 <- fitme(formula = mean_source_value ~ 1 + MAP + Matern(1|long + lat) + (1|SiteName),
                     family = gaussian(link = identity), data = GI1,
                     resid.model = list(formula= ~ 0 + offset(disp), family = Gamma(link = identity)),
                     prior.weights = n_source_value, control.dist = list(dist.method = "Earth"), method = "REML")

# MAT + MAP
GI1meanfit4 <- fitme(formula = mean_source_value ~ 1 + MAT + MAP + Matern(1|long + lat) + (1|SiteName),
                     family = gaussian(link = identity), data = GI1,
                     resid.model = list(formula= ~ 0 + offset(disp), family = Gamma(link = identity)),
                     prior.weights = n_source_value, control.dist = list(dist.method = "Earth"), method = "REML")

# MAT + MAP + MAT:MAP
GI1meanfit5 <- fitme(formula = mean_source_value ~ 1 + MAT + MAP + MAT:MAP+ Matern(1|long + lat) + (1|SiteName),
                     family = gaussian(link = identity), data = GI1,
                     resid.model = list(formula= ~ 0 + offset(disp), family = Gamma(link = identity)),
                     prior.weights = n_source_value, control.dist = list(dist.method = "Earth"), method = "REML")

# tempwarmest
GI1meanfit6 <- fitme(formula = mean_source_value ~ 1 + tempwarmest + Matern(1|long + lat) + (1|SiteName),
                     family = gaussian(link = identity), data = GI1,
                     resid.model = list(formula= ~ 0 + offset(disp), family = Gamma(link = identity)),
                     prior.weights = n_source_value, control.dist = list(dist.method = "Earth"), method = "REML")

# precipwarmest
GI1meanfit7 <- fitme(formula = mean_source_value ~ 1 + precip.warm.quart + Matern(1|long + lat) + (1|SiteName),
                     family = gaussian(link = identity), data = GI1,
                     resid.model = list(formula= ~ 0 + offset(disp), family = Gamma(link = identity)),
                     prior.weights = n_source_value, control.dist = list(dist.method = "Earth"), method = "REML")


# precipcoldest
GI1meanfit8 <- fitme(formula = mean_source_value ~ 1 + precip.cold.quart + Matern(1|long + lat) + (1|SiteName),
                     family = gaussian(link = identity), data = GI1,
                     resid.model = list(formula= ~ 0 + offset(disp), family = Gamma(link = identity)),
                     prior.weights = n_source_value, control.dist = list(dist.method = "Earth"), method = "REML")

# tempwarmest + precip.warm.quart
GI1meanfit9 <- fitme(formula = mean_source_value ~ 1 + tempwarmest + precip.warm.quart + Matern(1|long + lat) + (1|SiteName),
                     family = gaussian(link = identity), data = GI1,
                     resid.model = list(formula= ~ 0 + offset(disp), family = Gamma(link = identity)),
                     prior.weights = n_source_value, control.dist = list(dist.method = "Earth"), method = "REML")

# tempwarmest + precip.cold.quart
GI1meanfit10 <- fitme(formula = mean_source_value ~ 1 + tempwarmest + precip.cold.quart + Matern(1|long + lat) + (1|SiteName),
                      family = gaussian(link = identity), data = GI1,
                      resid.model = list(formula= ~ 0 + offset(disp), family = Gamma(link = identity)),
                      prior.weights = n_source_value, control.dist = list(dist.method = "Earth"), method = "REML")

# precip.cold.quart + precip.warm.quart
GI1meanfit11 <- fitme(formula = mean_source_value ~ 1 + precip.cold.quart + precip.warm.quart + Matern(1|long + lat) + (1|SiteName),
                      family = gaussian(link = identity), data = GI1,
                      resid.model = list(formula= ~ 0 + offset(disp), family = Gamma(link = identity)),
                      prior.weights = n_source_value, control.dist = list(dist.method = "Earth"), method = "REML")


# tempwarmest + precip.cold.quart +precip.warm.quart
GI1meanfit12 <- fitme(formula = mean_source_value ~ 1 + tempwarmest  + precip.cold.quart + precip.warm.quart + Matern(1|long + lat)
+ (1|SiteName),
                      family = gaussian(link = identity), data = GI1,
                      resid.model = list(formula= ~ 0 + offset(disp), family = Gamma(link = identity)),
                      prior.weights = n_source_value, control.dist = list(dist.method = "Earth"), method = "REML")


AIC(GI1meanfit1)
AIC(GI1meanfit2)
AIC(GI1meanfit3)
AIC(GI1meanfit4)
AIC(GI1meanfit5)
AIC(GI1meanfit6)
AIC(GI1meanfit7)
AIC(GI1meanfit8)
AIC(GI1meanfit9)
AIC(GI1meanfit10)
AIC(GI1meanfit11)
AIC(GI1meanfit12)



    # LAST GLACIAL TERMINATION FIXED EFFECTS MIXED MODELS ####
```

```
# MAT
GS2meanfit2 <- fitme(formula = mean_source_value ~ 1 + MAT + Matern(1|long + lat) + (1|SiteName),
                     family = gaussian(link = identity), data = GS2,
                     resid.model = list(formula= ~ 0 + offset(disp), family = Gamma(link = identity)),
                     prior.weights = n_source_value, control.dist = list(dist.method = "Earth"), method = "REML")
# MAP
GS2meanfit3 <- fitme(formula = mean_source_value ~ 1 + MAP + Matern(1|long + lat) + (1|SiteName),
                     family = gaussian(link = identity), data = GS2,
                     resid.model = list(formula= ~ 0 + offset(disp), family = Gamma(link = identity)),
                     prior.weights = n_source_value, control.dist = list(dist.method = "Earth"), method = "REML")

# MAT + MAP
GS2meanfit4 <- fitme(formula = mean_source_value ~ 1 + MAT + MAP + Matern(1|long + lat) + (1|SiteName),
                     family = gaussian(link = identity), data = GS2,
                     resid.model = list(formula= ~ 0 + offset(disp), family = Gamma(link = identity)),
                     prior.weights = n_source_value, control.dist = list(dist.method = "Earth"), method = "REML")

# MAT + MAP + MAT:MAP
GS2meanfit5 <- fitme(formula = mean_source_value ~ 1 + MAT + MAP + MAT:MAP+ Matern(1|long + lat) + (1|SiteName),
                     family = gaussian(link = identity), data = GS2,
                     resid.model = list(formula= ~ 0 + offset(disp), family = Gamma(link = identity)),
                     prior.weights = n_source_value, control.dist = list(dist.method = "Earth"), method = "REML")

# tempwarmest
GS2meanfit6 <- fitme(formula = mean_source_value ~ 1 + tempwarmest + Matern(1|long + lat) + (1|SiteName),
                     family = gaussian(link = identity), data = GS2,
                     resid.model = list(formula= ~ 0 + offset(disp), family = Gamma(link = identity)),
                     prior.weights = n_source_value, control.dist = list(dist.method = "Earth"), method = "REML")

# precipwarmest
GS2meanfit7 <- fitme(formula = mean_source_value ~ 1 + precip.warm.quart + Matern(1|long + lat) + (1|SiteName),
                     family = gaussian(link = identity), data = GS2,
                     resid.model = list(formula= ~ 0 + offset(disp), family = Gamma(link = identity)),
                     prior.weights = n_source_value, control.dist = list(dist.method = "Earth"), method = "REML")


# precipcoldest
GS2meanfit8 <- fitme(formula = mean_source_value ~ 1 + precip.cold.quart + Matern(1|long + lat) + (1|SiteName),
                     family = gaussian(link = identity), data = GS2,
                     resid.model = list(formula= ~ 0 + offset(disp), family = Gamma(link = identity)),
                     prior.weights = n_source_value, control.dist = list(dist.method = "Earth"), method = "REML")

# tempwarmest + precip.warm.quart
GS2meanfit9 <- fitme(formula = mean_source_value ~ 1 + tempwarmest + precip.warm.quart + Matern(1|long + lat) + (1|SiteName),
                     family = gaussian(link = identity), data = GS2,
                     resid.model = list(formula= ~ 0 + offset(disp), family = Gamma(link = identity)),
                     prior.weights = n_source_value, control.dist = list(dist.method = "Earth"), method = "REML")

# tempwarmest + precip.cold.quart
GS2meanfit10 <- fitme(formula = mean_source_value ~ 1 + tempwarmest + precip.cold.quart + Matern(1|long + lat) + (1|SiteName),
                      family = gaussian(link = identity), data = GS2,
                      resid.model = list(formula= ~ 0 + offset(disp), family = Gamma(link = identity)),
                      prior.weights = n_source_value, control.dist = list(dist.method = "Earth"), method = "REML")

# precip.cold.quart + precip.warm.quart
GS2meanfit11 <- fitme(formula = mean_source_value ~ 1 + precip.cold.quart + precip.warm.quart + Matern(1|long + lat) + (1|SiteName),
                      family = gaussian(link = identity), data = GS2,
                      resid.model = list(formula= ~ 0 + offset(disp), family = Gamma(link = identity)),
                      prior.weights = n_source_value, control.dist = list(dist.method = "Earth"), method = "REML")


# tempwarmest + precip.cold.quart +precip.warm.quart
GS2meanfit12 <- fitme(formula = mean_source_value ~ 1 + tempwarmest  + precip.cold.quart + precip.warm.quart + Matern(1|long + lat)
+ (1|SiteName),
                      family = gaussian(link = identity), data = GS2,
                      resid.model = list(formula= ~ 0 + offset(disp), family = Gamma(link = identity)),
                      prior.weights = n_source_value, control.dist = list(dist.method = "Earth"), method = "REML")


AIC(GS2meanfit1)
AIC(GS2meanfit2)
AIC(GS2meanfit3)
AIC(GS2meanfit4)
AIC(GS2meanfit5)
AIC(GS2meanfit6)
AIC(GS2meanfit7)
AIC(GS2meanfit8)
AIC(GS2meanfit9)
AIC(GS2meanfit10)
AIC(GS2meanfit11)
AIC(GS2meanfit12)



    # LAST GLACIAL MAXIMUM FIXED EFFECTS MIXED MODELS ####

# MAT
LGMmeanfit2 <- fitme(formula = mean_source_value ~ 1 + MAT + Matern(1|long + lat) + (1|SiteName),
                     family = gaussian(link = identity), data = LGM,
                     resid.model = list(formula= ~ 0 + offset(disp), family = Gamma(link = identity)),
                     prior.weights = n_source_value, control.dist = list(dist.method = "Earth"), method = "REML")
# MAP
LGMmeanfit3 <- fitme(formula = mean_source_value ~ 1 + MAP + Matern(1|long + lat) + (1|SiteName),
                     family = gaussian(link = identity), data = LGM,
```

```
                     resid.model = list(formula= ~ 0 + offset(disp), family = Gamma(link = identity)),
                     prior.weights = n_source_value, control.dist = list(dist.method = "Earth"), method = "REML")

# MAT + MAP
LGMmeanfit4 <- fitme(formula = mean_source_value ~ 1 + MAT + MAP + Matern(1|long + lat) + (1|SiteName),
                     family = gaussian(link = identity), data = LGM,
                     resid.model = list(formula= ~ 0 + offset(disp), family = Gamma(link = identity)),
                     prior.weights = n_source_value, control.dist = list(dist.method = "Earth"), method = "REML")

# MAT + MAP + MAT:MAP
LGMmeanfit5 <- fitme(formula = mean_source_value ~ 1 + MAT + MAP + MAT:MAP+ Matern(1|long + lat) + (1|SiteName),
                     family = gaussian(link = identity), data = LGM,
                     resid.model = list(formula= ~ 0 + offset(disp), family = Gamma(link = identity)),
                     prior.weights = n_source_value, control.dist = list(dist.method = "Earth"), method = "REML")

# tempwarmest
LGMmeanfit6 <- fitme(formula = mean_source_value ~ 1 + tempwarmest + Matern(1|long + lat) + (1|SiteName),
                     family = gaussian(link = identity), data = LGM,
                     resid.model = list(formula= ~ 0 + offset(disp), family = Gamma(link = identity)),
                     prior.weights = n_source_value, control.dist = list(dist.method = "Earth"), method = "REML")

# precipwarmest
LGMmeanfit7 <- fitme(formula = mean_source_value ~ 1 + precip.warm.quart + Matern(1|long + lat) + (1|SiteName),
                     family = gaussian(link = identity), data = LGM,
                     resid.model = list(formula= ~ 0 + offset(disp), family = Gamma(link = identity)),
                     prior.weights = n_source_value, control.dist = list(dist.method = "Earth"), method = "REML")


# precipcoldest
LGMmeanfit8 <- fitme(formula = mean_source_value ~ 1 + precip.cold.quart + Matern(1|long + lat) + (1|SiteName),
                     family = gaussian(link = identity), data = LGM,
                     resid.model = list(formula= ~ 0 + offset(disp), family = Gamma(link = identity)),
                     prior.weights = n_source_value, control.dist = list(dist.method = "Earth"), method = "REML")

# tempwarmest + precip.warm.quart
LGMmeanfit9 <- fitme(formula = mean_source_value ~ 1 + tempwarmest + precip.warm.quart + Matern(1|long + lat) + (1|SiteName),
                     family = gaussian(link = identity), data = LGM,
                     resid.model = list(formula= ~ 0 + offset(disp), family = Gamma(link = identity)),
                     prior.weights = n_source_value, control.dist = list(dist.method = "Earth"), method = "REML")

# tempwarmest + precip.cold.quart
LGMmeanfit10 <- fitme(formula = mean_source_value ~ 1 + tempwarmest + precip.cold.quart + Matern(1|long + lat) + (1|SiteName),
                      family = gaussian(link = identity), data = LGM,
                      resid.model = list(formula= ~ 0 + offset(disp), family = Gamma(link = identity)),
                      prior.weights = n_source_value, control.dist = list(dist.method = "Earth"), method = "REML")

# precip.cold.quart + precip.warm.quart
LGMmeanfit11 <- fitme(formula = mean_source_value ~ 1 + precip.cold.quart + precip.warm.quart + Matern(1|long + lat) + (1|SiteName),
                      family = gaussian(link = identity), data = LGM,
                      resid.model = list(formula= ~ 0 + offset(disp), family = Gamma(link = identity)),
                      prior.weights = n_source_value, control.dist = list(dist.method = "Earth"), method = "REML")


# tempwarmest + precip.cold.quart +precip.warm.quart
LGMmeanfit12 <- fitme(formula = mean_source_value ~ 1 + tempwarmest  + precip.cold.quart + precip.warm.quart + Matern(1|long + lat)
+ (1|SiteName),
                      family = gaussian(link = identity), data = LGM,
                      resid.model = list(formula= ~ 0 + offset(disp), family = Gamma(link = identity)),
                      prior.weights = n_source_value, control.dist = list(dist.method = "Earth"), method = "REML")


AIC(LGMmeanfit1)
AIC(LGMmeanfit2)
AIC(LGMmeanfit3)
AIC(LGMmeanfit4)
AIC(LGMmeanfit5)
AIC(LGMmeanfit6)
AIC(LGMmeanfit7)
AIC(LGMmeanfit8)
AIC(LGMmeanfit9)
AIC(LGMmeanfit10)
AIC(LGMmeanfit11)
AIC(LGMmeanfit12)


    # LATE OIS 3 FIXED EFFECTS MIXED MODELS ####

# MAT
LOIS3meanfit2 <- fitme(formula = mean_source_value ~ 1 + MAT + Matern(1|long + lat) + (1|SiteName),
                       family = gaussian(link = identity), data = LOIS3,
                       resid.model = list(formula= ~ 0 + offset(disp), family = Gamma(link = identity)),
                       prior.weights = n_source_value, control.dist = list(dist.method = "Earth"), method = "REML")
# MAP
LOIS3meanfit3 <- fitme(formula = mean_source_value ~ 1 + MAP + Matern(1|long + lat) + (1|SiteName),
                       family = gaussian(link = identity), data = LOIS3,
                       resid.model = list(formula= ~ 0 + offset(disp), family = Gamma(link = identity)),
                       prior.weights = n_source_value, control.dist = list(dist.method = "Earth"), method = "REML")

# MAT + MAP
LOIS3meanfit4 <- fitme(formula = mean_source_value ~ 1 + MAT + MAP + Matern(1|long + lat) + (1|SiteName),
                       family = gaussian(link = identity), data = LOIS3,
                       resid.model = list(formula= ~ 0 + offset(disp), family = Gamma(link = identity)),
                       prior.weights = n_source_value, control.dist = list(dist.method = "Earth"), method = "REML")

# MAT + MAP + MAT:MAP
```

```
LOIS3meanfit5 <- fitme(formula = mean_source_value ~ 1 + MAT + MAP + MAT:MAP+ Matern(1|long + lat) + (1|SiteName),
                       family = gaussian(link = identity), data = LOIS3,
                       resid.model = list(formula= ~ 0 + offset(disp), family = Gamma(link = identity)),
                       prior.weights = n_source_value, control.dist = list(dist.method = "Earth"), method = "REML")

# tempwarmest
LOIS3meanfit6 <- fitme(formula = mean_source_value ~ 1 + tempwarmest + Matern(1|long + lat) + (1|SiteName),
                       family = gaussian(link = identity), data = LOIS3,
                       resid.model = list(formula= ~ 0 + offset(disp), family = Gamma(link = identity)),
                       prior.weights = n_source_value, control.dist = list(dist.method = "Earth"), method = "REML")

# precipwarmest
LOIS3meanfit7 <- fitme(formula = mean_source_value ~ 1 + precip.warm.quart + Matern(1|long + lat) + (1|SiteName),
                       family = gaussian(link = identity), data = LOIS3,
                       resid.model = list(formula= ~ 0 + offset(disp), family = Gamma(link = identity)),
                       prior.weights = n_source_value, control.dist = list(dist.method = "Earth"), method = "REML")


# precipcoldest
LOIS3meanfit8 <- fitme(formula = mean_source_value ~ 1 + precip.cold.quart + Matern(1|long + lat) + (1|SiteName),
                       family = gaussian(link = identity), data = LOIS3,
                       resid.model = list(formula= ~ 0 + offset(disp), family = Gamma(link = identity)),
                       prior.weights = n_source_value, control.dist = list(dist.method = "Earth"), method = "REML")

# tempwarmest + precip.warm.quart
LOIS3meanfit9 <- fitme(formula = mean_source_value ~ 1 + tempwarmest + precip.warm.quart + Matern(1|long + lat) + (1|SiteName),
                       family = gaussian(link = identity), data = LOIS3,
                       resid.model = list(formula= ~ 0 + offset(disp), family = Gamma(link = identity)),
                       prior.weights = n_source_value, control.dist = list(dist.method = "Earth"), method = "REML")

# tempwarmest + precip.cold.quart
LOIS3meanfit10 <- fitme(formula = mean_source_value ~ 1 + tempwarmest + precip.cold.quart + Matern(1|long + lat) + (1|SiteName),
                        family = gaussian(link = identity), data = LOIS3,
                        resid.model = list(formula= ~ 0 + offset(disp), family = Gamma(link = identity)),
                        prior.weights = n_source_value, control.dist = list(dist.method = "Earth"), method = "REML")

# precip.cold.quart + precip.warm.quart
LOIS3meanfit11 <- fitme(formula = mean_source_value ~ 1 + precip.cold.quart + precip.warm.quart + Matern(1|long + lat) +
(1|SiteName),
                        family = gaussian(link = identity), data = LOIS3,
                        resid.model = list(formula= ~ 0 + offset(disp), family = Gamma(link = identity)),
                        prior.weights = n_source_value, control.dist = list(dist.method = "Earth"), method = "REML")


# tempwarmest + precip.cold.quart +precip.warm.quart
LOIS3meanfit12 <- fitme(formula = mean_source_value ~ 1 + tempwarmest  + precip.cold.quart + precip.warm.quart + Matern(1|long +
lat) + (1|SiteName),
                        family = gaussian(link = identity), data = LOIS3,
                        resid.model = list(formula= ~ 0 + offset(disp), family = Gamma(link = identity)),
                        prior.weights = n_source_value, control.dist = list(dist.method = "Earth"), method = "REML")


AIC(LOIS3meanfit1)
AIC(LOIS3meanfit2)
AIC(LOIS3meanfit3)
AIC(LOIS3meanfit4)
AIC(LOIS3meanfit5)
AIC(LOIS3meanfit6)
AIC(LOIS3meanfit7)
AIC(LOIS3meanfit8)
AIC(LOIS3meanfit9)
AIC(LOIS3meanfit10)
AIC(LOIS3meanfit11)
AIC(LOIS3meanfit12)




    # EARLY OIS 3FIXED EFFECTS MIXED MODELS ####

# MAT
EOIS3meanfit2 <- fitme(formula = mean_source_value ~ 1 + MAT + Matern(1|long + lat) + (1|SiteName),
                       family = gaussian(link = identity), data = EOIS3,
                       resid.model = list(formula= ~ 0 + offset(disp), family = Gamma(link = identity)),
                       prior.weights = n_source_value, control.dist = list(dist.method = "Earth"), method = "REML")
# MAP
EOIS3meanfit3 <- fitme(formula = mean_source_value ~ 1 + MAP + Matern(1|long + lat) + (1|SiteName),
                       family = gaussian(link = identity), data = EOIS3,
                       resid.model = list(formula= ~ 0 + offset(disp), family = Gamma(link = identity)),
                       prior.weights = n_source_value, control.dist = list(dist.method = "Earth"), method = "REML")

# MAT + MAP
EOIS3meanfit4 <- fitme(formula = mean_source_value ~ 1 + MAT + MAP + Matern(1|long + lat) + (1|SiteName),
                       family = gaussian(link = identity), data = EOIS3,
                       resid.model = list(formula= ~ 0 + offset(disp), family = Gamma(link = identity)),
                       prior.weights = n_source_value, control.dist = list(dist.method = "Earth"), method = "REML")

# MAT + MAP + MAT:MAP
EOIS3meanfit5 <- fitme(formula = mean_source_value ~ 1 + MAT + MAP + MAT:MAP+ Matern(1|long + lat) + (1|SiteName),
                       family = gaussian(link = identity), data = EOIS3,
                       resid.model = list(formula= ~ 0 + offset(disp), family = Gamma(link = identity)),
                       prior.weights = n_source_value, control.dist = list(dist.method = "Earth"), method = "REML")

# tempwarmest
EOIS3meanfit6 <- fitme(formula = mean_source_value ~ 1 + tempwarmest + Matern(1|long + lat) + (1|SiteName),
```

```
                       family = gaussian(link = identity), data = EOIS3,
                       resid.model = list(formula= ~ 0 + offset(disp), family = Gamma(link = identity)),
                       prior.weights = n_source_value, control.dist = list(dist.method = "Earth"), method = "REML")

# precipwarmest
EOIS3meanfit7 <- fitme(formula = mean_source_value ~ 1 + precip.warm.quart + Matern(1|long + lat) + (1|SiteName),
                       family = gaussian(link = identity), data = EOIS3,
                       resid.model = list(formula= ~ 0 + offset(disp), family = Gamma(link = identity)),
                       prior.weights = n_source_value, control.dist = list(dist.method = "Earth"), method = "REML")


# precipcoldest
EOIS3meanfit8 <- fitme(formula = mean_source_value ~ 1 + precip.cold.quart + Matern(1|long + lat) + (1|SiteName),
                       family = gaussian(link = identity), data = EOIS3,
                       resid.model = list(formula= ~ 0 + offset(disp), family = Gamma(link = identity)),
                       prior.weights = n_source_value, control.dist = list(dist.method = "Earth"), method = "REML")

# tempwarmest + precip.warm.quart
EOIS3meanfit9 <- fitme(formula = mean_source_value ~ 1 + tempwarmest + precip.warm.quart + Matern(1|long + lat) + (1|SiteName),
                       family = gaussian(link = identity), data = EOIS3,
                       resid.model = list(formula= ~ 0 + offset(disp), family = Gamma(link = identity)),
                       prior.weights = n_source_value, control.dist = list(dist.method = "Earth"), method = "REML")

# tempwarmest + precip.cold.quart
EOIS3meanfit10 <- fitme(formula = mean_source_value ~ 1 + tempwarmest + precip.cold.quart + Matern(1|long + lat) + (1|SiteName),
                        family = gaussian(link = identity), data = EOIS3,
                        resid.model = list(formula= ~ 0 + offset(disp), family = Gamma(link = identity)),
                        prior.weights = n_source_value, control.dist = list(dist.method = "Earth"), method = "REML")

# precip.cold.quart + precip.warm.quart
EOIS3meanfit11 <- fitme(formula = mean_source_value ~ 1 + precip.cold.quart + precip.warm.quart + Matern(1|long + lat) +
(1|SiteName),
                        family = gaussian(link = identity), data = EOIS3,
                        resid.model = list(formula= ~ 0 + offset(disp), family = Gamma(link = identity)),
                        prior.weights = n_source_value, control.dist = list(dist.method = "Earth"), method = "REML")


# tempwarmest + precip.cold.quart +precip.warm.quart
EOIS3meanfit12 <- fitme(formula = mean_source_value ~ 1 + tempwarmest  + precip.cold.quart + precip.warm.quart + Matern(1|long +
lat) + (1|SiteName),
                        family = gaussian(link = identity), data = EOIS3,
                        resid.model = list(formula= ~ 0 + offset(disp), family = Gamma(link = identity)),
                        prior.weights = n_source_value, control.dist = list(dist.method = "Earth"), method = "REML")


AIC(EOIS3meanfit1)
AIC(EOIS3meanfit2)
AIC(EOIS3meanfit3)
AIC(EOIS3meanfit4)
AIC(EOIS3meanfit5)
AIC(EOIS3meanfit6)
AIC(EOIS3meanfit7)
AIC(EOIS3meanfit8)
AIC(EOIS3meanfit9)
AIC(EOIS3meanfit10)
AIC(EOIS3meanfit11)
AIC(EOIS3meanfit12)


    # SET UP ENVIRONMENTAL DATAFRAME FOR PREDICTION ####

# file as raster bricks (one per variable)
MAT <- brick(beyerdata, varname = "BIO1")  # mean annual temperature
tempwarmest <- brick(beyerdata, varname = "BIO10") # mean temp of warmest quarter
MAP <- brick(beyerdata, varname = "BIO12") # mean annual precipitation
precip.warm.quart <- brick(beyerdata, varname = "BIO18") # precipitation of warmest quarter
precip.cold.quart <- brick(beyerdata, varname = "BIO19") # precipitation of coldest quarter

# create a raster stack for each time period of interest
stack.11k <- stack(MAP[[61]], MAT[[61]], precip.warm.quart[[61]], precip.cold.quart[[61]], tempwarmest[[61]])
stack.12k <- stack(MAP[[60]], MAT[[60]], precip.warm.quart[[60]], precip.cold.quart[[60]], tempwarmest[[60]])
stack.14k <- stack(MAP[[58]], MAT[[58]], precip.warm.quart[[58]], precip.cold.quart[[58]], tempwarmest[[58]])
stack.15k <- stack(MAP[[57]], MAT[[57]], precip.warm.quart[[57]], precip.cold.quart[[57]], tempwarmest[[57]])
stack.24k <- stack(MAP[[49]], MAT[[49]], precip.warm.quart[[49]], precip.cold.quart[[49]], tempwarmest[[49]])
stack.36k <- stack(MAP[[43]], MAT[[43]], precip.warm.quart[[43]], precip.cold.quart[[43]], tempwarmest[[43]])
stack.42k <- stack(MAP[[40]], MAT[[40]], precip.warm.quart[[40]], precip.cold.quart[[40]], tempwarmest[[40]])

# crop extent
stack.11k <- crop(stack.11k, extent(-10, 30, 35, 60))
stack.12k <- crop(stack.12k, extent(-10, 30, 35, 60))
stack.14k <- crop(stack.14k, extent(-10, 30, 35, 60))
stack.15k <- crop(stack.15k, extent(-10, 30, 35, 60))
stack.24k <- crop(stack.24k, extent(-10, 30, 35, 60))
stack.36k <- crop(stack.36k, extent(-10, 30, 35, 60))
stack.42k <- crop(stack.42k, extent(-10, 30, 35, 60))

# resample rasters and assign names
    EURDEM <- raster("EURDEM01.tif")
    # EH
    stack.11k <- resample(stack.11k, EURDEM, method="bilinear") # resample
    names(stack.11k) <- c("MAP", "MAT", "precip.warm.quart", "precip.cold.quart",
                          "tempwarmest")  # set names
    # GS1
    stack.12k <- resample(stack.12k, EURDEM, method="bilinear") # resample
    names(stack.12k) <- c("MAP", "MAT", "precip.warm.quart", "precip.cold.quart",
```

```
                    "tempwarmest")  # set names
  # GI1
  stack.14k <- resample(stack.14k, EURDEM, method="bilinear") # resample
  names(stack.14k) <- c("MAP", "MAT", "precip.warm.quart", "precip.cold.quart",
                    "tempwarmest")  # set names
  # GS2
  stack.15k <- resample(stack.15k, EURDEM, method="bilinear") # resample
  names(stack.15k) <- c("MAP", "MAT", "precip.warm.quart", "precip.cold.quart",
                    "tempwarmest")  # set names
  # LGM
  stack.24k <- resample(stack.24k, EURDEM, method="bilinear") # resample
  names(stack.24k) <- c("MAP", "MAT", "precip.warm.quart", "precip.cold.quart",
                    "tempwarmest")  # set names   set names
  # LOIS3
  stack.36k <- resample(stack.36k, EURDEM, method="bilinear") # resample
  names(stack.36k) <- c("MAP", "MAT", "precip.warm.quart", "precip.cold.quart",
                    "tempwarmest")  # set names
  # EOIS3
  stack.42k <- resample(stack.42k, EURDEM, method="bilinear") # resample
  names(stack.42k) <- c("MAP", "MAT", "precip.warm.quart", "precip.cold.quart",
                    "tempwarmest")  # set names

# extract covaraite data at prediction locations #
# scale environmental variables and set predictions locations
  #EH
  EH.env_data_coarse <- aggregate(stack.11k, fact=1)
  EH.env_data_pred <- na.omit(as.data.frame(EH.env_data_coarse, xy = TRUE))
  EH.env_data_pred <- EH.env_data_pred %>%
    mutate_at(c("MAP", "MAT", "precip.warm.quart", "precip.cold.quart", "tempwarmest"),
          ~(scale(.) %>% as.vector))
  EH.pred_locs <- cbind(EH.env_data_pred$x, EH.env_data_pred$y)
  EH.pred_pts <- SpatialPoints(EH.pred_locs)
  EH.env_data_pred$lat <- EH.env_data_pred$y
  EH.env_data_pred$long <- EH.env_data_pred$x
  EH.env_data_pred$SiteName <- EH.env_data_pred %>% group_indices(long, lat)

#GS1
  GS1.env_data_coarse <- aggregate(stack.12k, fact=1)
  GS1.env_data_pred <- na.omit(as.data.frame(GS1.env_data_coarse, xy = TRUE))
  GS1.env_data_pred <- GS1.env_data_pred %>%
   mutate_at(c("MAP", "MAT", "precip.warm.quart", "precip.cold.quart", "tempwarmest"),
          ~(scale(.) %>% as.vector))
  GS1.pred_locs <- cbind(GS1.env_data_pred$x, GS1.env_data_pred$y)
  GS1.pred_pts <- SpatialPoints(GS1.pred_locs)
  GS1.env_data_pred$lat <- GS1.env_data_pred$y
  GS1.env_data_pred$long <- GS1.env_data_pred$x
  GS1.env_data_pred$SiteName <- GS1.env_data_pred %>% group_indices(long, lat)

#GI1
  GI1.env_data_coarse <- aggregate(stack.14k, fact=1)
  GI1.env_data_pred <- na.omit(as.data.frame(GI1.env_data_coarse, xy = TRUE))
  GI1.env_data_pred <- GI1.env_data_pred %>%
    mutate_at(c("MAP", "MAT", "precip.warm.quart", "precip.cold.quart", "tempwarmest"),
          ~(scale(.) %>% as.vector))
  GI1.pred_locs <- cbind(GI1.env_data_pred$x, GI1.env_data_pred$y)
  GI1.pred_pts <- SpatialPoints(GI1.pred_locs)
  GI1.env_data_pred$lat <- GI1.env_data_pred$y
  GI1.env_data_pred$long <- GI1.env_data_pred$x
  GI1.env_data_pred$SiteName <- GI1.env_data_pred %>% group_indices(long, lat)

  #GS2
  GS2.env_data_coarse <- aggregate(stack.15k, fact=1)
  GS2.env_data_pred <- na.omit(as.data.frame(GS2.env_data_coarse, xy = TRUE))
  GS2.env_data_pred <- GS2.env_data_pred %>%
    mutate_at(c("MAP", "MAT", "precip.warm.quart", "precip.cold.quart", "tempwarmest"),
          ~(scale(.) %>% as.vector))
  GS2.pred_locs <- cbind(GS2.env_data_pred$x, GS2.env_data_pred$y)
  GS2.pred_pts <- SpatialPoints(GS2.pred_locs)
  GS2.env_data_pred$lat <- GS2.env_data_pred$y
  GS2.env_data_pred$long <- GS2.env_data_pred$x
  GS2.env_data_pred$SiteName <- GS2.env_data_pred %>% group_indices(long, lat)

  #LGM
  LGM.env_data_coarse <- aggregate(stack.24k, fact=1)
  LGM.env_data_pred <- na.omit(as.data.frame(LGM.env_data_coarse, xy = TRUE))
  LGM.env_data_pred <- LGM.env_data_pred %>%
    mutate_at(c("MAP", "MAT", "precip.warm.quart", "precip.cold.quart", "tempwarmest"),
          ~(scale(.) %>% as.vector))
  LGM.pred_locs <- cbind(LGM.env_data_pred$x, LGM.env_data_pred$y)
  LGM.pred_pts <- SpatialPoints(LGM.pred_locs)
  LGM.env_data_pred$lat <- LGM.env_data_pred$y
  LGM.env_data_pred$long <- LGM.env_data_pred$x
  LGM.env_data_pred$SiteName <- LGM.env_data_pred %>% group_indices(long, lat)

  #LOIS3
  LOIS3.env_data_coarse <- aggregate(stack.36k, fact=1)
  LOIS3.env_data_pred <- na.omit(as.data.frame(LOIS3.env_data_coarse, xy = TRUE))
  LOIS3.env_data_pred <- LOIS3.env_data_pred %>%
    mutate_at(c("MAP", "MAT", "precip.warm.quart", "precip.cold.quart", "tempwarmest"),
          ~(scale(.) %>% as.vector))
  LOIS3.pred_locs <- cbind(LOIS3.env_data_pred$x, LOIS3.env_data_pred$y)
  LOIS3.pred_pts <- SpatialPoints(LOIS3.pred_locs)
  LOIS3.env_data_pred$lat <- LOIS3.env_data_pred$y
  LOIS3.env_data_pred$long <- LOIS3.env_data_pred$x
  LOIS3.env_data_pred$SiteName <- LOIS3.env_data_pred %>% group_indices(long, lat)
```

```
    #EOIS3
    EOIS3.env_data_coarse <- aggregate(stack.42k, fact=1)
    EOIS3.env_data_pred <- na.omit(as.data.frame(EOIS3.env_data_coarse, xy = TRUE))
    EOIS3.env_data_pred <- EOIS3.env_data_pred %>%
      mutate_at(c("MAP", "MAT", "precip.warm.quart", "precip.cold.quart", "tempwarmest"),
            ~(scale(.) %>% as.vector))
    EOIS3.pred_locs <- cbind(EOIS3.env_data_pred$x, EOIS3.env_data_pred$y)
    EOIS3.pred_pts <- SpatialPoints(EOIS3.pred_locs)
    EOIS3.env_data_pred$lat <- EOIS3.env_data_pred$y
    EOIS3.env_data_pred$long <- EOIS3.env_data_pred$x
    EOIS3.env_data_pred$SiteName <- EOIS3.env_data_pred %>% group_indices(long, lat)


    # PREDICT ISOSCAPES WITH BEST PERFORMING MODEL ####

# EH
# Using env_data_pred and dispfit, we predict the residual dispersion variance for each prediction location:
EH.env_data_pred$disp <- predict(EH.dispfit, newdata = EH.env_data_pred)[, 1]
# We can then use the env_data_pred and meanfit to compute the predicted isotope value over all possible observations:
EH.predict.best <- predict(EHmeanfit4, newdata = EH.env_data_pred,
                          variances = list(predVar = TRUE, residVar = TRUE))
# Then extract the point predictions, variance and residual variance:
EH.pred.best <- EH.env_data_pred[, c("lat", "long")]
EH.pred.best$pred <- EH.predict.best[, 1]
EH.pred.best$predVar <- attr(EH.predict.best, "predVar")
EH.pred.best$residVar <- attr(EH.predict.best, "residVar")

# GS1
# Using env_data_pred and dispfit, we predict the residual dispersion variance for each prediction location:
GS1.env_data_pred$disp <- predict(GS1.dispfit, newdata = GS1.env_data_pred)[, 1]
# We can then use the env_data_pred and meanfit to compute the predicted isotope value over all possible observations:
GS1.predict.best <- predict(GS1meanfit6, newdata = GS1.env_data_pred,
                           variances = list(predVar = TRUE, residVar = TRUE))
# Then extract the point predictions, variance and residual variance:
GS1.pred.best <- GS1.env_data_pred[, c("lat", "long")]
GS1.pred.best$pred <- GS1.predict.best[, 1]
GS1.pred.best$predVar <- attr(GS1.predict.best, "predVar")
GS1.pred.best$residVar <- attr(GS1.predict.best, "residVar")

# GI1
# Using env_data_pred and dispfit, we predict the residual dispersion variance for each prediction location:
GI1.env_data_pred$disp <- predict(GI1.dispfit, newdata = GI1.env_data_pred)[, 1]
# We can then use the env_data_pred and meanfit to compute the predicted isotope value over all possible observations:
GI1.predict.best <- predict(GI1meanfit6, newdata = GI1.env_data_pred,
                           variances = list(predVar = TRUE, residVar = TRUE))
# Then extract the point predictions, variance and residual variance:
GI1.pred.best <- GI1.env_data_pred[, c("lat", "long")]
GI1.pred.best$pred <- GI1.predict.best[, 1]
GI1.pred.best$predVar <- attr(GI1.predict.best, "predVar")
GI1.pred.best$residVar <- attr(GI1.predict.best, "residVar")

# GS2
# Using env_data_pred and dispfit, we predict the residual dispersion variance for each prediction location:
GS2.env_data_pred$disp <- predict(GS2.dispfit, newdata = GS2.env_data_pred)[, 1]
# We can then use the env_data_pred and meanfit to compute the predicted isotope value over all possible observations:
GS2.predict.best <- predict(GS2meanfit2, newdata = GS2.env_data_pred,
                           variances = list(predVar = TRUE, residVar = TRUE))
# Then extract the point predictions, variance and residual variance:
GS2.pred.best <- GS2.env_data_pred[, c("lat", "long")]
GS2.pred.best$pred <- GS2.predict.best[, 1]
GS2.pred.best$predVar <- attr(GS2.predict.best, "predVar")
GS2.pred.best$residVar <- attr(GS2.predict.best, "residVar")

# LGM
# Using env_data_pred and dispfit, we predict the residual dispersion variance for each prediction location:
LGM.env_data_pred$disp <- predict(LGM.dispfit, newdata = LGM.env_data_pred)[, 1]
# We can then use the env_data_pred and meanfit to compute the predicted isotope value over all possible observations:
LGM.predict.best <- predict(LGMmeanfit11, newdata = LGM.env_data_pred,
                           variances = list(predVar = TRUE, residVar = TRUE))
# Then extract the point predictions, variance and residual variance:
LGM.pred.best <- LGM.env_data_pred[, c("lat", "long")]
LGM.pred.best$pred <- LGM.predict.best[, 1]
LGM.pred.best$predVar <- attr(LGM.predict.best, "predVar")
LGM.pred.best$residVar <- attr(LGM.predict.best, "residVar")

# LOIS3
# Using env_data_pred and dispfit, we predict the residual dispersion variance for each prediction location:
LOIS3.env_data_pred$disp <- predict(LOIS3.dispfit, newdata = LOIS3.env_data_pred)[, 1]
# We can then use the env_data_pred and meanfit to compute the predicted isotope value over all possible observations:
LOIS3.predict.best <- predict(LOIS3meanfit3, newdata = LOIS3.env_data_pred,
                             variances = list(predVar = TRUE, residVar = TRUE))
# Then extract the point predictions, variance and residual variance:
LOIS3.pred.best <- LOIS3.env_data_pred[, c("lat", "long")]
LOIS3.pred.best$pred <- LOIS3.predict.best[, 1]
LOIS3.pred.best$predVar <- attr(LOIS3.predict.best, "predVar")
LOIS3.pred.best$residVar <- attr(LOIS3.predict.best, "residVar")

# EOIS3
# Using env_data_pred and dispfit, we predict the residual dispersion variance for each prediction location:
EOIS3.env_data_pred$disp <- predict(EOIS3.dispfit, newdata = EOIS3.env_data_pred)[, 1]
# We can then use the env_data_pred and meanfit to compute the predicted isotope value over all possible observations:
EOIS3.predict.best <- predict(EOIS3meanfit2, newdata = EOIS3.env_data_pred,
                             variances = list(predVar = TRUE, residVar = TRUE))
# Then extract the point predictions, variance and residual variance:
```

```
EOIS3.pred.best <- EOIS3.env_data_pred[, c("lat", "long")]
EOIS3.pred.best$pred <- EOIS3.predict.best[, 1]
EOIS3.pred.best$predVar <- attr(EOIS3.predict.best, "predVar")
EOIS3.pred.best$residVar <- attr(EOIS3.predict.best, "residVar")


    # SET UP PREDICTION SURFACES ####

# EH
# tranform prediction surfaces into raster
EH.pred.best <- EH.pred.best[,c('long', 'lat', 'pred', 'predVar', 'residVar')]
EH.pred.rast.best  <- rasterFromXYZ(EH.pred.best)  #Convert first two columns as lon-lat and third as value
# crop prediction values to interpolation area
EH.cr.best  <- crop(EH.pred.rast.best, extent(EH.interpol.area), snap="out")
EH.fr.best  <- rasterize(EH.interpol.area, EH.cr.best)
EH.lr.best  <- raster::mask(x=EH.cr.best, mask=EH.fr.best)
# then convert raster to dataframe:
EH.pred.rast.df.best   <- as.data.frame(EH.lr.best , xy=TRUE)
# remove data with NAs
EH.pred.rast.df.best  <- na.omit(EH.pred.rast.df.best)

# GS1
# tranform prediction surfaces into raster
GS1.pred.best <- GS1.pred.best[,c('long', 'lat', 'pred', 'predVar', 'residVar')]
GS1.pred.rast.best  <- rasterFromXYZ(GS1.pred.best )  #Convert first two columns as lon-lat and third as value
# crop prediction values to interpolation area
GS1.cr.best  <- crop(GS1.pred.rast.best, extent(GS1.interpol.area), snap="out")
GS1.fr.best  <- rasterize(GS1.interpol.area, GS1.cr.best)
GS1.lr.best  <- raster::mask(x=GS1.cr.best, mask=GS1.fr.best)
# then convert raster to dataframe:
GS1.pred.rast.df.best   <- as.data.frame(GS1.lr.best , xy=TRUE)
# remove data with NAs
GS1.pred.rast.df.best  <- na.omit(GS1.pred.rast.df.best)

# GI1
# tranform prediction surfaces into raster
GI1.pred.best <- GI1.pred.best[,c('long', 'lat', 'pred', 'predVar', 'residVar')]
GI1.pred.rast.best  <- rasterFromXYZ(GI1.pred.best )  #Convert first two columns as lon-lat and third as value
# crop prediction values to interpolation area
GI1.cr.best  <- crop(GI1.pred.rast.best, extent(GI1.interpol.area), snap="out")
GI1.fr.best  <- rasterize(GI1.interpol.area, GI1.cr.best)
GI1.lr.best  <- raster::mask(x=GI1.cr.best, mask=GI1.fr.best)
# then convert raster to dataframe:
GI1.pred.rast.df.best   <- as.data.frame(GI1.lr.best , xy=TRUE)
# remove data with NAs
GI1.pred.rast.df.best  <- na.omit(GI1.pred.rast.df.best)

# GS2
# tranform prediction surfaces into raster
GS2.pred.best <- GS2.pred.best[,c('long', 'lat', 'pred', 'predVar', 'residVar')]
GS2.pred.rast.best  <- rasterFromXYZ(GS2.pred.best )  #Convert first two columns as lon-lat and third as value
# crop prediction values to interpolation area
GS2.cr.best  <- crop(GS2.pred.rast.best, extent(GS2.interpol.area), snap="out")
GS2.fr.best  <- rasterize(GS2.interpol.area, GS2.cr.best)
GS2.lr.best  <- raster::mask(x=GS2.cr.best, mask=GS2.fr.best)
# then convert raster to dataframe:
GS2.pred.rast.df.best   <- as.data.frame(GS2.lr.best , xy=TRUE)
# remove data with NAs
GS2.pred.rast.df.best  <- na.omit(GS2.pred.rast.df.best)

# LGM
# tranform prediction surfaces into raster
LGM.pred.best <- LGM.pred.best[,c('long', 'lat', 'pred', 'predVar', 'residVar')]
LGM.pred.rast.best  <- rasterFromXYZ(LGM.pred.best )  #Convert first two columns as lon-lat and third as value
# crop prediction values to interpolation area
LGM.cr.best  <- crop(LGM.pred.rast.best, extent(LGM.interpol.area), snap="out")
LGM.fr.best  <- rasterize(LGM.interpol.area, LGM.cr.best)
LGM.lr.best  <- raster::mask(x=LGM.cr.best, mask=LGM.fr.best)
# then convert raster to dataframe:
LGM.pred.rast.df.best   <- as.data.frame(LGM.lr.best , xy=TRUE)
# remove data with NAs
LGM.pred.rast.df.best  <- na.omit(LGM.pred.rast.df.best)

# LOIS3
# tranform prediction surfaces into raster
LOIS3.pred.best <- LOIS3.pred.best[,c('long', 'lat', 'pred', 'predVar', 'residVar')]
LOIS3.pred.rast.best  <- rasterFromXYZ(LOIS3.pred.best )  #Convert first two columns as lon-lat and third as value
# crop prediction values to interpolation area
LOIS3.cr.best  <- crop(LOIS3.pred.rast.best, extent(LOIS3.interpol.area), snap="out")
LOIS3.fr.best  <- rasterize(LOIS3.interpol.area, LOIS3.cr.best)
LOIS3.lr.best  <- raster::mask(x=LOIS3.cr.best, mask=LOIS3.fr.best)
# then convert raster to dataframe:
LOIS3.pred.rast.df.best   <- as.data.frame(LOIS3.lr.best , xy=TRUE)
# remove data with NAs
LOIS3.pred.rast.df.best  <- na.omit(LOIS3.pred.rast.df.best)

# EOIS3
# tranform prediction surfaces into raster
EOIS3.pred.best <- EOIS3.pred.best[,c('long', 'lat', 'pred', 'predVar', 'residVar')]
EOIS3.pred.rast.best  <- rasterFromXYZ(EOIS3.pred.best )  #Convert first two columns as lon-lat and third as value
# crop prediction values to interpolation area
EOIS3.cr.best  <- crop(EOIS3.pred.rast.best, extent(EOIS3.interpol.area), snap="out")
EOIS3.fr.best  <- rasterize(EOIS3.interpol.area, EOIS3.cr.best)
EOIS3.lr.best  <- raster::mask(x=EOIS3.cr.best, mask=EOIS3.fr.best)
# then convert raster to dataframe:
```

```
EOIS3.pred.rast.df.best   <- as.data.frame(EOIS3.lr.best , xy=TRUE)
# remove data with NAs
EOIS3.pred.rast.df.best  <- na.omit(EOIS3.pred.rast.df.best)




    # PLOT ISOSCAPES (FIGURE 6 and 7) ####

# EH prediction surface
(EHmappred.best  <- ggplot()+
  geom_polygon(data=pcoastEH.simp, aes(long, lat, group=group), fill="lightgrey") +
  geom_tile(data = EH.pred.rast.df.best, mapping=aes(x=x, y=y, fill=pred)) +
  scale_fill_stepsn(colours = col, limits = c(0,6), breaks=c(0.5,1,1.5,2,2.5,3,3.5,4,4.5,5,5.5)) +
  geom_path(data=eurmed, aes(long, lat, group=group), colour="black") +
  geom_polygon(data=iceEH, aes(long, lat, group=group), colour="black", fill="white") +
  geom_point(data = EH, aes(long, lat), colour="black", size=0.5) +
  coord_cartesian(xlim = c(-10,25), ylim = c(40, 60)) +
  xlab("Longitude") +
  ylab("Latitude") +
  labs(title="Early Holocene",
       fill=expression(paste(delta^{15}, "N (\u2030)"))) +
  theme_bw() +
  theme(legend.position="right"))


# EH prediciton variance
(EHmapvar.best <- ggplot()+
  geom_polygon(data=pcoastEH.simp, aes(long, lat, group=group), fill="lightgrey") +
  geom_tile(data = EH.pred.rast.df.best, mapping=aes(x=x, y=y, fill=predVar)) +
  scale_fill_stepsn(colours = col.var, breaks=c(0.5,1,1.5), limits = c(0, 3))+
  geom_path(data=eurmed, aes(long, lat, group=group), colour="black") +
  geom_polygon(data=iceEH, aes(long, lat, group=group), colour="black", fill="white") +
  geom_point(data = EH, aes(long, lat), colour="black", size=0.5) +
  coord_cartesian(xlim = c(-10,25), ylim = c(40, 60)) +
  xlab("Longitude") +
  ylab("Latitude") +
  labs(title="Early Holocene",
       fill=expression(paste(delta^{15}, "N (\u2030)"))) +
  theme(legend.position="right") +
  theme_bw())

# GS1 prediciton surface
(GS1mappred.best <- ggplot()+
  geom_polygon(data=pcoastGS1.simp, aes(long, lat, group=group), fill="lightgrey") +
  geom_tile(data = GS1.pred.rast.df.best, mapping=aes(x=x, y=y, fill=pred)) +
  scale_fill_stepsn(colours = col, limits = c(0,6), breaks=c(0.5,1,1.5,2,2.5,3,3.5,4,4.5,5,5.5)) +
  geom_path(data=eurmed, aes(long, lat, group=group), colour="black") +
  geom_polygon(data=iceGS1, aes(long, lat, group=group), colour="black", fill="white") +
  geom_point(data = GS1, aes(long, lat), colour="black", size=0.5) +
  coord_cartesian(xlim = c(-10,25), ylim = c(40, 60)) +
  xlab("Longitude") +
  ylab("Latitude") +
  labs(title="Younger Dryas",
       fill=expression(paste(delta^{15}, "N (\u2030)"))) +
  theme_bw() +
  theme(legend.position="none"))


# GS1 prediciton variance
(GS1mapvar.best <- ggplot()+
  geom_polygon(data=pcoastGS1.simp, aes(long, lat, group=group), fill="lightgrey") +
  geom_tile(data = GS1.pred.rast.df.best, mapping=aes(x=x, y=y, fill=predVar)) +
  scale_fill_stepsn(colours = col.var, breaks=c(0.5,1,1.5), limits = c(0, 3))+
  geom_path(data=eurmed, aes(long, lat, group=group), colour="black") +
  geom_polygon(data=iceGS1, aes(long, lat, group=group), colour="black", fill="white") +
  geom_point(data = GS1, aes(long, lat), colour="black", size=0.5) +
  coord_cartesian(xlim = c(-10,25), ylim = c(40, 60)) +
  xlab("Longitude") +
  ylab("Latitude") +
  labs(title="Younger Dryas",
       fill=expression(paste(delta^{15}, "N (\u2030)"))) +
  theme_bw() +
  theme(legend.position="none"))


# GI1 prediciton surface
(GI1mappred.best <- ggplot()+
  geom_polygon(data=pcoastGI1.simp, aes(long, lat, group=group), fill="lightgrey") +
  geom_tile(data = GI1.pred.rast.df.best, mapping=aes(x=x, y=y, fill=pred)) +
  scale_fill_stepsn(colours = col, limits = c(0,6), breaks=c(0.5,1,1.5,2,2.5,3,3.5,4,4.5,5,5.5)) +
  geom_path(data=eurmed, aes(long, lat, group=group), colour="black") +
  geom_polygon(data=iceGI1, aes(long, lat, group=group), colour="black", fill="white") +
  geom_point(data = GI1, aes(long, lat), colour="black", size=0.5) +
  coord_cartesian(xlim = c(-10,25), ylim = c(40, 60)) +
  xlab("Longitude") +
  ylab("Latitude") +
  labs(title="Late Glacial Interstadial",
       fill=expression(paste(delta^{15}, "N (\u2030)"))) +
  theme_bw() +
  theme(legend.position="none"))

# GI1 prediciton variance
(GI1mapvar.best <- ggplot()+
  geom_polygon(data=pcoastGI1.simp, aes(long, lat, group=group), fill="lightgrey") +
```

```
  geom_tile(data = GI1.pred.rast.df.best, mapping=aes(x=x, y=y, fill=predVar)) +
  scale_fill_stepsn(colours = col.var, breaks=c(0.5,1,1.5), limits = c(0, 3))+
  geom_path(data=eurmed, aes(long, lat, group=group), colour="black") +
  geom_polygon(data=iceGI1, aes(long, lat, group=group), colour="black", fill="white") +
  geom_point(data = GI1, aes(long, lat), colour="black", size=0.5) +
  coord_cartesian(xlim = c(-10,25), ylim = c(40, 60)) +
  xlab("Longitude") +
  ylab("Latitude") +
  labs(title="Late Glacial Interstadial",
       fill=expression(paste(delta^{15}, "N (\u2030)"))) +
  theme_bw() +
  theme(legend.position="none"))

# GS2 prediciton surface
(GS2mappred.best <- ggplot()+
  geom_polygon(data=pcoastGS2.simp, aes(long, lat, group=group), fill="lightgrey") +
  geom_tile(data = GS2.pred.rast.df.best, mapping=aes(x=x, y=y, fill=pred)) +
  scale_fill_stepsn(colours = col, limits = c(0,6), breaks=c(0.5,1,1.5,2,2.5,3,3.5,4,4.5,5,5.5)) +
  geom_path(data=eurmed, aes(long, lat, group=group), colour="black") +
  geom_polygon(data=iceGS2, aes(long, lat, group=group), colour="black", fill="white") +
  geom_point(data = GS2, aes(long, lat), colour="black", size=0.5) +
  coord_cartesian(xlim = c(-10,25), ylim = c(40, 60)) +
  xlab("Longitude") +
  ylab("Latitude") +
  labs(title="Last Glacial Termination",
       fill=expression(paste(delta^{15}, "N (\u2030)"))) +
  theme_bw() +
  theme(legend.position="none"))

# GS2 prediciton variance
(GS2mapvar.best <- ggplot()+
  geom_polygon(data=pcoastGS2.simp, aes(long, lat, group=group), fill="lightgrey") +
  geom_tile(data = GS2.pred.rast.df.best, mapping=aes(x=x, y=y, fill=predVar)) +
  scale_fill_stepsn(colours = col.var, breaks=c(0.5,1,1.5), limits = c(0, 3))+
  geom_path(data=eurmed, aes(long, lat, group=group), colour="black") +
  geom_polygon(data=iceGS2, aes(long, lat, group=group), colour="black", fill="white") +
  geom_point(data = GS2, aes(long, lat), colour="black", size=0.5) +
  coord_cartesian(xlim = c(-10,25), ylim = c(40, 60)) +
  xlab("Longitude") +
  ylab("Latitude") +
  labs(title="Last Glacial Termination",
       fill=expression(paste(delta^{15}, "N (\u2030)"))) +
  theme_bw() +
  theme(legend.position="none"))

# LGM prediciton surface
(LGMmappred.best <- ggplot()+
  geom_polygon(data=pcoastLGM.simp, aes(long, lat, group=group), fill="lightgrey") +
  geom_tile(data = LGM.pred.rast.df.best, mapping=aes(x=x, y=y, fill=pred)) +
  scale_fill_stepsn(colours = col, limits = c(0,6), breaks=c(0.5,1,1.5,2,2.5,3,3.5,4,4.5,5,5.5)) +
  geom_path(data=eurmed, aes(long, lat, group=group), colour="black") +
  geom_polygon(data=iceLGM, aes(long, lat, group=group), colour="black", fill="white") +
  geom_point(data = LGM, aes(long, lat), colour="black", size=0.5) +
  coord_cartesian(xlim = c(-10,25), ylim = c(40, 60)) +
  xlab("Longitude") +
  ylab("Latitude") +
  labs(title="Last Glacial Maximum",
       fill=expression(paste(delta^{15}, "N (\u2030)"))) +
  theme_bw() +
  theme(legend.position="none"))

# LGM prediciton variance
(LGMmapvar.best <- ggplot()+
  geom_polygon(data=pcoastLGM.simp, aes(long, lat, group=group), fill="lightgrey") +
  geom_tile(data = LGM.pred.rast.df.best, mapping=aes(x=x, y=y, fill=predVar)) +
  scale_fill_stepsn(colours = col.var, breaks=c(0.5,1,1.5), limits = c(0, 3))+
  geom_path(data=eurmed, aes(long, lat, group=group), colour="black") +
  geom_polygon(data=iceLGM, aes(long, lat, group=group), colour="black", fill="white") +
  geom_point(data = LGM, aes(long, lat), colour="black", size=0.5) +
  coord_cartesian(xlim = c(-10,25), ylim = c(40, 60)) +
  xlab("Longitude") +
  ylab("Latitude") +
  labs(title="Last Glacial Maximum",
       fill=expression(paste(delta^{15}, "N (\u2030)"))) +
  theme_bw() +
  theme(legend.position="none"))

# LOIS3 prediciton surface
(LOIS3mappred.best <- ggplot()+
  geom_polygon(data=pcoastLOIS3.simp, aes(long, lat, group=group), fill="lightgrey") +
  geom_tile(data = LOIS3.pred.rast.df.best, mapping=aes(x=x, y=y, fill=pred)) +
  scale_fill_stepsn(colours = col, limits = c(0,6), breaks=c(0.5,1,1.5,2,2.5,3,3.5,4,4.5,5,5.5)) +
  geom_path(data=eurmed, aes(long, lat, group=group), colour="black") +
  geom_polygon(data=iceLOIS3, aes(long, lat, group=group), colour="black", fill="white") +
  geom_point(data = LOIS3, aes(long, lat), colour="black", size=0.5) +
  coord_cartesian(xlim = c(-10,25), ylim = c(40, 60)) +
  xlab("Longitude") +
  ylab("Latitude") +
  labs(title="Late OIS 3",
       fill=expression(paste(delta^{15}, "N (\u2030)"))) +
  theme_bw() +
  theme(legend.position="none"))

# LOIS3 prediciton variance
(LOIS3mapvar.best <- ggplot()+
```

```
  geom_polygon(data=pcoastLOIS3.simp, aes(long, lat, group=group), fill="lightgrey") +
  geom_tile(data = LOIS3.pred.rast.df.best, mapping=aes(x=x, y=y, fill=predVar)) +
  scale_fill_stepsn(colours = col.var, breaks=c(0.5,1,1.5), limits = c(0, 3))+
  geom_path(data=eurmed, aes(long, lat, group=group), colour="black") +
  geom_polygon(data=iceLOIS3, aes(long, lat, group=group), colour="black", fill="white") +
  geom_point(data = LOIS3, aes(long, lat), colour="black", size=0.5) +
  coord_cartesian(xlim = c(-10,25), ylim = c(40, 60)) +
  xlab("Longitude") +
  ylab("Latitude") +
  labs(title="Late OIS 3",
       fill=expression(paste(delta^{15}, "N (\u2030)"))) +
  theme_bw() +
  theme(legend.position="none"))

# EOIS3 prediciton surface
(EOIS3mappred.best <- ggplot()+
  geom_polygon(data=pcoastEOIS3.simp, aes(long, lat, group=group), fill="lightgrey") +
  geom_tile(data = EOIS3.pred.rast.df.best, mapping=aes(x=x, y=y, fill=pred)) +
  scale_fill_stepsn(colours = col, limits = c(0,6), breaks=c(0.5,1,1.5,2,2.5,3,3.5,4,4.5,5,5.5)) +
  geom_path(data=eurmed, aes(long, lat, group=group), colour="black") +
  geom_polygon(data=iceEOIS3, aes(long, lat, group=group), colour="black", fill="white") +
  geom_point(data = EOIS3, aes(long, lat), colour="black", size=0.5) +
  coord_cartesian(xlim = c(-10,25), ylim = c(40, 60)) +
  xlab("Longitude") +
  ylab("Latitude") +
  labs(title="Early OIS 3",
       fill=expression(paste(delta^{15}, "N (\u2030)"))) +
  theme_bw() +
  theme(legend.position="none"))

# EOIS3 prediciton variance
(EOIS3mapvar.best <- ggplot()+
  geom_polygon(data=pcoastEOIS3.simp, aes(long, lat, group=group), fill="lightgrey") +
  geom_tile(data = EOIS3.pred.rast.df.best, mapping=aes(x=x, y=y, fill=predVar)) +
  scale_fill_stepsn(colours = col.var, breaks=c(0.5,1,1.5), limits = c(0, 3))+
  geom_path(data=eurmed, aes(long, lat, group=group), colour="black") +
  geom_polygon(data=iceEOIS3, aes(long, lat, group=group), colour="black", fill="white") +
  geom_point(data = EOIS3, aes(long, lat), colour="black", size=0.5) +
  coord_cartesian(xlim = c(-10,25), ylim = c(40, 60)) +
  xlab("Longitude") +
  ylab("Latitude") +
  labs(title="Early OIS 3",
       fill=expression(paste(delta^{15}, "N (\u2030)"))) +
  theme_bw() +
  theme(legend.position="none"))

# first get legends
legendPred <- get_legend(EHmappred.best)
# remove legend from EH ploy
EHmappred.best <- EHmappred.best + theme(legend.position="none")
# combine plots
predplots.best <- ggpubr::ggarrange(EHmappred.best,
                                    GS1mappred.best,
                                    GI1mappred.best,
                                    GS2mappred.best,
                                    LGMmappred.best,
                                    LOIS3mappred.best,
                                    EOIS3mappred.best,
                                    legendPred, ncol=2, nrow=4)

ggsave(file="Figure_6.pdf", predplots.best, width = 20, height = 30, units = c("cm"), dpi = 300)

# variance surfaces #

# first get legends
legendvar <- get_legend(EHmapvar.best)

EHmapvar.best <- EHmapvar.best + theme(legend.position="none")

varplots.best <- ggpubr::ggarrange(EHmapvar.best,
                                   GS1mapvar.best,
                                   GI1mapvar.best,
                                   GS2mapvar.best,
                                   LGMmapvar.best,
                                   LOIS3mapvar.best,
                                   EOIS3mapvar.best,
                                   legendvar, ncol=2, nrow=4)

ggsave(file="figure_7.pdf", varplots.best, width = 20, height = 30, units = c("cm"), dpi = 300)



    # COMPARE PREDICTED TO MODELLED VALUES (FIGURE 8) ####

# turn each dataframe into an spdf
d15N.EH.aggre.spdf <- EH
coordinates(d15N.EH.aggre.spdf) <- ~long+lat
proj4string(d15N.EH.aggre.spdf) <- latlon
d15N.GS1.aggre.spdf <- GS1
coordinates(d15N.GS1.aggre.spdf) <- ~long+lat
proj4string(d15N.GS1.aggre.spdf) <- latlon
d15N.GI1.aggre.spdf <- GI1
coordinates(d15N.GI1.aggre.spdf) <- ~long+lat
proj4string(d15N.GI1.aggre.spdf) <- latlon
```

```
d15N.GS2.aggre.spdf <- GS2
coordinates(d15N.GS2.aggre.spdf) <- ~long+lat
proj4string(d15N.GS2.aggre.spdf) <- latlon
d15N.LGM.aggre.spdf <- LGM
coordinates(d15N.LGM.aggre.spdf) <- ~long+lat
proj4string(d15N.LGM.aggre.spdf) <- latlon
d15N.LOIS3.aggre.spdf <- LOIS3
coordinates(d15N.LOIS3.aggre.spdf) <- ~long+lat
proj4string(d15N.LOIS3.aggre.spdf) <- latlon
d15N.EOIS3.aggre.spdf <- EOIS3
coordinates(d15N.EOIS3.aggre.spdf) <- ~long+lat
proj4string(d15N.EOIS3.aggre.spdf) <- latlon

#overlay rasters and extract predicted values
EH_predicted_data <- raster::extract(EH.lr.best, d15N.EH.aggre.spdf) # extract data
d15N.EH.aggre.spdf <- cbind(d15N.EH.aggre.spdf, EH_predicted_data) # merge to the main data
colnames(d15N.EH.aggre.spdf@data)
# convert back to df and remove NAs
d15N.EH.aggre.df <- as.data.frame(d15N.EH.aggre.spdf)
d15N.EH.aggre.df <- d15N.EH.aggre.df %>% drop_na(pred)

# Calculate RMSE
rmse.EH <- round(modelRMSE(o = d15N.EH.aggre.df$mean_source_value, p = d15N.EH.aggre.df$pred), 2)
rmse.EH <- as.character(rmse.EH)
rmse.EH

#overlay rasters and extract predicted values
GS1_predicted_data <- raster::extract(GS1.lr.best, d15N.GS1.aggre.spdf) # extract data
d15N.GS1.aggre.spdf <- cbind(d15N.GS1.aggre.spdf, GS1_predicted_data) # merge to the main data
colnames(d15N.GS1.aggre.spdf@data)
# convert back to df and remove NAs
d15N.GS1.aggre.df <- as.data.frame(d15N.GS1.aggre.spdf)
d15N.GS1.aggre.df <- d15N.GS1.aggre.df %>% drop_na(pred)

# Calculate RMSE
rmse.GS1 <- round(modelRMSE(o = d15N.GS1.aggre.df$mean_source_value, p = d15N.GS1.aggre.df$pred), 2)
rmse.GS1 <- as.character(rmse.GS1)
rmse.GS1

#overlay rasters and extract predicted values
GI1_predicted_data <- raster::extract(GI1.lr.best, d15N.GI1.aggre.spdf) # extract data
d15N.GI1.aggre.spdf <- cbind(d15N.GI1.aggre.spdf, GI1_predicted_data) # merge to the main data
colnames(d15N.GI1.aggre.spdf@data)
# convert back to df and remove NAs
d15N.GI1.aggre.df <- as.data.frame(d15N.GI1.aggre.spdf)
d15N.GI1.aggre.df <- d15N.GI1.aggre.df %>% drop_na(pred)

# Calculate RMSE
rmse.GI1 <- round(modelRMSE(o = d15N.GI1.aggre.df$mean_source_value, p = d15N.GI1.aggre.df$pred), 2)
rmse.GI1 <- as.character(rmse.GI1)
rmse.GI1

#overlay rasters and extract predicted values
GS2_predicted_data <- raster::extract(GS2.lr.best, d15N.GS2.aggre.spdf) # extract data
d15N.GS2.aggre.spdf <- cbind(d15N.GS2.aggre.spdf, GS2_predicted_data) # merge to the main data
colnames(d15N.GS2.aggre.spdf@data)
# convert back to df and remove NAs
d15N.GS2.aggre.df <- as.data.frame(d15N.GS2.aggre.spdf)
d15N.GS2.aggre.df <- d15N.GS2.aggre.df %>% drop_na(pred)

# Calculate RMSE
rmse.GS2 <- round(modelRMSE(o = d15N.GS2.aggre.df$mean_source_value, p = d15N.GS2.aggre.df$pred), 2)
rmse.GS2 <- as.character(rmse.GS2)
rmse.GS2

#overlay rasters and extract predicted values
LGM_predicted_data <- raster::extract(LGM.lr.best, d15N.LGM.aggre.spdf) # extract data
d15N.LGM.aggre.spdf <- cbind(d15N.LGM.aggre.spdf, LGM_predicted_data) # merge to the main data
colnames(d15N.LGM.aggre.spdf@data)
# convert back to df and remove NAs
d15N.LGM.aggre.df <- as.data.frame(d15N.LGM.aggre.spdf)
d15N.LGM.aggre.df <- d15N.LGM.aggre.df %>% drop_na(pred)

# Calculate RMSE
rmse.LGM <- round(modelRMSE(o = d15N.LGM.aggre.df$mean_source_value, p = d15N.LGM.aggre.df$pred), 2)
rmse.LGM <- as.character(rmse.LGM)
rmse.LGM

#overlay rasters and extract predicted values
LOIS3_predicted_data <- raster::extract(LOIS3.lr.best, d15N.LOIS3.aggre.spdf) # extract data
d15N.LOIS3.aggre.spdf <- cbind(d15N.LOIS3.aggre.spdf, LOIS3_predicted_data) # merge to the main data
colnames(d15N.LOIS3.aggre.spdf@data)
# convert back to df and remove NAs
d15N.LOIS3.aggre.df <- as.data.frame(d15N.LOIS3.aggre.spdf)
d15N.LOIS3.aggre.df <- d15N.LOIS3.aggre.df %>% drop_na(pred)

# Calculate RMSE
rmse.LOIS3 <- round(modelRMSE(o = d15N.LOIS3.aggre.df$mean_source_value, p = d15N.LOIS3.aggre.df$pred), 2)
rmse.LOIS3 <- as.character(rmse.LOIS3)
rmse.LOIS3


#overlay rasters and extract predicted values
EOIS3_predicted_data <- raster::extract(EOIS3.lr.best, d15N.EOIS3.aggre.spdf) # extract data
d15N.EOIS3.aggre.spdf <- cbind(d15N.EOIS3.aggre.spdf, EOIS3_predicted_data) # merge to the main data
```

```
colnames(d15N.EOIS3.aggre.spdf@data)
# convert back to df and remove NAs
d15N.EOIS3.aggre.df <- as.data.frame(d15N.EOIS3.aggre.spdf)
d15N.EOIS3.aggre.df <- d15N.EOIS3.aggre.df %>% drop_na(pred)

# Calculate RMSE
rmse.EOIS3 <- round(modelRMSE(o = d15N.EOIS3.aggre.df$mean_source_value, p = d15N.EOIS3.aggre.df$pred), 2)
rmse.EOIS3 <- as.character(rmse.EOIS3)
rmse.EOIS3



(EH.pred.ob.best <- ggplot(data = d15N.EH.aggre.df, mapping=aes(x=pred, y=mean_source_value))+
  geom_point() +
  geom_abline(slope=1, intercept=0)+
  geom_smooth(method = "lm", se=TRUE, color="red", formula = y ~ x) +
  stat_poly_eq(aes(label = paste(..rr.label.., sep = "~~~")),
               label.x.npc = "right", label.y.npc = 0.15,
               formula = y ~ x, parse = TRUE, size = 4) +
  stat_poly_eq(aes(label = paste(..p.value.label.., sep = "~~~")),
               label.x.npc = "right", label.y.npc = "bottom",
               formula = y ~ x, parse = TRUE, size = 4) +
  annotate("text", x=10.5, y=3, label = paste("RMSE = ", rmse.EH))+
  ggtitle("Early Holocene")+
  labs(x = expression(paste(delta^{15}, "N predicted (\u2030)")),
       y = expression(paste(delta^{15}, "N observed (\u2030)"))) +
  scale_y_continuous(limits = c(0,12), breaks = seq(0, 12, by = 2)) +
  scale_x_continuous(limits = c(0,12),breaks = seq(0, 12, by = 2)) +
  theme_bw())

(GS1.pred.ob.best <- ggplot(data = d15N.GS1.aggre.df, mapping=aes(x=pred, y=mean_source_value))+
  geom_point() +
  geom_abline(slope=1, intercept=0)+
  geom_smooth(method = "lm", se=TRUE, color="red", formula = y ~ x) +
  stat_poly_eq(aes(label = paste(..rr.label.., sep = "~~~")),
               label.x.npc = "right", label.y.npc = 0.15,
               formula = y ~ x, parse = TRUE, size = 4) +
  stat_poly_eq(aes(label = paste(..p.value.label.., sep = "~~~")),
               label.x.npc = "right", label.y.npc = "bottom",
               formula = y ~ x, parse = TRUE, size = 4) +
    annotate("text", x=10.5, y=3, label = paste("RMSE = ", rmse.GS1))+
  ggtitle("Younger Dryas")+
  labs(x = expression(paste(delta^{15}, "N predicted (\u2030)")),
       y = expression(paste(delta^{15}, "N observed (\u2030)"))) +
    scale_y_continuous(limits = c(0,12), breaks = seq(0, 12, by = 2)) +
    scale_x_continuous(limits = c(0,12),breaks = seq(0, 12, by = 2)) +
  theme_bw())



(GI1.pred.ob.best <- ggplot(data = d15N.GI1.aggre.df, mapping=aes(x=pred, y=mean_source_value))+
  geom_point() +
  geom_abline(slope=1, intercept=0)+
  geom_smooth(method = "lm", se=TRUE, color="red", formula = y ~ x) +
  stat_poly_eq(aes(label = paste(..rr.label.., sep = "~~~")),
               label.x.npc = "right", label.y.npc = 0.15,
               formula = y ~ x, parse = TRUE, size = 4) +
  stat_poly_eq(aes(label = paste(..p.value.label.., sep = "~~~")),
               label.x.npc = "right", label.y.npc = "bottom",
               formula = y ~ x, parse = TRUE, size = 4) +
  annotate("text", x=10.5, y=3, label = paste("RMSE = ", rmse.GI1))+
  ggtitle("Late Glacial Interstadial")+
  labs(x = expression(paste(delta^{15}, "N predicted (\u2030)")),
       y = expression(paste(delta^{15}, "N observed (\u2030)"))) +
  scale_y_continuous(limits = c(0,12), breaks = seq(0, 12, by = 2)) +
  scale_x_continuous(limits = c(0,12),breaks = seq(0, 12, by = 2)) +
  theme_bw())


# plot predicted versus observed values
(GS2.pred.ob.best <- ggplot(data = d15N.GS2.aggre.df, mapping=aes(x=pred, y=mean_source_value))+
  geom_point() +
  geom_abline(slope=1, intercept=0)+
  geom_smooth(method = "lm", se=TRUE, color="red", formula = y ~ x) +
  stat_poly_eq(aes(label = paste(..rr.label.., sep = "~~~")),
               label.x.npc = "right", label.y.npc = 0.15,
               formula = y ~ x, parse = TRUE, size = 4) +
  stat_poly_eq(aes(label = paste(..p.value.label.., sep = "~~~")),
               label.x.npc = "right", label.y.npc = "bottom",
               formula = y ~ x, parse = TRUE, size = 4) +
  annotate("text", x=10.5, y=3, label = paste("RMSE = ", rmse.GS2))+
  ggtitle("Last Glacial Termination")+
  labs(x = expression(paste(delta^{15}, "N predicted (\u2030)")),
       y = expression(paste(delta^{15}, "N observed (\u2030)"))) +
  scale_y_continuous(limits = c(0,12), breaks = seq(0, 12, by = 2)) +
  scale_x_continuous(limits = c(0,12),breaks = seq(0, 12, by = 2)) +
  theme_bw())


(LGM.pred.ob.best <- ggplot(data = d15N.LGM.aggre.df, mapping=aes(x=pred, y=mean_source_value))+
  geom_point() +
  geom_abline(slope=1, intercept=0)+
  geom_smooth(method = "lm", se=TRUE, color="red", formula = y ~ x) +
  stat_poly_eq(aes(label = paste(..rr.label.., sep = "~~~")),
               label.x.npc = "right", label.y.npc = 0.15,
```

```
               formula = y ~ x, parse = TRUE, size = 4) +
  stat_poly_eq(aes(label = paste(..p.value.label.., sep = "~~~")),
               label.x.npc = "right", label.y.npc = "bottom",
               formula = y ~ x, parse = TRUE, size = 4) +
    annotate("text", x=10.5, y=3, label = paste("RMSE = ", rmse.LGM))+
  ggtitle("Last Glacial Maximum")+
  labs(x = expression(paste(delta^{15}, "N predicted (\u2030)")),
       y = expression(paste(delta^{15}, "N observed (\u2030)"))) +
  scale_y_continuous(limits = c(0,12), breaks = seq(0, 12, by = 2)) +
  scale_x_continuous(limits = c(0,12),breaks = seq(0, 12, by = 2)) +
  theme_bw())

(LOIS3.pred.ob.best <- ggplot(data = d15N.LOIS3.aggre.df, mapping=aes(x=pred, y=mean_source_value))+
  geom_point() +
  geom_abline(slope=1, intercept=0)+
  geom_smooth(method = "lm", se=TRUE, color="red", formula = y ~ x) +
  stat_poly_eq(aes(label = paste(..rr.label.., sep = "~~~")),
               label.x.npc = "right", label.y.npc = 0.15,
               formula = y ~ x, parse = TRUE, size = 4) +
  stat_poly_eq(aes(label = paste(..p.value.label.., sep = "~~~")),
               label.x.npc = "right", label.y.npc = "bottom",
               formula = y ~ x, parse = TRUE, size = 4) +
    annotate("text", x=10.5, y=3, label = paste("RMSE = ", rmse.LOIS3))+
  ggtitle("Late OIS 3")+
  labs(x = expression(paste(delta^{15}, "N predicted (\u2030)")),
       y = expression(paste(delta^{15}, "N observed (\u2030)"))) +
  scale_y_continuous(limits = c(0,12), breaks = seq(0, 12, by = 2)) +
  scale_x_continuous(limits = c(0,12),breaks = seq(0, 12, by = 2)) +
  theme_bw())


# plot predicted versus observed values
(EOIS3.pred.ob.best <- ggplot(data = d15N.EOIS3.aggre.df, mapping=aes(x=pred, y=mean_source_value))+
  geom_point() +
  geom_abline(slope=1, intercept=0)+
  geom_smooth(method = "lm", se=TRUE, color="red", formula = y ~ x) +
  stat_poly_eq(aes(label = paste(..rr.label.., sep = "~~~")),
               label.x.npc = "right", label.y.npc = 0.15,
               formula = y ~ x, parse = TRUE, size = 4) +
  stat_poly_eq(aes(label = paste(..p.value.label.., sep = "~~~")),
               label.x.npc = "right", label.y.npc = "bottom",
               formula = y ~ x, parse = TRUE, size = 4) +
    annotate("text", x=10.5, y=3, label = paste("RMSE = ", rmse.EOIS3))+
  ggtitle("Early OIS 3")+
  labs(x = expression(paste(delta^{15}, "N predicted (\u2030)")),
       y = expression(paste(delta^{15}, "N observed (\u2030)"))) +
  scale_y_continuous(limits = c(0,12), breaks = seq(0, 12, by = 2)) +
  scale_x_continuous(limits = c(0,12),breaks = seq(0, 12, by = 2)) +
  theme_bw())

pred.ob.best.comparison <- ggpubr::ggarrange(EH.pred.ob.best,
                                        GS1.pred.ob.best,
                                        GI1.pred.ob.best,
                                        GS2.pred.ob.best,
                                        LGM.pred.ob.best,
                                        LOIS3.pred.ob.best,
                                        EOIS3.pred.ob.best,
                                        ncol=2, nrow=4)

ggsave(file="Figure_8.pdf", pred.ob.best.comparison, width = 20, height = 30, units = c("cm"), dpi = 300)

  ## SPECIES SPECIFIC ISOSCAPES ####
    # PLOT DIISTRIBUTION OF KEY SPECIES BY TIME BIN (FIGURE S6.1) ####

isoscape_data$finalagebin  <- factor(isoscape_data$finalagebin, levels = c("EH", "YD", "LGI", "LGT", "LGM", "LOIS3", "EOIS3"))

(species.plot <- isoscape_data %>%
  filter(FaunalCatCombi == "Equus" | FaunalCatCombi == "Rangifer" | FaunalCatCombi == "Cervus elaphus") %>%
  ggplot() +
  geom_point(aes(x=Longitude, y=Latitude), colour="red") +
  geom_path(data=eurmed, aes(long, lat, group=group), colour="black") +
  facet_grid(finalagebin ~  FaunalCatCombi) +
  xlim(-10,25) + ylim(35,60) +
  theme_bw())


ggsave(file="Figure_S5_1.pdf", species.plot, width = 20, height = 30, units = c("cm"), dpi = 300)
ggsave(file="Figure_S5_1.png", species.plot, width = 20, height = 30, units = c("cm"), dpi = 300)

# subset data
isoscape_data_horse <- isoscape_data %>%
  filter(FaunalCatCombi == "Equus")
isoscape_data_reindeer <- isoscape_data %>%
  filter(FaunalCatCombi == "Rangifer")
isoscape_data_reddeer <- isoscape_data %>%
  filter(FaunalCatCombi == "Cervus elaphus")

    # HORSE ####
# aggregate data at each location by time slice
d15N.data.aggre.horse <- isoscape_data_horse %>%
  dplyr::group_by(SiteName, finalagebin, add=TRUE) %>%
  dplyr::summarise(n_source_value = n(),
                   mean_source_value = mean(d15Ncoll),
                   var_source_value = var(d15Ncoll),
                   lat = mean(Latitude),
```

```
                  long = mean(Longitude)) %>%
  ungroup()

# set var_source_value to NA where it equals zero
d15N.data.aggre.horse$var_source_value[d15N.data.aggre.horse$var_source_value == 0] <- NA

# split df by age bin

GI1.horse <- d15N.data.aggre.horse %>% filter(finalagebin == "LGI")
GS2.horse <- d15N.data.aggre.horse %>% filter(finalagebin == "LGT")

# GI-1 model
# Fit the residual dispersion model
GI1.horse.dispfit <- fitme(formula = var_source_value ~ 1 + Matern(1|long + lat) + (1|SiteName),
                            family = Gamma(link = log), data = GI1.horse, fixed = list(phi = 2),
                            prior.weights = n_source_value - 1, control.dist = list(dist.method = "Earth"), method = "REML")

# predict Ïg of the expected square of the residual error in each location using the fit of the residual dispersion model:
GI1.horse$disp <- predict(GI1.horse.dispfit, newdata = GI1.horse)[,1]
# Fit mean model
GI1meanfit.horse <- fitme(formula = mean_source_value ~ 1 + Matern(1|long + lat) + (1|SiteName),
                           family = gaussian(link = identity), data = GI1.horse,
                           resid.model = list(formula= ~ 0 + offset(disp), family = Gamma(link = identity)),
                           prior.weights = n_source_value, control.dist = list(dist.method = "Earth"), method = "REML")


## GS-2 model
# Fit the residual dispersion model
GS2.horse.dispfit <- fitme(formula = var_source_value ~ 1 + Matern(1|long + lat) + (1|SiteName),
                            family = Gamma(link = log), data = GS2.horse, fixed = list(phi = 2),
                            prior.weights = n_source_value - 1, control.dist = list(dist.method = "Earth"), method = "REML")
# predict Ïg of the expected square of the residual error in each location using the fit of the residual dispersion model:
GS2.horse$disp <- predict(GS2.horse.dispfit, newdata = GS2.horse)[,1]
# Fit mean model
GS2meanfit.horse <- fitme(formula = mean_source_value ~ 1 + Matern(1|long + lat) + (1|SiteName),
                           family = gaussian(link = identity), data = GS2.horse,
                           resid.model = list(formula= ~ 0 + offset(disp), family = Gamma(link = identity)),
                           prior.weights = n_source_value, control.dist = list(dist.method = "Earth"), method = "REML")

# create blank raster to predict in to
GI1.raster.horse <- raster(ncol=400, nrow=250, xmn=-10, xmx=30, ymn=35, ymx=60)
projection(GI1.raster.horse) <- "+proj=longlat +datum=WGS84"
GI1_data_pred.horse <- as.data.frame(GI1.raster.horse , xy = TRUE)
GI1.pred_locs.horse <- cbind(GI1_data_pred.horse$x, GI1_data_pred.horse$y)
GI1.pred_pts.horse <- SpatialPoints(GI1.pred_locs.horse)
GI1_data_pred.horse$lat <- GI1_data_pred.horse$y
GI1_data_pred.horse$long <- GI1_data_pred.horse$x
GI1_data_pred.horse$SiteName <- GI1_data_pred.horse %>% group_indices(long, lat)

GS2.raster.horse <- raster(ncol=400, nrow=250, xmn=-10, xmx=30, ymn=35, ymx=60)
projection(GS2.raster.horse) <- "+proj=longlat +datum=WGS84"
GS2_data_pred.horse <- as.data.frame(GS2.raster.horse , xy = TRUE)
GS2.pred_locs.horse <- cbind(GS2_data_pred.horse$x, GS2_data_pred.horse$y)
GS2.pred_pts.horse <- SpatialPoints(GS2.pred_locs.horse)
GS2_data_pred.horse$lat <- GS2_data_pred.horse$y
GS2_data_pred.horse$long <- GS2_data_pred.horse$x
GS2_data_pred.horse$SiteName <- GS2_data_pred.horse %>% group_indices(long, lat)

## predict isoscapes ##

# GI1
# Using env_data_pred and dispfit, we predict the residual dispersion variance for each prediction location:
GI1_data_pred.horse$disp <- predict(GI1.horse.dispfit, newdata = GI1_data_pred.horse)[, 1]
# We can then use the env_data_pred and meanfit to compute the predicted isotope value over all possible observations:
GI1.horse.predict <- predict(GI1meanfit.horse, newdata = GI1_data_pred.horse,
                              variances = list(predVar = TRUE, residVar = TRUE))
# Then extract the point predictions, variance and residual variance:
GI1.horse.pred <- GI1_data_pred.horse[, c("lat", "long")]
GI1.horse.pred$pred <- GI1.horse.predict[, 1]
GI1.horse.pred$predVar <- attr(GI1.horse.predict, "predVar")
GI1.horse.pred$residVar <- attr(GI1.horse.predict, "residVar")

# GS2
# Using env_data_pred and dispfit, we predict the residual dispersion variance for each prediction location:
GS2_data_pred.horse$disp <- predict(GS2.horse.dispfit, newdata = GS2_data_pred.horse)[, 1]
# We can then use the env_data_pred and meanfit to compute the predicted isotope value over all possible observations:
GS2.horse.predict <- predict(GS2meanfit.horse, newdata = GS2_data_pred.horse,
                              variances = list(predVar = TRUE, residVar = TRUE))
# Then extract the point predictions, variance and residual variance:
GS2.horse.pred <- GS2_data_pred.horse[, c("lat", "long")]
GS2.horse.pred$pred <- GS2.horse.predict[, 1]
GS2.horse.pred$predVar <- attr(GS2.horse.predict, "predVar")
GS2.horse.pred$residVar <- attr(GS2.horse.predict, "residVar")


## PLOTS ISOSCAPES ##

# SET UP INTERPOLATION AREA ##

# # GI1
# first create a buffered convex hull around points
GI1.horse.x <- GI1.horse$long
GI1.horse.y <- GI1.horse$lat
GI1.horse.d15N.sample_xy <- cbind(GI1.horse.x,GI1.horse.y)
GI1.horse.d15N.sample_pts <- SpatialPoints(GI1.horse.d15N.sample_xy)
```

```
GI1.horse.ch <- chull(GI1.horse.d15N.sample_xy)
GI1.horse.d15N.sample_bound <- GI1.horse.d15N.sample_xy[c(GI1.horse.ch, GI1.horse.ch[1]), ]  # closed polygon
GI1.horse.outer <- SpatialPolygons(list(Polygons(list(Polygon(GI1.horse.d15N.sample_bound)), ID=1)))
GI1.horse.outer <- gBuffer(GI1.horse.outer, 100000, byid=TRUE)
# then define interpolation area
GI1.horse.plotarea <- gDifference(GI1.horse.outer, pcoastGI1.simp)
GI1.horse.interpol.area <- gDifference(GI1.horse.outer, GI1.horse.plotarea)

# GS2
# first create a buffered convex hull around points
GS2.horse.x <- GS2.horse$long
GS2.horse.y <- GS2.horse$lat
GS2.horse.d15N.sample_xy <- cbind(GS2.horse.x,GS2.horse.y)
GS2.horse.d15N.sample_pts <- SpatialPoints(GS2.horse.d15N.sample_xy)
GS2.horse.ch <- chull(GS2.horse.d15N.sample_xy)
GS2.horse.d15N.sample_bound <- GS2.horse.d15N.sample_xy[c(GS2.horse.ch, GS2.horse.ch[1]), ]  # closed polygon
GS2.horse.outer <- SpatialPolygons(list(Polygons(list(Polygon(GS2.horse.d15N.sample_bound)), ID=1)))
GS2.horse.outer <- gBuffer(GS2.horse.outer, 100000, byid=TRUE)
# then define interpolation area
GS2.horse.plotarea <- gDifference(GS2.horse.outer, pcoastGS2.simp)
GS2.horse.interpol.area <- gDifference(GS2.horse.outer, GS2.horse.plotarea)


## set up prediction surfaces ##
##
# GI1
# tranform prediction surfaces into raster
GI1.horse.pred <- GI1.horse.pred[,c('long', 'lat', 'pred', 'predVar', 'residVar')]
GI1.horse.pred.rast <- rasterFromXYZ(GI1.horse.pred)  #Convert first two columns as lon-lat and third as value
# crop prediction values to interpolation area
GI1.horse.cr <- crop(GI1.horse.pred.rast, extent(GI1.horse.interpol.area), snap="out")
GI1.horse.fr <- rasterize(GI1.horse.interpol.area, GI1.horse.cr)
GI1.horse.lr <- raster::mask(x=GI1.horse.cr, mask=GI1.horse.fr)
# then convert raster to dataframe:
GI1.horse.pred.rast.df  <- as.data.frame(GI1.horse.lr, xy=TRUE)
# remove data with NAs
GI1.horse.pred.rast.df <- na.omit(GI1.horse.pred.rast.df)

# GS2
# tranform prediction surfaces into raster
GS2.horse.pred <- GS2.horse.pred[,c('long', 'lat', 'pred', 'predVar', 'residVar')]
GS2.horse.pred.rast <- rasterFromXYZ(GS2.horse.pred)  #Convert first two columns as lon-lat and third as value
# crop prediction values to interpolation area
GS2.horse.cr <- crop(GS2.horse.pred.rast, extent(GS2.horse.interpol.area), snap="out")
GS2.horse.fr <- rasterize(GS2.horse.interpol.area, GS2.horse.cr)
GS2.horse.lr <- raster::mask(x=GS2.horse.cr, mask=GS2.horse.fr)
# then convert raster to dataframe:
GS2.horse.pred.rast.df  <- as.data.frame(GS2.horse.lr, xy=TRUE)
# remove data with NAs
GS2.horse.pred.rast.df <- na.omit(GS2.horse.pred.rast.df)

## plots ##
# set up colour schemes
col <- rev(brewer.pal(11,"Spectral"))
col.var <- c("honeydew1", "palegreen", "mediumseagreen", "seagreen4")

# GI1 prediciton surface
(GI1mappred.horse <- ggplot()+
  geom_polygon(data=pcoastGI1.simp, aes(long, lat, group=group), fill="lightgrey") +
  geom_tile(data = GI1.horse.pred.rast.df, mapping=aes(x=x, y=y, fill=pred))+
  scale_fill_stepsn(colours = col, limits = c(0,6), breaks=c(0.5,1,1.5,2,2.5,3,3.5,4,4.5,5,5.5))+
  geom_path(data=eurmed, aes(long, lat, group=group), colour="black") +
  geom_polygon(data=iceGI1, aes(long, lat, group=group), colour="black", fill="white") +
  geom_point(data = GI1.horse, aes(long, lat), colour="black", size=0.5) +
  coord_cartesian(xlim = c(-10,25), ylim = c(40, 60)) +
  xlab("Longitude") +
  ylab("Latitude") +
  labs(title="Late Glacial Interstadial: Horse",
       fill=expression(paste(delta^{15}, "N (\u2030)"))) +
  theme_bw() +
  theme(legend.position="right"))


# GI1 prediciton variance
(GI1mapvar.horse <- ggplot()+
  geom_polygon(data=pcoastGI1.simp, aes(long, lat, group=group), fill="lightgrey") +
  geom_tile(data = GI1.horse.pred.rast.df, mapping=aes(x=x, y=y, fill=predVar)) +
  scale_fill_stepsn(colours = col.var, breaks=c(0.5,1,1.5), limits = c(0, 3))+
  geom_path(data=eurmed, aes(long, lat, group=group), colour="black") +
  geom_polygon(data=iceGI1, aes(long, lat, group=group), colour="black", fill="white") +
  geom_point(data = GI1.horse, aes(long, lat), colour="black", size=0.5) +
  coord_cartesian(xlim = c(-10,25), ylim = c(40, 60)) +
  xlab("Longitude") +
  ylab("Latitude") +
  labs(title="Late Glacial Interstadial: Horse",
       fill=expression(paste(delta^{15}, "N (\u2030)"))) +
  theme_bw() +
  theme(legend.position="right"))


# GS2 prediciton surface
(GS2mappred.horse <- ggplot()+
  geom_polygon(data=pcoastGS2.simp, aes(long, lat, group=group), fill="lightgrey") +
  geom_tile(data = GS2.horse.pred.rast.df, mapping=aes(x=x, y=y, fill=pred)) +
  scale_fill_stepsn(colours = col, limits = c(0,6), breaks=c(0.5,1,1.5,2,2.5,3,3.5,4,4.5,5,5.5)) +
```

```
  geom_path(data=eurmed, aes(long, lat, group=group), colour="black") +
  geom_polygon(data=iceGS2, aes(long, lat, group=group), colour="black", fill="white") +
  geom_point(data = GS2.horse, aes(long, lat), colour="black", size=0.5) +
  coord_cartesian(xlim = c(-10,25), ylim = c(40, 60)) +
  xlab("Longitude") +
  ylab("Latitude") +
  labs(title="Last Glacial Termination: Horse",
       fill=expression(paste(delta^{15}, "N (\u2030)"))) +
  theme_bw() +
  theme(legend.position="none"))

# GS2 prediciton variance
(GS2mapvar.horse <- ggplot()+
  geom_polygon(data=pcoastGS2.simp, aes(long, lat, group=group), fill="lightgrey") +
  geom_tile(data = GS2.horse.pred.rast.df, mapping=aes(x=x, y=y, fill=predVar)) +
  scale_fill_stepsn(colours = col.var, breaks=c(0.5,1,1.5), limits = c(0, 3))+
  geom_path(data=eurmed, aes(long, lat, group=group), colour="black") +
  geom_polygon(data=iceGS2, aes(long, lat, group=group), colour="black", fill="white") +
  geom_point(data = GS2.horse, aes(long, lat), colour="black", size=0.5) +
  coord_cartesian(xlim = c(-10,25), ylim = c(40, 60)) +
  xlab("Longitude") +
  ylab("Latitude") +
  labs(title="Last Glacial Termination: Horse",
       fill=expression(paste(delta^{15}, "N (\u2030)"))) +
  theme_bw() +
  theme(legend.position="none"))


    # REINDEER ####
# aggregate data at each location by time slice
d15N.data.aggre.reindeer <- isoscape_data_reindeer %>%
  dplyr::group_by(SiteName, finalagebin, add=TRUE) %>%
  dplyr::summarise(n_source_value = n(),
                   mean_source_value = mean(d15Ncoll),
                   var_source_value = var(d15Ncoll),
                   lat = mean(Latitude),
                   long = mean(Longitude)) %>%
  ungroup()

# set var_source_value to NA where it equals zero
d15N.data.aggre.reindeer$var_source_value[d15N.data.aggre.reindeer$var_source_value == 0] <- NA

# split df by age bin

GI1.reindeer <- d15N.data.aggre.reindeer %>% filter(finalagebin == "LGI")
GS2.reindeer <- d15N.data.aggre.reindeer %>% filter(finalagebin == "LGT")

# GI-1 model
# Fit the residual dispersion model
GI1.reindeer.dispfit <- fitme(formula = var_source_value ~ 1 + Matern(1|long + lat) + (1|SiteName),
                            family = Gamma(link = log), data = GI1.reindeer, fixed = list(phi = 2),
                            prior.weights = n_source_value - 1, control.dist = list(dist.method = "Earth"), method = "REML")

# predict Ïg of the expected square of the residual error in each location using the fit of the residual dispersion model:
GI1.reindeer$disp <- predict(GI1.reindeer.dispfit, newdata = GI1.reindeer)[,1]
# Fit mean model
GI1meanfit.reindeer <- fitme(formula = mean_source_value ~ 1 + Matern(1|long + lat) + (1|SiteName),
                           family = gaussian(link = identity), data = GI1.reindeer,
                           resid.model = list(formula= ~ 0 + offset(disp), family = Gamma(link = identity)),
                           prior.weights = n_source_value, control.dist = list(dist.method = "Earth"), method = "REML")


## GS-2 model
# Fit the residual dispersion model
GS2.reindeer.dispfit <- fitme(formula = var_source_value ~ 1 + Matern(1|long + lat) + (1|SiteName),
                            family = Gamma(link = log), data = GS2.reindeer, fixed = list(phi = 2),
                            prior.weights = n_source_value - 1, control.dist = list(dist.method = "Earth"), method = "REML")
# predict Ïg of the expected square of the residual error in each location using the fit of the residual dispersion model:
GS2.reindeer$disp <- predict(GS2.reindeer.dispfit, newdata = GS2.reindeer)[,1]
# Fit mean model
GS2meanfit.reindeer <- fitme(formula = mean_source_value ~ 1 + Matern(1|long + lat) + (1|SiteName),
                           family = gaussian(link = identity), data = GS2.reindeer,
                           resid.model = list(formula= ~ 0 + offset(disp), family = Gamma(link = identity)),
                           prior.weights = n_source_value, control.dist = list(dist.method = "Earth"), method = "REML")

# create blank raster to predict in to
GI1.raster.reindeer <- raster(ncol=400, nrow=250, xmn=-10, xmx=30, ymn=35, ymx=60)
projection(GI1.raster.reindeer) <- "+proj=longlat +datum=WGS84"
GI1_data_pred.reindeer <- as.data.frame(GI1.raster.reindeer , xy = TRUE)
GI1.pred_locs.reindeer <- cbind(GI1_data_pred.reindeer$x, GI1_data_pred.reindeer$y)
GI1.pred_pts.reindeer <- SpatialPoints(GI1.pred_locs.reindeer)
GI1_data_pred.reindeer$lat <- GI1_data_pred.reindeer$y
GI1_data_pred.reindeer$long <- GI1_data_pred.reindeer$x
GI1_data_pred.reindeer$SiteName <- GI1_data_pred.reindeer %>% group_indices(long, lat)

GS2.raster.reindeer <- raster(ncol=400, nrow=250, xmn=-10, xmx=30, ymn=35, ymx=60)
projection(GS2.raster.reindeer) <- "+proj=longlat +datum=WGS84"
GS2_data_pred.reindeer <- as.data.frame(GS2.raster.reindeer , xy = TRUE)
GS2.pred_locs.reindeer <- cbind(GS2_data_pred.reindeer$x, GS2_data_pred.reindeer$y)
GS2.pred_pts.reindeer <- SpatialPoints(GS2.pred_locs.reindeer)
GS2_data_pred.reindeer$lat <- GS2_data_pred.reindeer$y
GS2_data_pred.reindeer$long <- GS2_data_pred.reindeer$x
GS2_data_pred.reindeer$SiteName <- GS2_data_pred.reindeer %>% group_indices(long, lat)

## predict isoscapes ##
```

```
# GI1
# Using env_data_pred and dispfit, we predict the residual dispersion variance for each prediction location:
GI1_data_pred.reindeer$disp <- predict(GI1.reindeer.dispfit, newdata = GI1_data_pred.reindeer)[, 1]
# We can then use the env_data_pred and meanfit to compute the predicted isotope value over all possible observations:
GI1.reindeer.predict <- predict(GI1meanfit.reindeer, newdata = GI1_data_pred.reindeer,
                             variances = list(predVar = TRUE, residVar = TRUE))
# Then extract the point predictions, variance and residual variance:
GI1.reindeer.pred <- GI1_data_pred.reindeer[, c("lat", "long")]
GI1.reindeer.pred$pred <- GI1.reindeer.predict[, 1]
GI1.reindeer.pred$predVar <- attr(GI1.reindeer.predict, "predVar")
GI1.reindeer.pred$residVar <- attr(GI1.reindeer.predict, "residVar")

# GS2
# Using env_data_pred and dispfit, we predict the residual dispersion variance for each prediction location:
GS2_data_pred.reindeer$disp <- predict(GS2.reindeer.dispfit, newdata = GS2_data_pred.reindeer)[, 1]
# We can then use the env_data_pred and meanfit to compute the predicted isotope value over all possible observations:
GS2.reindeer.predict <- predict(GS2meanfit.reindeer, newdata = GS2_data_pred.reindeer,
                             variances = list(predVar = TRUE, residVar = TRUE))
# Then extract the point predictions, variance and residual variance:
GS2.reindeer.pred <- GS2_data_pred.reindeer[, c("lat", "long")]
GS2.reindeer.pred$pred <- GS2.reindeer.predict[, 1]
GS2.reindeer.pred$predVar <- attr(GS2.reindeer.predict, "predVar")
GS2.reindeer.pred$residVar <- attr(GS2.reindeer.predict, "residVar")


## PLOTS ISOSCAPES ##

# SET UP INTERPOLATION AREA ##

# # GI1
# first create a buffered convex hull around points
GI1.reindeer.x <- GI1.reindeer$long
GI1.reindeer.y <- GI1.reindeer$lat
GI1.reindeer.d15N.sample_xy <- cbind(GI1.reindeer.x,GI1.reindeer.y)
GI1.reindeer.d15N.sample_pts <- SpatialPoints(GI1.reindeer.d15N.sample_xy)
GI1.reindeer.ch <- chull(GI1.reindeer.d15N.sample_xy)
GI1.reindeer.d15N.sample_bound <- GI1.reindeer.d15N.sample_xy[c(GI1.reindeer.ch, GI1.reindeer.ch[1]), ]  # closed polygon
GI1.reindeer.outer <- SpatialPolygons(list(Polygons(list(Polygon(GI1.reindeer.d15N.sample_bound)), ID=1)))
GI1.reindeer.outer <- gBuffer(GI1.reindeer.outer, 100000, byid=TRUE)
# then define interpolation area
GI1.reindeer.plotarea <- gDifference(GI1.reindeer.outer, pcoastGI1.simp)
GI1.reindeer.interpol.area <- gDifference(GI1.reindeer.outer, GI1.reindeer.plotarea)

# GS2
# first create a buffered convex hull around points
GS2.reindeer.x <- GS2.reindeer$long
GS2.reindeer.y <- GS2.reindeer$lat
GS2.reindeer.d15N.sample_xy <- cbind(GS2.reindeer.x,GS2.reindeer.y)
GS2.reindeer.d15N.sample_pts <- SpatialPoints(GS2.reindeer.d15N.sample_xy)
GS2.reindeer.ch <- chull(GS2.reindeer.d15N.sample_xy)
GS2.reindeer.d15N.sample_bound <- GS2.reindeer.d15N.sample_xy[c(GS2.reindeer.ch, GS2.reindeer.ch[1]), ]  # closed polygon
GS2.reindeer.outer <- SpatialPolygons(list(Polygons(list(Polygon(GS2.reindeer.d15N.sample_bound)), ID=1)))
GS2.reindeer.outer <- gBuffer(GS2.reindeer.outer, 100000, byid=TRUE)
# then define interpolation area
GS2.reindeer.plotarea <- gDifference(GS2.reindeer.outer, pcoastGS2.simp)
GS2.reindeer.interpol.area <- gDifference(GS2.reindeer.outer, GS2.reindeer.plotarea)


## set up prediction surfaces ##
##
# GI1
# tranform prediction surfaces into raster
GI1.reindeer.pred <- GI1.reindeer.pred[,c('long', 'lat', 'pred', 'predVar', 'residVar')]
GI1.reindeer.pred.rast <- rasterFromXYZ(GI1.reindeer.pred)  #Convert first two columns as lon-lat and third as value
# crop prediction values to interpolation area
GI1.reindeer.cr <- crop(GI1.reindeer.pred.rast, extent(GI1.reindeer.interpol.area), snap="out")
GI1.reindeer.fr <- rasterize(GI1.reindeer.interpol.area, GI1.reindeer.cr)
GI1.reindeer.lr <- raster::mask(x=GI1.reindeer.cr, mask=GI1.reindeer.fr)
# then convert raster to dataframe:
GI1.reindeer.pred.rast.df  <- as.data.frame(GI1.reindeer.lr, xy=TRUE)
# remove data with NAs
GI1.reindeer.pred.rast.df <- na.omit(GI1.reindeer.pred.rast.df)

# GS2
# tranform prediction surfaces into raster
GS2.reindeer.pred <- GS2.reindeer.pred[,c('long', 'lat', 'pred', 'predVar', 'residVar')]
GS2.reindeer.pred.rast <- rasterFromXYZ(GS2.reindeer.pred)  #Convert first two columns as lon-lat and third as value
# crop prediction values to interpolation area
GS2.reindeer.cr <- crop(GS2.reindeer.pred.rast, extent(GS2.reindeer.interpol.area), snap="out")
GS2.reindeer.fr <- rasterize(GS2.reindeer.interpol.area, GS2.reindeer.cr)
GS2.reindeer.lr <- raster::mask(x=GS2.reindeer.cr, mask=GS2.reindeer.fr)
# then convert raster to dataframe:
GS2.reindeer.pred.rast.df  <- as.data.frame(GS2.reindeer.lr, xy=TRUE)
# remove data with NAs
GS2.reindeer.pred.rast.df <- na.omit(GS2.reindeer.pred.rast.df)

## plots ##
# set up colour schemes
col <- rev(brewer.pal(11,"Spectral"))
col.var <- c("honeydew1", "palegreen", "mediumseagreen", "seagreen4")

# GI1 prediciton surface
(GI1mappred.reindeer <- ggplot()+
  geom_polygon(data=pcoastGI1.simp, aes(long, lat, group=group), fill="lightgrey") +
```

```
  geom_tile(data = GI1.reindeer.pred.rast.df, mapping=aes(x=x, y=y, fill=pred))+
  scale_fill_stepsn(colours = col, limits = c(0,6), breaks=c(0.5,1,1.5,2,2.5,3,3.5,4,4.5,5,5.5))+
  geom_path(data=eurmed, aes(long, lat, group=group), colour="black") +
  geom_polygon(data=iceGI1, aes(long, lat, group=group), colour="black", fill="white") +
  geom_point(data = GI1.reindeer, aes(long, lat), colour="black", size=0.5) +
  coord_cartesian(xlim = c(-10,25), ylim = c(40, 60)) +
  xlab("Longitude") +
  ylab("Latitude") +
  labs(title="Late Glacial Interstadial: Reindeer",
       fill=expression(paste(delta^{15}, "N (\u2030)"))) +
  theme_bw() +
  theme(legend.position="right"))


# GI1 prediciton variance
(GI1mapvar.reindeer <- ggplot()+
  geom_polygon(data=pcoastGI1.simp, aes(long, lat, group=group), fill="lightgrey") +
  geom_tile(data = GI1.reindeer.pred.rast.df, mapping=aes(x=x, y=y, fill=predVar)) +
  scale_fill_stepsn(colours = col.var, breaks=c(0.5,1,1.5), limits = c(0, 3))+
  geom_path(data=eurmed, aes(long, lat, group=group), colour="black") +
  geom_polygon(data=iceGI1, aes(long, lat, group=group), colour="black", fill="white") +
  geom_point(data = GI1.reindeer, aes(long, lat), colour="black", size=0.5) +
  coord_cartesian(xlim = c(-10,25), ylim = c(40, 60)) +
  xlab("Longitude") +
  ylab("Latitude") +
  labs(title="Late Glacial Interstadial: Reindeer",
       fill=expression(paste(delta^{15}, "N (\u2030)"))) +
  theme_bw() +
  theme(legend.position="right"))


# GS2 prediciton surface
(GS2mappred.reindeer <- ggplot()+
  geom_polygon(data=pcoastGS2.simp, aes(long, lat, group=group), fill="lightgrey") +
  geom_tile(data = GS2.reindeer.pred.rast.df, mapping=aes(x=x, y=y, fill=pred)) +
  scale_fill_stepsn(colours = col, limits = c(0,6), breaks=c(0.5,1,1.5,2,2.5,3,3.5,4,4.5,5,5.5)) +
  geom_path(data=eurmed, aes(long, lat, group=group), colour="black") +
  geom_polygon(data=iceGS2, aes(long, lat, group=group), colour="black", fill="white") +
  geom_point(data = GS2.reindeer, aes(long, lat), colour="black", size=0.5) +
  coord_cartesian(xlim = c(-10,25), ylim = c(40, 60)) +
  xlab("Longitude") +
  ylab("Latitude") +
  labs(title="Last Glacial Termination: Reindeer",
       fill=expression(paste(delta^{15}, "N (\u2030)"))) +
  theme_bw() +
  theme(legend.position="none"))


# GS2 prediciton variance
(GS2mapvar.reindeer <- ggplot()+
  geom_polygon(data=pcoastGS2.simp, aes(long, lat, group=group), fill="lightgrey") +
  geom_tile(data = GS2.reindeer.pred.rast.df, mapping=aes(x=x, y=y, fill=predVar)) +
  scale_fill_stepsn(colours = col.var, breaks=c(0.5,1,1.5), limits = c(0, 3))+
  geom_path(data=eurmed, aes(long, lat, group=group), colour="black") +
  geom_polygon(data=iceGS2, aes(long, lat, group=group), colour="black", fill="white") +
  geom_point(data = GS2.reindeer, aes(long, lat), colour="black", size=0.5) +
  coord_cartesian(xlim = c(-10,25), ylim = c(40, 60)) +
  xlab("Longitude") +
  ylab("Latitude") +
  labs(title="Last Glacial Termination: Reindeer",
       fill=expression(paste(delta^{15}, "N (\u2030)"))) +
  theme_bw() +
  theme(legend.position="none"))

    # RED DEER ####
# aggregate data at each location by time slice
d15N.data.aggre.reddeer <- isoscape_data_reddeer %>%
  dplyr::group_by(SiteName, finalagebin, add=TRUE) %>%
  dplyr::summarise(n_source_value = n(),
                   mean_source_value = mean(d15Ncoll),
                   var_source_value = var(d15Ncoll),
                   lat = mean(Latitude),
                   long = mean(Longitude)) %>%
  ungroup()

# set var_source_value to NA where it equals zero
d15N.data.aggre.reddeer$var_source_value[d15N.data.aggre.reddeer$var_source_value == 0] <- NA

# split df by age bin

GI1.reddeer <- d15N.data.aggre.reddeer %>% filter(finalagebin == "LGI")
GS2.reddeer <- d15N.data.aggre.reddeer %>% filter(finalagebin == "LGT")

# GI-1 model
# Fit the residual dispersion model
GI1.reddeer.dispfit <- fitme(formula = var_source_value ~ 1 + Matern(1|long + lat) + (1|SiteName),
                             family = Gamma(link = log), data = GI1.reddeer, fixed = list(phi = 2),
                             prior.weights = n_source_value - 1, control.dist = list(dist.method = "Earth"), method = "REML")

# predict Ïg of the expected square of the residual error in each location using the fit of the residual dispersion model:
GI1.reddeer$disp <- predict(GI1.reddeer.dispfit, newdata = GI1.reddeer)[,1]
# Fit mean model
GI1meanfit.reddeer <- fitme(formula = mean_source_value ~ 1 + Matern(1|long + lat) + (1|SiteName),
                            family = gaussian(link = identity), data = GI1.reddeer,
                            resid.model = list(formula= ~ 0 + offset(disp), family = Gamma(link = identity)),
```

```
                              prior.weights = n_source_value, control.dist = list(dist.method = "Earth"), method = "REML")


## GS-2 model
# Fit the residual dispersion model
GS2.reddeer.dispfit <- fitme(formula = var_source_value ~ 1 + Matern(1|long + lat) + (1|SiteName),
                               family = Gamma(link = log), data = GS2.reddeer, fixed = list(phi = 2),
                               prior.weights = n_source_value - 1, control.dist = list(dist.method = "Earth"), method = "REML")
# predict Ïg of the expected square of the residual error in each location using the fit of the residual dispersion model:
GS2.reddeer$disp <- predict(GS2.reddeer.dispfit, newdata = GS2.reddeer)[,1]
# Fit mean model
GS2meanfit.reddeer <- fitme(formula = mean_source_value ~ 1 + Matern(1|long + lat) + (1|SiteName),
                              family = gaussian(link = identity), data = GS2.reddeer,
                              resid.model = list(formula= ~ 0 + offset(disp), family = Gamma(link = identity)),
                              prior.weights = n_source_value, control.dist = list(dist.method = "Earth"), method = "REML")

# create blank raster to predict in to
GI1.raster.reddeer <- raster(ncol=400, nrow=250, xmn=-10, xmx=30, ymn=35, ymx=60)
projection(GI1.raster.reddeer) <- "+proj=longlat +datum=WGS84"
GI1_data_pred.reddeer <- as.data.frame(GI1.raster.reddeer , xy = TRUE)
GI1.pred_locs.reddeer <- cbind(GI1_data_pred.reddeer$x, GI1_data_pred.reddeer$y)
GI1.pred_pts.reddeer <- SpatialPoints(GI1.pred_locs.reddeer)
GI1_data_pred.reddeer$lat <- GI1_data_pred.reddeer$y
GI1_data_pred.reddeer$long <- GI1_data_pred.reddeer$x
GI1_data_pred.reddeer$SiteName <- GI1_data_pred.reddeer %>% group_indices(long, lat)

GS2.raster.reddeer <- raster(ncol=400, nrow=250, xmn=-10, xmx=30, ymn=35, ymx=60)
projection(GS2.raster.reddeer) <- "+proj=longlat +datum=WGS84"
GS2_data_pred.reddeer <- as.data.frame(GS2.raster.reddeer , xy = TRUE)
GS2.pred_locs.reddeer <- cbind(GS2_data_pred.reddeer$x, GS2_data_pred.reddeer$y)
GS2.pred_pts.reddeer <- SpatialPoints(GS2.pred_locs.reddeer)
GS2_data_pred.reddeer$lat <- GS2_data_pred.reddeer$y
GS2_data_pred.reddeer$long <- GS2_data_pred.reddeer$x
GS2_data_pred.reddeer$SiteName <- GS2_data_pred.reddeer %>% group_indices(long, lat)

## predict isoscapes ##

# GI1
# Using env_data_pred and dispfit, we predict the residual dispersion variance for each prediction location:
GI1_data_pred.reddeer$disp <- predict(GI1.reddeer.dispfit, newdata = GI1_data_pred.reddeer)[, 1]
# We can then use the env_data_pred and meanfit to compute the predicted isotope value over all possible observations:
GI1.reddeer.predict <- predict(GI1meanfit.reddeer, newdata = GI1_data_pred.reddeer,
                                 variances = list(predVar = TRUE, residVar = TRUE))
# Then extract the point predictions, variance and residual variance:
GI1.reddeer.pred <- GI1_data_pred.reddeer[, c("lat", "long")]
GI1.reddeer.pred$pred <- GI1.reddeer.predict[, 1]
GI1.reddeer.pred$predVar <- attr(GI1.reddeer.predict, "predVar")
GI1.reddeer.pred$residVar <- attr(GI1.reddeer.predict, "residVar")

# GS2
# Using env_data_pred and dispfit, we predict the residual dispersion variance for each prediction location:
GS2_data_pred.reddeer$disp <- predict(GS2.reddeer.dispfit, newdata = GS2_data_pred.reddeer)[, 1]
# We can then use the env_data_pred and meanfit to compute the predicted isotope value over all possible observations:
GS2.reddeer.predict <- predict(GS2meanfit.reddeer, newdata = GS2_data_pred.reddeer,
                                 variances = list(predVar = TRUE, residVar = TRUE))
# Then extract the point predictions, variance and residual variance:
GS2.reddeer.pred <- GS2_data_pred.reddeer[, c("lat", "long")]
GS2.reddeer.pred$pred <- GS2.reddeer.predict[, 1]
GS2.reddeer.pred$predVar <- attr(GS2.reddeer.predict, "predVar")
GS2.reddeer.pred$residVar <- attr(GS2.reddeer.predict, "residVar")


## PLOTS ISOSCAPES ##

# SET UP INTERPOLATION AREA ##

# # GI1
# first create a buffered convex hull around points
GI1.reddeer.x <- GI1.reddeer$long
GI1.reddeer.y <- GI1.reddeer$lat
GI1.reddeer.d15N.sample_xy <- cbind(GI1.reddeer.x,GI1.reddeer.y)
GI1.reddeer.d15N.sample_pts <- SpatialPoints(GI1.reddeer.d15N.sample_xy)
GI1.reddeer.ch <- chull(GI1.reddeer.d15N.sample_xy)
GI1.reddeer.d15N.sample_bound <- GI1.reddeer.d15N.sample_xy[c(GI1.reddeer.ch, GI1.reddeer.ch[1]), ]  # closed polygon
GI1.reddeer.outer <- SpatialPolygons(list(Polygons(list(Polygon(GI1.reddeer.d15N.sample_bound)), ID=1)))
GI1.reddeer.outer <- gBuffer(GI1.reddeer.outer, 100000, byid=TRUE)
# then define interpolation area
GI1.reddeer.plotarea <- gDifference(GI1.reddeer.outer, pcoastGI1.simp)
GI1.reddeer.interpol.area <- gDifference(GI1.reddeer.outer, GI1.reddeer.plotarea)

# GS2
# first create a buffered convex hull around points
GS2.reddeer.x <- GS2.reddeer$long
GS2.reddeer.y <- GS2.reddeer$lat
GS2.reddeer.d15N.sample_xy <- cbind(GS2.reddeer.x,GS2.reddeer.y)
GS2.reddeer.d15N.sample_pts <- SpatialPoints(GS2.reddeer.d15N.sample_xy)
GS2.reddeer.ch <- chull(GS2.reddeer.d15N.sample_xy)
GS2.reddeer.d15N.sample_bound <- GS2.reddeer.d15N.sample_xy[c(GS2.reddeer.ch, GS2.reddeer.ch[1]), ]  # closed polygon
GS2.reddeer.outer <- SpatialPolygons(list(Polygons(list(Polygon(GS2.reddeer.d15N.sample_bound)), ID=1)))
GS2.reddeer.outer <- gBuffer(GS2.reddeer.outer, 100000, byid=TRUE)
# then define interpolation area
GS2.reddeer.plotarea <- gDifference(GS2.reddeer.outer, pcoastGS2.simp)
GS2.reddeer.interpol.area <- gDifference(GS2.reddeer.outer, GS2.reddeer.plotarea)
```

```
## set up prediction surfaces ##
##
# GI1
# tranform prediction surfaces into raster
GI1.reddeer.pred <- GI1.reddeer.pred[,c('long', 'lat', 'pred', 'predVar', 'residVar')]
GI1.reddeer.pred.rast <- rasterFromXYZ(GI1.reddeer.pred)  #Convert first two columns as lon-lat and third as value
# crop prediction values to interpolation area
GI1.reddeer.cr <- crop(GI1.reddeer.pred.rast, extent(GI1.reddeer.interpol.area), snap="out")
GI1.reddeer.fr <- rasterize(GI1.reddeer.interpol.area, GI1.reddeer.cr)
GI1.reddeer.lr <- raster::mask(x=GI1.reddeer.cr, mask=GI1.reddeer.fr)
# then convert raster to dataframe:
GI1.reddeer.pred.rast.df  <- as.data.frame(GI1.reddeer.lr, xy=TRUE)
# remove data with NAs
GI1.reddeer.pred.rast.df <- na.omit(GI1.reddeer.pred.rast.df)

# GS2
# tranform prediction surfaces into raster
GS2.reddeer.pred <- GS2.reddeer.pred[,c('long', 'lat', 'pred', 'predVar', 'residVar')]
GS2.reddeer.pred.rast <- rasterFromXYZ(GS2.reddeer.pred)  #Convert first two columns as lon-lat and third as value
# crop prediction values to interpolation area
GS2.reddeer.cr <- crop(GS2.reddeer.pred.rast, extent(GS2.reddeer.interpol.area), snap="out")
GS2.reddeer.fr <- rasterize(GS2.reddeer.interpol.area, GS2.reddeer.cr)
GS2.reddeer.lr <- raster::mask(x=GS2.reddeer.cr, mask=GS2.reddeer.fr)
# then convert raster to dataframe:
GS2.reddeer.pred.rast.df  <- as.data.frame(GS2.reddeer.lr, xy=TRUE)
# remove data with NAs
GS2.reddeer.pred.rast.df <- na.omit(GS2.reddeer.pred.rast.df)

## plots ##
# set up colour schemes
col <- rev(brewer.pal(11,"Spectral"))
col.var <- c("honeydew1", "palegreen", "mediumseagreen", "seagreen4")

# GI1 prediciton surface
(GI1mappred.reddeer <- ggplot()+
  geom_polygon(data=pcoastGI1.simp, aes(long, lat, group=group), fill="lightgrey") +
  geom_tile(data = GI1.reddeer.pred.rast.df, mapping=aes(x=x, y=y, fill=pred))+
  scale_fill_stepsn(colours = col, limits = c(0,6), breaks=c(0.5,1,1.5,2,2.5,3,3.5,4,4.5,5,5.5))+
  geom_path(data=eurmed, aes(long, lat, group=group), colour="black") +
  geom_polygon(data=iceGI1, aes(long, lat, group=group), colour="black", fill="white") +
  geom_point(data = GI1.reddeer, aes(long, lat), colour="black", size=0.5) +
  coord_cartesian(xlim = c(-10,25), ylim = c(40, 60)) +
  xlab("Longitude") +
  ylab("Latitude") +
  labs(title="Late Glacial Interstadial: Red Deer",
       fill=expression(paste(delta^{15}, "N (\u2030)"))) +
  theme_bw() +
  theme(legend.position="right"))


# GI1 prediciton variance
(GI1mapvar.reddeer <- ggplot()+
  geom_polygon(data=pcoastGI1.simp, aes(long, lat, group=group), fill="lightgrey") +
  geom_tile(data = GI1.reddeer.pred.rast.df, mapping=aes(x=x, y=y, fill=predVar)) +
  scale_fill_stepsn(colours = col.var, breaks=c(0.5,1,1.5), limits = c(0, 3))+
  geom_path(data=eurmed, aes(long, lat, group=group), colour="black") +
  geom_polygon(data=iceGI1, aes(long, lat, group=group), colour="black", fill="white") +
  geom_point(data = GI1.reddeer, aes(long, lat), colour="black", size=0.5) +
  coord_cartesian(xlim = c(-10,25), ylim = c(40, 60)) +
  xlab("Longitude") +
  ylab("Latitude") +
  labs(title="Late Glacial Interstadial: Red Deer",
       fill=expression(paste(delta^{15}, "N (\u2030)"))) +
  theme_bw() +
  theme(legend.position="right"))

# GS2 prediciton surface
(GS2mappred.reddeer <- ggplot()+
  geom_polygon(data=pcoastGS2.simp, aes(long, lat, group=group), fill="lightgrey") +
  geom_tile(data = GS2.reddeer.pred.rast.df, mapping=aes(x=x, y=y, fill=pred)) +
  scale_fill_stepsn(colours = col, limits = c(0,6), breaks=c(0.5,1,1.5,2,2.5,3,3.5,4,4.5,5,5.5)) +
  geom_path(data=eurmed, aes(long, lat, group=group), colour="black") +
  geom_polygon(data=iceGS2, aes(long, lat, group=group), colour="black", fill="white") +
  geom_point(data = GS2.reddeer, aes(long, lat), colour="black", size=0.5) +
  coord_cartesian(xlim = c(-10,25), ylim = c(40, 60)) +
  xlab("Longitude") +
  ylab("Latitude") +
  labs(title="Last Glacial Termination: Red Deer",
       fill=expression(paste(delta^{15}, "N (\u2030)"))) +
  theme_bw() +
  theme(legend.position="none"))

# GS2 prediciton variance
(GS2mapvar.reddeer <- ggplot()+
  geom_polygon(data=pcoastGS2.simp, aes(long, lat, group=group), fill="lightgrey") +
  geom_tile(data = GS2.reddeer.pred.rast.df, mapping=aes(x=x, y=y, fill=predVar)) +
  scale_fill_stepsn(colours = col.var, breaks=c(0.5,1,1.5), limits = c(0, 3))+
  geom_path(data=eurmed, aes(long, lat, group=group), colour="black") +
  geom_polygon(data=iceGS2, aes(long, lat, group=group), colour="black", fill="white") +
  geom_point(data = GS2.reddeer, aes(long, lat), colour="black", size=0.5) +
  coord_cartesian(xlim = c(-10,25), ylim = c(40, 60)) +
  xlab("Longitude") +
  ylab("Latitude") +
  labs(title="Last Glacial Termination: Red Deer",
       fill=expression(paste(delta^{15}, "N (\u2030)"))) +
```

```
  theme_bw() +
  theme(legend.position="none"))


## COMBINE PLOTS (FIGURES 9 AND 10) ####
species.comp.isoscapes <- ggarrange(GI1mappred.horse, GI1mappred.reindeer, GI1mappred.reddeer, GS2mappred.horse,
GS2mappred.reindeer, GS2mappred.reddeer, ncol=3, nrow=2, common.legend = TRUE, legend="bottom")
ggsave(file="Fig_10.pdf", species.comp.isoscapes, width = 40, height = 25, units = c("cm"), dpi = 300)

species.comp.var.isoscapes <- ggarrange(GI1mapvar.horse, GI1mapvar.reindeer, GI1mapvar.reddeer, GS2mapvar.horse, GS2mapvar.reindeer,
GS2mapvar.reddeer, ncol=3, nrow=2, common.legend = TRUE, legend="bottom")
ggsave(file="Fig_S5_3.pdf", species.comp.var.isoscapes, width = 40, height = 25, units = c("cm"), dpi = 300)
```
